# Lithocholic acid targets TULP3 to activate sirtuins and AMPK to retard ageing

**DOI:** 10.1101/2023.12.07.570558

**Authors:** Qi Qu, Yan Chen, Yu Wang, Shating Long, Weiche Wang, Heng-Ye Yang, Jianfeng Wu, Mengqi Li, Xiao Tian, Xiaoyan Wei, Yan-Hui Liu, Shengrong Xu, Chunyan Yang, Zhenhua Wu, Xi Huang, Changchuan Xie, Yaying Wu, Zheni Xu, Cixiong Zhang, Baoding Zhang, Jin-Wei Feng, Junjie Chen, Liyun Lin, ZK Xie, Beibei Sun, Yong Yu, Hai-Long Piao, Xiao-Song Xie, Xianming Deng, Chen-Song Zhang, Sheng-Cai Lin

**Affiliations:** State Key Laboratory for Cellular Stress Biology, School of Life Sciences, Xiamen University, Fujian, China; Laboratory Animal Research Centre, Xiamen University, Fujian, China; Analysis and Measurement Centre, School of Pharmaceutical Sciences, Xiamen University, Fujian, China; CAS Key Laboratory of Separation Science for Analytical Chemistry, Dalian Institute of Chemical Physics, Chinese Academy of Sciences, Liaoning, China; McDermott Center of Human Growth and Development MC8591, University of Texas Southwestern Medical Center, TX, USA

**Author notes:** These authors contributed equally: Qi Qu, Yan Chen, Yu Wang, Shating Long, Weiche Wang.

## Abstract

Lithocholic acid (LCA), accumulated in the body during calorie restriction (CR), can confer administered metazoans with the ability to activate AMP-activated protein kinase (AMPK) and retard ageing. However, how LCA is signalled to activate AMPK and elicit the biological effects is unclear. Here, we show that LCA can enhance sirtuins (SIRTs) to deacetylate and subsequently inhibit vacuolar H^+^-ATPase (v-ATPase), thereby triggering AMPK activation via the lysosomal glucose-sensing pathway. Through proteomic analysis of SIRT1-coimmunoprecipitated proteins, we identify and validate that TUB like protein 3 (TULP3) is a constitutive component of SIRTs. Surprisingly, we found that TULP3 is an LCA receptor, and that the LCA-bound TULP3 activates SIRTs. The activated SIRTs in turn deacetylate the V1E1 subunit of v-ATPase on K52, K99 and K191 residues. Muscle-specific expression of the 3KR mutant of V1E1, mimicking the deacetylated state, dominantly activates AMPK and rejuvenates muscles in aged mice. Moreover, LCA once administered also activates AMPK and extends lifespan and healthspan in nematodes and flies, depending on the TULP3 homologues *tub-1* and *ktub*, respectively. Our study thus elucidates that LCA triggers the TULP3-sirtuin-v-ATPase- AMPK route to manifest benefits of calorie restriction.

---

Calorie restriction (CR), a dieting habit for reducing calorie intake, has been shown to extend lifespan and healthspan in diverse species, such as worms, fruit flies, and primates^1^. CR can also elicit a myriad of molecular, cellular and physiological changes to maintain mitochondrial quality and quantity, reduce oxidative damage, and suppress inflammation, which are commonly dysregulated in aged animals^2–4^. CR can thereby help slow down age [associated metabolic disorders and diseases including central obesity, insulin resistance, muscle degeneration, and cancer^5^. AMPK, a master regulator for maintaining metabolic homeostasis, plays an essential role in mediating these benefits of CR^6^. Our preceding paper has identified that the single metabolite LCA accumulated after CR can activate AMPK in mice, and nematodes and flies as well. Importantly, AMPK is required for LCA to arouse various benefits including rejuvenation of muscle in aged mice, and extends lifespan as well as healthspan in nematodes and flies. However, how AMPK is activated by LCA to manifest benefits remains unknown.

AMPK can be activated in multiple ways, depending on the severities of cellular nutrient and energy stresses or Ca^2+^ levels^7^. Once cellular energy levels are low, such as during ischaemia^8^, the increased AMP and ADP (higher AMP:ATP and ADP:ATP ratios) lead to canonical, allosteric activation in AMPK^9–12^. Upon global increase in intracellular Ca^2+^ levels, for instance, under VEGF treatment^13^, AMPK is activated in a calcium/calmodulin-dependent protein kinase kinase 2 (CaMKK2)-dependent manner^14–16^. Under falling levels of glucose and hence lower glycolytic intermediate fructose-1,6-bisphosphate (FBP)^17^, AMPK is activated at the lysosome via the glucose sensing pathway independently of increase of AMP levels^18^. As shown in the preceding paper, we found that LCA activates AMPK without elevating AMP:ATP and ADP:ATP ratios or causing global Ca^2+^ increase, indicating that LCA activates AMPK in an AMP/ADP- or CaMKK2-independent manner. This leaves the glucose sensing pathway, or the lysosomal AMPK pathway, as a possible route for LCA to activate AMPK. In this pathway, lack of FBP is directly sensed by aldolase associated with the lysosomal proton pump v-ATPase, which in turn blocks the endoplasmic reticulum (ER)-localised calcium channel transient receptor potential V (TRPVs), converting low cellular glucose/FBP to low local Ca^2+^ signal at the ER-lysosome contact site^17,19^. TRPV then interacts with v-ATPase, causing a re-configuration of the aldolase-v-ATPase complex, resulting in inhibition of v-ATPase^19^. As a result, AXIN utilises v-ATPase and its associated Ragulator (comprised of 5 LAMTOR subunits, LAMTOR1-5) as docking sites to tether liver kinase B1 (LKB1), an AMPK upstream kinase^18,20^. This leads to phosphorylation of threonine-172 (Thr172) on the α subunit of AMPK and its activation. In addition to the aldolase-TRPV axis that senses low glucose, it was shown that metformin at clinically relevant doses binds to PEN2, which in turn binds and inhibits v-ATPase to trigger the lysosomal pathway for AMPK activation and lifespan extension^21–23^.

We therefore tested whether LCA activates AMPK via the lysosomal pathway. We found that as low as 1 μM LCA, similar to that detected in the serum of CR mice, induced AXIN in complex with LKB1 to translocate to the lysosome in MEFs, as evidenced by immunofluorescent co-staining with the lysosomal marker LAMP2 (Fig. 1a). As a result, the complex formation between AXIN/LKB1 with the v-ATPase-Ragulator complex was increased by LCA (Fig. 1b). We also showed that LCA inhibited the activity of v-ATPase in mouse embryonic fibroblasts (MEFs), as determined by the signal from LysoSensor Green DND-189 dye which decreases with increased lysosomal pH^24^ (Fig. 1c). Consistently, knocking down the *v0c* subunit (*ATP6v0c*), which renders v-ATPase devoid of the docking site for AXIN and LKB1 (ref. ^18^), blocked the AMPK activation by LCA, as assessed by the levels of the phosphorylation of AMPK α (p-AMPK α) and its substrate ACC1/2 (p-ACC1/2) (Fig. 1d). Knockout of *AXIN* (AXIN1, the only AXIN protein expressed in MEFs^25^) or *LAMTOR1* in MEFs also blocked the LCA-mediated AMPK activation (Fig. 1e, f and Extended Data Fig. 1a-c). We also delineated the requirement of the components of the lysosomal AMPK pathway in mice by treating with LCA (coated with 2-hydroxypropyl)- β-cyclodextrin) at 1 g/l in drinking water, which led to an accumulation of approximately 1.1 μM LCA in the serum (Fig. 2a of the preceding paper), and found that knockdown of *LAMTOR1* or *AXIN* (*AXIN1* in the liver, or both *AXIN1* and *AXIN2* in muscles as both AXIN1 and AXIN2 are expressed in the muscle that redundantly regulate AMPK^25–27^) abolished LCA-mediated AMPK activation in the mouse tissues (Fig. 1g, h and Extended Data Fig. 1d, e). The results above indicate that LCA inhibits v-ATPase to trigger the lysosomal pathway for AMPK activation.

**Fig. 1.**
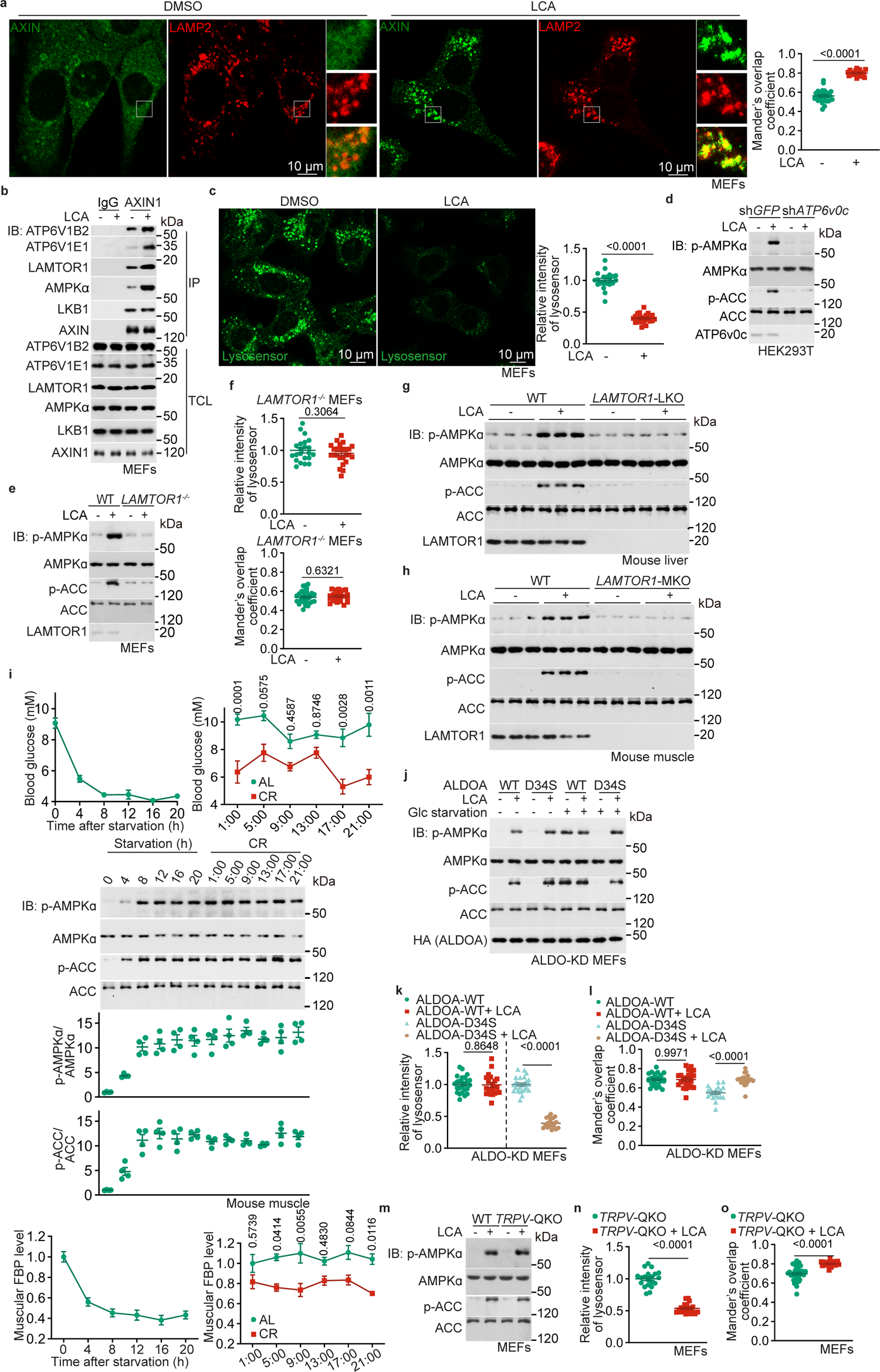
LCA activates AMPK through the lysosomal pathway. **a**, LCA triggers lysosomal translocation of AXIN. MEFs were incubated with 1 μM LCA for 4 h, followed by immunostaining to determine the lysosomal translocation of AXIN (accessed by the co-localisation, i.e., the Mander’s overlap coefficients, between AXIN and the lysosomal marker LAMP2). Results are shown as mean ± s.e.m.; *n* = 29 (DMSO) or 20 (LCA) cells, and *P* value by two-sided Student’s *t*-test with Welch’s correction. **b**, LCA induces the formation of the lysosomal AMPK-activating complex. MEFs were treated as **a**, and the cell lysates were immunoprecipitated (IP) with an antibody against AXIN. Immunoblotting (IB) reveals that AXIN forms a complex with v-ATPase, Ragulator, AXIN and LKB1. TCL, total cell lysate. **c**, LCA inhibits v-ATPase. MEFs were incubated with 1 μM LCA for 4 h, followed by determination of the activity of v-ATPase (accessed by the decreased intensities of the lysosensor to monitor the deacidification of lysosomes; representative images are shown on the left panel, and the statistical analysis data on the right as mean ± s.e.m., normalised to the DMSO group; *n* = 21 (DMSO) or 23 (LCA) cells, and *P* value by two-sided Mann-Whitney test). **d-h**, The v-ATPase and LAMTORs are required for AXIN lysosomal translocation in response to LCA. HEK293T cells with *ATP6v0c* (*v0c*) knockdown (sh*ATP6v0c*; **d**), MEFs with *LAMTOR1* knockout (*LAMTOR1*^-/-^; **e**, **f**), or mice with liver- or muscle-specific *LAMTOR1* knockout (*LAMTOR1*-LKO, in **g**; or *LAMTOR1*-MKO, in **h**), were treated with LCA, either at 1 μM for 4 h (**d-f**) or at 1 g/l after coating with (2-hydroxypropyl)- β-cyclodextrin and supplied in the drinking water for 1 month (**g**, **h**), followed by determination for AMPK activation (**d**, **e**, **g**, **h**; the levels of p-AMPK α and p-ACC) by immunoblotting, the v-ATPase inhibition (**f**, upper panel; see Extended Data Fig. 1b for the representative images), and the lysosomal translocation of AXIN evidenced by colocalisation with LAMP2 (**f**, lower panel; see Extended Data Fig. 1c for representative images). Results in **f** are shown as mean ± s.e.m., with data in upper panel normalised to the DMSO group; *n* = 22 (DMSO group of upper panel of **f**), 25 (LCA group of upper panel of **f**), 29 (DMSO group of lower panel of **f**) or 23 (LCA group of lower panel of **f**) cells, and *P* value by two-sided Student’s *t*-test. **i**, The glucose decline in CR mice does not suffice to cause AMPK activation. Levels of blood glucose (upper panel), muscle FBP (lower panel), and the activity of muscle AMPK (middle panel) were determined in mice subjected to starvation or CR (at different times of the day). Results are shown as mean ± s.e.m.; *n* = 5 (blood glucose of CR and starvation groups) or 4 (others) mice for each time point, and *P* value by two-way ANOVA followed by Tukey’s test. **j-o**, LCA triggers the lysosomal AMPK pathway downstream of the glucose sensor aldolase and TRPVs. MEFs with knockdown of aldolase A-C (ALDO-KD) and re-introduced with ALDOA-D34S that mimics high glucose/FBP condition (**j-l**; as described and validated previously^17^), or MEFs with TRPV1-4 knockout (**m-o**; TRPV-QKO; generated as described previously^19^), were incubated in normal (**j**, **m-o**) or glucose-free (**j-l**) medium (Glc starvation) containing 1 μM LCA for 4 h, followed by determination for the AMPK activation (**j**, **m**), the v-ATPase inhibition (**k**, **n**; representative images are shown in Extended Data Fig. 1g, i), and the lysosomal translocation of AXIN (**l**, **o**; representative images are shown in Extended Data Fig. 1h, j). Results are shown as mean ± s.e.m., where **k** and **n** were normalised to the DMSO group; *n* = 25 (**k**, ALDOA-WT + DMSO, ALDOA-D34S + DMSO), 24 (**l**, ALDOA-WT + DMSO), 23 (**l**, ALDOA-WT + LCA), 21 (**n** and **o**, LCA), 33 (**o**, DMSO) or 20 (others) cells; and *P* value by two-sided Student’s *t*-test (**k**, ALDOA-WT), two-sided Student’s *t*-test with Welch’s correction (**k**, ALDOA-D34S; and **n**), or two-way ANOVA followed by Tukey’s test (**l**). Experiments in this figure were performed three times, except **a**, **c** four times.

**Fig. 2.**
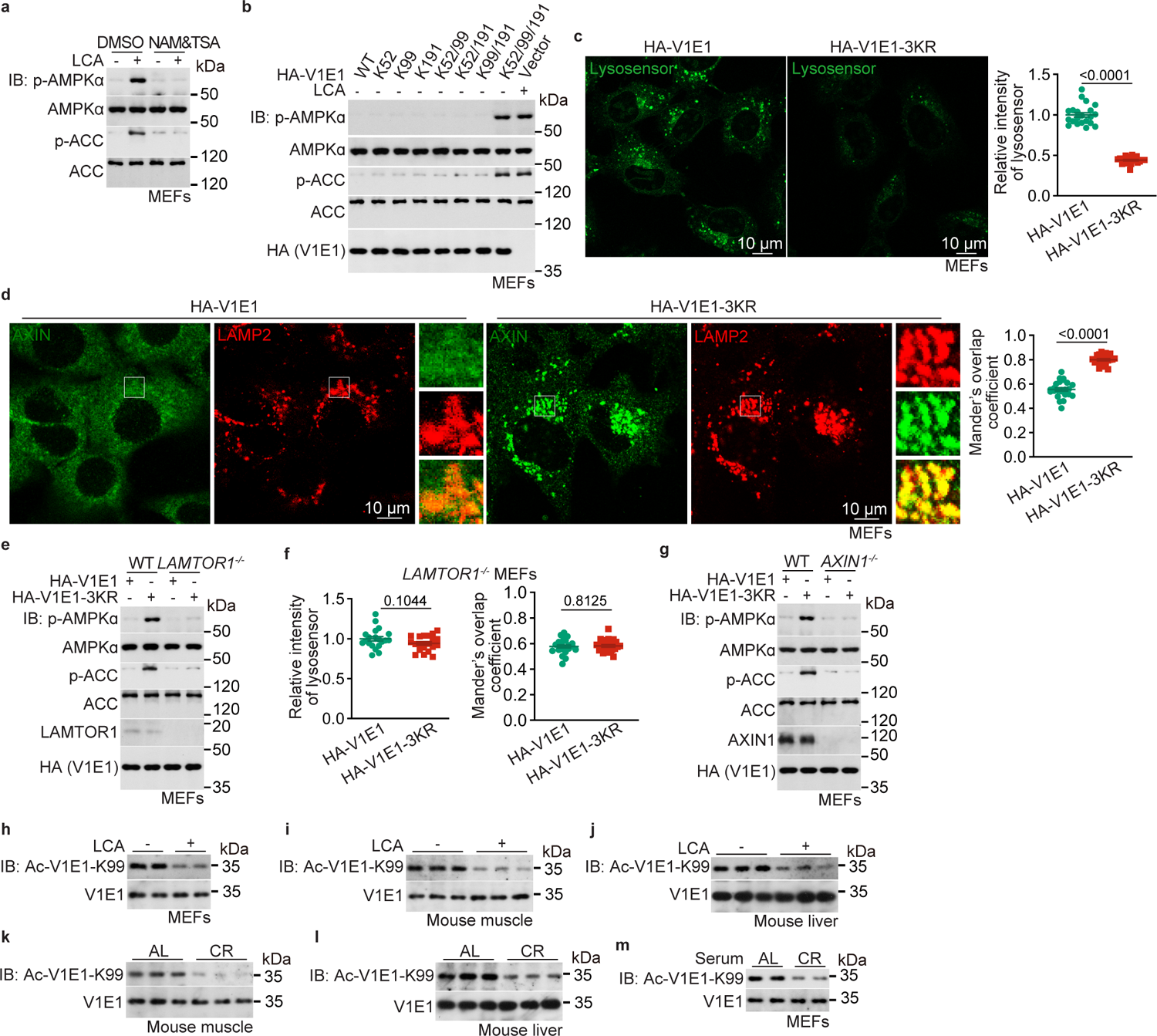
LCA causes deacetylation of v-ATPase to enter the lysosomal AMPK pathway. **a**, Deacetylation is required for LCA-induced activation of AMPK. MEFs were pre-treated with 2 μM TSA and 8 mM NAM for 12 h, followed by incubating with 1 μM LCA for 4 h. Cells were then lysed, and the AMPK activities were determined by immunoblotting. **b-d**, LCA leads to deacetylation of V1E1 at K52, K99 and K191 residues. MEFs were infected with lentivirus carrying the HA-tagged, triple mutation of K52, K99 and K191 of V1E1 to arginine (V1E1-3KR; mimicking the deacetylated state of V1E1), followed by determining the activation of AMPK (**b**), inhibition of v-ATPase (**c**; representative images are shown on the left panel), and the lysosomal translocation of AXIN (**d**; representative images are shown on the left panel). Statistical analysis data in **c** and **d** are shown as mean ± s.e.m. on the right panels, with data in **c** normalised to the V1E1 group; *n* = 21 (**c**), 24 (**d**, V1E1) or 23 (**d**, V1E1-3KR) cells, and *P* value by two-sided Mann-Whitney test (**c**) or two-sided Student’s *t*-test with Welch’s correction (**d**). **e-g**, V1E1-3KR activates AMPK. *LAMTOR1*^-/-^ (**e**, **f**) and *AXIN1*^-/-^ (**g**; AXIN knockout) MEFs, and their wildtype control MEFs, were infected with lentivirus carrying HA-tagged V1E1-3KR, followed by determination for AMPK activation (**e**, **g**), the activity of v-ATPase (**f**, left panel; representative images are shown in the upper panel of Extended Data Fig. 3b), and the lysosomal translocation of AXIN (**f**, right panel; representative images are shown on the lower panel of Extended Data Fig. 3b). Statistical analysis data in **f** are shown as mean ± s.e.m.; *n* = 20 (left panel), 22 (V1E1 of right panel) or 23 (V1E1-3KR of right panel) cells, and *P* value by two-sided Student’s *t*-test. **h-m**, CR or LCA treatment causes the deacetylation of V1E1 in vivo. MEFs treated with 1 μM LCA for 4 h (**h**), cultured in DMEM with FBS supplemented in the medium replaced with an equal volume of serum from 4-month CR mice (**m**), or mice subjected to CR for 4 months (**k**, **l**) and fed with at 1 g/l (2-hydroxypropyl)- β-cyclodextrin-coated LCA in drinking water, for 1 month (**i**, **j**) were lysed or sacrificed, followed by determining the levels of acetylated K99 of V1E1 (Ac-V1E1-K99) in cells (**h**, **m**), muscles (**i**, **k**) and livers (**j**, **l**). Experiments in this figure were performed three times, except **b** five times.

We next explored how LCA inhibits v-ATPase. As AMPK activation displays an linear regression relationship with glucose and hence FBP levels (ref. ^17^), and that both LCA and CR can lower blood glucose to some extent (Fig. 1i, and Fig. 2n of the preceding paper), we determined whether the reduction of glucose/FBP levels observed during CR or under LCA treatment could be a cause of v-ATPase inhibition. We observed that blood glucose levels of ad libitum fed mice dropped from approximately postprandial 9 mM to 5.5 mM after 4 h of fasting, which further decreased to 4.5 mM after 8 h (Fig. 1i). For AMPK activation, we observed a slight increase in muscular p-AMPK α and p-ACC at 4 h of fasting, which increased and plateaued after 8 h in these mice (Fig. 1i). In comparison, in mice after 4 months of CR treatment blood glucose levels never decreased to below 5.5 mM throughout the day, although the activation of AMPK in the CR mice is constant at a level comparable to that in ad libitum fed mice at 8-h fasting (Fig. 1i and Extended Data Fig. 1f). Similarly, levels of FBP were greatly reduced during fasting state of the ad libitum fed mice, but the change was small in CR mice (Fig. 1i). These data suggest that the decline in glucose/FBP levels alone is not the primary factor responsible for triggering AMPK activation in calorie-restricted mice. In addition, we found that LCA could still lead to the activation of AMPK in MEFs expressing the D34S mutant of aldolase A (ALDOA), a mutant that is constitutively bound with FBP and hence blocks AMPK activation even in low glucose^17,28^ (Fig. 1j-l and Extended Data Fig. 1g, h), indicating that LCA activates AMPK via a different entry, downstream of aldolase, into the lysosomal AMPK activation route. Similarly, knockout of TRPV1-4 in MEFs (MEFs have little expression of other TRPVs, as validated previously^19^) that are required for the signalling of low glucose/FBP to AMPK activation^19^, did not affect the LCA-mediated AMPK activation (Fig. 1m-o and Extended Data Fig. 1i, j). We also ruled out the PEN2-ATP6AP1 shunt used by metformin to activate the lysosomal AMPK, as knockout of *PEN2* or by re-introduction of ATP6AP1^Δ420–440^ mutants that cannot bind PEN2 into *ATP6AP1*^-/-^ MEFs^23^, did not affect the inhibition of v-ATPase and the activation of AMPK by LCA (Extended Data Fig. 2a-f).

The above data indicate that LCA acts downstream of aldolase and TRPV to enter lysosomal AMPK pathway. We wondered whether a post-translational modification(s) of v-ATPase itself might be the initiating point in LCA-mediated AMPK activation. To this end, we determined the modifications of v-ATPase by immunoprecipitating all of the HA-tagged 20 subunits of v-ATPase from HEK293T cells, followed by mass spectrometry (MS) analysis. We found that among the posttranslational modifications, acetylation is most prominent, with a total of 263 residues identified (Supplementary Table 1, and summarised in Extended Data Table 1). Importantly, the addition of nicotinamide (NAM) and Trichostatin A (TSA) that inhibit deacetylases^29,30^ prevented the LCA-mediated AMPK activation (Fig. 2a), pointing to deacetylation as a possible mechanism for LCA. We then determined acetylation and deacetylation of which residue(s) is involved in the modulation of AMPK activation by mutating all the 263 acetylated lysine residues across the 21 subunits of v-ATPase to arginine (mimicking deacetylated state) in different combinations. As summarised in Extended Data Table 1, we found that single mutations to arginine of K52, K99 or K191 of the V1E1 (ATP6V1E1), and K60 or K118 of V1E2 (ATP6V1E2) subunits of v-ATPase led to constitutive activation of AMPK. As V1E2 is specifically expressed in spermatocytes, spermatids and mature sperm of the testis^31,32^, we focused on the role of V1E1 deacetylation in AMPK activation hereafter. Triple mutation of K52, K99 and K191 to arginine (V1E1-3KR) led to a pronounced activation of AMPK, similar to the extent by LCA treatment (Fig. 2b-d and Extended Data Fig. 3a). We also found that AMPK activation by V1E1-3KR in MEFs was abrogated by knockout of *AXIN* or *LAMTOR1* (Fig. 2e-g and Extended Data Fig. 3b), connecting to the lysosomal AMPK pathway. We next made attempts to generate the site-specific antibodies against each of the three acetylated residues of V1E1, and successfully obtained an antibody against acetylated K99 (Ac-V1E1-K99; Extended Data Fig. 3c). Using this antibody, we observed that K99 is deacetylated in MEFs, mouse muscles and livers, after LCA administration (Fig. 2h-j). Similarly, deacetylation of K99 could be observed in muscle and liver tissues from CR mice (Fig. 2k, l) or in MEFs treated with the serum of CR mice (Fig. 2m). Data above demonstrate that it is the deacetylation of V1E1 that triggers the lysosomal pathway for AMPK activation by LCA.

We next explored the mechanism by which LCA leads to V1E1 deacetylation. Ectopic expression of any member of the sirtuin deacetylase family (SIRT1 to SIRT7), but not their dominant-negative mutants sufficiently led to the deacetylation of V1E1 as determined by anti-Ac-V1E1-K99 antibody in HEK293T cells (Fig. 3a). In comparison, none of the HDACs (histone deacetylases; HDAC1 to HDAC11) could deacetylate K99 (Extended Data Fig. 4a). These results indicate that sirtuins but not the HDACs participate in the deacetylation of V1E1. Consistently, ectopic expression of individual sirtuins also led to the inhibition of v-ATPase and activation of AMPK (Fig. 3a, b and Extended Data Fig. 4b). In addition, depletion of sirtuins by hepta-knockout of SIRT1 to SIRT7 (*SIRT1*-*7*^-/-^; validated in Extended Data Fig. 4c) in MEFs blocked the LCA-induced V1E1 deacetylation, v-ATPase inhibition, as well as AMPK activation (Fig. 3c-e and Extended Data Fig. 4d, e). Since *SIRT1*-*7*^-/-^ MEFs grew much slower than wildtype MEFs (Extended Data Fig. 4f), possibly owing to the requirement of sirtuin activity for the control of transcriptional processes during cell cycle and cell proliferation^33,34^, we generated SIRT2 to SIRT7 hexa-knockout MEFs (*SIRT2*-*7*^-/-^; validated in Extended Data Fig. 4g) and added SIRT1 inhibitor EX-527 (ref. ^35^) to the *SIRT2*-*7*^-/-^ MEFs, and found again that LCA failed to activate AMPK in these cells (Fig. 3f-h, Extended Data Fig. 4h, i). Re-introduction of any of the sirtuins into *SIRT1*-*7*^-/-^ MEFs partially restored the activation of AMPK by LCA, with SIRT1 being the most potent (Fig. 3i). The LCA-enhanced activity of sirtuins was also seen with other substrates, such as acetylated histone H3 (on K9 residue, which can be deacetylated by SIRT1, SIRT2 and SIRT6, as described previously^36–38^) (Fig. 3j).

**Fig. 3.**
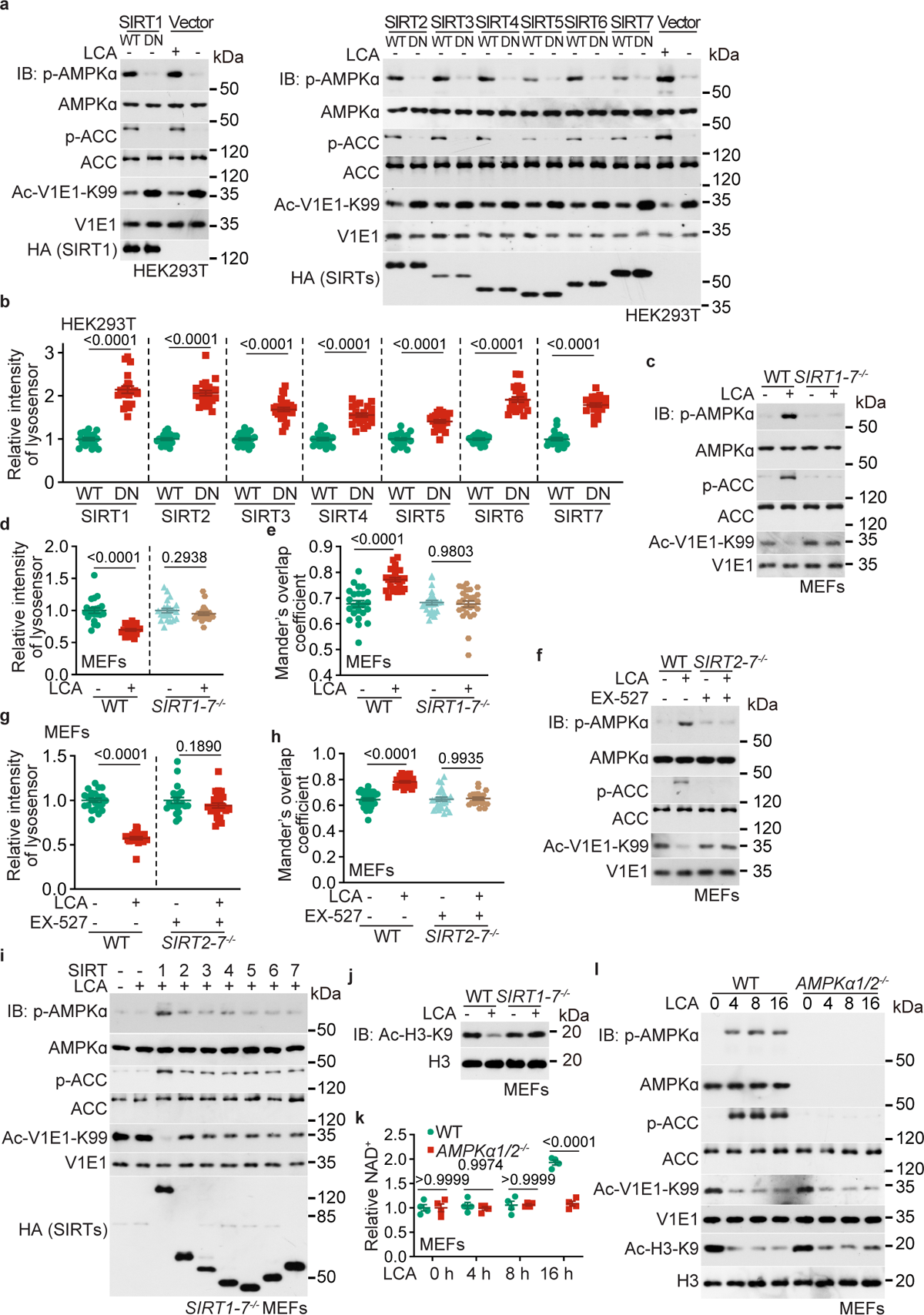
Sirtuins are required for the deacetylation of v-ATPase. **a, b**, Sirtuins are responsible for the deacetylation of V1E1 and AMPK activation. HEK293T cells were infected with lentiviruses carrying each sirtuin (HA-tagged SIRT1 to SIRT7), or their dominant negative (DN) form, followed by determining V1E1 acetylation and AMPK activity (**a**) and the v-ATPase activity (**b**; results are shown as mean ± s.e.m., normalised to the WT group of each sirtuin; *n* = 21 (SIRT2-DN and SIRT4-DN), 26 (SIRT3-DN), 23 (SIRT5-DN), 22 (SIRT6-DN) or 20 (others), and *P* value by two-sided Student’s *t*-test (SIRT4, SIRT5 and SIRT7) or two-sided Student’s *t*-test with Welch’s correction (others); the representative images are shown in Extended Data Fig. 4b). **c-h**, Sirtuins are required for the LCA-induced V1E1 deacetylation. MEFs with sirtuins depletion, either by hepta-knockout of SIRT1 to SIRT7 (**c-e**; *SIRT1*-*7*^-/-^, see validation data in Extended Data Fig. 4c), or by hexa-knockout of SIRT2 to SIRT7 (*SIRT2*-*7*^-/-^, validated in Extended Data Fig. 4g) and treatment with 10 μM EX-527 for 12 h to inhibit SIRT1 (**f-h**), were treated with 1 μM LCA for 4 h, followed by determination for the AMPK activation (**c**, **f**), the activity of v-ATPase (**d**, **g**; representative data in Extended Data Fig. 4d, h, and statistical analysis data as mean ± s.e.m., normalised to the DMSO group of each genotype; *n* = 22 (*SIRT1*-*7*^-/-^, LCA of **d**; WT, DMSO of **g**; and *SIRT2*-*7*^-/-^ + EX-527, DMSO of **g**), 25 (*SIRT2*-*7*^-/-^ + EX-527, LCA of **g**), or 21 (others), and *P* value by two-sided Student’s *t*-test with Welch’s correction (WT of **d**), two-sided Mann-Whitney test (*SIRT1*-*7*^-/-^ of **d**) or two-sided Student’s *t*-test (**g**)), and the translocation of AXIN (**e**, **h**; representative data in Extended Data Fig. 4e, i, and statistical analysis data as mean ± s.e.m.; *n* = 26 (*SIRT1*-*7*^-/-^, LCA of **e**; and *SIRT2*-*7*^-/-^ + EX-527, DMSO of **h**), 25 (WT, LCA of **h**), 30 (WT, DMSO of **h**), 21 (*SIRT2*-*7*^-/-^ + EX-527, LCA of **h**) or 24 (others), and *P* value by two-way ANOVA followed by Tukey’s test). **i**, Sirtuin members play complementary roles in mediating the LCA-induced V1E1 deacetylation and AMPK activation. The *SIRT1*-*7*^-/-^ MEFs were individually infected with lentivirus carrying 7 members of sirtuin family, followed by treating with 1 μM LCA for 4 h. The activity of AMPK and the acetylation of V1E1 were then determined by immunoblotting. **j-l**, LCA stimulates the activity of sirtuins before the elevation of NAD^+^. Wildtype, *SIRT1*-*7*^-/-^ and *AMPK*α*1*/*2*^-/-^ MEFs were treated with 1 μM LCA for 4 h (**j**) or indicated time periods (**k**, **l**), followed by determining the acetylation of histone H3 (on K9 residue, or Ac-H3-K9; **j**, **l**), the acetylation of V1E1 (**l**), the activity of AMPK (**l**), and the levels of NAD^+^ (**k**; results are shown as mean ± s.e.m.; *n* = 4 samples, and *P* value by two-way ANOVA followed by Tukey’s test). Experiments in this figure were performed three times, except **c** four times.

We then explored the mechanism of how LCA activates sirtuins. We first determined the levels of NAD^+^ which is a cosubstrate of sirtuins (ref. ^36,39^), and found that intracellular levels of NAD^+^ were increased only after 16 h of LCA treatment although LCA treatment for 4 h significantly stimulated the activity of sirtuins in MEFs (Fig. 3j-l). These results indicate that sirtuins are activated by LCA ahead of the increase of NAD^+^. We further found that the induction of NAD^+^ by LCA was dependent on AMPK (Fig. 3l), consistent with the previous findings that NAD^+^ upregulation is a downstream event of AMPK activation^40^. We next tested whether LCA could directly activate sirtuins by using a cell-free system in which purified bacterially expressed SIRT1, as a representative of sirtuins, was incubated with LCA at various concentrations, and found that LCA did not activate SIRT1 in such a reconstituted condition (Fig. 4a; see Extended Data Fig. 4j for the validation of SIRT1 activity assay). In addition, we found that LCA did not bind to bacterially expressed SIRT1, as determined by an affinity pull-down experiment using a synthetic LCA probe (Fig. 4b, see also Extended Data Fig. 5a-d for the synthesis procedures of LCA probe, and Fig. 4b and Extended Data Fig. 5e for the schematic depicting the procedure of the pull-down experiment using the LCA probe). However, after pre-incubation with lysates from MEFs, SIRT1 activity was greatly increased upon LCA addition (Fig. 4c). These observations suggested that LCA activates sirtuins via an unknown partner(s) in the cell lysate. We therefore wondered whether such an unknown component might interact with SIRT1 by pulling down HA-tagged-SIRT1 from the cell lysate and performed mass spectrometry on the co-pulled proteins. As listed in Supplementary Table 2, a total of 1,655 potential SIRT1-binding proteins were hit. We then engineered expression plasmids for all these 1,655 proteins, and verified that 157 of them could be co-immunoprecipitated by the HA-tagged SIRT1 when individually expressed in HEK293T cells (Supplementary Table 2). We then individually knocked down those 157 proteins in MEFs through lentivirus-mediated shRNA silencing, and found that TULP3, formerly known as a bipartite transcription regulator that binds DNA and phosphatidylinositol-phosphate^41–44^ and also identified as a hit among SIRT1-interacting proteins^45–47^, rendered the cells insensitive to LCA treatment, as assessed by levels of p-AMPK α, the acetylation of V1E1 and histone H3, and the activity of v-ATPase (Extended Data Fig. 6a, b). We also generated MEFs with *TULP3* knockout, and confirmed the requirement of TULP3 for the LCA-induced activation of AMPK and sirtuins (Fig. 4d, e and Extended Data Fig. 6d; see Extended Data Fig. 6c for the validation of *TULP3* knockout MEFs). We further found that the association between TULP3 and SIRT1 is constitutive, independently of LCA (Extended Data Fig. 6e). In support of such a notion was that the two purified bacterially expressed proteins of TULP3 and SIRT1 after incubation were co-eluted on size exclusion chromatography regardless of LCA addition (Fig. 4f). We also found that TULP3 interacted with all 7 members of the sirtuin family independently of LCA (Extended Data Fig. 6e). Additionally, in the presence of TULP3, LCA was able to activate the bacterially expressed SIRT1 (Fig. 4g), indicating that TULP3 bridges LCA to SIRT1. TULP3 was determined to bind LCA with a *K*_d_ at 15 μM by the competition assay (Fig. 4h; where the binding of the LCA probe to TULP3 was competed by LCA, to completion at approximately 40 μM). In silico docking assays revealed 4 residues of TULP3 (Y193, P195, K333 and P336) that comprise the potential binding pocket for LCA (Extended Data Fig. 6f). Indeed, mutation of these residues to glycine (TULP3-4G) blocked the LCA-mediated activation of SIRTs and AMPK in MEFs (Fig. 4i-m; Extended Data Fig. 6g, h). Different from TULP3, other known binding partners/targets of LCA and its derivatives, including FXR^48–50^, FXRb^51^, PXR^52,53^, VDR^54^, CAR^55–57^, LXRa^58^, LXRb^58^, S1PR2 (ref. ^59^), TGR5 (ref. ^60^), CHRM2 (ref. ^61^), CHRM3 (ref. ^62^), PKC ζ^63,64^, FAS^65,66^, FPR1^67^ and YES1^68^, did not mediate the activation of AMPK by LCA, as knockout of any of them in MEFs did not impair the activation of AMPK by LCA (Extended Data Fig. 7a-o). As an additional control, because some of the receptors share virtually identical structures of ligand-binding pockets^69,70^, we generated LXRa/b double-knockout MEFs, and found that LCA could still activate AMPK in these cells (MEFs; Extended Data Fig. 7p). For a similar reason, we generated double-knockouts of FXR and FXRb^51^, PXR and CAR^71^, and CHRM2 and 3 (ref. ^61,62^) in MEFs, and still found intact AMPK activation by LCA (Extended Data Fig. 7q-s). Together, it is TULP3 that binds LCA to activates SIRT1.

**Fig. 4.**
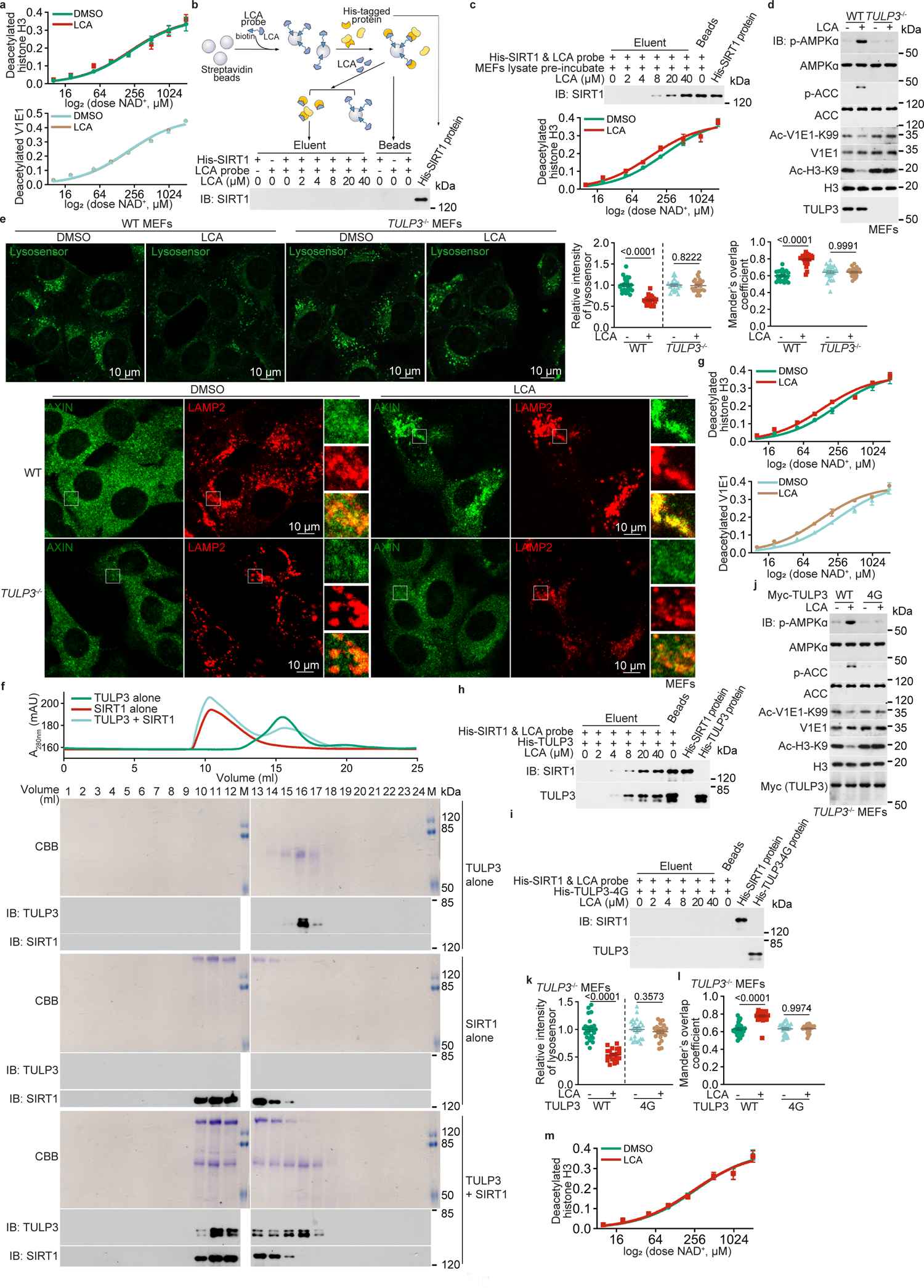
TULP3 is the binding protein of LCA for activation of sirtuins. **a-c**, Purified bacterially expressed SIRT1 requires a cytosolic partner(s) for binding to LCA. His-tagged SIRT1, pre-incubated with (**c**) or without (**a**, **b**) cellular lysates of MEFs, was incubated with close-to-intracellular concentrations of LCA (5 μM). The deacetylation activities of such pre-incubated SITR1 towards histone H3 and V1E1 (**a**, lower panel of **c**) were determined through a cell-free assay (see Methods section for details, and validation data in Extended Data Fig. 4j; results are shown as mean ± s.e.m., *n* = 4 samples), and the affinity of SIRT1 towards LCA through an affinity pull-down assay that used LCA probe as a bait, followed by competitive elution with LCA (**b**, upper panel of **c**; LCA probe was synthesised and purified as described in Extended Data Fig. 5a-d; the procedure of this assay was depicted in the upper panel of **b** and Extended Data Fig. 5e). **d**, **e**, TULP3 is required for SIRT1-mediated AMPK activation by LCA. TULP3 was identified as a SIRT1-interacting protein through MS on the hits of proteins co-pulled with SIRT1 (see Supplementary Table 2). The roles of TULP3 in mediating the LCA-induced AMPK activation were determined using MEFs with *TULP3* knockout (clone #1 of *TULP3*^-/-^ MEFs, and the same hereafter for all *TULP3* knockout experiments; validated in the upper panel of Extended Data Fig 6c; see also data for the clone #2 of *TULP3*^-/-^ MEFs in the lower panel of Extended Data Fig. 6c, and Extended Data Fig. 6d). After treating with 1 μM LCA for 4 h, the AMPK activation (**d**), the V1E1 and histone H3 acetylation (**d**), the v-ATPase activity (**e**, upper panel), and the lysosomal translocation of AXIN (**e**, lower panel) were determined. Statistical analysis data of **e** are shown as mean ± s.e.m., of which the v-ATPase activities were normalised to the DMSO group of each genotype; *n* = 22 (WT, DMSO of AXIN translocation), 23 (WT, LCA and *TULP3*^-/-^, DMSO of v-ATPase activity), 24 (*TULP3*^-/-^, LCA of v-ATPase activity), 26 (*TULP3*^-/-^, LCA of AXIN translocation) or 25 (others) cells; and *P* value by two-sided Student’s *t*-test (*TULP3*^-/-^, v-ATPase activity), two-sided Student’s *t*-test with Welch’s correction (WT, v-ATPase activity), or two-way ANOVA followed by Tukey’s test (AXIN translocation). **f**, TULP3 is constitutively associated with SIRT1 independent of LCA. TULP3 and SIRT1, both bacterially expressed and purified, were co-incubated, followed by subjecting to size exclusion chromatography. The elution chromatograms of SIRT1 alone (red), TULP3 alone (green), or SIRT1 co-incubated with TULP3 (cyan) were shown (**f**, upper panel). The presence of SIRT1-TULP3 complex was confirmed by the shift of the peak retention volume of TULP3 from 16 ml to 11 ml after incubating with SIRT1 (**f**, upper panel), and the presence of both SIRT1 and TULP3 in the fractions of the shifted peak (**f**, lower panel; analysed by SDS-PAGE and immunoblotting). CBB, Coomassie brilliant blue staining. **g**, **h**, TULP3 binds LCA and mediates activation of SIRT1. Experiments were performed as in **a** (for **g**; results are shown as mean ± s.e.m.; *n* = 4 samples) and **b** (for **h**), except that the bacterially expressed His-tagged SIRT1 was co-eluted with bacterially expressed His-tagged TULP3 on a Superdex column before each experiment. **i-m**, TULP3-4G unable to bind LCA blocks the LCA-induced activation of sirtuin and AMPK. The LCA-binding affinity (**i**) was determined as in **h**, except that the His-tagged TULP3-4G was used. The LCA-mediated AMPK activation was determined in **j-l**, in which the *TULP3*^-/-^ MEFs were infected with lentivirus carrying TULP3-4G mutant or its wildtype control, followed by treatment with 1 μM LCA for 4 h. The activity of AMPK (**j**), the acetylation of V1E1 (**j**), the activity of v-ATPase (**k**, left panel; representative images are shown in Extended Data Fig. 6g), and the lysosomal translocation of AXIN (**l**, right panel; representative images are shown in Extended Data Fig. 6h) were then determined. The effects of LCA on SIRT1 activity were determined in **m** (as in **g**, except that the TULP3-4G was co-eluted with SIRT1). The results are shown as mean ± s.e.m. *n* = 4 (**m**), 21 (WT, LCA of **k**), 26 (WT, LCA of **l**), 28 (WT, DMSO of **l**), 23 (4G, DMSO of **l**), 30 (4G, LCA of **l**) or 25 (others) samples; and *P* value by two-sided Student’s *t*-test (**k**, 4G), two-sided Student’s *t*-test with Welch’s correction (**k**, WT), or two-way ANOVA followed by Tukey’s test (**l**). Experiments in this figure were performed three times, except **d** four times.

We next determined whether the TULP3-sirtuins-v-ATPase axis mediates the rejuvenation and anti-ageing effects of LCA or CR in animal models. We found that muscle-specific expression (induced by tamoxifen) of V1E1-3KR, a deacetylated version of the V1E1 subunit of v-ATPase, led to AMPK activation (Extended Data Fig. 8a) and promoted muscle functions in aged mice by facilitating the transition of glycolytic fibres to oxidative fibres (Fig. 5a), independent of LCA. We also found that V1E1-3KR increased muscular NAD^+^ levels (Fig. 5b) and elevated muscular mitochondrial content (the mtDNA:nDNA ratios (Fig. 5c), the expression of mitochondrial genes and the OXPHOS proteins (Extended Data Fig. 8b)) and respiratory function (oxygen consumption rates (OCR); Fig. 5d) in aged mice. Consistently, a significant elevation of energy expenditure (Fig. 5e and Extended Data Fig. 8c, d), running distance, running duration and grip strength were observed in these mice (Fig. 5f, g). Muscle-specific knockout of AMPK impaired all benefits brought about by the expression of V1E1-3KR (Fig. 5a-g, and Extended Data Fig. 8a-d). To test the molecular mechanisms in the regulation of lifespan, we used *C*. *elegans* and *D*. *melanogaster* as surrogate models administered with LCA. For TULP3, we found that knockout of *tub-1* (TULP3 homologue in *C*. *elegans*) in the wildtype (N2) nematodes impaired the activation of AMPK by LCA (Extended Data Fig. 8e; LCA was absorbed into nematodes as effectively as into mouse muscles, as validated in the Extended Data Fig. 5a of the preceding paper) and abrogated the effects of LCA in lifespan extension (Extended Data Fig. 8f; see also statistical analyses on Extended Data Table 2, and for all the other lifespan data hereafter). Note that knockout of *tub-1* led to an extension of lifespan (Extended Data Fig. 8f), possibly owing to the inhibition of insulin signalling as described previously^72^. We also observed similar phenotypes in flies with *ktub* (TULP3 homologue in *D*. *melanogaster*) knocked down (Extended Data Fig. 8e, f; flies were able to absorb LCA, to a similar level as by mouse muscles, as validated in Extended Data Fig. 5b of the preceding paper). Re-introduction of wildtype TULP3, but not TULP3-4G (defective in binding to LCA), restored the extent of lifespan extension by LCA in nematodes and flies (Fig. 5h; see Extended Data Fig. 8g for the data on AMPK activation). Similarly, depletion of sirtuins in nematodes (by knocking down *sir*-*2.3* and *sir*-*2.4* in *sir*-*2.1*/*sir*-*2.2*-double knockout nematodes) and flies (by knocking out *Sir2*) impaired LCA-induced AMPK activation (Extended Data Fig. 8h) and lifespan extension (Fig. 5i). To test the role of the deacetylation of v-ATPase, nematodes and flies expressing V1E1-3KR showed AMPK activation, and extended lifespan (Fig. 5j and Extended Data Fig. 8i). We also found that the LCA-mediated extension of healthspan was impaired in nematodes and flies with expression of TULP3-4G or depletion of sirtuins, as they all impaired the LCA-induced resistance to oxidative stress and starvation (Fig. 5k, l and Extended Data Fig. 8j, k), the increase of mitochondrial contents and functions (Fig. 5m, n and Extended Data Fig. 8l-o), or the increase of NAD^+^ levels (Fig. 5o, p). In contrast, expression of V1E1-3KR extended healthspan independently of LCA (Fig. 5q-s and Extended Data Fig. 8p-r). The above results demonstrate that the TULP3-sirtuins-v-ATPase axis relays the signal of LCA to AMPK activation via deacetylation of the V1E1 subunit of the v-ATPase, which ultimately elicit the beneficial effects of CR in nematodes, flies and mice.

**Fig. 5.**
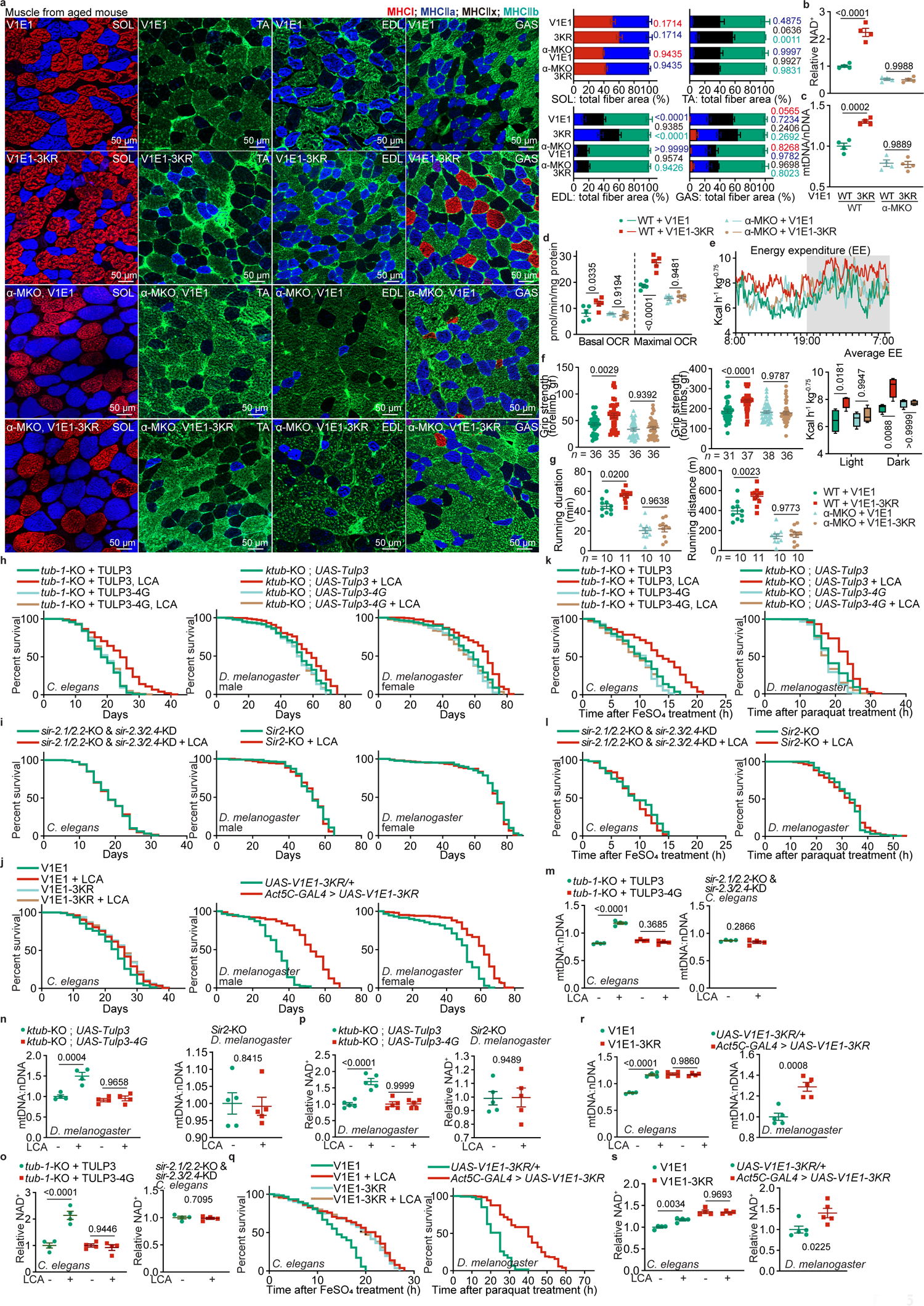
LCA-TULP3-sirtuins-v-ATPase axis exerts rejuvenating effects and retards ageing. **a**, V1E1-3KR induces oxidative fibre conversion in the muscle of aged mice. Muscle fibre type from wildtype or muscle-specific *AMPK*α knockout (α-MKO) mice with muscle-specific expression of V1E1-3KR (induced by tamoxifen at 18 months old, for 8 weeks; see “mouse strains” sections of the Methods section for the procedures for constructing this strain) were determined. The representative images from whole-muscle cross sections for soleus (SOL), extensor digitorum longus (EDL), tibialis anterior (TA) and gastrocnemius (GAS) muscle are shown on the upper panel. Statistical analysis data are shown as mean ± s.e.m.; *n* = 6 mice for each genotype/treatment, and *P* value by two-way ANOVA followed by Tukey’s test. **b**, V1E1-3KR induces elevation of muscle NAD^+^ in aged mice. Mice were treated as in **a**, followed by determining NAD^+^ levels in the gastrocnemius muscle. Results are shown as mean ± s.e.m.; *n* = 4 mice for each genotype/treatment, and *P* value by two-way ANOVA followed by Tukey’s test. **c**, **d**, V1E1-3KR elevates mitochondrial contents and mitochondrial respiratory function in the muscles of aged mice. Mice were treated as in **a**, followed by determining the ratios of mtDNA:nDNA (**c**), and the OCR (**d**) in the gastrocnemius muscle. Results are shown on the right panel as mean ± s.e.m.; *n* = 4 (**c**) or 5 (**d**) mice for each genotype/treatment, and *P* value by two-way ANOVA followed by Tukey’s test. The mRNA and protein levels of the OXPHOS complex in these mouse muscles are shown in Extended Data Fig. 8b. **e**, V1E1-3KR elevates energy expenditure (EE) in aged mice. Mice were treated as in **a**, followed by determining EE. Data are shown as mean (upper panel; at 5-min intervals during a 24-h course after normalisation to the body weight (kg^0.75^)), or as box-and-whisker plots (lower panel, in which the lower and upper bounds of the box represent the first and the third quartile scores, the centre line represents the median, and the lower and upper limits denote minimum and maximum scores, respectively; and the same hereafter for all box-and-whisker plots; *n* = 4 mice for each genotype/treatment, and *P* value by two-way ANOVA followed by Tukey’s test). The respiratory quotient (RQ) and the ambulatory activity data generated in this experiment are shown in Extended Data Fig. 8d. **f**, **g**, V1E1-3KR promotes muscle strength and running duration in aged mice. Mice were treated as in **a**, followed by determining the grip strength (**f**, results are mean ± s.e.m.), running duration (**g**, left panel; results are mean ± s.e.m.) and maximal running distance (**g**, right panel; results are mean ± s.e.m., generated according to the left panel of **g**). The *n* numbers are labelled in each panel, and *P* value by two-way ANOVA followed by Tukey’s test. **h-j**, TULP3, sirtuins and v-ATPase are required for lifespan extension by LCA. The *TULP3* knockout (*tub*-*1*-KO) nematodes re-introduced with TULP3-4G (**h**, left panel; see “*Caenorhabditis elegans* strains” sections of the Methods section for the procedures for constructing this strain), sirtuin depletion nematodes (**i**, left panel; *sir*-*2.1*/*sir*-*2.2*-double knockout (KO) nematodes with *sir*-*2.3* and *sir*-*2.4* knockdown (KD); see Methods section for details), the V1E1-3KR expressing nematodes (**j**, left panel), and their control nematodes were cultured in the medium containing LCA at 100 μM which led to an activation of AMPK in a similar way to that in mice (validated in Extended Data Fig. 5a of the preceding paper). Lifespan data are shown as Kaplan-Meier curves (see also statistical analyses in Extended Data Table 2, and the same hereafter for all lifespan data). Data obtained from experiments using flies with *TULP3* knockout (*ktub*-KO) re-introduced with TULP3-4G (*UAS*-*TULP3*-*4G*) (**h**, middle and right panels; see “*Drosophila melanogaster* strains” for details, and the same for other fly strains below), sirtuin knockout (*sir2*-KO) (**i**, middle and right panels), and the V1E1-3KR expressing (*Act5C*-*GAL4* > *UAS*-*V1E1*-*3KR*) (**j**, middle and right panels), during which flies were cultured in the med um containing 100 μM LCA that activates AMPK in a similar way to that in mice (see validation data in Extended Data Fig. 5b of the preceding paper), were also shown. **k**, **l**, **q**, The TULP3-sirtuins-v-ATPase axis mediates LCA to improve oxidative stress resistance. Nematodes and flies with TULP3-4G re-introduction in the *TULP3*-knockout background (**k**), sirtuins depletion (**l**), and V1E1-3KR expressing (**q**) were treated with 100 μM LCA for 2 days (left panels of **k**, **l**, **q**) or 30 days (right panels of **k**, **l**, **q**), followed by transferring to media containing 15 mM FeSO_4_ (left panels of **k**, **l**, **q**) or 20 mM paraquat (right panels of **k**, **l**, **q**), both elicited oxidative stress. Lifespan data are shown as Kaplan-Meier curves. See also results of experiments using 5% H_2_O_2_, as another oxidative stress inducer, in Extended Data Fig. 8j. **m**, **n**, **r**, The LCA-TULP3-sirtuins-v-ATPase axis elevates mitochondrial contents and mitochondrial respiratory function in nematodes and flies. Nematodes and flies with TULP3-4G re-introduction in the *TULP3*-knockout background (left panels of **m**, **n**), sirtuins depletion (right panels of **m**, **n**), and V1E1-3KR expressing (**r**) were treated with 100 μM LCA for 2 days (**m**, and left panel of **r**) or 30 days (**n**, and right panel of **r**), followed by determining the ratios of mtDNA:nDNA. Results are mean ± s.e.m.; *n* = 5 (right panels of **n**, **r**) or 4 (others) samples for each genotype/condition, and *P* and *P* value by two-way ANOVA followed by Tukey’s test (left panels of **m**, **n**, **r**) or two-sided Student’s *t*-test (right panels of **m**, **n**, **r**). The mRNA levels of the OXPHOS are shown in Extended Data Fig. 8l, n, q, and OCR in Extended Data Fig. 8m, o, r. **o**, **p**, **s**, The LCA-TULP3-sirtuins-v-ATPase axis elevates NAD^+^ levels in nematodes and flies. Nematodes and flies with TULP3-4G re-introduction in the *TULP3*-knockout background (left panels of **o**, **p**), sirtuins depletion (right panels of **o**, **p**), and V1E1-3KR expressing (**s**) were treated with 100 μM LCA for 2 days (**o**, and left panel of s) or 30 days (**p**, and right panel of s), followed by determining the ratios of NAD^+^. Results are mean ± s.e.m.; *n* = 5 (**p**, and right panel of **s**) or 4 (others) samples for each genotype/condition, and *P* value by two-way ANOVA followed by Tukey’s test (left panels of **o**, **p**, **s**) or two-sided Student’s *t*-test (right panels of **o**, **p**, **s**). Experiments in this figure were performed three times.

## Discussion

In this study, we have delineated how AMPK is activated by physiological levels of LCA seen in CR mice. We have identified TULP3 as the binding partner for LCA. Importantly, knockout of *TULP3*, or introduction of TULP3-4G mutant that cannot bind LCA, renders LCA entirely unable to activate AMPK in mice, supporting that TULP3 is required for LCA signalling. Once bound to LCA, TULP3 activates sirtuins, even before the increase of cellular levels of NAD^+^, which is increased only after prolonged activation of AMPK, as observed in this study (Fig. 3k) and others^40^. Therefore, through activating AMPK, LCA-TULP3 elicits a positive feedback loop for the production of NAD^+^ and the activities of sirtuins, particularly for those sirtuins whose *K*_m_ values for NAD^+^ are close to the intracellular levels of NAD^+^ in ad libitum fed tissues, such as SIRT1 (ref. ^73^). As for the other sirtuin members that have *K*_m_ values for NAD^+^ much lower than the regular cellular levels of NAD^+^, and are constantly saturated with NAD^+^, the CR-induced increase of NAD^+^ may not be essential for the activation of those sirtuins members (reviewed in ref. ^74^), such as SIRT2 (ref. ^73,75^), SIRT4 (ref. ^76^), and SIRT6 (ref. ^77^). Nevertheless, TULP3 is required for LCA to activate sirtuins, as shown in TULP3 knockout cells or animals and, given that TULP3 is a constitutive component of all 7 sirtuins. The common requirement of TULP3 for the activation of sirtuins is in line with the previous observations that all members of the sirtuin family participate in the mediation of the physiological effects of CR^78–85^. Apart from deacetylation^76,83,86^, some sirtuins, such as SIRT4 and SIRT5, also possess other activities including demethylglutarylation, dehydroxymethylglutarylation, demethylglutaconylation, desuccinylation, demalonylation, and deglutarylation activities^87–89^. It is therefore interesting to know whether these de-acylation activities may have other physiological functions than regulation of v-ATPase and AMPK.

The importance of the activation of sirtuins can be supported by the finding that LCA treatment can lead to deacetylation of the V1E1 subunit of v-ATPase at K52, K99 and K191 residues. Importantly, deacetylated version of V1E1 results in inhibition of the v-ATPase, which is a prerequisite for the lysosomal AMPK activation. These findings are consistent with the previous observation that when v-ATPase holoenzyme is reconstituted with some of V1 subunits, including V1E1, expressed and purified from *E*. *coli*, showed 5-fold lower proton pumping activity compared to the v-ATPase reconstituted with all components purified from calf brain^90^. Once inhibited, v-ATPase acts as an anchor, along with the Ragulator complex, for AXIN/LKB1 to translocate to the surface of the lysosome, where LKB1 phosphorylates and activates AMPK (Extended Data Fig. 8m). The intersection of the LCA-TULP3-sirtuins axis to the lysosomal AMPK pathway is further supported by the required role of AXIN and Ragulator (LAMTOR1) for response to LCA treatment. It is also interesting to note that the v-ATPase complex for AMPK activation can have multiple signalling routes converge upon it: in addition to the LCA-TULP3-sirtuin axis, the low glucose sensor aldolase-TRPV axis, and metformin-PEN2 axis also act upon the v-ATPase. The v-ATPase, therefore, appears to be a rotary for multiple pro-longevity inputs to intersect.

It is noteworthy that, despite LCA not being synthesised in nematodes or flies, the existence of the TULP3-sirtuin axis in these animals enables the administered LCA to exert life-extending roles. Our results thus point to a conserved signalling route involving sirtuins, NAD^+^ and AMPK. However, it remains to be elucidated how LCA extends lifespan in yeast, as it can benefit from LCA in terms of extending the chronological lifespan^91^, although it does not have conserved TULP3 orthologues. Finally, it must also be noted that CR or LCA treatment, at least as far as the late stage of CR is concerned, does not decrease blood glucose to below 5 mM which is necessary for the activation of AMPK, as seen during fasting or starvation of the ad libitum fed mice. Such a notion is also supported by the observations that disruption of the low glucose-sensing aldolase-TRPV axis, through the expression of ALDOA-D34S or knockout of TRPVs that are effective in blocking low glucose-induced AMPK activation, does not affect the LCA-mediated AMPK activation. The absence of hypoglycaemia in calorie-restricted mice has been observed in mice and humans by other researchers as well^92–95^, possibly owing to the enhanced gluconeogenesis, as well as the decreased glucose oxidation at the expense of increased fat and protein oxidation in these animals^96,97^.

Together, we have identified that TULP3 is a receptor of LCA, and upon binding to LCA, TULP3 activates sirtuins, which in turn deacetylates and inhibits v-ATPase, transmitting the LCA signal to the activation of the lysosomal AMPK pathway and further stimulation of NAD^+^ levels, thereby leading to anti-ageing effects in metazoans.

## Online content

Any methods, additional references, Nature Portfolio reporting summaries, source data, extended data, supplementary information, acknowledgements, details of author contributions and competing interests; and statements of data and code availability are available at https://doi.org/10.1038/.

### Publisher’s note

Springer Nature remains neutral with regard to jurisdictional claims in published maps and institutional affiliations.

## Supporting information

Uncropped gel images

Source data and statistical analysis data for all graphs

A complete list of the post-translational modifications identified for v-ATPase

List of the proteins co-immunoprecipitated with SIRT1

## Methods

### Data reporting

The chosen sample sizes were similar to those used in this field: *n* = 4-5 samples were used to evaluate the levels of metabolites in serum^98,99^, cells^17,100^, tissues^17,25,100,101^, nematodes^102–104^ and flies^105–107^; *n* = 4-5 samples to determine OCR in tissues^100,108^ and nematodes^109–111^; *n* = 3-4 samples to determine the mRNA levels of a specific gene^18^; *n* = 2-6 samples to determine the expression levels and phosphorylation levels of a specific protein^18^; *n* = 20-33 cells to determine AXIN translocation and lysosomal pH^17,19^; *n* = 4 replicates to determine SIRT1 activity^112,113^; *n* = 200 worms to determine lifespan^114–116^; *n* = 60 worms to determine healthspan^117–119^; *n* = 200 flies, male or female, to determine lifespan^120–122^; *n* = 60 flies, male or female, to determine healthspan^123–125^; *n* = 4-8 mice for EE and RQ^100^; *n* = 5 mice for body composition^100^; *n* = 6 mice for muscle fibre type^94,126,127^; *n* = 10-11 mice for running duration^19,100^; and *n* = 35-38 mice for grasp strength^100^. No statistical methods were used to predetermine sample size. All experimental findings were repeated as stated in figure legends, and all additional replication attempts were successful. For animal experiments, mice, nematodes and flies were housed under the same condition or place. For cell experiments, cells of each genotype were cultured in the same CO_2_ incubator and were seeded in parallel. Each experiment was designed and performed along with proper controls, and samples for comparison were collected and analysed under the same conditions. Randomisation was applied wherever possible. For example, during MS analyses (for metabolites and proteins), samples were processed and subjected to the MS in random orders. For animal experiments, sex-matched (for mice and flies), age-matched litter-mate animals in each genotype were randomly assigned to LCA or vehicle treatments. In cell experiments, cells of each genotype were parallel seeded and randomly assigned to different treatments. Otherwise, randomisation was not performed. For example, when performing immunoblotting, samples needed to be loaded in a specific order to generate the final figures. Blinding was applied wherever possible. For example, samples, cages or agar plates/vials during sample collection and processing were labelled as code names that were later revealed by the individual who picked and treated animals or cells, but did not participate in sample collection and processing, until assessing outcome. Similarly, during microscopy data collection and statistical analyses, the fields of view were chosen on a random basis, and are often performed by different operators, preventing potentially biased selection for desired phenotypes. Otherwise, blinding was not performed, such as the measurement of OCR and SIRT1 activity in vitro, as different reagents were added for particular reactions.

### Mouse strains

Wildtype C57BL/6J mice (#000664) were obtained from The Jackson Laboratory. *AXIN*^F/F^ (*AXIN1*^F/F^) and *LAMTOR1*^F/F^ mice were generated and validated as described previously^18^. *AMPK*α*1*^F/F^ (#014141) and *AMPK*α*2*^F/F^ mice (#014142) were obtained from Jackson Laboratory, provided by Dr. Sean Morrison, and *PKC*ζ^F/F^ mice (#024417) by Dr. Richard Huganir. *AXIN2*^F/F^ mice (Cat. T008456) were purchased from Shanghai Model Organisms Center, Inc. *LAMTOR1*-MKO and *LAMTOR1*-LKO mice were generated by crossing *LAMTOR1*^F/F^ mice with *Mck*-*Cre* and *Alb*-*Cre* mice as described and validated previously^18^, *AXIN*-LKO mice by crossing *AXIN*^F/F^ mice with *Alb*-*Cre* mice (validated in ref. ^21^), and *AXIN1*/*2*-MKO mice by crossing *AXIN1*/*2*^F/F^ mice with *HSA*-*CreERT2* mice (#025750; Jackson Laboratory). Muscular AXIN1 and AXIN2 from the *AXIN1*/*2*^F/F^, *HSA*-*CreERT2* mice were removed by intraperitoneally injecting mice with tamoxifen (dissolved in corn oil) at 200 mg/kg, 3 times a week.

Wildtype V1E1 or V1E1-3KR was introduced to the muscle of *AMPK*α*1*/*2*^F/F^ mice through the *Rosa26*-LSL(LoxP-Stop-LoxP) system^128^, followed by crossing with the *HSA-CreERT2* mice and then intraperitoneally injecting with tamoxifen. To introduce V1E1 or V1E1-3KR into *AMPK*α*1*/*2*^F/F^ mice, cDNA fragments encoding V1E1 or V1E1-3KR were inserted into the *Rosa26*-CTV vector^129^, followed by purification of the plasmids using CsCl density gradient ultracentrifugation method. Some 100 μg of plasmid was then diluted with 500 μl of di-distilled water, followed by concentrating via centrifuge at 14,000*g* at room temperature in a 30-kDa-cutoff filter (UFC503096, Millipore) to 50 μl of solution. The solution was diluted with 450 μl of di-distilled water, followed by another two rounds of dilution/concentration cycles. The plasmid was then mixed with 50 μl of di-distilled water to a final volume of 100 μl, followed by mixing with 10 μl of NaAc solution (3 M stock concentration, pH 5.2). The mixture was then mixed with 275 μl of ethanol, followed by incubating at room temperature for 30 min to precipitate plasmid. The precipitated plasmid was collected by centrifuge at 16,000*g* for 10 min at room temperature, followed by washing with 800 μl of 75% (v/v) ethanol (in di-distilled water) twice. After evaporating ethanol by placing the plasmid next to an alcohol burner lamp for 10 min, plasmid was dissolved in 100 μl of nuclease-free water. The plasmid, along with SpCas9 mRNA and the sgRNAs against the mouse *Rosa26* locus, was then microinjected into the in vitro fertilised (IVF) embryos of the *AMPK*α*1*/*2*^F/F^ mice. To generate the SpCas9 mRNA, 1 ng of pcDNA3.3-hCas9 plasmid (constructed by inserting the Cas9 fragment released from Addgene #41815 (ref. ^130^), into the pcDNA3.3 vector; diluted to 1 ng/ μl) was amplified using the Phusion High-Fidelity DNA Polymerase kit on a thermocycler (T100, Bio-Rad) with the following programmes: pre-denaturing at 98 °C for 30 s; denaturing at 98 °C for 10 s, annealing at 68 °C for 25 s, then extending at 72 °C for 2 min in each cycle; and final extending at 72 °C for 2 min; cycle number: 33. The following primer pairs were used: 5’-CACCGACTGAGCTCCTTAAG-3’, and 5’-TAGTCAAGCTTCCATGGCTCGA-3’. The PCR product was then purified using the MinElute PCR Purification Kit following the manufacturer’s instructions. The purified SpCas9 PCR product was then subjected to in vitro transcription using the mMESSAGE mMACHINE T7 Transcription Kit following the manufacturer’s instruction (with minor modifications). Briefly, 5.5 μl (300 ng/ μl) of SpCas9 PCR product as the template was mixed with 10 μl of 2× NTP/ARCA solution, 2 μl of 10×T7 Reaction Buffer, 0.5 μl of RNase inhibitor, 2 μl of T7 Enzyme Mix, and 4.5 μl of nuclease-free water, followed by incubating at 37 °C for 2 h. The mixture was then mixed with 1 μl of Turbo DNase, followed by incubating at 37 °C for 20 min to digest the template. The mixture was then mixed with 20 μl of 5× *E*-PAP Buffer, 10 μl of 25 mM MnCl_2_, 10 μl of 10 mM ATP, 4 μl of *E*-PAP enzyme, and 36 μl of nuclease-free water, followed by incubating at 37 °C for 20 min for poly(A) tailing. The tailed product was then purified using the MEGAclear Transcription Clean-Up Kit following the manufacturer’s instruction (with minor modifications). Briefly, some 20 μl of tailed RNA was mixed with 20 μl of Elution Solution, followed by mixing with 350 μl of Binding Solution Concentrate. Some 250 μl of ethanol was then added to the mixture, followed by passing the mixture through the Filter Cartridge and washing with 250 μl of Wash Solution twice. The RNA was then eluted with 50 μl of pre-warmed (at 90 °C) Elution Solution. The sgRNAs was prepared as in the SpCas9 mRNA preparation, except that: a) the gRNA Cloning Vector (Addgene, #41824, ref. ^130^) was used as template, and the following programmes: pre-denaturing at 98 °C for 30 s; denaturing at 98 °C for 10 s, annealing at 60 °C for 25 s, then extending at 72 °C for 20 s in each cycle; and final extending at 72 °C for 2 min; cycle number: 33; and the following primers: 5’-GAAATTAATACGACTCACTATAGGCGCCCATCTTCTAGAAAGACGTTTTA GAGCTAGAAATAGC-3’, and 5’-AAAAGCACCGACTCGGTGCC-3’; were used; b) in vitro transcription was performed using the MEGAshortscript T7 Transcription Kit, in which the mixture containing: 7.5 μl (100 ng/ μl) of purified PCR product, 2 μl of T7 10× T7 Reaction Buffer, 2 μl of T7 ATP solution, 2 μl of T7 CTP solution, 2 μl of T7 GTP solution, 2 μl of T7 UTP solution, 0.5 μl of RNase inhibitor, 2 μl of T7 Enzyme Mix, and 7.5 μl of nuclease-free water; was prepared. In addition, the poly(A) tailing assay was not performed.

The prepared *Rosa26*-CTV-V1E1, SpCas9mRNA and *Rosa26* sgRNA plasmids were then microinjected into each of the zygotes of *AMPK*α*1*/*2*^F/F^ mice. To prepare the zygotes, the *AMPK*α*1*^/^*2*^F/F^ mice (according to ref. ^131^, with modifications) were first subjected to IVF. Briefly, the 4-week-old *AMPK*α*1*/*2*^F/F^ female mice were intraperitoneally injected with pregnant mare’s serum gonadotrophin (PMSG) at a dose of 10 U/mouse. At 46 h after the PMSG injection, 10 U/mouse human chorionic gonadotrophin (hCG) was intraperitoneally injected. At 12 h after the hCG injection, oocytes from the oviducts of female mice, along with sperms from cauda epididymides and vasa deferentia of 16-week-old, proven s^tud^ *AMPK*α*1*/*2*^F/F^male mice, were isolated. To isolate oocytes, oviducts were briefly left on a filter paper, followed by incubating in a human tubal fluid medium (HTF)/GSH drop on an IVF dish (prepared by placing 200 μl of HTF solution supplemented with 125 mM GSH on a 35-mm dish to form a drop, followed by covering the drop with mineral oil and pre-balancing in a humidified incubator containing 5% CO_2_ at 37 °C for 0.5 h before use). The ampulla was then teared down by forceps, and the cumulus oocyte masses inside was collected and transferred to another HTF/GSH drop. To isolate sperms, cauda epididymides and vasa deferentia were briefly left on a filter paper, followed by penetrating with a 26 G needle on the cauda epididymides for 5 times. Sperms were then released to an HTF drop on sperm capacitation dish (prepared by placing 200 μl of HTF solution on a 35-mm dish to form a drop, followed by covering the drop with mineral oil and pre-balancing in a humidified incubator containing 5% CO_2_ at 37 °C for 12 h before use) by slightly pressing/squeezing the cauda epididymides, followed by incubating in a humidified incubator containing 5% CO_2_ at 37 °C for 0.5 h. The capacitated, motile sperms (located on the edge of each HTF drop) were then collected, followed by adding to the oocyte masses soaked in the HTF/GSH drop, 8 μl per drop. The IVF dishes containing oocyte masses and sperms were then cultured in a humidified incubator containing 5% CO_2_ at 37 °C for 4 h, followed by collecting and washing oocytes in a KSOM drop (freshly prepared by placing 20 μl of KSOM medium on a 35-mm dish to form a drop, followed by covering the drop with mineral oil and pre-balancing in a humidified incubator containing 5% CO_2_ at 37 °C for 0.5 h) twice. The oocytes were then cultured in an HTF/GSH drop on an IVF dish for another 12 h in a humidified incubator containing 5% CO_2_ at 37 °C. The presumptive zygotes (in which 2 pronuclei and an extruded, second polar body could be observed) were then picked up. Some 10 pl of DNA mixture comprising *Rosa26*-CTV-V1E1 plasmid (20 ng/μl final concentration), SpCas9 mRNA (120 ng/μl final concentration) and *Rosa26* sgRNA (100 ng/μl), was microinjected into each of the zygote, and were cultured in KSOM medium at 37 °C in a humidified incubator containing 5% CO_2_. At 16 h of culturing, the zygotes/embryos at two-cell stage were picked up and transplanted into pseudopregnant ICR female mice (8-10 weeks old, >26 g; prepared by breeding the in-estrus female with a 14-week-old, vasectomised male at a day before the transplantation), 20 zygotes/embryos per mouse, and the offspring carrying the LSL-V1E1 or LSL-V1E1-3KR allele were further outcrossed 6 times to C57BL/6 mice before crossing with the *HSA-CreERT2* mice. Mice were validated by genotyping. For genotyping the *Rosa26* locus, the following programmes were used: pre-denaturing at 98 °C for 300 s; denaturing at 95 °C for 30 s, annealing at 64 °C for 30 s, then extending at 72 °C for 45 s in each cycle for 5 cycles; denaturing at 95 °C for 30 s, annealing at 61 °C for 30 s, then extending at 72 °C for 45 s in each cycle for 5 cycles; denaturing at 95 °C for 30 s, annealing at 58 °C for 30 s, then extending at 72 °C for 45 s in each cycle for 5 cycles; denaturing at 95 °C for 30 s, annealing at 55 °C for 30 s, then extending at 72 °C for 45 s in each cycle for 5 cycles; and final extending at 72 °C for 10 min. For genotyping other genes and elements, the following programmes were used: pre-denaturing at 95 °C for 300 s; denaturing at 95 °C for 30 s, annealing at 58 °C for 40 s, then extending at 72 °C for 30 s in each cycle; and final extending at 72 °C for 10 min; cycle number: 35. The following primers: 5’-CAGGTAGGGCAGGAGTTGG-3’ and 5’-TTTGCCCCCTCCATATAACA-3’ for *HSA*-*Cre*; 5’-AGTGGCCTCTTCCAGAAATG-3’ and 5’-TGCGACTGTGTCTGATTTCC-3’ for the control of *HSA*-*Cre*; 5’-CCCACCATCACTCCATCTCT-3’ and 5’-AGCCTGCTTGGCACACTTAT-3’ for *Prkaa1*, 5’-GCAGGCGAATTTCTGAGTTC-3’ and 5’-TCCCCTTGAACAAGCATACC-3’ for *Prkaa2*, 5’-TCTCCCAAAGTCGCTCTGAG-3’, 5’-AAGACCGCGAAGAGTTTGTC-3’, and 5’-ATGCTCTGTCTAGGGGTTGG-3’ for *Rosa26*, 5’-CCACACAGGCATAGAGTGTCT-3’ and 5’-TTTGCACAAGCCGACCTTTC-3’ for 5’-terminus of *Rosa26*-*V1E1* (the presence of *Rosa26*-*V1E1* recombination), and 5’-CTCCACACAGGCATAGAGTGT-3’ and 5’-TTGCACAAGCCGACCTTTCT-3’ for 3’-terminus of *Rosa26*-*V1E1*, 5’-AGAGAATTCGGATCCATGGCTCTCAGCGATGCTGA-3’ and 5’-CTTCCATGGCTCGAGGTCCAAAAACTTCCTGTTGGC-3’ for generating PCR products for sequencing V1E1.

### CR and fasting treatments of mice

Protocols for all rodent experiments were approved by the Institutional Animal Care and the Animal Committee of Xiamen University (XMULAC20180028 and XMULAC20220050). Unless stated otherwise, mice were housed with free access to water and standard diet (65% carbohydrate, 11% fat, 24% protein) under specific pathogen-free conditions. The light was on from 8:00 to 20:00, with the temperature kept at 21-24 °C and humidity at 40-70%. Only male mice were used in the study, and male littermate controls were used throughout the study.

Mice were individually caged for 1 week before each treatment. For fasting, the diet was withdrawn from the cage at 5 p.m., and mice were sacrificed at desired time points by cervical dislocation. For CR, each mouse was fed with 2.5 g of standard diet (approximately 70% of ad libitum food intake for a mouse at 4 months old and older) at 5 p.m. of each day. For creating the diabetic mouse model, mice were fed a HFD (60% calories from fat; D12492, Research Diets) for desired time periods starting at 4 weeks old.

The following ages of mice were used: a) for immunoblotting and the measurement of adenylates: wildtype mice aged 4weeks (treated with LCA for 1 week starting from 4 weeks old), or the *AXIN*-LKO, *AXIN*-MKO, *LAMTOR1*-LKO and *LAMTOR1*-MKO mice aged 7 weeks (treated with LCA for 1 week starting from 6 weeks old); b) for immunohistochemistry: the V1E1-3KR- or wildtype V1E1-expressing mice aged 18 months (into which tamoxifen was injected at 16 months old for 2 months); c) for determination of mouse healthspan: V1E1-3KR- or wildtype V1E1-expressing mice aged 18 months (into which tamoxifen was injected at 16 months old for 2 months); and d) for all the other experiments, mice aged 4 weeks.

### Formulation of LCA

LCA was formulated as described in our preceding paper. Briefly, for cell-based experiments, LCA powder was dissolved in DMSO to a stock concentration of 500 mM, and was aliquoted and stored at −20 °C. The solution was placed at room temperature for 10 min (until no precipitate was visible) before adding to the culture medium. Note that any freeze-thaw cycle was not allowed to avoid the re-crystallisation of LCA (otherwise forms in sheet-like, insoluble crystals) in the stock solution.

For mouse experiments, LCA was coated with (2-hydroxypropyl)- β-cyclodextrin before treating animals. To coat LCA, the LCA powder was dissolved in 100 ml of methanol to a concentration of 0.01 g/ml, followed by mixing with 308 ml of (2-hydroxypropyl)- β-cyclodextrin solution (by dissolving (2-hydroxypropyl)- β-cyclodextrin in 30% (v/v, in water) methanol to 0.04 g/ml, followed by a 30 min of sonication). The control vehicle was similarly prepared, with no LCA added to the (2-hydroxypropyl)- β-cyclodextrin solution. After evaporating at 50 [, 90 r.p.m. by the rotary evaporator (Rotavator R-300, Vacuum Pump V-300, BUCHI), the coated powder was stored at 4 °C for no more than 2 weeks and was freshly dissolved in free drinking water to 1 g/l before feeding mice.

For nematode experiments, LCA at desired concentrations was freshly dissolved in DMSO, and was added to warm (cooled to approximately 60 °C after autoclaving) nematode growth medium^132^ (NGM, containing 0.3% (wt/vol) NaCl, 0.25% (wt/vol) bacteriological peptone, 1 mM CaCl_2_, 1 mM MgSO_4_, 25 mM KH_2_PO_4_-K_2_HPO_4_, pH 6.0, 0.02% (wt/vol) streptomycin, and 5 μg/ml cholesterol. The medium was used to make the NGM plate by adding 1.7% (wt/vol) agar. The plates were stored at 20 °C for no more than 3 days.

For fly experiments, LCA was coated and dissolved in water as in the mouse experiments, and was added to the BDSC Standard Cornmeal Medium^133^ (for regular culture). The BDSC Standard Cornmeal Medium was prepared as described previously, with minor modifications^133^. Briefly, 60.5 g of dry yeast, 35 g of soy flour, 255.5 g of cornmeal, 20 g of agar, and 270 ml of corn syrup were mixed with 3,500 ml of water in a stockpot. The mixture was thoroughly stirred using a long-handled soup spoon, and then boiled, during which lumps formed were pressed out using the back of the spoon. After cooling to approximately 60 °C, 16.8 ml of propionic acid was added to the medium, followed by stirring with the spoon. The medium was then dispensed into the culture vials, 6 ml each. The media vials were covered with a single-layer gauze, followed by blowing with the breeze from a fan at room temperature overnight. Some 100 μl of LCA solution (dissolved in the (2-hydroxypropyl)- β-cyclodextrin as for mice) at desired concentration was then layered (added dropwise) onto the medium surface of each vial, followed by blowing with the breeze for another 8 h at room temperature. The media vials were kept at 4 °C (for no more than 3 days) before experiment.

### Determination of mouse running capacity and grip strength

The maximal running capacity was determined as described previously^19,134^, with minor modifications. Briefly, mice were trained on Rodent Treadmill NG (UGO Basile, cat. 47300) for 3 days with normal light-dark cycle, and tests were performed during the dark period. Before the experiment, mice were fasted for 2 h. The treadmill was set at 5° incline, and the speed of treadmill was set to increase in a ramp-mode (commenced at a speed of 5 m/min followed by an increase to a final speed of 25 m/min within 120 min). Mice were considered to be exhausted, and removed from the treadmill, following the accumulation of 5 or more shocks (0.1 mA) per minute for two consecutive minutes. The distances travelled were recorded as the running capacity.

Grip strength was determined on a grip strength meter (Ugo Basile, cat. 47200) following the protocol described previously^119^. Briefly, the mouse was held by its tail and lowered (“landed”) until forelimb or all four limbs grasped the T[bar connected to a digital force gauge. The mouse was further lowered to the extent that the body was horizontal to the apparatus, and was then slowly, steady drawn away from the T[bar until forelimb or all four limbs were removed from the bar, which gave rise to the peak force in grams. Each mouse was repeated 5 times with 5 min intervals between measurements.

### Determination of body composition

Lean and fat body mass were measured by quantitative magnetic resonance (EchoMRI-100H Analyzer; Echo Medical Systems) as described previously^100^. Briefly, the system was calibrated with oil standard before the measurement. Mice were individually weighted and inserted into a restrainer tube, and were immobilised by gently inserting a plunger. The mouse was then positioned to a gesture that curled up like a donut, with its head against the end of the tube. Body composition of each mouse was measured with 3 repeated runs, and the average values were taken for further analysis.

### Determination of energy expenditure

Mouse energy expenditure (EE) was determined by a metabolic cage system (Promethion Line, CAB-16-1-EU; Sable Systems International) as described previously^135^. Briefly, the system was maintained in a condition identical to that for housing mice. Each metabolic cage in the 16-cage system consisted of a cage with standard bedding, a food hopper and water bottle, connected to load cells for continuous monitoring. To minimise the stress of new environment, mice were acclimatised (by individually housing in the gas-calibrated chamber) for 1 week before data collection. Mice treated with LCA or vehicle control were randomly assigned/housed to prevent systematic errors in measurement. Body weights and fat proportion of mice were determined before and after the acclimation, and the food intake and water intake daily. Mice found not acclimatised to the metabolic cage (e.g., resist to eat and drink) were removed from the study. Data acquisition (5-min intervals each cage) and instrument control were performed using MetaScreen software (v.2.3.15.12, Sable Systems) and raw data processed using Macro Interpreter (v.2.32, Sable Systems). Ambulatory activity and position were monitored using XYZ beam arrays with a beam spacing of 0.25 cm (beam breaks), and the mouse pedestrial locomotion (walking distance) within the cage were calculated accordingly. Respiratory gases were measured using the GA-3 gas analyser (Sable Systems) equipped with a pull-mode, negative-pressure system. Air flow was measured and controlled by FR-8 (Sable Systems), with a set flow rate of 2,000 ml/min. Oxygen consumption (VO_2_) and carbon dioxide production (VCO_2_) were reported in ml per minute. Water vapour was measured continuously and its dilution effect on O_2_ and CO_2_ was compensated mathematically in the analysis stream. Energy expenditure (EE) was calculated using: kcal/h = 60 × (0.003941 × VO_2_ + 0.001106 × VCO_2_) (Weir Equation). Differences of average EE were analysed by analysis of covariance (ANCOVA) using body weight as the covariate. Respiratory quotient (RQ) was calculated as VCO_2_/VO_2_.

### Histology

Muscle fibre types were determined as described previously^94,136^. Briefly, muscle tissues were excised, followed by freezing in isopentane (pre-chilled in liquid nitrogen) for 2 min (until they appeared chalky white). The tissues were then quickly transferred to embedding molds containing O.C.T. Compound, and were frozen in liquid nitrogen for another 10 min. The embedded tissues were then sectioned into 6- μm slices at −20 °C using a CM1950 Cryostat (Leica), followed by fixing in 4% paraformaldehyde for 10 min, and washing with running water for 5 min at room temperature. After incubating with PBST (PBS supplemented with 5% (v/v) Triton X-100) for 10 min, the sections were blocked with BSA Solution (PBS containing 5% (m/v) BSA) for 30 min at room temperature. Muscle fibres were stained with the antibody against MHCIIb (6 μg/ml, diluted in BSA Solution) overnight at 4 °C, followed by washing with PBS for 3 times, 5 min each at room temperature. The sections were then incubated with Alexa Fluor 488-conjugated, goat anti-mouse IgM antibody (1:200 diluted in BSA Solution) for 1 h at room temperature in a dark humidified chamber, followed by washing with PBS for 3 times, 5 min each, incubated with 4% paraformaldehyde for 2 min, and then washed with PBS twice, 5 min each, all at room temperature. The sections were then incubated with antibody against MHCI (6 μg/ml, diluted in BSA Solution) for 3 h at room temperature in a dark humidified chamber, followed by washing with PBS buffer for 3 times, 5 min each at room temperature, and then incubated with Alexa Fluor 594-conjugated, goat anti-mouse IgG2b antibody (1:200 diluted in BSA Solution) for another 1 h at roo temperature in a dark humidified chamber, followed by washing with PBS buffer for 3 times, 5 min each at room temperature. After fixing with 4% paraformaldehyde for 2 min and washing with PBS twice, 5 min each at room temperature, the sections were incubated with the antibody against MHCIIa (6 μg/ml, diluted in BSA Solution) for 3 h at room temperature in a dark humidified chamber, followed by washing with PBS buffer for 3 times, 5 min each at room temperature, and then incubated in Alexa Fluor 647-conjugated goat anti-mouse IgG1 antibody (1:200 diluted in BSA Solution) for another 1 h at room temperature in a dark humidified chamber, followed by washing with PBS buffer for 3 times, 5 min each at room temperature. Tissue sections were mounted with 90% glycerol and visualised on an LSM980 microscope (Zeiss). Images were processed and analysed on Zen 3.4 software (Zeiss), and formatted on Photoshop 2023 software (Adobe).

### Caenorhabditis elegans strains

Nematodes (hermaphrodites) were maintained on NGM plates spread with *E. coli* OP50 as standard food. All worms were cultured at 20 °C. Wildtype (N2 Bristol) and *sir*-*2.1* (VC199) strains were obtained from *Caenorhabditis* Genetics Center, and *sir*-*2.2* (tm2673) from National BioResource Project. All mutant strains were outcrossed 6 times to N2 before the experiments. Unless stated otherwise, worms were maintained on NGM plates spread with *Escherichia coli* OP50 as standard food. The administration of LCA was initiated at the L4 stage.

The *sir*-*2.1*/*sir*-*2.2*-double knockout strain was generated by crossing *sir*-*2.1*-knockout with *sir*-*2.2*-knockout strains. Before crossing, *sir*-*2.1*-knockout hermaphrodites were synchronised: worms were washed off from agar plates with 15 ml of M9 buffer (22.1 mM KH_2_PO_4_, 46.9 mM Na_2_HPO_4_, 85.5 mM NaCl and 1 mM MgSO_4_) supplemented with 0.05% (v/v) Triton X-100 per plate, followed by centrifugation at 1,000*g* for 2 min. The worm sediment was suspended with 6 ml of M9 buffer containing 50% synchronising bleaching solution (by mixing 25 ml of NaClO solution (5% active chlorine), 8.3 ml of 25% (w/v) NaOH, and 66.7 ml of M9 buffer, for a total of 100 ml), followed by vigorous shaking for 2 min, and centrifugation for 2 min at 1,000*g*. The sediment was washed with 12 ml of M9 buffer twice, then suspended with 6 ml of M9 buffer, followed by rotating at 20 °C, 30 r.p.m. for 12 h. The synchronised worms were then transferred to the NGM plate and cultured to the L4 stage, followed by heat-shocking at 28 °C for 12 h. The heat-shocked worms were then cultured at 20 °C for 4 days, and the males were picked up for mating with *sir*-*2.1*-knockout hermaphrodites for another 36 h. The mated hermaphrodites were transferred to new NGM plates and allowed to give birth to more *sir*-*2.1*-knockout males for another 4 days at 20 °C. The *sir*-*2.1*-knockout males were then picked up and co-cultured with *sir*-*2.2*-knockout hermaphrodites at a 1:2 ratio (e.g., 4 males and 8 hermaphrodites on a 10-cm NGM plate) for mating for 36 h at 20 °C, and the mated hermaphrodites (*sir*-*2.2*-knockout) were picked up for culturing for another 2 days. The offspring were then picked up and were individually cultured on the 35-mm NGM plate, followed by being individually subjected for genotyping after egg-laying (after culturing for approximately 2 days). For genotyping, individual worms were lysed with 5 μl of Single Worm lysis buffer (50 mM HEPES, pH 7.4, 1 mM EGTA, 1 mM MgCl_2_, 100 mM KCl, 10% (v/v) glycerol, 0.05% (v/v) NP-40, 0.5 mM DTT and protease inhibitor cocktail). The lysates were then frozen at −80 °C for 12 h, followed by incubating at 65 °C for 1 h and 95 °C for 15 min on a thermocycler (XP Cycler, Bioer). The lysates were then cooled to room temperature, followed by amplifying genomic DNA on a thermocycler with the following programmes: pre-denaturing at 95 °C for 10 min; denaturing at 95 °C for 10 s, then annealing and extending at 60 °C for 30 s in each cycle; cycle number: 35. Primer sequences are as follows: *C*. *elegans sir*-*2.1*, 5’-GAATCGGCTCGTTGCAAGTC-3’ and 5’-AGTTGTGGAATGTCATGGATCCT-3’; and *C*. *elegans sir*-*2.2*, 5’-TACCGCTCGAAAGATGTGGG-3’ and 5’-CTGGAGCCACGTGTTCTTCT-3’. The offspring generated from *sir*-*2.1*- and *sir*-*2.2*-knockout-assured individuals were then outcrossed 6 times to the N2 strain.

The *sir*-*2.3* and *sir*-*2.4* genes were then knocked down in the *sir*-*2.1*/*sir*-*2.2*-double knockout strain following the previously described procedures^23^. Briefly, synchronised worms (around the L1 stage) were placed on the RNAi plates (NGM containing 1 mg/ml IPTG and 50 μg/ml carbenicillin) spread with HT115 *E*. *coli* stains containing RNAi against *sir*-*2.3* and *sir*-*2.4* (well C10 on plate X6, and well K04 on plate I9 from the Ahringer *C*. *elegans* RNAi Collection) for 2 days. The knockdown efficiency was then examined by determining the levels of *sir*-*2.3* and *sir*-*2.4* mRNAs by qPCR, in which approximately 1,000 worms were washed off from an RNAi plate with 15 ml of M9 buffer containing Triton X-100, followed by centrifugation for 2 min at 1,000*g*. The sediment was then washed with 1 ml of M9 buffer twice, and then lysed with 1 ml of TRIzol. The worms were then frozen in liquid nitrogen, thawed at room temperature and then subjected to repeated freeze-thaw for another two times. The worm lysates were then placed at room temperature for 5 min, then mixed with 0.2 ml of chloroform followed by vigorous shaking for 15 s. After 3 min, lysates were centrifuged at 20,000*g* at 4 °C for 15 min, and 450 μl of the aqueous phase (upper layer) was transferred to a new RNase-free centrifuge tube (Biopur, Eppendorf), followed by mixing with 450 μl of isopropanol, then centrifuged at 20,000*g* at 4 °C for 10 min. The sediments were washed with 1 ml of 75% ethanol (v/v) followed by centrifugation at 20,000*g* for 10 min, and then with 1 ml of anhydrous ethanol followed by centrifugation at 20,000*g* for 10 min. The sediments were then dissolved with 20 μl of RNase-free water after the ethanol was evaporated. The dissolved RNA was then reverse-transcribed to cDNA using ReverTra Ace qPCR RT master mix with a gDNA Remover kit, followed by performing real-time qPCR using Maxima SYBR Green/ROX qPCR master mix on a CFX96 thermocycler (Bio-Rad) with the programmes as described in genotyping the *sir*-*2.1*/*2.2*-knockout strain. Primer sequences are as follows: *C*. *elegans sir*-*2.3*, 5’-ACTCTCTCGCCTGTGCAAAT-3’ and 5’-ACTTCCAACGATGTCCCAAG-3’; and *C*. *elegans sir*-*2.4*, 5’-GCCGTAAAAAGTTTGAGCCC-3’ and 5’-TTTCCATGCTTTTCGGATT-3’. Data were analysed using CFX Manager software (v.3.1, Bio-Rad). Knockdown efficiency was evaluated according to the CT values obtained.

The *tub*-*1* knockout nematode strains expressing human TULP3 or TULP3-4G were established through a 3-step strategy as described previously^23^, with minor modifications: a) TULP3 or TULP3-4G was first introduced to the N2 strain; b) such generated strains were then subjected to knockout of the *tub*-*1* gene; and c) the *tub*-*1*-knockout worms were then picked up for the further outcrossing with N2 strain. Briefly, to generate N2 strain expressing TULP3 or TULP3-4G, cDNAs of them were inserted into a pJM1 vector, with GFP as a selection marker, between the *Nhe* I and *Kpn* I sites (expressed under control by a *sur*-*5* promoter), and then injected into the syncytial gonad of the worm (200 ng/µl, 0.5 µl per worm) following the procedure as described previously^23^. The injected worms were then recovered on a NGM plate for 2 days, and the F_1_ GFP-expressing hermaphrodites were selected for further culture. The extrachromosomally TULP3 or TULP-S4G expression plasmid was then integrated into the nematode genome using UV irradiation to establish non-mosaic transgenic strains as described previously^137^, with minor modifications. Briefly, 70 TULP3- or TULP3-4G-expressing worms at L4 stage were picked up and incubated with 600 µl of M9 buffer, followed by adding 10 µl of TMP solution (3 mg/ml stock concentration in DMSO) and rotating at 30 rpm for 15 min in the dark. Worms were then transferred to a 10-cm NGM plate without OP50 bacteria in the dark, followed by irradiating with UV at a total dose of 35 J/cm^2^ (exposed for 35 s) on a UV crosslinker (CL-508; UVITEC). The irradiated worms were fed with 1 ml of OP50 bacteria at 10^13^/ml concentration, and then cultured at 20 °C for 5 h in the dark, followed by individually cultured on 35-mm NGM plate for 1 week without transferring to any new NGM plate (to make sure that F_1_ was under starvation before further selection). The F_1_ GFP-expressing hermaphrodites were selected and individually cultured for another 2 days, and those F_2_ with 100% GFP-expressing hermaphrodite were selected for further culture. The genomic sequence encoding *tub*-*1* was then knocked out from this strain by injecting a mixture of a pDD122 (P*eft*-*3*::*Cas9* + *ttTi5605* sgRNA) vector carrying sgRNAs targeting *tub*-*1* (5’-ATAGCTGATCAAAAGTTCTA-3’ for intron 2 and 5’-GAGAGCGGTCAGTGACACGG-3’ for intron 7; inserted into the pDD122 vector with the *ttTi5605* sgRNA sequence replaced) that are designed using the CHOPCHOP website http://chopchop.cbu.uib.no/), into young adult worms. The F_1_ hermaphrodite worms were individually cultured on an NGM plate. After egg-laying, worms were lysed using Single Worm lysis buffer, followed by PCR with the programmes as described in genotyping the *sir*-*2.1*/*2.2*-knockout strain, and the primers 5’-GAGTAATTTTCGGCATTTGTGC-3’ and 5’-CGAGAAGCTCATTTCAAGGTTT-3’ were used. The offspring generated from knockout-assured individuals were outcrossed 6 times to the N2 strain, and the expression levels of TULP3 or TULP3-4G were examined by immunoblotting. Strains expressing TULP3 or TULP3-4G at similar levels were chosen for further experiments.

Nematode strain (N2) with human V1E1 or V1E1-3KR expression was constructed as described above, except that cDNAs of human V1E1 or V1E1-3KR were used, and that the *sur*-*5* promoter on the pJM1 vector was replaced by the promoter of V1E1 (vha-8) itself (by replacing the sequence between *Asc* I and *Fse* I sites with the annealed primer pair 5’-CTGACTGGGCCGGCCCTTAGAGATAGACTGTGGTC-3’ and 5’-CTCTAGAGGCGCGCCGACATTTAATAAAATAATCATTTTTC-3’). The presence of V1E1 or V1E1-3KR was validated by sequencing using the primer 5’-ATGGCTCTCAGCGATGCTGA-3’. For all nematode experiment, worms at L4 stage were used.

### Evaluation of nematode lifespan and healthspan

To determine the lifespan of nematodes, the worms were first synchronised. Synchronised worms were cultured to L4 stage before transfer to desired agar plates for determining lifespan. Worms were transferred to new plates every 2 d. Live and dead worms were counted during the transfer. Worms that displayed no movement upon gentle touching with a platinum picker were judged as dead. Kaplan-Meier curves were graphed by Prism 9 (GraphPad Software), and the statistical analysis data by SPSS 27.0 (IBM).

The resistance of nematodes to the oxidative stress was determined as described previously^117^. Briefly, synchronised worms were cultured to L4 stage after which LCA was administered. After 2 days of LCA treatment, 20 worms were transferred to an NGM plate containing 15 mM FeSO_4_. Worms were then cultured at 20 [on such a plate, during which the live and dead worms were counted at every 1 h.

### Drosophila melanogaster strains

All flies were cultured at 25 °C and 60% humidity with a 12-hour light and dark cycle. Adult flies were cultured in Bloomington *Drosophila* Stock Center (BDSC) Standard Cornmeal Medium (for regular culture) agar diet. Larvae and the crossed fly strains were reared on Semi-Defined, Rich Medium, which was prepared as described previously^138^, with minor modifications. Briefly, 10 g of agar, 80 g of dry yeast, 20 g of yeast extract, 20 g of peptone, 30 g of sucrose, 60 g of glucose, 0.5 g of MgSO_4_·6H_2_O and 0.5g of CaCl_2_·6H_2_O were dissolved in 1,000 ml of di-distilled water, and then boiled, followed by cooling to 60 °C. Some 6 ml of propionic acid was then added to the medium, and the medium was dispensed into culture vials, 6 ml each. The media vials were covered with gauze and blown with the breeze as in BDSC diet, and were kept at 20 °C (for no more than 3 days) before experiment. Fly embryos were prepared using the grape juice plates. The plates were prepared by mixing 30 g of agar in 1 l of water, followed by boiling in a microwave for 3 rounds, 3 min per round. Some 360 ml of grape juice, 2 g of methyl paraben and 30 g of sucrose were then sequentially added into the agar solution, with each component mixed thoroughly on a heated magnetic stirrer. The medium was dispensed into 100-cm Petri dishes, 10 ml each, and were kept at 4 °C (for no more than 1 week) before experiment. The yeast paste used for collecting embryo was freshly prepared by thoroughly mixing 7 g of dry yeast with 9 ml of water in a 50-ml conical flask with a metal spatula, until it reached the consistency of peanut butter. The paste was then dabbed at the centre of each grape juice plate before experiment.

The wildtype fly strain (*w*^111^*^8^*; #3605), *Sir2^4.5^* strain (*Sirt1^4.5^ cn^1^/SM6b*, *P*{*ry^+t7.2^=eve-lacZ8.0*}*SB^1^*; #32568), *Sir2^5.^*^26^ strain (*Sirt1^5.^*^26^ *cn^1^*; #32657), Cas9-expressing strain (*y*^1^ *sc*\* *v*^1^ *sev*^21^; *P*{*y*^+t7.7^ *v*^+t1.8^=*nanos*-*Cas9*.R}*attP2*; #78782), Bloomington DB strain (*w*^1118^; *wg^Sp-1^*/*CyO*; *MKRS*/*TM6B*, *Tb^1^*; #76357) and the *GAL4*-expressing strain (*y^1^ w\**; *P*{*Act5C-GAL4-w*}*E1/CyO*; #25374) were obtained from the BDSC. The *GAL4*-induced, *ktub* RNAi-carrying strain (*w*^1118^; *P*{*GD14210*}*v29111*; #29111) was obtained from the Vienna *Drosophila* Resource Center (VDRC). The *yw*R13S strain (*yw*; *Sp*/*CyO*; *MKRS*/*TM2*), CAS DB strain (*w^1118^*; *BL1*/*CyO*; *TM2*/*TM6B*), *attp*3# (68A4) strain (*y*^1^ *M*{*vas*-*int*.*Dm*}*ZH*-*2A w*\*; P{*CaryP*}*attP2*) and the *attp*2# (25C6) strain (*y*^1^ *M*{*vas*-*int*.*Dm*}*ZH*-*2A w*\*; P{*CaryP*}*attP40*) were obtained from Core Facility of *Drosophila* Resource and Technology, Chinese Academy of Sciences. Files with *Sir2* knockout were obtained by crossing the *Sir2^4.5^* and the *Sir2^5.26^* strains, followed by picking the F_1_ offspring with red eyes and straight wings (*Sir2^4.5^/Sir2^5.26^*). Flies with *GAL4* expressed on the *w^1118^* background (*w^1118^*; *P*{*Act5C-GAL4-w*}*E1/CyO*) was first generated as described in our preceding paper by crossing the *y^1^ w\**; *P*{*Act5C-GAL4-w*}*E1/CyO* males with *w^1118^*; *Sp/CyO* females, followed by crossing the F1 males with straight wings (*w^1118^*; *P*{*Act5C-GAL4-w*}*E1/Sp*) with *w^1118^*; *Sp/CyO* females.

To generate *TULP3* (*ktub*, *CG9398*) knockout flies, the synchronised (see “Evaluation of lifespan and healthspan of flies” for details), Cas9-expressing flies were housed in a collection cage (cat. 59-101; Genesee Scientific) containing a grape juice plate for 2 days. Flies were then allowed to lay embryos for 3 rounds, with each round lasting for 1 h, on a fresh grape juice plate that contains a dab of yeast paste at the centre. Some 800 embryos were then collect by filling each grape juice plate with water, followed by gently dislodging the embryos with a paint brush, and then filtered by passing through a 70- μm Cell Strainer (cat. 350350; Falcon). The embryos in the strainer were then vigorously rinsed with tap water at 2 ml/s for 3 times, 30 s each, followed by roughly dried on a paper towel. The embryos were them dechorionated by dipping the strainer into freshly prepared 50% bleach solution (diluted by water) filled in a 10-cm petri dish for 1 min, followed by rinsing with tap water at 2 ml/s for 3 times, 30 s each.

After roughly dried on a paper towel, embryos were aligned with their ventral side sitting on a segment of double-sided tape (cat. 665; 3MScotch) that was stuck onto a slide, followed by covering with a drop of mineral oil (1:1 - Halocarbon oil 700:Halocarbon oil 27). Some 2 nl of 500 ng/μl gRNAs targeting the exon 3 of *ktub* (5’-CGCATTACGCGGGACAGGAA-3’ and 5’-CGGAGATCTCATCGCACGAGT-3’) were then injected into the posterior pole of the embryo following the procedures as described previously^23^. The injected embryos were cultured at 18 °C for 48 h on an IHC transparent humidified chamber, and the larvae were then transferred to the Semi-Defined, Rich Medium and cultured at 25 °C for another week to for the F_0_ adults. The F_0_ adults were then individually mated with *yw*R13S adults, and any F_1_ offspring with straight wings were discarded. The remaining F_1_ female offspring with curled wings were then screened for the presence of *ktub* knockout through genotyping following the procedures outlines in the “analysis of mitochondrial DNA copy numbers in mice, nematodes and flies” section, with following primers: 5’-AGATCAATCGACCCATGTCCG-3’ and 5’-CTTGCCGTAGTCACGTTCCA-3’, and the F_1_ male, curled-wing offspring (possibly with the *yw*; *ktub*^-/-^/*CyO*; +/*TM2* genotype) from those female counterparts with *ktub* knockout were individually mated with virgin females of the CAS DB strain. After 4 days of mating, each F_1_ males was subjected to genotype the knockout *ktub* knockout, and the F_2_ male offspring (possibly the *w^1118^*; *ktub*^-/-^/*CyO*; +/*TM6B*) generated from those *ktub*-knockout-assured F_1_ males were picked and mated again with virgin females of CAS DB strain. The F_3_ male, curled-wing offspring (possibly the *w^1118^*; *ktub*^-/-^/*CyO*; +/*TM6B*) were then individually mated with virgin females of F_3_ curled-wing offspring. After 10 days of mating, each pair of F_3_ male and female were subjected to examine the knockout of *ktub*, and the F_4_ offspring that generated from the breeding of those F_3_ males and females with *ktub* knockout were selected to breed with each other, leading to the generation of the *ktub*^-/-^ (*w^1118^*; *ktub*^-/-^/*ktub*^-/-^) flies.

Human TULP3-4G was re-introduced to the *ktub*^-/-^ flies by expressing TULP3-4G in the *attp*3# flies (with TULP3 inserted into the chromosome III). The modified flies were then sequentially crossed with *yw*R13S and DB flies to backcross to a *w^1118^* background, followed by crossing with the *ktub*^-/-^ flies. Briefly, to generate *attp*3# flies with TULP3-4G expression, cDNAs of TULP3-4G were inserted into the pUAST-attB vector between the *Xho* I and *Xba* I sites. The construct was then injected into the embryos of *attp*3# flies as in the sgRNA injection, except the concentration at 300 ng/ μl. The F_0_ adults were then individually mated with *yw*R13S flies, and the F_1_ males with curled wings and orange eyes (possibly the *yw*; +/*CyO*; *TULP3*-*4G*/*TM2*) were mated with the virgin females of the CAS DB strain. The F_2_ males with curled wings and white eyes (*w^1118^*; +/*CyO*; *TULP3*-*4G*/*TM6B*) were mated with virgin females of the CAS DB strain again, followed by crossing the F_3_ males with curled wings and white eyes with F_3_ virgin females with curled wings and white eyes to generate the TULP3-4G-expressing (*w^1118^*; +/+; *TULP3*-*4G*/*TM6B*) flies. The TULP3-4G-expressing strains and the *ktub*^-/-^ strains were separately crossed with Bloomington DB flies. The F_1_ offspring of TULP3-4G with curled wings (*w^1118^*; +/*CyO*; *TULP3*-*4G*/*MKRS*) and the F_1_ offspring of *ktub*^-/-^ with additional bristles in the humerus (*w^1118^*; *ktub*^-/-^/*wg^Sp-1^*; +/*TM6B*, *Tb^1^*), were then mated together, and the F_2_ offspring with curled wings and additional bristles in the humerus (possibly the *w^1118^*; *ktub*^-/-^/*Cyo*; *TULP3*-*4G*/*TM6B*, *Tb^1^*) were picked. The expression of TULP3-4G was achieved by crossing the picked F_2_ offspring with the *GAL4*-expressing strain (*w^1118^*; *P*{*Act5C-GAL4-w*}*E1/CyO*), and the F_3_ offspring with straight wings and regular bristles (*w^1118^*; *P*{*Act5C-GAL4-w*}*E1/ktub*^-/-^; *TULP3*-*4G*/+) were used for further experiments. The TULP3-re-introduced, *ktub*^-/-^ flies were similarly generated, except the TULP3 cDNA were used.

The human V1E1-3KR was introduced to the wildtype flies as in TULP3, except that the *attp*2# strain (with V1E1 inserted into the chromosome II) was used as the acceptor. The F_0_ adults were then sequentially mated with *yw*R13S flies, CAS DB flies, the F_2_ offspring themselves and the *GAL4*-expressing strain as in TULP3. The *w^1118^*; *P*{*Act5C-GAL4-w*}*E1/V1E1*-*3KR* were used for further experiments.

### Evaluation of lifespan and healthspan of flies

Fly lifespan was determined as described previously^139^, with minor modifications. Before the experiment, flies were synchronised: approximately 200 pairs of flies, housed 10 pairs per tube, were cultured in Semi-Defined, Rich Medium and allowed to lay eggs for a day. After discarding the parent flies, the embryos were cultured for another 10 days, and the flies eclosed at day 12 were anaesthetised and collected with CO_2_ (those emerged before day 12 were discarded), followed by transferring to the BDSC Standard Cornmeal Medium and cultured for another 2 days. The male and female adults were then sorted by briefly anaesthetised with CO_2_ on the anaesthetic pad using a homemade feather brush (by attaching the apical region of a vane from the secondary coverts of an adult goose, to a plastic balloon stick), and some 200 adults of each group/gender were randomly assigned to the BDSC Standard Cornmeal Medium, containing LCA or not, 20 flies per tube. The files were transferred to new medium tubes every two days without anaesthesia until the last survivor was dead. During each tube transfer, the sum of dead flies in the old tubes and the dead flies carried to the new tubes were recorded as the numbers of deaths, and the escaped or accidentally died flies (i.e., died within 3 days of same-sex culturing, or squeezed by the tube plugs) were censored from the experiments. Kaplan-Meier curves were graphed by Prism 9 (GraphPad Software), and the statistical analysis data by SPSS 27.0 (IBM).

The resistance of flies to the oxidative stress was determined as described previously^140^. Briefly, synchronised adults were treated with LCA for 30 days, followed by transferring to vials (20 flies each), each containing a filter paper soaked with 20 mM paraquat or 5% (m/v) H_2_O_2_ dissolved/diluted in 5% (w/v, in water) glucose solution. To determine the resistance of flies to starvation (food deprivation), flies treated with LCA for 30 days were transferred to vials with culture medium replaced by the same volume of 1.5% agarose to remove food supply. Dead files were recorded every 2 h until the last survivor was dead.

### Quantification of mRNA levels of mitochondrial genes in mice, nematodes and flies

Mice treated with LCA were sacrificed by cervical dislocation, immediately followed by dissecting gastrocnemius muscle. The muscle tissue was roughly sliced to cubes having edge lengths of approximately 2 mm, and then soaked in RNAprotect Tissue Reagent (1 ml per 100 mg of tissue) for 24 h at room temperature. The tissue was then incubated in 1 ml of TRIzol, followed by three rounds of freeze/thaw cycles, and was then homogenised. The homogenate was centrifuged at 12,000g for 15 min at 4 °C, and 900 µl of clear supernatant (not the lipid layer on the top) was transferred to an RNase-free tube. The supernatant was then added with 200 µl of chloroform, followed by vigorous vortex for 15 s. After centrifugation at 12,000*g* for 15 min at 4 °C, 450 µl of the upper aqueous layer was transferred to an RNase free tube. The RNA was then precipitated by adding 450 µl of isopropanol, followed with centrifugation at 12,000*g* for 30 min at 4 °C. The pellet was washed twice with 75% ethanol, and once with 100% ethanol, and was dissolved with 20 µl of DEPC-treated water. The concentration of RNA was determined by a NanoDrop 2000 spectrophotometer (Thermo). Some 1 µg of RNA was diluted with DEPC-treated water to a final volume of 10 µl, heated at 65 °C for 5 min, and chilled on ice immediately. The Random Primer Mix, Enzyme Mix and 5× RT buffer (all from the ReverTra Ace qPCR RT Master Mix) were then added to the RNA solution, followed by incubation at 37 °C for 15 min, and then at 98 °C for 5 min on a thermocycler. The reverse-transcribed cDNA was quantified with Maxima SYBR Green/ROX qPCR Master Mix on a LightCycler 480 II System (Roche) with following programmes: pre-denaturing at 95 °C for 10 min; denaturing at 95 °C for 10 s, then annealing and extending at 65 °C for 30 s in each cycle (determined according to the amplification curves, melting curves, and bands on agarose gel of serial pilot reactions (in which a serial annealing temperature was set according to the estimated annealing temperature of each primer pair) of each primer pair, and same hereafter), for a total of 45 cycles. Primer pairs for mouse *Nd1*, *Nd2*, *Nd3*, *Nd4*, *Nd4l*, *Nd5*, *Nd6*, *Ndufab1*, *Cytb*, *Uqcrc1*, *Uqcrc2*, *Atp5f1b*, *Cox6a1*, *Atp6*, *Atp8*, *Cox1* and *Cox3* were generated as described previously^141^, and others using the Primer-BLAST website (https://www.ncbi.nlm.nih.gov/tools/primer-blast/index.cgi). Primer sequence are as follows: mouse *Gapdh*, 5’-GACTTCAACAGCAACTCCCAC-3’ and 5’-TCCACCACCCTGTTGCTGTA-3’; mouse *Nd1*, 5’-TGCACCTACCCTATCACTCA-3’ and 5’-C GGCTCATCCTGATCATAGAATGG-3’; mouse *Nd2*, 5’- ATACTAGCAATTACTTCTATTTTCATAGGG-3’ and 5’-GAGGGATGGGTTGTAAGGAAG-3’; mouse *Nd3*, 5’-AAGCAAATCCATATGAATGCGG-3’ and 5’-GCTCATGGTAGTGGAAGTAGAAG-3’; mouse *Nd4*, 5’-CCTCAGACCCCCTATCCACA-3’ and 5’-GTTTGGTTCCCTCATCGGGT-3’; mouse *Nd4l*, 5’-CCAACTCCATAAGCTCCATACC-3’ and 5’-GATTTTGGACGTAATCTGTTCCG-3’; mouse *Nd5*, 5’-ACGAAAATGACCCAGACCTC-3’ and 5’-GAGATGACAAATCCTGCAAAGATG-3’; mouse *Nd6*, 5’-TGTTGGAGTTATGTTGGAAGGAG-3’ CAAAGATCACCCAGCTACTACC-3’; and mouse 5’-*Tfam*, 5’-GGTCGCATCCCCTCGTCTAT-3’ and 5’-TTGGGTAGCTGTTCTGTGGAA-3’; mouse *Cs*, 5’-CTCTACTCACTGCAGCAACCC-3’ and 5’-TTCATGCCTCTCATGCCACC-3’; mouse *Ndufs8*, 5’-TGGCGGCAACGTACAAGTAT-3’ and 5’-GTAGTTGATGGTGGCAGGCT-3’; mouse *Ndufab1*, 5’-GGACCGAGTTCTGTATGTCTTG-3’ and 5’-AAACCCAAATTCGTCTTCCATG-3’; mouse *Ndufb10*, 5’-TGCCAGATTCTTGGGACAAGG-3’ and 5’-GTCGTAGGCCTTCGTCAAGT-3’; mouse *Ndufv3*, 5’-GTGTGCTCAAAGAGCCCGAG-3’ and 5’-TCAGTGCCGAGGTGACTCT-3’; mouse *Ndufa8*, 5’-GCGGAGCCTTTCACAGAGTA-3’ and 5’-TCAATCACAGGGTTGGGCTC-3’; mouse *Ndufs3*, 5’-CTGACTTGACGGCAGTGGAT-3’ and 5’-CATACCAATTGGCCGCGATG-3’; mouse *Ndufa9*, 5’-TCTGTCAGTGGAGTTGTGGC-3’ and 5’-CCCATCAGACGAAGGTGCAT-3’; mouse *Ndufa10*, 5’-CAGCGCGTGGGACGAAT-3’ and 5’-ACTCTATGTCGAGGGGCCTT-3’; mouse *Sdha*, 5’-AGGGTTTAATACTGCATGCCTTA-3’ and 5’-TCATGTAATGGATGGCATCCT-3’; mouse *Sdhb*, 5’-AGTGCGGACCTATGGTGTTG-3’ and 5’-AGACTTTGCTGAGGTCCGTG-3’; mouse *Sdhc*, 5’-TGAGACATGTCAGCCGTCAC-3’ and 5’-GGGAGACAGAGGACGGTTTG-3’; mouse *Sdhd*, 5’-TGGTACCCAGCACATTCACC-3’ and 5’-GGGTGTCCCCATGAACGTAG-3’; mouse *Cytb*, 5’-CCCACCCCATATTAAACCCG-3’ and 5’-GAGGTATGAAGGAAAGGTATTAGGG-3’; mouse *Uqcrc1*, 5’-ATCAAGGCACTGTCCAAGG-3’ and 5’-TCATTTTCCTGCATCTCCCG-3’; mouse *Uqcrc2*, 5’-TTCCAGTGCAGATGTCCAAG-3’ and 5’-CTGTTGAAGGACGGTAGAAGG-3’; mouse *Atp5f1b*, 5’-CCGTGAGGGCAATGATTTATAC-3’ and 5’-GTCAAACCAGTCAGAGCTACC-3’ mouse *Cox6a1*, 5’-GTTCGTTGCCTACCCTCAC-3’ and 5’-TCTCTTTACTCATCTTCATAGCCG-3’; mouse *Atp6*, 5’-TCCCAATCGTTGTAGCCATC-3’ and 5’-TGTTGGAAAGAATGGAGTCGG-3’; mouse *Atp8*, 5’-GCCACAACTAGATACATCAACATG-3’ and 5’-TGGTTGTTAGTGATTTTGGTGAAG-3’; mouse *Atp5f1a*, 5’-CATTGGTGATGGTATTGCGC-3’ and 5’-TCCCAAACACGACAACTCC-3’; mouse *Cox1*, 5’-CCCAGATATAGCATTCCCACG-3’ and 5’-ACTGTTCATCCTGTTCCTGC-3’; mouse *Cox2*, 5’-TCTACAAGACGCCACATCCC-3’ and 5’-ACGGGGTTGTTGATTTCGTCT-3’; mouse *Cox3*, 5’-CGTGAAGGAACCTACCAAGG-3’ and 5’-CGCTCAGAAGAATCCTGCAA-3’; mouse *Cox5b*, 5’-AGCTTCAGGCACCAAGGAAG-3’ and 5’-TGGGGCACCAGCTTGTAATG-3’.

The mRNA level was then calculated using the comparative Δ Δct method using the LightCycler software (v.96 1.1, Roche; same hereafter for all qPCR experiments). Nematodes at L4 stage treated with LCA for 1 day were used for analysis of mitochondrial gene expression. Some 1,000 worms were collected with 15 ml of M9 buffer containing 0.05% Triton X-100 (v/v), followed by centrifugation for 2 min at 1,000*g*. The sediment was then washed with 1 ml of M9 buffer twice, and then lysed with 1 ml of TRIzol. Worms were then frozen in liquid nitrogen, thawed at room temperature, and then repeated freeze-thaw for another 2 times. The worm lysates were then placed at room temperature for 5 min, then mixed with 0.2 ml of chloroform, followed by vigorous shaking for 15 s. After centrifugation at 12,000*g* for 15 min at 4 °C, 450 µl of the upper aqueous layer was transferred to an RNase free tube. The RNA was then precipitated by adding 450 µl of isopropanol, followed with centrifugation at 12,000*g* for 30 min at 4 °C. The pellet was washed twice with 75% ethanol, and once with 100% ethanol, and was dissolved with 20 µl of DEPC-treated water. The concentration of RNA was determined by a NanoDrop 2000 spectrophotometer (Thermo). Some 1 µg of RNA was diluted with DEPC-treated water to a final volume of 10 µl, heated at 65 °C for 5 min, and chilled on ice immediately. The Random Primer Mix, Enzyme Mix and 5× RT buffer (all from the ReverTra Ace qPCR RT Master Mix) were then added to the RNA solution, followed by incubation at 37 °C for 15 min, and then at 98 °C for 5 min on a thermocycler. The reverse-transcribed cDNA was quantified with Maxima SYBR Green/ROX qPCR Master Mix on a LightCycler 480 II System (Roche) with following programmes: pre-denaturing at 95 °C for 10 min; denaturing at 95 °C for 10 s, then annealing and extending at 65 °C for 30 s in each cycle (determined according to the amplification curves, melting curves, and bands on agarose gel of serial pilot reactions (in which a serial annealing temperature was set according to the estimated annealing temperature of each primer pair) of each primer pair, and same hereafter), for a total of 45 cycles. Primer pairs used for qPCR are as previously described^142,143^, except that *C*. *elegans ctb-1* were designed using the Primer-BLAST website. Primer sequences are as follows: *C*. *elegans ama-1*, 5’-GACATTTGGCACTGCTTTGT-3’ and 5’-ACGATTGATTCCATGTCTCG-3’; *C*. *elegans nuo-6*, 5’-CTGCCAGGACATGAATACAATCTGAG-3’ and 5’-GCTATGAGGATCGTATTCACGACG-3’; *C*. *elegans nuaf-1*, 5’-GAGACA TAACGAGGCTCGTGTTG-3’ and 5’-GAAGCCTTCTTTCCAATCACTATCG-3’; *C*. *elegans sdha-1*, 5’-TTACCAGCGTGCTTTCGGAG-3’ and 5’-AGGGTGTGGAGAAGAGAATGACC-3’; *C*. *elegans sdhb-1*, 5’-GCTGAACGTGATCGTCTTGATG-3’ and 5’-GTAGGATGGGCATGACGTGG-3’; *C*. *elegans cyc-2.1*, 5’-CGGA GTTATCGGACGTACATCAG-3’ and 5’-GTCTCGCGGGTCCAGACG-3’; *C*. *elegans isp-1*, 5’-GCAGAAAGATGAATGGTCCGTTG-3’ and 5’-ATCCGTGACAAGGGCAGTAATAAC-3’; *C. elegans cco-1*, 5’-GCTGGAGATGATCGTTACGAG-3’ and 5’-GCATCCAATGATTCTGAAGTCG-3’; *C. elegans cco-2*, 5’-GTGATACCGTCTACGCCTACATTG-3’ and 5’-GCTCTGGCACGAAGAATTCTG-3’; *C. elegans atp-3*, 5’-GTCCTCGACCCAACTCTCAAG-3’ and 5’-GTCCAAGGAAGTTTCCAGTCTC-3’; *C*. *elegans nduo-1*, 5’-AGCGTCATTTATTGGGAAGAAGAC-3’ and 5’-AAGCTTGTGCTAATCCCATAAATGT-3’; *C*. *elegans nduo-2*, 5’-TCTT TGTAGAGGAGGTCTATTACA-3’ and 5’-ATGTTAAAAACCACATTAGCCCA-3’; *C. elegans nduo-4*, 5’-GCACACGGTTATACATCTACACTTATG-3’ and 5’-GATGTATGATAAAATTCACCAATAAGG-3’; *C*. *elegans nduo-5*, 5’-AGATGAGATTTATTGGGTATTTCTAG-3’ and 5’-CACCTAGACGATTAGTTAATGCTG-3’; *C*. *elegans ctc-1*, 5’-GCAGCAGGGTTAAGATCTATCTTAG-3’ and 5’-CTGTTACAAATACAGTTCAAACAAAT-3’; *C*. *elegans ctc-2*, 5’-GTAGTTTATTGTTGGGAGTTTTAGTG-3’ and 5’-CACAATAATTCACCAAACTGATACTC-3’; *C*. *elegans atp-6*, 5’-TGCTGCTGTAGCGTGATTAAG-3’ and 5’-ACTGTTAAAGCAAGTGGACGAG-3’; *C*. *elegans ctb-1*, 5’-TGGTGTTACAGGGGCAACAT-3’ and 5’-TGGCCTCATTATAGGGTCAGC-3’.

The mRNA level was then calculated using the comparative Δ Δct method using the LightCycler software (v.96 1.1, Roche; same hereafter for all qPCR experiments). *Drosophila* adults treated with LCA for 30 days were used to determine the expression of mitochondrial genes. For each sample, some 20 adults were used. The adults were anesthetised, transferred to a 1.5-ml Eppendorf tube, followed by quickly freezing in liquid nitrogen, and then homogenised using a pellet pestle (cat. Z359963-1EA, Sigma). The homogenate was then lysed in 1 ml of TRIzol for 5 min at room temperature, followed by centrifuged at 12,000*g* for 15 min at 4 °C. Some 900 μl of supernatant (without the lipid layer) was transferred to an RNase-free tube, followed by mixing with 200 μl of chloroform. After vigorous vortexing for 15 s, the mixture was centrifuged at 12,000*g* for 15 min at 4°C, and some 450 μl of the upper aqueous layer was transferred to an RNase-free tube. The RNA was then precipitated by adding 450 μl of isopropanol, followed by centrifugation at 12,000*g* for 30 min at 4 °C. The pellet was washed twice with 75% (v/v, in water) ethanol, and was dissolved with 20 μl of DEPC-treated water. The concentration of RNA was determined by a NanoDrop 2000 spectrophotometer (Thermo). Some 1 µg of RNA was diluted with DEPC-treated water to a final volume of 10 µl, heated at 65 °C for 5 min, and chilled on ice immediately. The Random Primer Mix, Enzyme Mix and 5× RT buffer (all from the ReverTra Ace qPCR RT Master Mix) were then added to the RNA solution, followed by incubation at 37 °C for 15 min, and then at 98 °C for 5 min on a thermocycler. The reverse-transcribed cDNA was quantified with Maxima SYBR Green/ROX qPCR Master Mix on a LightCycler 480 II System (Roche) with following programmes: pre-denaturing at 95 °C for 5 min; denaturing at 95 °C for 10 s, then annealing at 60 °C for 20 s, and then extending at 72 °C for 20 s in each cycle, for a total of 40 cycles. Primer pairs used for qPCR are as previously described^144^, and are listed as follows: *D*. *melanogaster CG9172*, 5’-CGTGGCTGCGATAGGATAAT-3’ and 5’-ACCACATCTGGAGCGTCTTC-3’; *D*. *melanogaster CG9762*, 5’-AGTCACCGCATTGGTTCTCT-3’ and 5’-GAGATGGGGTGCTTCTCGTA-3’; *D*. *melanogaster CG17856*, 5’-ACCTTTCCATGACCAAGACG-3’ and 5’-CTCCATTCCTCACGCTCTTC-3’; *D*. *melanogaster CG18809*, 5’-AAGTGAAGACGCCCAATGAGA-3’ and 5’-GCCAGGTACAACGACCAGAAG-3’; *D*. *melanogaster CG5389*, 5’-ATGGCTACAGCATGTGCAAG-3’ and 5’-GACAGGGAGGCATGAAGGTA-3’; *C. melanogaster Act5C*, 5’-GCAGCAACTTCTTCGTCACA-3’ and 5’-CATCAGCCAGCAGTCGTCTA-3’.

### Analysis of mitochondrial DNA copy numbers in mice, nematodes and flies

Mouse mitochondrial DNA copy numbers were determined as described previously^100^. Briefly, mouse tissue DNA was extracted with the Biospin tissue genomic DNA extraction kit (BioFlux) following the manufacturer’s instruction, with minor modifications. Briefly, mice treated with LCA were sacrificed by cervical dislocation, quickly followed by dissecting gastrocnemius muscle. The muscle tissue was then grinded on a ceramic mortar in liquid nitrogen. Some 50 mg of grinded tissue was then transferred to a 1.5-ml Eppendorf tube, followed by addition of 600 µl of FL buffer and 10 µl of PK solution containing 2 µl of 100 mg/ml RNase A. The mixture was then incubated at 56 [for 15 min, followed by centrifuge at 12,000*g* for 3 min.

Some 500 µl of supernatant was transferred to a 2-ml Eppendorf tube, followed by mixing with 700 µl of binding buffer and 300 µl of absolute ethanol. The mixture was then loaded onto a Spin column, and was centrifuged at 10,000*g* for 1 min. The flowthrough was discarded, and 500 µl of the PW buffer was added to the Spin column, followed by centrifuge at 10,000*g* for 30 sec. Some 600 µl of washing buffer was then added to the spin column, followed by centrifuge at 10,000*g* for 30 sec, and then repeated once. The Spin column was then centrifuged for 1 min at 10,000*g* to completely remove the washing buffer, and the DNA on the column was eluted by 100 µl of Elution buffer (added to Spin column, followed by incubation at room temperature for 5 min, and then centrifuged at 12,000*g* for 1 min). Total DNA was quantified with Maxima SYBR Green/ROX qPCR Master Mix on a LightCycler 480 II System (Roche) with following programmes: 70 ng of DNA was pre-denatured at 95 °C for 10 min, and then subjected to PCR for a total of 45 cycles: denaturing at 95 °C for 10 s, annealing and extending at 65 °C for 30 s in each cycle. Primer pairs used for qPCR are as previously described^145^ (mouse *Hk2*, 5’-GCCAGCCTCTCCTGATTTTAGTGT-3’ and 5’-GGGAACACAAAAGACCTCTTCTGG-3’; mouse *Nd1*, 5’-CTAGCAGAAACAAACCGGGC-3’ and 5’-CCGGCTGCGTATTCTACGTT-3’).

Nematode mitochondrial DNA copy numbers were determined from worm lysates as described previously^100^. Briefly, 30 synchronised early L4 worms were collected, and were lysed with 10 μl of Single Worm lysis buffer. The worm lysate was frozen at −80 [overnight, followed by incubating at 65 [for 1 h and 95 [for 15 min. Nematode DNA was then quantified with Maxima SYBR Green/ROX qPCR Master Mix on a LightCycler 480 II System (Roche) with following programmes: pre-denaturing at 95 °C for 10 min and then for a total of 45 cycles of denaturing at 95 °C for 10 s, and annealing and extending at 65 °C for 30 s in each cycle. Primer pairs used for qPCR are designed as described previously^118^ (*C*. *elegans nd-1*, 5’-AGCGTCATTTATTGGGAAGAAGAC-3’ and 5’-AAGCTTGTGCTAATCCCATAAATGT-3’; *C. elegans act-3*, 5’-TGCGACATTGATATCCGTAAGG-3’ and 5’-GGTGGTTCCTCCGGAAAGAA-3’).

*Drosophila* DNA copy numbers were determined as described previously^146^, with minor modifications. Briefly, some 20 anaesthetised adults were homogenised in 100 μl of Fly Lysis Buffer (75 mM NaCl, 25 mM EDTA, 25 mM HEPES, pH7.5) containing proteinase K (100 μg/ml). The homogenate was then frozen at −80 [for 12 h, followed by incubating at 65 [for 1 h and 95 [for another 15 min. The fly DNA was then quantified using the Maxima SYBR Green/ROX qPCR Master Mix on a LightCycler 480 II System (Roche) with following programmes: pre-denaturing at 95 °C for 5 min and then for a total of 40 cycles of denaturing at 95 °C for 10 s, and annealing 60 [for 20 s and extending at 72 °C for 20 s in each cycle. Primer pairs used for qPCR are as previously described^146^ (*D. melanogaster 16S rRNA*, 5’-TCGTCCAACCATTCATTCCA-3’ and 5’-TGGCCGCAGTATTTTGACTG-3’; *D. melanogaster RpL32*, 5’-AGGCCCAAGATCGTGAAGAA-3’ and 5’-TGTGCACCAGGAACTTCTTGAA-3’).

### Measurement of adenylates and NAD^+^

ATP, ADP, AMP, and NAD^+^ from cells, tissues or flies were analysed by CE-MS as described previously^17^, with minor modifications^147^. Briefly, each measurement required MEFs collected from one 10-cm dish (60-70% confluence) 100 mg of liver or muscle tissue dissected by freeze clamp, or 20 anesthetised adult flies were used. Before CE-MS analysis, cells were rinsed with 20 ml of 5% (m/v) mannitol solution (dissolved in water) and instantly frozen in liquid nitrogen. Cells were then lysed with 1 ml of methanol containing IS1 (50 µM L-methionine sulfone, 50 µM D-campher-10-sulfonic acid, dissolved in water; 1:500 (v/v) added to the methanol and used to standardise the metabolite intensity and to adjust the migration time), and were scraped from the dish. For analysis of metabolites in liver and muscle, mice were anaesthetised after indicated treatments. The tissue was then quickly excised by freeze-clamping, and then ground in 1 ml of methanol with IS1. For analysis of metabolites in flies, some 20 adult flies were anesthetised, followed by grounding in 1 ml of methanol with IS1 after freezing by liquid nitrogen. The lysate was then mixed with 1 ml of chloroform and 400 μl of water by 20 s of vortexing. After centrifugation at 15,000*g* for 15 min at 4 °C, 450 μl of aqueous phase was collected and was then filtrated through a 5-kDa cutoff filter (cat. OD003C34, PALL) by centrifuging at 12,000*g* for 3 h at 4 °C. In parallel, quality control samples were prepared by combining 10 μl of the aqueous phase from each sample and then filtered alongside the samples. The filtered aqueous phase was then freeze-dried in a vacuum concentrator at 4 °C, and then dissolved in 100 μl of water containing IS2 (50 µM 3-aminopyrrolidine dihydrochloride, 50 µM N,N-diethyl-2-phenylacetamide, 50 µM trimesic acid, 50 µM 2-naphtol-3,6-disulfonic acid disodium salt, dissolved in methanol; used to adjust the migration time). A total of 20 μl of re-dissolved solution was then loaded into an injection vial (cat. 9301-0978, Agilent; equipped with a snap cap (cat. 5042-6491, Agilent)). Before CE-MS analysis, the fused-silica capillary (cat. TSP050375, i.d. 50 µm × 80 cm; Polymicro Technologies) was installed in a CE/MS cassette (cat. G1603A, Agilent) on the CE system (Agilent Technologies 7100). The capillary was then pre-conditioned with Conditioning Buffer (25 mM ammonium acetate, 75 mM diammonium hydrogen phosphate, pH 8.5) for 30 min, followed by balanced with Running Buffer (50 mM ammonium acetate, pH 8.5; freshly prepared) for another 1 h. CE-MS analysis was run at anion mode, during which the capillary was washed by Conditioning Buffer, followed by injection of the samples at a pressure of 50 mbar for 25 s, and then separation with a constant voltage at −30 kV for another 40 min. Sheath Liquid (0.1 μM hexakis(1H, 1H, 3H-tetrafluoropropoxy)phosphazine, 10 μM ammonium trifluoroacetate, dissolved in methanol/water (50% v/v); freshly prepared) was flowed at 1 ml/min through a 1:100 flow splitter (Agilent Technologies 1260 Infinity II; actual flow rate to the MS: 10 μl/min) throughout each run. The parameters of MS (Agilent Technologies 6545) were set as: a) ion source: Dual AJS ESI; b) polarity: negative; c) nozzle voltage: 2,000 V; d) fragmentor voltage: 110 V; e) skimmer voltage: 50 V; f) OCT RFV: 500 V; g) drying gas (N_2_) flow rate: 7 L/min; h) drying gas (N_2_) temperature: 300 °C; i) nebulizer gas pressure: 8 psig; j) sheath gas temperature: 125 °C; k) sheath gas (N_2_) flow rate: 4 L/min; l) capillary voltage (applied onto the sprayer): 3,500 V; m) reference (lock) masses: m/z 1,033.988109 for hexakis(1H, 1H, 3H-tetrafluoropropoxy)phosphazine, and m/z 112.985587 for trifluoroacetic acid; n) scanning range: 50-1,100 m/z; and o) scanning rate: 1.5 spectra/s. Data were collected using MassHunter LC/MS acquisition 10.1.48 (Agilent Technologies), and were processed using Qualitative Analysis B.06.00 (Agilent Technologies). Levels of AMP, ADP, ATP and NAD^+^ were measured using full scan mode with m/z values of 346.0558, 426.0221, 505.9885 and 662.1019. Note that a portion of ADP and ATP could lose one phosphate group during in-source fragmentation, thus leaving the same m/z ratios as AMP and ADP, and should be corrected according to their different retention times in the capillary. Therefore, the total amount of ADP is the sum of the latter peak of the m/z 346.0558 spectrogramme and the former peak of the m/z 426.0221 spectrogramme, and the same is applied for ATP. Note that the retention time of each metabolite may vary between each run, and can be adjusted by isotope-labelled standards (dissolved in individual cell or tissue lysates) run between each samples, so do IS1 and IS2.

Levels of ATP, ADP, AMP and NAD^+^ in nematodes were analysed using the HPLC-MS as described previously^100^. Briefly, some 150 nematodes maintained on NGM (containing LCA or not) for 2 days were washed with ice-cold M9 buffer containing Triton X-100, followed by removing bacteria by quickly centrifuging the slurry at 100*g* for 5 s, and then instantly lysed in 1 ml of methanol. The lysates were then mixed with 1 ml of chloroform and 400 µl of water (containing 4 µg/ml [U-^13^C]-glutamine), followed with 20 s of vortexing. After centrifugation at 15,000*g* for another 15 min at 4 °C, 800 µl of aqueous phase was collected, lyophilised in a vacuum concentrator at 4 °C, and then dissolved in 30 µl of 50% (v/v, in water) acetonitrile. Some 30 µl of supernatant was then loaded into an injection vial (cat. 5182-0714, Agilent Technologies; with an insert (cat. HM-1270, Zhejiang Hamag Technology)) equipped with a snap cap (cat. HM-2076, Zhejiang Hamag Technology). Measurements of adenylate and NAD^+^ levels were based on ref. ^148^ using a QTRAP MS (QTRAP 5500, SCIEX) interfaced with a UPLC system (ExionLC AD, SCIEX). Some 2 µl of each sample were loaded onto a HILIC column (ZIC-pHILIC, 5 μm, 2.1 × 100 mm, PN: 1.50462.0001, Millipore). The mobile phase consisted of 15 mM ammonium acetate containing 3 ml/l ammonium hydroxide (>28%, v/v) in the LC-MS grade water (mobile phase A) and LC-MS grade 90% (v/v) acetonitrile in LC-MS grade water (mobile phase B) run at a flow rate of 0.2 ml/min. Metabolites were separated with the following HPLC gradient elution programme: 95% B held for 2 min, then to 45% B in 13 min, held for 3 min, and then back to 95% B for 4 min. The MS was run on a Turbo V ion source in negative mode with a spray voltage of −4,500 V, source temperature at 550 °C, Gas No.1 at 50 psi, Gas No.2 at 55 psi, and curtain gas at 40 psi. Metabolites were measured using the multiple reactions monitoring mode (MRM), and declustering potentials and collision energies were optimised through using analytical standards. The following transitions were used for monitoring each compound: 505.9/158.9 and 505.9/408.0 for ATP; 425.9/133.9, 425.9/158.8, 425.9/328.0 for ADP; 345.9/79.9, 345.9/96.9, 345.9/133.9 for AMP, 662.0/540.1 for NAD^+^, and 149.9/114 for [U-^13^C]-glutamine. Data were collected using Analyst software (v.1.7.1, SCIEX), and the relative amounts of metabolites were analysed using MultiQuant software (v.3.0.3, SCIEX). Similar to the CE-MS analysis, a portion of ADP and ATP could lose one or two phosphate groups during in-source-fragmentation thus leaving same m/z ratios as AMP and ADP, which was corrected according to their different retention times in column.

### Reagents

Rabbit polyclonal antibody against acetylated V1E1-K99 (Ac-V1E1-K99; 1:1,000 dilution for immunoblotting (IB)) was raised using the peptide CARDDLITDLLNEA(AcK) of human V1E1 conjugated to the KLH immunogen (linked to the cysteine residue). A rabbit was then biweekly immunised with 300 µg of KLH-conjugated antigen, which is pre-incubated with 1.5 mg manganese adjuvant (kindly provided by Dr. Zhengfan Jiang from Peking University, see ref. ^149^) for 5 min and then mixed with PBS to a total volume of 1.5 ml, for 4 times, followed by collecting antiserum. The Ac-V1E1-K99 antibody was then purified from the antiserum using the CARDDLITDLLNEA(AcK) peptide-conjugated SulfoLink Coupling resin/column supplied in the SulfoLink Immobilization Kit. To prepare the column, 1 mg of peptide was first dissolved with 2 ml of Coupling Buffer, followed by addition of 0.1 ml of TCEP (25 mM stock concentration) and then incubation at room temperature for 30 min. The mixture was then incubated with SulfoLink Resin in a column, which is pre-calibrated by 2 ml of Coupling Buffer for 2 times, on a rotator at room temperature for 15 min, followed by incubating at room temperature for 30 min without rotating. The excess peptide was then removed, and the resin was washed with 2 ml of Wash Solution for 3 times, followed by 2 ml of Coupling Buffer 2 times. The nonspecific-binding sites on the resin was then blocked by incubating with 2 ml of cysteine solution (by dissolving 15.8 mg of L-cysteine-HCl in 2 ml of Coupling Buffer to make a concentration of 50 mM cysteine) on a rotator for 15 min at room temperature, followed by incubating for another 30 min without rotating at room temperature. After removing the cysteine solution, the resin was washed with 6 ml of Binding/Wash Buffer, followed by incubating with 2 ml of antiserum mixed with 0.2 ml of Binding/Wash Buffer for 2 h on a rotator. The resin was then washed with 1 ml of Binding/Wash Buffer for 5 times, and the antibody was eluted with 2 ml of Elution Buffer. The eluent was then mixed with 100 μl of Neutralization Buffer.

The antibody against basal V1E1 exists in the crude antibody eluent was then removed through a previously described membrane-based affinity purification method^150^. Briefly, the bacterially purified, His-tagged V1E1 was subjected to SDS-PAGE, followed by transferring to a PVDF membrane (see details in “IP and IB assays” section). The V1E1-bound-membrane was incubated in 5% (w/v) non-fat milk dissolved in TBST (40 mM Tris, 275 μM NaCl, 0.2% (v/v) Tween-20, pH 7.6) for 2 h, then incubated with the crude antibody preparation for 2 days, and then repeated for another 2 times.

Rabbit anti-phospho-AMPK α-Thr172 (cat. #2535, RRID: AB_331250; 1:1,000 for immunoblotting (IB)), anti-AMPK α (cat. #2532, RRID: AB_330331; 1:1,000 for IB), anti-phospho-ACC-Ser79 (cat. #3661, RRID: AB_330337; 1:1,000 for IB), anti-ACC (cat. #3662, RRID: AB_2219400; 1:1,000 for IB), anti-LKB1 (cat. #3047, RRID: AB_2198327; 1:1,000 for IB), anti-His-tag (cat. #12698, RRID: AB_2744546; 1:1,000 for IB), anti-Myc-tag (cat. #2278, RRID: AB_490778; 1:120 for immunofluorescence (IF)), anti-AXIN1 (cat. #2074, RRID: AB_2062419; 1:1,000 for IB), anti-SIRT1 (cat. #9475, RRID: AB_2617130; 1:1,000 for IB), anti-SIRT2 (cat. #12650, RRID: AB_2716762; 1:1,000 for IB), anti-SIRT3 (cat. #5490, RRID: AB_10828246; 1:1,000 for IB), anti-SIRT5 (cat. #8782, RRID: AB_2716763; 1:1,000 for IB), anti-SIRT6 (cat. #12486, RRID: AB_2636969; 1:1,000 for IB), anti-SIRT7 (cat. #5360, RRID: AB_2716764; 1:1,000 for IB), anti-histone H3 (cat. #4499, RRID: AB_10544537; 1:1,000 for IB), anti-acetyl-histone H3-Lys9 (cat. #9649, RRID: AB_823528; 1:1,000 for IB), anti-LAMTOR1 (cat. #8975, RRID: AB_10860252; 1:1,000 for IB), anti-GAPDH (cat. #5174; RRID: AB_10622025; 1:1,000 for IB), and HRP-conjugated mouse anti-rabbit IgG (conformation-specific, cat. #5127, RRID: AB_10892860; 1:2,000 for IB) antibodies were purchased from Cell Signaling Technology. Mouse anti-HA-tag (cat. sc-7392, RRID: AB_2894930; 1:500 for IP or 1:120 for IF), goat anti-AXIN (cat. sc-8567, RRID: AB_22277891;: 100 for IP (immunoprecipitation) and 1:120 for IF) and mouse anti-goat IgG-HRP (cat. sc-2354, RRID: AB_628490; 1:2,000 for IB) antibodies were purchased from Santa Cruz Biotechnology. Mouse anti-total OXPHOS (cat. ab110413, RRID: AB_2629281; 1:5,000 for IB), rat anti-LAMP2 (cat. ab13524, RRID: AB_2134736; 1:120 for IF), rabbit anti-laminin (cat. ab11575, RRID: AB_298179; 1:200 for IF), anti-ATP6V1B2 (cat. ab73404, RRID: AB_1924799; 1:1,000 for IB), anti-PEN2 (cat. ab154830, 1:1,000 for IB), and goat anti-SIRT4 (cat. ab10140, RRID: AB_2188769; 1:1,000 for IB) antibodies were purchased from Abcam. Mouse anti-eMHC (cat. BF-G6, RRID: AB_10571455; 1:100 for IHC), anti-Pax7 (cat. Pax-7, RRID: AB_2299243; 1:100 for IHC), anti-MHCIIa (cat. SC71, RRID: AB_2147165; 1:100 for IHC), anti-MHCIIb (cat. BF-F3, RRID: AB_2266724; 1:100 for IHC), and anti-MHCI (cat. C6B12, RRID: AB_528351; 1:100 for IHC) antibodies were purchased from Developmental Studies Hybridoma Bank. Rabbit anti-tubulin (cat. 10068-1-AP, RRID: AB_2303998; 1:1,000 for IB nematode tubulin), anti-ATP6V1E1 (cat. 15280-1-AP, RRID: AB_2062545; 1:1,000 for IB), anti-TULP3 (cat. #13637-1-AP, RRID: AB_2211547, 1:20,000 for IB), and mouse anti-tubulin (cat. 66031-1-Ig, RRID: AB_11042766; 1:20,000 for IB mammalian tubulin), anti-HA-tag (cat. 66006-2-Ig, RRID: AB_2881490; 1:20,000 for IB) antibodies were purchased from Proteintech. Rabbit anti-ATP6v0c (cat. NBP1-59654, RRID: AB_11004830; 1:1,000 for IB) antibody was purchased from Novus Biologicals. Mouse anti β-ACTIN (cat. A5316, RRID: AB_476743; 1:1,000 for IB) was purchased from Sigma. Donkey anti-Goat IgG (H+L) Highly Cross-Adsorbed Secondary Antibody, Alexa Fluor Plus 488 (cat. A-32814, RRID: AB_2762838; 1:100 for IF), Donkey anti-Rat IgG (H+L) Highly Cross-Adsorbed Secondary Antibody, Alexa Fluor 594 (cat. A-21209, RRID: AB_2535795; 1:100 for IF), Goat anti-Mouse IgM (Heavy chain) Cross-Adsorbed Secondary Antibody, Alexa Fluor 488 (cat. A-21042, RRID: AB_2535711; 1:200 for IHC), Goat anti-Mouse IgG2b Cross-Adsorbed Secondary Antibody, Alexa Fluor 594 (cat. A-21145, RRID: AB_2535781; 1:200 for IHC), Goat anti-Mouse IgG1 Cross-Adsorbed Secondary Antibody, Alexa Fluor 647 (cat. A-21240, RRID: AB_2535809; 1:200 for IHC), Goat anti-Mouse IgG1 Cross-Adsorbed Secondary Antibody, Alexa Fluor 488 (cat. A-21121, RRID: AB_2535764; 1:200 for IHC), and Goat anti-Rabbit IgG (H+L) Cross-Adsorbed Secondary Antibody, Alexa Fluor 594 (cat. A-11012, RRID: AB_2534079; 1:200 for IHC) were purchased from Thermo. The horseradish peroxidase (HRP)-conjugated goat anti-mouse IgG (cat. 115-035-003, RRID: AB_10015289; 1:5,000 dilution for IB) and goat anti-rabbit IgG (cat. 111-035-003, RRID: AB_2313567; 1:5,000 dilution for IB) antibodies were purchased from Jackson ImmunoResearch.

DMSO (cat. D2650), LCA (cat. L6250), (2-hydroxypropyl)- β-cyclodextrin (cat. C0926), methanol (cat. 646377), ethanol (cat. 459836), chloroform (cat. C7559), PBS (cat. P5493), Triton X-100 (cat. T9284), sodium acetate (NaAc; cat. S5636), nuclease-free water (cat. W4502), human tubal fluid (HTF) medium (cat. MR-070-D), KSOM medium (cat. MR-121-D), L-glutathione reduced (GSH; cat. G4251), NaCl (cat. S7653), CaCl_2_ (cat. C5670), MgSO_4_ (cat. M2643), H_2_O_2_ (cat. H1009), KH_2_PO_4_ (cat. P5655), K_2_HPO_4_ (cat. P9666), streptomycin (cat. 85886), cholesterol (cat. C3045), agar (cat. A1296), propionic acid (cat. P5561), sucrose (cat. S7903), glucose (cat. G7021), 2-methylbutane (isopentane; cat. M32631), paraformaldehyde (cat. 158127), Canada balsam (cat. C1795), BSA (cat. A2153), glycerol (cat. G5516), Na_2_HPO_4_ (cat. S7907), sodium hypochlorite solution (NaClO; cat. 239305), NaOH (cat. S8045), Iron(II) sulfate heptahydrate (FeSO_4_; cat. F8633), HEPES (cat. H4034), EDTA (cat. E6758), EGTA (cat. E3889), MgCl_2_ (cat. M8266), KCl (cat. P9333), IGEPAL CA-630 (NP-40; cat. I3021), dithiothreitol (DTT; cat. 43815), IPTG (cat. I6758), carbenicillin (cat. C1613), D-mannitol (cat. M4125), glycine (cat. G8898), isopropanol (cat. 34863), diethylpyrocarbonate (DEPC)-treated water (cat. 693520), Trioxsalen (TMP; cat. T6137), methyl 4-hydroxybenzoate (methyl paraben; cat. H3647), mineral oil (cat. M5310), Halocarbon oil 700 (cat. H8898), Halocarbon oil 27 (cat. H8773), paraquat (cat. 36541), H_2_O_2_ (cat. H1009), agarose (cat. A9539), proteinase K (cat. P6556), L-methionine sulfone (cat. M0876), D-campher-10-sulfonic acid (cat. 1087520), acetonitrile (cat. 34888), ammonium acetate (cat. 73594), ammonium hydroxide solution (cat. 338818), 3-aminopyrrolidine dihydrochloride (cat. 404624), N,N-diethyl-2-phenylacetamide (cat. 384011), trimesic acid (cat. 482749), diammonium hydrogen phosphate (cat. 1012070500), ammonium trifluoroacetate (cat. 56865), formic acid (cat. 5.43804), diammonium hydrogen phosphate (cat. 1012070500), ammonium trifluoroacetate (cat. 56865), Tween-20 (cat. P9416), hexadimethrine bromide (polybrene; cat. H9268), octyl β-D-glucopyranoside (ODG; cat. O8001), Trizma base (Tris; cat. T1503), sodium pyrophosphate (cat. P8135), β-glycerophosphate (cat. 50020), SDS (cat. 436143), sodium deoxycholate (cat. S1827), β-mercaptoethanol (cat. M6250), formaldehyde solution (formalin; F8775), chloroform-*d* (with 0.05% (v/v) tetramethylsilane (TMS); cat. 612200), N,N-dimethylformamide (DMF; cat. 33120), N,N-diisopropylethylamine (DIEA; cat. 199818), dichloromethane (DCM; cat. 650463), Na_2_SO_4_ (cat. 746363), IPTG (cat. I6758), imidazole (cat. I5513), HIS-Select Nickel Affinity Gel (cat. P6611), Coomassie Brilliant Blue R-250 (cat. 1.12553), acetic acid (cat. 27225), TCEP (cat. C4706), TBTA (cat. 678937), CuSO_4_ (cat. C1297), trichloroacetic acid (cat. 91228), ammonium formate (cat. 70221), FCCP (cat. C2920), sodium azide (NaN_3_; cat. S2002), gentamycin (cat. 345814), collagenase A (cat. 11088793001) and oligomycin A (cat. 75351) were purchased from Sigma. TRIzol (cat. 15596018), Phusion High-Fidelity DNA Polymerase kit (cat. F530N), mMESSAGE mMACHINE T7 Transcription Kit (cat. AM1344), MEGAclear Transcription Clean-Up Kit (cat. AM1908), MEGAshortscript T7 Transcription Kit (cat. AM1354), Maxima SYBR Green/ROX qPCR master mix (cat. K0223), SulfoLink Immobilization Kit for Peptides (cat. 44999), DMEM, high glucose (DMEM; cat. 12800082), FBS (cat. 10099141C), penicillin-streptomycin (cat. 15140163), Lipofectamine 2000 (cat. 11668500), MEM non-essential amino acids solution (cat. 11140050), GlutaMAX (cat. 35050061), sodium pyruvate (cat. 11360070), ProLong Diamond antifade mountant (cat. P36970), ProLong Live Antifade reagent (cat. P36975), Streptavidin Magnetic Beads (cat. 88817; 1:100 for IP), Prestained Protein MW Marker (cat. 26612), and LysoSensor Green DND-189 (cat. L7535) were purchased from Thermo. MinElute PCR Purification Kit (cat. 28004) were purchased from Qiagen. Biospin Tissue Genomic DNA extraction Kit (cat. BSC04M1) was purchased from BioFlux. hCG (cat. 110900282) and PMSG (cat. 110904564) were purchased from Sansheng Biological Technology (Ningbo, China). WesternBright ECL and peroxide solutions (cat. 210414-73) were purchased from Advansta. Bacteriological peptone (cat. LP0037) and yeast extract (cat. LP0021) were purchased from Oxoid. Protease inhibitor cocktail (cat. 70221) was purchased from Roche. 3-hydroxynaphthalene-2,7-disulfonic acid disodium salt (2-naphtol-3,6-disulfonic acid disodium salt; cat. H949580) was purchased from Toronto Research Chemicals. Hexakis(1H,1H,3H-perfluoropropoxy)phosphazene (hexakis(1H, 1H, 3H-tetrafluoropropoxy)phosphazine; cat. sc-263379) was purchased from Santa Cruz Biotechnology. Polyethylenimine (PEI; cat. 23966) was purchased from Polysciences. Nonfat dry milk (cat. #9999) and normal goat serum (NGS; cat. #5425) were purchased from Cell Signaling Technology. 2-(3-(but-3-yn-1-yl)-3H-diazirin-3-yl)ethan-1-amine and 1H-benzotriazol-1-yloxytris(dimethylamino)phosphonium hexafluorophosphate (BOP) were purchased from Bidepharm. O.C.T. Compound (cat. 4583) was purchased from Sakura. ReverTra Ace qPCR RT Master Mix with gDNA Remover (cat. FSQ-301) was purchased from Toyobo. PrimeSTAR HS polymerase (cat. R40A) was purchased from Takara. Dry yeast (cat. FLY804020F) and cornmeal (cat. FLY801020) were purchased from LabScientific. Soy flour (cat. 62116) was purchased from Genesee Scientific. Light corn syrup was purchased from Karo. Grape juice was purchased from Welch’s. Cell Counting Kit-8 (CCK-8) was purchased from ApexBio. [U-^13^C]-glutamine (cat. 184161-19-1) was purchased from Cambridge Isotope Laboratories. rProtein A Sepharose Fast Flow (cat. 17127904), Protein G Sepharose 4 Fast Flow (cat. 17061806) and Superdex 200 Increase 10/300 GL (cat. 28990944) were purchased from Cytiva.

### Cell lines

In this study, no cell line used is on the list of known misidentified cell lines maintained by the International Cell Line Authentication Committee (https://iclac.org/databases/cross-contaminations/). HEK293T cells (cat. CRL-3216) were purchased from ATCC. *LAMTOR1*^F/F^, *AXIN*^F/F^ and *PKC*ζ^F/F^ MEFs were established by introducing SV40 T antigen using lentivirus into cultured primary embryonic cells from mouse litters. *LAMTOR1*^-/-^ MEFs were generated by infecting *LAMTOR1*^F/F^ MEFs with adenoviruses expressing the Cre recombinase for 12 h, as for *AXIN*^-/-^ MEFs and *PKC*ζ^-/-^ MEFs. The infected cells were then incubated in fresh DMEM for another 12 h before further treatments. The *ALDO*-TKD^17^, *TRPV*-QKO^19^, *PEN2*^-/-^ (ref. ^23^), *ATP6AP1*^-/-^ (ref. ^23^) and *AMPK*α*1*/*2*^-/-^ (ref. ^151^) MEFs, and si*ATP6v0c*^18^ HEK293T cells were generated and validated as previously described. HEK293T cells and MEFs were maintained in DMEM supplemented with 10% FBS, 100 IU penicillin, 100 mg/ml streptomycin at 37 °C in a humidified incubator containing 5% CO_2_. All cell lines were verified to be free of mycoplasma contamination. HEK293T cells were authenticated by STR sequencing. PEI at a final concentration of 10 μM was used to transfect HEK293T cells. Total DNA to be transfected for each plate was adjusted to the same amount by using relevant empty vector. Transfected cells were harvested at 24 h after transfection.

Lentiviruses, including those for knockdown or stable expression, were packaged in HEK293T cells by transfection using Lipofectamine 2000. At 30 h post transfection, medium (DMEM supplemented with 10% FBS and MEM non-essential amino acids; approximately 2 ml) was collected and centrifuged at 5,000*g* for 3 min at room temperature. The supernatant was mixed with 10 μg/ml polybrene, and was added to MEFs or HEK293T cells, followed by centrifuging at 3000*g* for 30 min at room temperature (spinfection). Cells were incubated for another 24 h (MEFs) or 12 h (HEK293T cells) before further treatments. Cell numbers were determined using the CCK-8 kit according to the manufacturer’s instruction. The sequence of siRNA used to knockdown mouse *TULP3* is: 5’-GCATCTTGAGTAGTGTGAACTATGA-3’.

The genes (*SIRT1* to *SIRT7*, *FXR*, *FXRb*, *PXR*, *VDR*, *CAR*, *LXRa*, *LXRb*, *S1PR2*, *TGR5*, *CHRM2*, *CHRM3*, *FAS*, *FPR1*, *YES1* and *TULP3*) were deleted from MEFs using the CRISPR-Cas9system. Nucleotides were annealed to their complements containing the cloning tag aaac, and inserted into the back-to-back BsmB I restriction sites of lentiCRISPRv2 vector. The sequence for each sgRNA is as follows: 5’-CGGTATCTATGCTCGCCTTG-3’ and 5’-CAAGGCGAGCATAGATACCG-3’ for *SIRT1*, 5’-AAGGACGGGGAACTTACACG-3’ and 5’-CGTGTAAGTTCCCCGTCCTT-3’ for *SIRT2*, 5’-AACATCGACGGGCTTGAGAG-3’ and 5’-CTCTCAAGCCCGTCGATGTT-3’ for *SIRT3*, 5’-GGCGGCACAAATAACCCCGA-3’ and 5’-TCGGGGTTATTTGTGCCGCC-3’ for *SIRT4*, 5’-CATTGACGAGTTGCATCGCA-3’ and 5’-TGCGATGCAACTCGTCAATG-3’ for *SIRT5*, 5’-CGAGGGCCGAGCATTCTCGA-3’ and 5’-TCGAGAATGCTCGGCCCTCG-3’ for *SIRT6*, 5’-CCGACTTCGACCCTGCAGCT-3’ and 5’-AGCTGCAGGGTCGAAGTCGG-3’ for *SIRT7*, 5’-ACCAGTCTTCCGGTTGTTGG-3’ and 5’-CCAACAACCGGAAGACTGGT-3’ for *FXR* (#1), 5’-CGAATGGCCGCGGCATCGGC-3’ and 5’-GCCGATGCCGCGGCCATTCGC-3’ for *FXR* (#2), 5’-ACCAGTCTTCCGGTTGTTGG-3’ and 5’-CCAACAACCGGAAGACTGGT-3’ for *FXRb* (#1), 5’-GGACTCGACGCTCGAGAATC-3’ and 5’-GATTCTCGAGCGTCGAGTCC-3’ for *FXRb* (#2), 5’-TGAAACGCAATGTCCGGCTG-3’ and 5’-CAGCCGGACATTGCGTTTCA-3’ for *PXR* (#1), 5’-GATCATGTCCGATGCCGCTG-3’ and 5’-CAGCGGCATCGGACATGATC-3’ for *PXR* (#2), 5’-ACTTTGACCGGAATGTGCCT-3’ and 5’-AGGCACATTCCGGTCAAAGT-3’ for *VDR* (#1), 5’-TGGAGATTGCCGCATCACCA-3’ and 5’-TGGTGATGCGGCAATCTCCA-3’ for *VDR* (#2), 5’-CGGCCCATATTCTTCTTCAC-3’ and 5’-GTGAAGAAGAATATGGGCCG-3’ for *CAR* (#1), 5’-GGGGCCCACACTCGCCATGT-3’ and 5’-ACATGGCGAGTGTGGGCCCC-3’ for *CAR* (#2), 5’-TTCCGCCGCAGTGTCATCAA-3’ and 5’-TTGATGACACTGCGGCGGAA-3’ for *LXRa*, 5’-GCCGGGCGCTATGCCTGTCG-3’ and 5’-CGACAGGCATAGCGCCCGGC-3’ for *LXRb* (#1), 5’-CATAGCGCCCGGCCCCACCG-3’ and 5’-CGGTGGGGCCGGGCGCTATG-3’ for *LXRb* (#2), 5’-CGTGCAGTGGTTTGCCCGAG-3’ and 5’-CTCGGGCAAACCACTGCACG-3’ for *S1PR2* (#1), 5’-AAGACGGTCACCATCGTACT-3’ and 5’-AGTACGATGGTGACCGTCTT-3’ for *S1PR2* (#2), 5’-TAGTGGTGGGCGACGCTCAT-3’ and 5’-ATGAGCGTCGCCCACCACTA-3’ for *TGR5* (#1), 5’-GGCTGCGCAAGTGGCGGTCC-3’ and 5’-GGACCGCCACTTGCGCAGCC-3’ for *TGR5* (#2), 5’-GCATGATGATTGCAGCTGCG-3’ and 5’-CGCAGCTGCAATCATCATGC-3’ for *CHRM2*, 5’-GCCGTGCCGAAGGTGATGGT-3’ and 5’-ACCATCACCTTCGGCACGGC-3’ for *CHRM3* (#1), 5’-AGCCGGTGTGATGATTGGTC-3’ and 5’-GACCAATCATCACACCGGCT-3’ for *CHRM3* (#2), 5’-TCTCCGAGAGTTTAAAGCTG-3’ and 5’-CAGCTTTAAACTCTCGGAGA-3’ for *FAS* (#1), 5’-TGCTCAGAAGGATTATATCA-3’ and 5’-TGATATAATCCTTCTGAGCA-3’ for *FAS* (#2), 5’-AGAAGGTAATCATCGTACCC-3’ and 5’-GGGTACGATGATTACCTTCT-3’ for *FPR1* (#1), 5’-GGGCAACGGGCTCGTGATCT-3’ and 5’-AGATCACGAGCCCGTTGCCC-3’ for *FPR1* (#2), 5’-TCTAGTCGCAATGATTCTCG-3’ and 5’-CGAGAATCATTGCGACTAGA-3’ for *YES1*, 5’-CTCCCCGTCGGCTCGCTCAG-3’ and 5’-CTGAGCGAGCCGACGGGGAG-3’ for *TULP3* (#1), 5’-ACGTCGCTGCGAGGCATCTG-3’ and 5’-CAGATGCCTCGCAGCGACGT-3’ for *TULP3* (#2). The constructs were then subjected to lentivirus packaging using HEK293T cells that were transfected with 2 µg of DNA in Lipofectamine 2000 transfection reagent per well of a 6-well plate. At 30 h post transfection, the virus (approximately 2 ml) was collected and for infecting MEFs as described above, except cells cultured to 15% confluence were incubated with virus for 72 h. When cells were approaching to confluence, they were single-cell sorted into 96-well dishes. Clones were expanded and evaluated for knockout status by sequencing.

### Plasmids

Full-length cDNAs used in this study were obtained either by PCR using cDNA from MEFs, or by purchasing from Origene or Sino Biological. The dominant-negative mutants of SIRT1 to SIRT7, i.e., SIRT1-H363Y^152^, SIRT2-H150Y^153^, SIRT3-H248Y^80^, SIRT4-H161Y^154^, SIRT5-H158Y^155^, SIRT6-H133Y^38^ and SIRT7-H187Y^85^ were generated according to previous reports. Mutations of V1E1, SIRT1-7, and TULP3 were performed by PCR-based site-directed mutagenesis using PrimeSTAR HS polymerase. Expression plasmids for various epitope-tagged proteins were constructed in the pcDNA3.3 vector (#K830001, Thermo) for transfection (ectopical expression) in mammalian cells, in the pBOBI vector for lentivirus packaging (stable expression) in mammalian cells, in the in pLVX-IRES (for ALDOA; #631849, Takara) for doxycycline-inducible expression in mammalian cells, or in the pET-28a vector (#69864-3, Novagen) for bacterial expression. PCR products were verified by sequencing (Invitrogen, China). All expression plasmids constructed in this study have been deposited to Addgene (https://www.addgene.org/Sheng-cai_Lin/). The lentivirus-based vector pLV-H1-EF1a-puro (#SORT-B19, Biosettia) was used for expression of siRNA in MEFs. *Escherichia coli* strain DH5 α (cat. #PTA-1977) was purchased from ATCC and Stbl3 (cat. #C737303) from Thermo. All plasmids were amplified in *E*. *coli* strain DH5 α, except those mutagenesis in Stbl3. All plasmids used in this study were purified by CsCl density gradient ultracentrifugation method.

### IP and IB assays

For determining the AMPK-activating complex formation, endogenous AXIN was immunoprecipitated and analysed as described previously^18^. Briefly, four 15-cm dishes of MEFs (grown to 80% confluence) were collected for IP of AXIN. Cells were lysed with 750 μl/dish of ice cold ODG buffer (50 mM Tris-HCl, pH 8.0, 50 mM NaCl, 1 mM EDTA, 2% (w/v) ODG, 5 mM β-mercaptoethanol with protease inhibitor cocktail), followed by sonication and centrifugation at 4 °C for 15 min. Cell lysates were incubated with anti-AXIN antibody overnight. Overnight protein aggregates were pre-cleared by centrifugation at 20,000*g* for 10 min, and protein A/G beads (1:250, balanced with ODG buffer) were then added into the lysate/antibody mixture for another 3 h at 4 °C. The beads were centrifuged and washed with 100 times volume of ODG buffer for 3 times (by centrifuging at 2,000*g*) at 4 °C and then mixed with an equal volume of 2× SDS sample buffer and boiled for 10 min before immunoblotting.

To determine the interaction between ectopically expressed TULP3 and sirtuins, a 6 cm-dish of HEK293T cells was transfected with different expression plasmids. At 24 h after transfection, cells were collected and lysed in 500 µl of ice-cold Triton lysis buffer (20 mM Tris-HCl, pH 7.5, 150 mM NaCl, 1 mM EDTA, 1 mM EGTA, 1% (v/v) Triton X-100, 2.5 mM sodium pyrophosphate, 1 mM β-glycerophosphate, with protease inhibitor cocktail), followed by sonication and centrifugation at 4 °C for 15 min. Anti-HA-tag (1:100) or anti-Myc-tag (1:100) antibodies, along with protein A/G beads (1:100, pre-balanced in Triton lysis buffer) was added into the supernatant and mixed for 4 h at 4 °C. The beads were washed with 200 times volume of ice-cold Triton lysis buffer wash buffer for 3 times at 4 °C and then mixed with an equal volume of 2× SDS sample buffer and boiled for 10 min before immunoblotting.

To analyse the levels of p-AMPK α, p-ACC and Ac-V1E1-K99 in HEK293T cells and MEFs, cells grown to 70-80% confluence in a well of a 6-well dish were lysed with 250 μl of ice-cold Triton lysis buffer. The lysates were then centrifuged at 20,000*g* for 10 min at 4 °C and an equal volume of 2× SDS sample buffer was added into the supernatant. Samples were then boiled for 10 min and then directly subjected to immunoblotting. To analyse the levels of p-AMPK α, p-ACC and Ac-V1E1-K99 in muscle and liver tissues, mice were anesthetised after indicated treatments. Freeze-clamped tissues were immediately lysed with ice-cold Triton lysis buffer (10 μl/mg tissue weight for liver, and 5 μl/mg tissue weight for muscle), followed by homogenisation and centrifugation as described above. The lysates were then mixed with 2× SDS sample buffer, boiled, and subjected to immunoblotting. To analyse the levels of p-AMPK α, p-ACC in flies, 20 adults or third instar larvae were lysed with 200 μl ice-cold RIPA buffer (50 mM Tris-HCl, pH 7.5, 150 mM NaCl, 1% NP-40, 0.5% sodium deoxycholate, with protease inhibitor cocktail) containing 0.1% SDS, followed by homogenisation and centrifugation as described above. The lysates were then mixed with 5× SDS sample buffer, boiled, and subjected to immunoblotting. To analyse the levels of p-AMPK α and p-ACC in nematodes, some 150 nematodes cultured on the NGM plate were collected for each sample. Worms were quickly washed with ice-cold M9 buffer containing Triton X-100, and were lysed with 150 μl of ice-cold lysis buffer. The lysates were then mixed with 5× SDS sample buffer, followed by homogenisation and centrifugation as described above, and then boiled before being subjected to IB. All samples were subjected to IB on the same day of preparation, and any freeze–thaw cycles were avoided.

For IB, the SDS-polyacrylamide gels were prepared in house as described previously^23^. The thickness of gels used in this study was 1.0 mm. Samples of less than 10 μl were loaded into wells, and the electrophoresis was run at 100 V (by PowerPac HC High-Current Power Supply, Bio-Rad) in a Mini-PROTEAN Tetra Electrophoresis Cell (Bio-Rad). In this study, all samples were resolved on 8% resolving gels, except those for H3, H3-Ac-K9, LAMTOR1, OXPHOS proteins and ATP6v0c, which were on 15% gels (prepared as those for 8%, except that a final concentration of 15% Acryl/Bis was added to the resolving gel solution), and β-ACTIN, GAPDH, ALDOA, V1E1 and V1E1-Ac-K99, which were on 10% gels.

The resolved proteins were then transferred to the PVDF membrane (0.45 μm, cat. IPVH00010, Merck) as described previously^23^. The PVDF membrane was then blocked by 5% (w/v) BSA (for all antibodies against phosphorylated proteins) or 5% (w/v) non-fat milk (for all antibodies against total proteins) dissolved in TBST for 2 h on an orbital shaker at 60 r.p.m. at room temperature, followed by rinsing with TBST for twice, 5 min each. The PVDF membrane was then incubated with desired primary antibody overnight at 4 °C on an orbital shaker at 60 r.p.m., followed by rinsing with TBST for three times, 5 min each at room temperature, and then the secondary antibodies for 3 h at room temperature with gentle shaking. The secondary antibody was then removed, and the PVDF membrane was further washed with TBST for 3 times, 5 min each at room temperature. PVDF membrane was incubated in ECL mixture (by mixing equal volumes of ECL solution and Peroxide solution for 5 min), then life with Medical X-Ray Film (FUJIFILM). The films were then developed with X-OMAT MX Developer (Carestream), and X-OMAT MX Fixer and Replenisher solutions (Carestream) on a Medical X-Ray Processor (Carestream) using Developer (Model 002, Carestream). The developed films were scanned using a Perfection V850 Pro scanner (Epson) with an Epson Scan software (v.3.9.3.4), and were cropped using Photoshop 2023 software (Adobe). Levels of total proteins and phosphorylated proteins were analysed on separate gels, and representative immunoblots are shown. Uncropped immunoblots are shown in Supplementary Fig. 1. The band intensities on developed films were quantified using ImageJ software (v.1.8.0, National Institutes of Health Freeware).

### Confocal microscopy

For determining the lysosomal localisation of AXIN, cells grown to 80% confluence on coverslips in 6-well dishes were fixed for 20 min with 4% (v/v) formaldehyde in PBS at room temperature. The coverslips were rinsed twice with PBS and permeabilised with 0.1% (v/v) Triton X-100 in PBS for 5 min at 4 °C. After rinsing twice with PBS, the sections were blocked with PBS containing 5% BSA for 30 min at room temperature. Then the coverslips were incubated with anti-AXIN and anti-LAMP2 antibodies (both at 1:100, diluted in PBS) overnight at 4 °C. The cells were then rinsed 3 times with 1 ml of PBS, and then incubated with secondary antibody for 8 h at room temperature in the dark. Cells were washed for another 4 times with 1 ml of PBS, and then mounted on slides using ProLong Diamond antifade mountant. Confocal microscopy images were taken using a STELLARIS 8 FALCON (Leica) systems equipped with HyD SMD detectors and an HC PL APO CS2 63x/1.40 OIL objective (Leica).

For detecting the pH of lysosomes, cells were grown on 35-mm glass-bottom dishes, and were cultured to 60-80% confluence. Cells were treated with 1 μM (final concentration) LysoSensor Green DND-189 for 30 min, then washed twice with PBS and incubated in fresh medium for another 30 min. In the meantime, ProLong Live antifade reagent, was added into the medium for staining the nucleus before taking images. During imaging, live cells were kept at 37 °C, 5% CO_2_ in a humidified incubation chamber (Zeiss, Incubator PM S1). Images were taken using a Zeiss LSM 980 with a ×63, 1.4 NA oil objective.

### Synthesis of LCA probe

All reagents and solvents were obtained from commercial suppliers and described in “Reagents” section. LCA probe were purified by preparative HPLC (Sail 1000, Welch Materials) equipped with an RD-C18 column (30.0 × 250 mm, 5 μm; Welch Materials). MS spectra for compound were recorded on 3100 Mass Detector (Waters).

High resolution MS spectra were obtained on Q-Exactive Orbitrap MS (Thermo). The purity of LCA probe was determined to be >95% by HPLC with UV detection at 220 nm (HD-C18 column, 5 μm, 4.6 × 250 mm, Agilent Technologies; the gradients were as follows: t = 0 min, 20% solvent B (methanol) and 80% solvent A (H_2_O); t = 10 min, 100% solvent B (methanol) and 0% solvent A (H_2_O); t = 30 min, 100% B (methanol) and 0% solvent A (H_2_O) with a constant flow rate at 1 ml/min). NMR spectra were recorded with a Bruker Avance III 600 MHz NMR spectrometer (^1^H: 600 MHz; ^13^C: 151 MHz), spectrometer instrument in Chloroform-*d*, with TMS as the internal standard. Chemical shift was reported in δ (ppm), multiplicities (s = singlet, d = doublet, t = triplet, q = quartet, p = pentet, dd = doublet of doublets, td = triplet of doublets, tt = triplet of triplets, and m = multiplet), integration and coupling constants (*J* in Hz). ^1^H and ^13^C chemical shift are relative to the solvent: δ_H_ 7.26 and δ_C_ 77.2 for Chloroform-*d*.

The biotin-N_3_ linker was synthesised as previously described^156^, and the LCA probe was synthesised through an amide condensation reaction between LCA and 2-(3-(but-3-yn-1-yl)-3H-diazirin-3-yl)ethan-1-amine. Briefly, LCA (55 mg, 0.146 mmol, 1.0 eq) was dissolved in 2 ml of DMF, followed by mixing with 97.3 mg of BOP (0.219 mmol, 1.5 eq). After 30 min of stirring at room temperature, some 24 mg of 2-(3-(but-3-yn-1-yl)-3H-diazirin-3-yl)ethan-1-amine (0.175 mmol, 1.2 eq) and 0.073 ml of DIEA (0.438 mmol, 3.0 eq) were added to the mixture, followed by stirring for another 12 h at room temperature. The mixture was then diluted with 20 ml of H_2_O, followed by extracting with 30 ml of DCM for three rounds. The organic phase obtained from the three rounds of extraction was pooled, dried by anhydrous Na_2_SO_4_, and then evaporated on a rotary evaporator (Rotavapor R-300, BUCHI) at 90 r.p.m., room temperature. The powder was then dissolved in 1 ml of DMSO, followed by purifying using the preparative HPLC with gradients as follows: t = 0 min, 50% solvent B (methanol) and 50% solvent A (H_2_O); t = 15 min, 80% solvent B (methanol) and 2% solvent A (H_2_O); t = 25 min, 100% B (methanol) and 0% solvent A (H_2_O); t = 45 min, 100% B (methanol) and 0% solvent A (H_2_O) with a constant flow rate at 20 ml/min. LCA probe was obtained as a white solid with yield of 80.7% (58.4 mg), and was dissolved in DMSO to a stock concentration of 50 mM. The solution was stored at 4 °C for no more than 1 month.

LCA probe: ^1^H NMR (600 MHz, Chloroform-*d*) δ 5.71 (t, J = 5.9 Hz, 1H), 3.62 (tt, J = 10.6, 4.6 Hz, 1H), 3.10 (q, J = 6.5 Hz, 2H), 2.24 (ddd, J = 14.5, 10.6, 5.1 Hz, 1H), 2.10 – 2.05 (m, 1H), 2.04 – 2.01 (m, 3H), 1.98 – 1.94 (m, 1H), 1.89 – 1.77 (m, 5H), 1.77 – 1.72 (m, 1H), 1.69 (t, J = 6.7 Hz, 2H), 1.67 – 1.63 (m, 3H), 1.60 – 1.54 (m, 1H), 1.53 – 1.48 (m, 1H), 1.44 – 1.35 (m, 6H), 1.35 – 1.29 (m, 2H), 1.28 – 1.19 (m, 3H), 1.17 – 1.12 (m, 1H), 1.09 (dd, J = 11.5, 7.8 Hz, 2H), 1.07 – 1.03 (m, 2H), 0.97 (td, J = 14.3, 3.5 Hz, 1H), 0.93 (d, J = 6.8 Hz, 3H), 0.92 (s, 3H), 0.64 (s, 3H); ^13^C NMR (151 MHz, Chloroform-*d*) δ 173.7, 82.7, 71.8, 69.4, 56.5, 56.0, 42.8, 42.1, 40.5, 40.2, 36.5, 35.9, 35.5, 35.4, 34.6, 34.3, 33.6, 32.6, 32.2, 31.7, 30.6, 28.2, 27.2, 26.9, 26.4, 24.2, 23.4, 20.8, 18.4, 13.2, 12.1. MS (ESI) m/z: 496 [M + H]^+^. HRMS (ESI): [M + H]^+^ calculated for C_31_H_50_O_2_N_3_, 496.3898; found, 496.3910. HPLC analysis: retention time = 14.55 min; peak area, 98.02 % (λ = 220 nm).

### Protein expression

The cDNAs encoding human SIRT1 and TULP3 were inserted into the pET-28a vector for expressing His-tagged recombinant proteins. The pET-28a plasmids were transformed into the *E*. *coli* strain BL21 (DE3) (cat. EC0114, Thermo), followed by culturing in LB medium in a shaker at 200 r.p.m. at 37 °C. The cultures of transformed cells were induced with 0.1 mM IPTG at an OD_600_ of 1.0. After incubating for another 12 h at 160 r.p.m. at 16 °C, the cells were collected. Cells were then homogenised in a His-binding Buffer (50 mM sodium phosphate, pH 7.0, 150 mM NaCl, 1% Triton X-100, 5% glycerol, and 10 mM imidazole), The homogenates were then sonicated on ice, and were subjected to ultracentrifugation at 150,000*g* for 30 min at 4 °C, followed by incubating with Nickel Affinity Gel (pre-balanced with His-binding buffer). The Nickel Affinity Gel was then washed with 100 times the volume of ice-cold His-washing Buffer (50 mM sodium phosphate, pH 7.0, 150 mM NaCl, and 20 mM imidazole), and the proteins were eluted from the resin by His-elution Buffer (50 mM sodium phosphate, pH 7.0, 150 mM NaCl, and 250 mM imidazole) at 4 °C. Proteins were then concentrated to approximately 3 mg/ml by ultrafiltration (Millipore, UFC905096) at 4 °C, then purified by gel filtration on a Superdex 200 column (Cytiva) balanced with the Superdex Buffer containing 50 mM Tris-HCl, pH 7.0 and 300 mM NaCl.

For determining the interaction between TULP3 and SIRT1 in vitro, the purified, His-tagged TULP3 and SIRT1, 4 μg of each, were pre-incubated in 100 μl (final volume) of Superdex Buffer on ice for 30 min. The mixture was then subjected to the Superdex 200 column balanced with the Superdex Buffer. The fraction size was set at 1 ml, with the mobile phase Superdex Buffer and a flow rate 0.5 ml/min. Samples were mixed with 5× SDS sample buffer, boiled, and subjected to SDS-PAGE followed by Coomassie Brilliant Blue staining (see details in the “Determination of the binding sites of TULP3 for LCA” section) or immunoblotting.

### Determination of the binding affinity of LCA to TULP3

To determining the binding affinity of LCA for TULP3 or SIRT1 by the affinity pull-down assay, the Streptavidin Magnetic Beads bound with biotinylated LCA probe were first prepared (described in ref. ^23^, with minor modifications). In brief, some 10 μM LCA probe was dissolved in 100 μl of ice-cold Triton lysis buffer at 4 °C, followed by mixing with 1 mM TCEP, 0.1 mM TBTA, 1 mM CuSO_4_, and 1 mM biotin-N_3_ linker (all final concentrations) at 4 °C for another 1 h. After centrifuge at 20,000*g*, 4 °C for 30 min, the supernatant was incubated with 10 μl of Streptavidin Magnetic Beads at 4 °C for 30 min, followed by washing with 100× volume of ice-cold Triton lysis buffer for 3 times. The biotinylated LCA probe-bound beads were then incubated with 3 μg of His-tagged TULP3, SIRT1, or TULP3-SIRT1 complex (prepared as described in the “Protein expression” section) in 100 μl of ice-cold Triton lysis buffer at 4 °C for 2 h, followed by washing with 100× volume of ice-cold Triton lysis buffer for 3 times, and then incubating in 100 μl of ice-cold Triton lysis buffer containing different concentrations of LCA at 4 °C for 0.5 h. The supernatant was then mixed with an equal volume of 2× SDS sample buffer, followed by immunoblotting to determine the amount of TULP3 or SIRT1 inside.

### Determination of the binding sites of TULP3 for LCA

The binding sites of TULP3 for LCA were determined through a two-step approach using MS combined in silico docking assays, as described previously^23^. Briefly, 10 μM LCA probe was incubated with 3 μg of His-tagged TULP3 (purified as described in the “Protein expression” section) in 100 μl of ice-cold Triton lysis buffer at 4 °C for 2 h, and then exposed to 365-nm wavelength UV (CX-2000, UVP) for 10 min. The mixture was then adjusted to final concentrations of 1 mM TCEP, 0.1 mM TBTA, 1 mM CuSO_4_ and 1 mM biotin-N_3_ linker, and were incubated at 4 °C for another 1 h. Protein aggregates were cleared by centrifugation at 20,000*g* for 15 [min, and 10 μl Streptavidin Magnetic Beads were then added to the supernatant for 2 h with gentle rotation. Beads were then washed with 100× volume of Triton lysis buffer for 3 times at 4 [°C, followed by incubating in 100 μl of ice-cold Triton lysis buffer containing different concentrations of LCA at 4 °C for 0.5 h. The supernatant was then mixed with an equal volume of 2× SDS sample buffer, followed by subjected to SDS-PAGE. After staining with Coomassie Brilliant Blue R-250 dye (5% (m/v) dissolved in 45% (v/v) methanol and 5% (v/v) acetic acid in water), gels were decoloured (in 45% (v/v) methanol and 5% (v/v) acetic acid in water) and the excised gel segments were subjected to in-gel chymotrypsin digestion, and then dried. Samples were analysed on a nanoElute (Bruker) coupled to a timsTOF Pro (Bruker) equipped with a CaptiveSpray source. Peptides were dissolved in 10 μl of 0.1% formic acid (v/v) and were loaded onto a homemade C18 column (35 cm × 75 μm, ID of 1.9 μm, 100 Å). Samples were then eluted with linear gradients of 3-35% acetonitrile (v/v, in 0.1% formic acid) at a flow rate of 0.3 μl/min, for 60 min. MS data were acquired with a timsTOF Pro MS (Bruker) operated in PASEF mode, and were analysed using Peaks Studio Xpro software (PEAKS Studio 10.6 build 20201221, Bioinformatics Solutions). The human UniProt Reference Proteome database was used for data analysis, during which the parameters were set as: a) precursor and fragment mass tolerances: 20 ppm and 0.05 Da; b) semi-specific digest mode: allowed; c) maximal missed cleavages per peptide: 3; d) variable modifications: oxidation of methionine, acetylation of protein N-termini, and phosphorylation of serine, threonine and tyrosine; e) fixed modification: carbamidomethylation of cysteine, and 467.3763 for LCA modification.

According to the MS results, the cleft comprising amino acids S194 and K333 might be the binding site of TULP3 for LCA. The in silico docking assay was then performed with the AutoDock vina software^157^, during which the structure of LCA and the AlphaFold-predicted TULP3 structure (https://alphafold.ebi.ac.uk/entry/O75386)^158,159^ were used. Data were then illustrated using the PyMOL (ver. 2.5, Schrodinger) software. The amino acid residues Y193, P195, K333 and P336 of TULP3 were then mutated (all to glycine) to generate the TULP3-4G mutant.

### Determination of SIRT1 activity

The deacetylase activity of SIRT1 was determined using an HPLC-MS-based method as described previously^36^, with some modifications. Briefly, some 4 μg of His-tagged SIRT1 (purified as described in the “Protein expression” section), either co-eluted with 4 μg of TULP3, or bound on 5 μl of Nickel Affinity Gel and incubated with lysates collected from two 10-cm dishes of MEFs (at 60-70% confluence, lysed in 1 ml of Triton lysis buffer as for immunoblotting) at 4 °C for 1 h and then washed with 1 ml of Triton lysis buffer twice, was incubated in 45 μl of Reaction Buffer containing 50 mM Tris-HCl, pH 9.0, 4 mM MgCl_2_, 0.2 mM DTT, 5 μM LCA (for LCA-treatment group only), and NAD^+^ at desired concentrations at 25 °C. The reaction was initiated by adding 5 μl of acetylated histone H3 peptide (QTAR(AcK)STGG) or acetylated V1E1 peptide (LNEA(AcK)QRLS), both dissolved in the Reaction Buffer to a stock concentration of 2 μg/ μl to the mixture, followed by incubating at 25 °C for another 15 min. Some 25 μl of 100% trichloroacetic acid and 50 μl of distilled water were then added to the mixture, and the mixture was then incubated at −20 °C for 12 h, followed by centrifuge at 18,000*g* at 4 °C for 30 min. The pellets were then dissolved with 70 μl of 10% (v/v, in water) acetonitrile, followed by votexing for 30s, and then centrifuge at 18,000*g* at 4 °C for another 10 min. Some 30 μl of supernatant was loaded into an injection vial (cat. 5182-0714, Agilent Technologies; with an insert (cat. HM-1270, Zhejiang Hamag Technology)) equipped with a snap cap (cat. HM-2076, Zhejiang Hamag Technology), and 4 μl of supernatant was injected into an HILIC column (HILIC Silica 3 μm, 2.1 × 150 mm, SN: 186002015; Atlantis) on a 1290 Infinity II LC System (Agilent Technologies) which is interfaced with a 6545 MS (Agilent Technologies). The mobile phase consisted of 10 mM ammonium formate containing 0.1% (v/v) formic acid in the LC-MS-grade water (mobile phase A) and LC-MS-grade acetonitrile containing 0.1% (v/v) formic acid (mobile phase B), and run at a flow rate of 0.3 ml/min. The HPLC gradient was as follows: 70%B for 1 min, then to 30% B at the 12^th^ min, hold for 3 min, and then back to 70% B at the 15.4^th^ min, hold for another 2 min. The parameters of 6545 MS (Agilent Technologies) were set as: a) ion source: Dual AJS ESI; b) polarity: positive; c) nozzle voltage: 500 V; d) fragmentor voltage: 175 V; e) skimmer voltage: 65 V; f) VCap: 500 V; g) drying gas (N_2_) flow rate: 8 l/min; h) drying gas (N_2_) temperature: 280 °C; i) nebulizer gas pressure: 35 psig; j) sheath gas temperature: 125 °C; k) sheath gas (N_2_) flow rate: 11 l/min; l) scanning range: 50-1,100 m/z. The deacetylated:acetylated peptide ratios of histone H3 and V1E1 were used to determine the activities of SIRT1 towards these substrates. The peak areas of each kind of peptides were calculated using following m/z values: a) acetylated histone H3: 453.2403 + 905.4805; b) deacetylated histone H3: 453.2403 + 905.4805; c) acetylated V1E1: 550.8071 + 1100.6064; and d) deacetylated V1E1: 529.8019 + 1058.5959.

### Protein and peptide MS

To determine the modification(s) of v-ATPase, the 21 HA-tagged subunits of v-ATPase, including ATP6V1A, ATP6V1B2, ATP6V1C1, ATP6V1C2, ATP6V1D, ATP6V1E1, ATP6V1E2, ATP6V1F, ATP6V1G1, ATP6V1G2, ATP6V1G3, ATP6V1H, ATP6V0A4, ATP6V0A2, ATP6V0B, ATP6V0C, ATP6V0D1, ATP6V0D2, ATP6V0E1, ATP6AP1 and ATP6AP2, were individually expressed in HEK293T cells. The immunoprecipitants (immunoprecipitated from 25 10-cm dishes of HEK293T cells ectopically expressing a certain subunit) were subjected to SDS-PAGE, and were processed as described in the “Determination of the binding sites of TULP3 for LCA” section. Samples were analysed on a nanoElute (Bruker) coupled to a timsTOF Pro (Bruker) equipped with a CaptiveSpray source. Peptides were dissolved in 10 μl of 0.1% formic acid (v/v) and were loaded onto a homemade C18 column (35 cm × 75 μm, ID of 1.9 μm, 100 Å). Samples were then eluted with linear gradients of 3-35% acetonitrile (v/v, in 0.1% formic acid) at a flow rate of 0.3 μl/min, for 60 min. MS data were acquired with a timsTOF Pro MS (Bruker) operated in PASEF mode, and were analysed using Peaks Studio Xpro (PEAKS Studio 10.6 build 20201221, Bioinformatics Solutions). The human UniProt Reference Proteome database was used for data analysis, during which the parameters were set as: a) precursor and fragment mass tolerances: 20 ppm and 0.05 Da; b) semi-specific digest mode: allowed; c) maximal missed cleavages per peptide: 3; d) variable modifications: oxidation of methionine, acetylation of protein N-termini, phosphorylation of serine, threonine and tyrosine, and acetylation of lysine; e) fixed modification: carbamidomethylation of cysteine.

To determine the SIRT1-interacting proteins, the HA-tagged SIRT1 immunoprecipitants (immunoprecipitated from twenty 10-cm dishes of MEFs stably expressing HA-tagged SIRT1) were subjected to SDS-PAGE, and were processed as described above. Data acquisition was performed as described above, except that an EASY-nLC 1200 System (Thermo) coupled to an Orbitrap Fusion Lumos Tribrid spectrometer (Thermo) equipped with an EASY-Spray Nanosource was used. Data were analysed using Proteome Discoverer (v.2.2, Thermo) against the mouse UniProt Reference Proteome database.

### Determination of oxygen consumption rates

The OCR of nematodes was measured as described previously^160^. Briefly, nematodes were washed with M9 buffer for 3 times. Some 15 to 25 nematodes were then suspended in 200 μl of M9 buffer, and were added to a well on a 96-well Seahorse XF Cell Culture Microplate. The measurement was performed in Seahorse XFe96 Analyzer (Agilent Technologies) at 20 °C following the manufacturer’s instruction, with a Seahorse XFe96 sensor cartridge (Agilent Technologies) pre-equilibrated in Seahorse XF Calibrant solution in a CO_2_-free incubator at 37 °C overnight.

Concentrations of respiratory chain inhibitors used during the assay were: FCCP at 10 μM and sodium azide at 40 mM. At the end of the assay, the exact number of nematode in each well was determined on a Cell Imaging Multi-Mode Reader (Cytation 1, BioTek) and was used for normalising/correcting OCR results. Data were collected using Wave 2.6.1 Desktop software (Agilent Technologies) and exported to Prism 9 (GraphPad) for further analysis according to manufacturer’s instructions.

The OCR of intact muscle tissue was measured as described previously^108,161^, with modifications. In brief, mice were starved for desired durations, and were sacrificed through cervical dislocation. The gastrocnemius muscles from two hindlegs were then excised, followed by incubating in 4 ml of dissociation media (DM; by dissolving 50 μg/ml gentamycin, 2% (v/v) FBS, 4 mg/ml collagenase A in DMEM containing HEPES) in a 35-mm culture dish in a humidified chamber at 37 °C, 5% CO_2_, for 1.5 h. The digested muscle masses were then washed with 4 ml of pre-warmed collagenase A-free DM, incubated in 0.5 ml of pre-warmed collagenase A-free DM, and dispersed by passing through a 20 G needle for 6 times. Some 20 μl of muscle homogenates was transferred to a well of a Seahorse XF24 Islet Capture Microplate (Agilent Technologies). After placing an islet capture screen by a Seahorse Capture Screen Insert Tool (Agilent Technologies) into the well, 480 μl of pre-warmed aCSF medium (120 mM NaCl, 3.5 mM KCl, 1.3 mM CaCl_2_, 0.4 mM KH_2_PO_4_, 1 mM MgCl_2_, 5 mM HEPES, 15 mM glucose, 1× MEM non-essential amino acids, 1 mM sodium pyruvate, and 1 mM GlutaMAX; adjust to pH 7.4 before use) was added, followed by equilibrating in a CO_2_-free incubator at 37 °C for 1 h. OCR was then measured at 37 °C in an XFe24 Extracellular Flux Analyzer (Agilent Technologies), with a Seahorse XFe24 sensor cartridge (Agilent Technologies) pre-equilibrated in Seahorse XF Calibrant solution (Agilent Technologies) in a CO_2_-free incubator at 37 °C overnight. Respiratory chain inhibitor used during the assay was oligomycin at 10 μM of final concentration. Data were collected using Wave 2.6.3 Desktop software (Agilent Technologies) and exported to Prism 9 (GraphPad) for further analysis according to the manufacturer’s instructions.

## Statistical analysis

Statistical analyses were performed using Prism 9 (GraphPad Software), except for the survival curves, which were analysed using SPSS 27.0 (IBM). Each group of data was subjected to Kolmogorov-Smirnov test, Anderson-Darling test, D’Agostino-Pearson omnibus test or Shapiro-Wilk test for normal distribution when applicable. An unpaired two-sided Student’s *t*-test was used to determine significance between two groups of normally distributed data. Welch’s correction was used for groups with unequal variances. An unpaired two-sided Mann-Whitney test was used to determine significance between data without a normal distribution. For comparisons between multiple groups with a fixed factor, an ordinary one-way ANOVA was used, followed by Tukey, Sidak, Dunnett or Dunn as specified in the legends. The assumptions of homogeneity of error variances were tested using F-test (*P* > 0.05). For comparison between multiple groups with two fixed factors, an ordinary two-way ANOVA was used, followed by Tukey’s or Sidak’s multiple comparisons test as specified in the legends. Geisser-Greenhouse’s correction was used where applicable. The adjusted means and s.e.m., or s.d., were recorded when the analysis met the above standards. Differences were considered significant when *P* &3C; 0.05, or *P* > 0.05 with large differences of observed effects (as suggested in refs. ^162,163^).

## Data availability

The data supporting the findings of this study are available within the paper and its Supplementary Information files. The MS proteomics data have been deposited to the ProteomeXchange Consortium (http://proteomecentral.proteomexchange.org) through the iProX partner repository^164,165^ with the dataset identifier IPX0007019000. Materials, reagents or other experimental data are available upon request. Full immunoblots are provided as Supplementary Information Fig. 1. Source data are provided with this paper.

## Code availability

The analysis was performed using standard protocols with previously described computational tools. No custom code was used in this study.

## Acknowledgements

We thank Dr. Sean Morrison (University of Texas Southwestern Medical Center) for providing the *AMPK*α*1*^F/F^ (The Jackson Laboratory, 014141), and *AMPK*α*2*^F/F^ (The Jackson Laboratory, 014142) mice and Richard Huganir (Johns Hopkins University School of Medicine) for the *PKC*ζ^F/F^ mice (The Jackson Laboratory, 024417); Mingliang Zhang (Shanghai Jiao Tong University Affiliated Sixth People’s Hospital) for the technical assistance of hyperinsulinemia euglycemic clamp; Su-Qin Wu, Ying He and Jing Song (Xiamen University) for mouse in vitro fertilisation; Ying Liu (Peking University) for nematode experiments; Bo Liu and Kewei Zheng (Xiamen University) for fly strains and experiments; Wei Wu (Chinese Academy of Sciences) fly embryo microinjection; Yiming Zheng (Xiamen University) for fly strains; Qiqi Guo from Zhenji Gan laboratory (Nanjing University) for the analysis of fibre types of muscle tissues; Xuan Guo (Jinzhou Medical University) for the helps of importing the fly strains; and all the other members of the S.-C.L. laboratory for technical assistance. We also acknowledge the *Caenorhabditis* Genetics Center and National BioResource Project for supplying nematode strains, and Bloomington *Drosophila* Stock Center, Vienna *Drosophila* Resource Center, and the Core Facility of *Drosophila* Resource and Technology, Center for Excellence in Molecular Cell Science, Chinese Academy of Sciences for providing fly strains and reagents. This work was supported by grants from the National Key R&D Program of China (2020YFA0803402, 2022YFA0806501), the National Natural Science Foundation of China (#92057204, #82088102, #32070753, and #92157001), the Fundamental Research Funds for the Central Universities (#20720200069), the Project “111” sponsored by the State Bureau of Foreign Experts and Ministry of Education of China (#BP2018017), the Joint Funds for the Innovation of Science and Technology, Fujian province (2021Y9232 and 2021Y9227), the Fujian provincial health technology project (2022ZD01005 and 2022ZQNZD009), the XMU Open Innovation Fund and Training Programme of Innovation and Entrepreneurship for Undergraduates (KFJJ-202103 and S202210384682), and the Agilent Applications and Core Technology - University Research Grant (#4769).

## Author contributions

Q.Q., C.-S.Z. and S.-C.L. conceived the study and designed the experiments. Q.Q. verified the requirement of the lysosomal pathway for LCA-mediated AMPK activation with the assistance from M.L. and X.T. Q.Q. identified the acetylated sites of v-ATPase required for AMPK activation with the assistance from J.-W.F., and determined the sirtuin requirement of LCA-mediated AMPK activation. Y.C. performed mouse and fly experiments, while Yu.W. conducted nematode experiments. S.L. and W.W. identified TULP3 as an acceptor of LCA. S.L. and H.-Y.Y. verified the binding affinity of TULP3 towards LCA, and the interaction between TULP3 and SIRT1 in vitro with the assistance from J.C. and B.S. S.L., H.-Y.Y. and C.Y. determined the activity of SIRT1. J.W. generated the V1E1-3KR-expressing mice. X.W. determined the OCR of mouse muscles. Y.-H.L. and S.X. determined the mRNA levels of mitochondrial OXPHOS complex. Z.W. and X.D. designed and synthesised the LCA probe, and X.H. generated the Ac-K99-V1E1 antibody. C.X., Yaying.W. and Z.X. performed the protein MS. C.Z. determined metabolites utilising HPLC-MS, and H.-L.P. CE-MS. B.Z. and X.D. performed the in silico docking assays. L.L. generated the TULP3-4G expressing nematodes, and ZK.X. the V1E1-3KR nematodes, both under the guidance from Y.Y. X.-S.X. helped interpret the v-ATPase acetylation data. C.-S.Z. and S.-C.L. wrote the manuscript.

## Additional information

### Supplementary information

The online version contains supplementary material available at https://doi.org/10.1038/.

Correspondence and requests for materials should be addressed to Sheng-Cai Lin.

**Peer review information** Nature thanks anonymous reviewer(s) for their contribution to the peer review of this work. Peer reviewer reports are available.

**Reprints and permissions information** is available at http://www.nature.com/reprints.

## Extended Data Figure legends

**Extended Data Fig. 1 LCA activates AMPK through the lysosomal AMPK pathway, downstream of the low glucose sensor aldolase-TRPV axis. a-e**, The lysosomal AMPK pathway is required for the LCA-induced AMPK activation. MEFs with *AXIN* (*AXIN1*) knockout (*AXIN*^-/-^; **a**), MEFs with *LAMTOR1* knockout (*LAMTOR1*^-/-^; **b**, **c**), or mice with hepatic (*AXIN1*-LKO; **d**) or muscular (*AXIN1***/***2*-MKO; **e**) AXIN knockout, were treated with LCA, either at 1 μM for 4 h (**a-c**), or coated with (2-hydroxypropyl)- β-cyclodextrin and supplied at 1 g/l in drinking water for 1 week (**d**, **e**), followed by determining the activity of AMPK (**a**, **d**, **e**), the activity of v-ATPase (**b**; statistical analysis data are shown in the upper panel of Fig. 1f), and the lysosomal translocation of AXIN (**c**; statistical analysis data are shown in the lower panel of Fig. 1f). **f**, CR leads to a constitutive activation of AMPK in muscle. Mice were subjected to CR for 4 months, followed by determining the muscular AMPK at different times of the day. **g-j**, The aldolase-TRPV axis that is triggered by low glucose is dispensable for the LCA-triggered lysosomal pathway. The aldolase knockdown MEFs re-introduced with ALDOA-D34S (**g**, **h**), and the TRPV-QKO MEFs (**i**, **j**) were treated with 1 μM LCA for 4 h, followed by determining the activity of v-ATPase (**g**, **i**; statistical analysis data are shown in Fig. 1k and 1n), and the lysosomal translocation of AXIN (**j**, **h**; statistical analysis data are shown in Fig. 1o and 1l). Experiments in this figure were performed three times.

**Extended Data Fig. 2 LCA triggers the lysosomal AMPK pathway unrelated to the PEN2-ATP6AP1 axis. a-f**, LCA could still activate AMPK in MEFs depleted of the PEN2-ATP6AP1 axis. The *PEN2* knockout (*PEN2*^-/-^) MEFs (**a-c**) and the *ATP6AP1* knockout (*ATP6AP1*^-/-^) MEFs with ATP6AP1^Δ420–440^ re-introduction (**d-f**), both are deficient in the PEN2-ATP6AP1 axis^23^, were treated with 1 μM LCA for 4 h. The activation of AMPK (**a**, **d**), the activity of v-ATPase (**b**, **e**), and the lysosomal translocation of AXIN (**c**, **f**) were then determined. Statistical analysis data of **b**, **c**, **e**, **f** are shown as mean ± s.e.m., with **b** and **e** normalised to the DMSO group. The *n* numbers were labelled in each panel, and *P* value by two-sided Student’s *t*-test (**c**), two-sided Student’s *t*-test with Welch’s correction (**b**), two-sided Mann-Whitney test (**e**), or two-way ANOVA followed by Tukey’s test (**f**). Experiments in this figure were performed three times.

**Extended Data Fig. 3 LCA triggers the lysosomal AMPK pathway through deacetylation of v-ATPase. a**, **b**, V1E1-3KR renders the lysosomal AMPK pathway constitutively active. HEK293T with ectopic expression of V1E1 or V1E1-3KR were lysed, followed by determining the activity of AMPK (**a**), the activity of v-ATPase (upper panel of **b**), and the translocation of AXIN (lower panel of **b**). The statistical analysis data for **b** are shown in Fig. 2f. **c**, Validation of the antibody specifically against K99-acetylated V1E1. HEK293T cells were transfected with V1E1-K99R, or V1E1-K52R, V1E1-K191R and the wildtype V1E1 as controls, followed by determining the reaction specificity of the Ac-K99-V1E1 antibody by immunoblotting. Experiments in this figure were performed three times, except **a** four times.

**Extended Data Fig. 4 LCA promotes sirtuins to deacetylase v-ATPase. a**, Sirtuins, but not HDACs, deacetylate V1E1 and activate AMPK. HEK293T cells were infected with lentiviruses carrying HA-tagged HDAC1 to HDAC11, followed by determining the acetylation of V1E1 and the activation of AMPK. The effects of sirtuins on AMPK activation are shown in Fig. 3a. **b**, Sirtuins inhibit v-ATPase. HEK293T cells infected with lentiviruses carrying each sirtuin (HA-tagged SIRT1 to SIRT7), or its dominant negative (DN) form, were subjected to determine the activity of v-ATPase by immunostaining (statistical analysis data are shown in Fig. 3b). **c-e**, Depletion of all members of the sirtuin family blocks the activation of AMPK by LCA. MEFs with hepta-knockout of SIRT1 to SIRT7 (*SIRT1*-*7*^-/-^; knockout strategy and the validation data are shown in **c**) were treated with 1 μM LCA for 4 h, followed by determining the activity of v-ATPase (**d**; statistical analysis data are shown in Fig. 3d), and the lysosomal translocation of AXIN (**e**; statistical analysis data are shown in Fig. 3e). **f**, The *SIRT1*-*7*^-/-^ MEFs grow much slower than wildtype MEFs. The *SIRT1*-*7*^-/-^ MEFs and the wildtype control MEFs were regularly cultured, followed by determining the proliferation rates by CCK8 assays. Results are shown as mean ± s.e.m.; *n* = 6 biological replicates, and *P* value by two-way ANOVA followed by Tukey’s test. **g-i**, Inhibition of SIRT1 in *SIRT2*-*7*^-/-^ MEFs blocks the activation of AMPK by LCA. The *SIRT2*-*7*^-/-^ MEFs (validated in **g**) were pre-treated with 10 μM EX-527 for 12 h, followed by treatment with 1 μM LCA for 4 h. The activity of v-ATPase (**h**; and statistical analysis data in Fig. 3g), and the lysosomal translocation of AXIN (**i**; and statistical analysis data in Fig. 3h). **j**, Validation data for the SIRT1 activity assay. Experiments were performed as described in the “Determination of SIRT1 activity” of Methods section. The representative spectrograms of acetylated histone H3 (left panel) and acetylated V1E1 (right panel) peptides before (-SIRT1) and after (+SIRT1) incubating with SIRT1 are shown. The presence of deacetylated histone H3 (m/z 453.2403 and 905.4805) and V1E1 (m/z 529.8019 and 1058.5959) peaks indicate that SIRT1 effectively catalysed the deacetylation reaction. Experiments in this figure were performed three times, except **a** four times.

**Extended Data Fig. 5 Identification of binding partners for the LCA probe. a-d**, Synthesis and purification of a photoactive LCA probe (WZH-15-080). Reactions for conjugating 2-(3-(but-3-yn-1-yl)-3H-diazirin-3-yl)ethan-1-amine to LCA that introduced a diazirine with a terminal alkyne moiety the carboxyl group of LCA (**a**; see detailed procedures in the “synthesis of LCA probe” of Methods section). The product was then purified on a preparative HPLC (**a**), and the spectrums of ^1^H NMR (**b**), ^13^C NMR (**c**), and the HSMS (**d**) are shown. **e**, Procedures for forming the complexes of biotinylated LCA probe and cellular proteins. To determine the binding affinity of LCA for TULP3 by the competition assay (left panel), the LCA probe was first biotinylated by mixing with Cu(II) salt which catalyses a [3 + 2] azide-alkyne cycloaddition with a biotin-azide linker, followed by incubating with Streptavidin beads, and then the TULP3 protein. The TULP3 protein captured by the biotinylated LCA probe-bound Streptavidin beads was then competitively eluted by different concentrations of LCA. To determine the potential binding site of LCA on TULP3 (right panel), the TULP3 protein was incubated with the LCA probe, followed by exposure to UV light. The LCA probe-protein conjugates were then mixed with Cu(II) salt and the biotin-azide linker, thus biotinylating probe-target complexes, allowing for the pull-down of such complexes for MS analysis with Streptavidin beads. Experiments in this figure were performed three times.

**Extended Data Fig. 6 TULP3 mediates LCA to activate sirtuins. a**, **b**, TULP3 is required for the LCA-mediated activation of SIRT1 and AMPK. MEFs with *TULP3* knocked down were treated with 1 μM LCA for 4 h, followed by determining the activity of AMPK (**a**), the acetylation of V1E1 (**a**), and the activity of v-ATPase (**b**). Statistical analysis data of **b** are shown as mean ± s.e.m., normalised to the DMSO group; *n* = 21 (DMSO) or 24 (LCA) cells, and *P* value by two-sided Student’s *t*-test. **c**, Knockout strategy and validation data of *TULP3*^-/-^ MEFs (both clone #1 and #2). **d**, Knockout of TULP3 abrogates the LCA-induced activation of SIRT1 and AMPK. The clone #2 of *TULP3*^-/-^ MEFs were treated with 1 μM LCA for 4 h, followed by determining the activity of AMPK by immunoblotting. **e**, TULP3 interacts with all 7 members of the sirtuin family independently of LCA. HEK293T cells were transfected with Myc-tagged TULP3 and each HA-tagged sirtuin (SIRT1 to SIRT7). Cells were then lysed, and a concentration of 5 μM LCA was added to the lysate. After incubation for 1 h, TULP3 was immunoprecipitated, and the co-immunoprecipitated sirtuins were probed through immunoblotting. **f**, In silico modelling of LCA bound to TULP3. Modelling was performed according to the results of MS on purified TULP3 conjugated to the LCA-probe, and the AlphaFold-predicted TULP3 structure (https://alphafold.ebi.ac.uk/entry/O75386; coloured in green). The Y193, P195, K333 and P336 residues (coloured in yellow) that comprise a hydrophobic pocket for LCA (coloured in red) binding as indicated were mutated to glycine (TULP3-4G) to create a TULP3 mutant that is defective in binding to LCA. **g**, **h**, TULP3-4G, unable to bind LCA, blocked the LCA-mediated activation of AMPK. *TULP3*^-/-^ MEFs were infected with lentivirus carrying TULP3-4G mutant or its wildtype control, followed by treatment with 1 μM LCA for 4 h. The activity of v-ATPase (**g**) and the lysosomal translocation of AXIN (**h**) were then determined. The statistical analysis data of **g** and **h** are shown in Fig. 4k and 4l. Experiments in this figure were performed three times.

**Extended Data Fig. 7 Other known binding partners of LCA are not required for the activation of AMPK. a-s**, MEFs with *FXR* (**a**, clone #1 on the left or upper panel, and clone #2 right or lower, and the same hereafter in this figure), *FXRb* (**b**), *PXR* (**c**), *VDR* (**d**), *CAR* (**e**), *LXRa* (**f**), *LXRb* (**g**), *S1PR2* (**h**), *TGR5* (**i**), *CHRM2* (**j**), *CHRM3* (**k**), *PKC*ζ (**l**), *FAS* (**m**), *FPR* (**n**), or *YES* (**o**) knockout, or with *LXRa* and *LXRb* (**p**), *FXRa* and *FXRb* (**q**), *PXR* and *CAR* (**r**), or *CHRM2* and *CHRM3* (**s**) double knockout, were treated with 1 μM LCA for 4 h, followed by determining the activity of AMPK. Experiments in this figure were performed three times.

**Extended Data Fig. 8 The LCA-TULP3-sirtuins-v-ATPase axis retards ageing. a-d**, V1E1-3KR improves muscle function in aged mice. WT or muscle-specific *AMPK*α knockout (α-MKO) mice with muscle V1E1-3KR or wildtype V1E1 expression (induced by tamoxifen at 18 months old, for 4 weeks) were subjected to the analysis of muscular AMPK activation (**a**), the mRNA and protein levels of the OXPHOS complex (**b**), the body composition (**c**), and the RQ and ambulatory activity (**d**). Data in **b** are shown as mean ± s.e.m., normalised to the WT + V1E1 group (*n* = 4 mice for each genotype, and *P* value by two-way ANOVA followed by Tukey’s test); in **c** as mean ± s.e.m. (*n* = 5 mice for each genotype, and *P* value by two-way ANOVA followed by Tukey’s test); and in **d** as mean (at 5-min intervals during a 24-h course; *n* = 4 mice for each genotype), and the others as box-and-whisker plots (*n* = 4 mice for each genotype, and *P* value by two-way ANOVA followed by Tukey’s test). **e-i**, The TULP3-sirtuins-v-ATPase axis is required for LCA to activate AMPK and extend lifespan in nematodes and flies. Nematodes and flies, either with *TULP3* knockout (upper panel of **e**, and left panel of **f**) or knockdown (*Act5C*-*GAL4* > *ktub* RNAi; lower panel of **e**, and right panel of **f**), with TULP3-4G reintroduction into the *TULP3*-knockout background (**g**), with sirtuin depletion (**h**), or with V1E1-3KR expression (**i**) were cultured in the medium containing LCA at 100 μM, followed by determining the activation of AMPK (**e**, **g**, **h**, **i**) and lifespan (**f**; shown as Kaplan-Meier curves). **j**, **k**, **p**, The TULP3-sirtuins-v-ATPase axis is required for LCA to promote resistance to oxidative stress and starvation. Flies, either with TULP3-4G reintroduction into the *TULP3*-knockout background (left panel of **j**, **k**), with sirtuin depletion (right panel of **j**, **k**), or with V1E1-3KR expression (**p**) were cultured in the medium containing LCA at 100 μM, followed by transferring to media containing 5% H_2_O_2_ (**j**, and left panel of **p**) or deprived of food (**k**, and right panel of **p**). Lifespan data are shown as Kaplan-Meier curves. **l-o**, **q**, **r**, The TULP3-sirtuins-v-ATPase axis elevates mitochondrial contents and respiratory functions in nematodes and flies. Nematodes and flies with TULP3-4G re-introduction in the background of *TULP3*-knockout (**l**, **m**), with sirtuins depletion (**n**, **o**), or with V1E1-3KR expression (**q**, **r**) were treated with LCA, followed by determining the mRNA levels of OXPHOS complex (**l**, **n**, **q**; results are mean ± s.e.m.; *n* = 4 samples for each genotype/treatment, and *P* value by two-way ANOVA followed by Tukey’s test, except those of flies in **n**, **q** two-sided Student’s *t*-test), and OCR (**m**, **o**, **r**; results are mean ± s.e.m.; *n* = 4 samples for each genotype/treatment, and *P* value by two-way ANOVA followed by Tukey’s test, except **o** two-sided Student’s *t*-test). **s**, Schematic diagram showing that the v-ATPase serves as a common entry into the lysosomal AMPK pathway. LCA, elevated after CR, binds to TULP3 and activates sirtuins, which in turn deacetylate the V1E1 subunit of v-ATPase (on K52, K99 and K191 residues) and inhibits v-ATPase, which along with Ragulator allows for AXIN/LKB1 to translocate to the surface of the lysosome, where LKB1 phosphorylates and activates AMPK. In response to low glucose, the FBP-unoccupied aldolase interacts and inhibits the cation channel TRPV, which in turn interacts and reconfigures v-ATPase, allowing for AXIN/LKB1 to translocate to the lysosome to activate AMPK. Likewise, metformin binds to its target PEN2, and the metformin-bound PEN2 is recruited to v-ATPase via interacting with ATP6AP1, thereby inhibiting v-ATPase. The v-ATPase hence acts as a common node for the signalling of low glucose, metformin and LCA to intersect, all leading to AMPK activation and manifestation of anti-aging effects. Experiments in this figure were performed three times.

**Extended Data Table 1 A summary of acetylated residues of v-ATPase**. The table contains the acetylated residues for each of the 21 subunits of v-ATPase (ATP6V1A, ATP6V1B2, ATP6V1C1, ATP6V1C2, ATP6V1D, ATP6V1E1, ATP6V1E2, ATP6V1F, ATP6V1G1, ATP6V1G2, ATP6V1G3, ATP6V1H, ATP6V0A4, ATP6V0A2, ATP6V0B, ATP6V0C, ATP6V0D1, ATP6V0D2, ATP6V0E1, ATP6AP1 and ATP6AP2) detected.

**Extended Data Table 2 Summary of lifespan and analysis in nematodes and flies.**

**Supplementary Table 1 A complete list of the post-translational modifications identified for v-ATPase**. See each sheet for the raw data of all peptides with post-translational modifications hit from each of the 21 subunits of v-ATPase.

**Supplementary Table 2 List of the proteins co-immunoprecipitated with SIRT1**. MEFs with stable expression of HA-tagged SIRT1 were subjected to immunoprecipitation using anti-HA antibody, followed by MS. On the list of the “SIRT1-specific” proteins shows a total of 1,655 co-immunoprecipitated with SIRT1, after subtracting hits of the proteins in the “IP IgG” sheet from the “IP SIRT1” sheet. After engineering the expression plasmids for all these 1,655 proteins, 157 of them could be co-immunoprecipitated by SIRT1 when individually expressed in HEK293T cells, as listed in the “SIRT1-interacting proteins” sheet. The “SIRT1-interacting proteins” sheet also includes the results of AMPK activation by LCA treatment in MEFs with these 157 proteins individually knocked down.

