## Supplementary material for "Lithocholic acid targets TULP3 to activate sirtuins and AMPK to retard ageing": ED figures 1-8 and ED tables 1-2

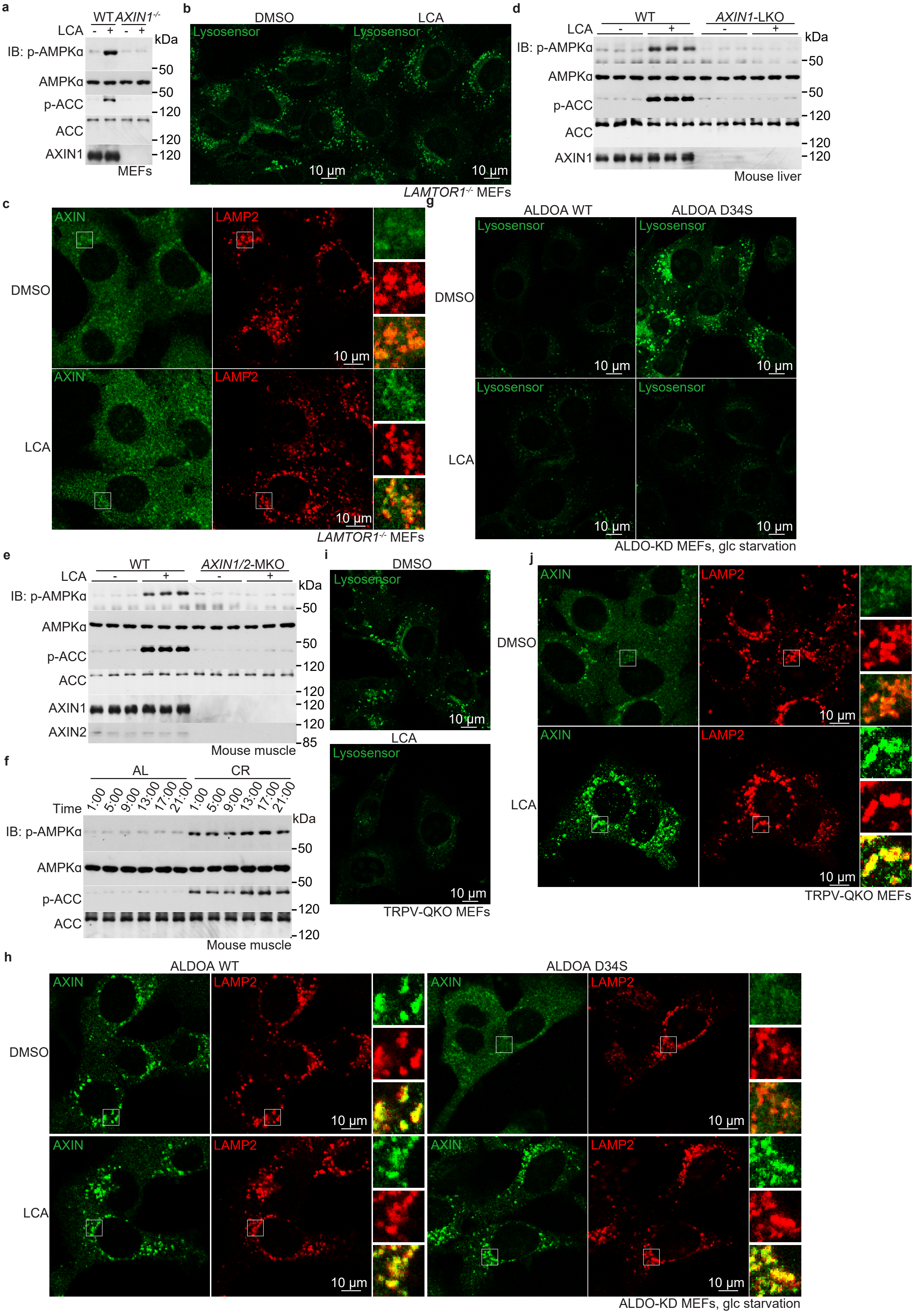


**Extended Data Fig. 1 | LCA activates AMPK through the lysosomal AMPK pathway, downstream of the low glucose sensor aldolase-TRPV axis.**

**a-e**, The lysosomal AMPK pathway is required for the LCA-induced AMPK activation. MEFs with *AXIN* (*AXIN1*) knockout (*AXIN*$^{-/-}$; **a**), MEFs with *LAMTOR1* knockout (*LAMTOR1*$^{-/-}$; **b**, **c**), or mice with hepatic (*AXIN1*-LKO; **d**) or muscular (*AXIN1/2*-MKO; **e**) AXIN knockout, were treated with LCA, either at 1 μM for 4 h (**a-c**), or coated with (2-hydroxypropyl)-β-cyclodextrin and supplied at 1 g/l in drinking water for 1 week (**d**, **e**), followed by determining the activity of AMPK (**a**, **d**, **e**), the activity of v-ATPase (**b**; statistical analysis data are shown in the upper panel of Fig. 1f), and the lysosomal translocation of AXIN (**c**; statistical analysis data are shown in the lower panel of Fig. 1f).

**f**, CR leads to a constitutive activation of AMPK in muscle. Mice were subjected to CR for 4 months, followed by determining the muscular AMPK at different times of the day.

**g-j**, The aldolase-TRPV axis that is triggered by low glucose is dispensable for the LCA-triggered lysosomal pathway. The aldolase knockdown MEFs re-introduced with ALDOA-D34S (**g**, **h**), and the TRPV-QKO MEFs (**i**, **j**) were treated with 1 μM LCA for 4 h, followed by determining the activity of v-ATPase (**g**, **i**; statistical analysis data are shown in Fig. 1k and 1n), and the lysosomal translocation of AXIN (**j**, **h**; statistical analysis data are shown in Fig. 1o and 1l).

Experiments in this figure were performed three times.

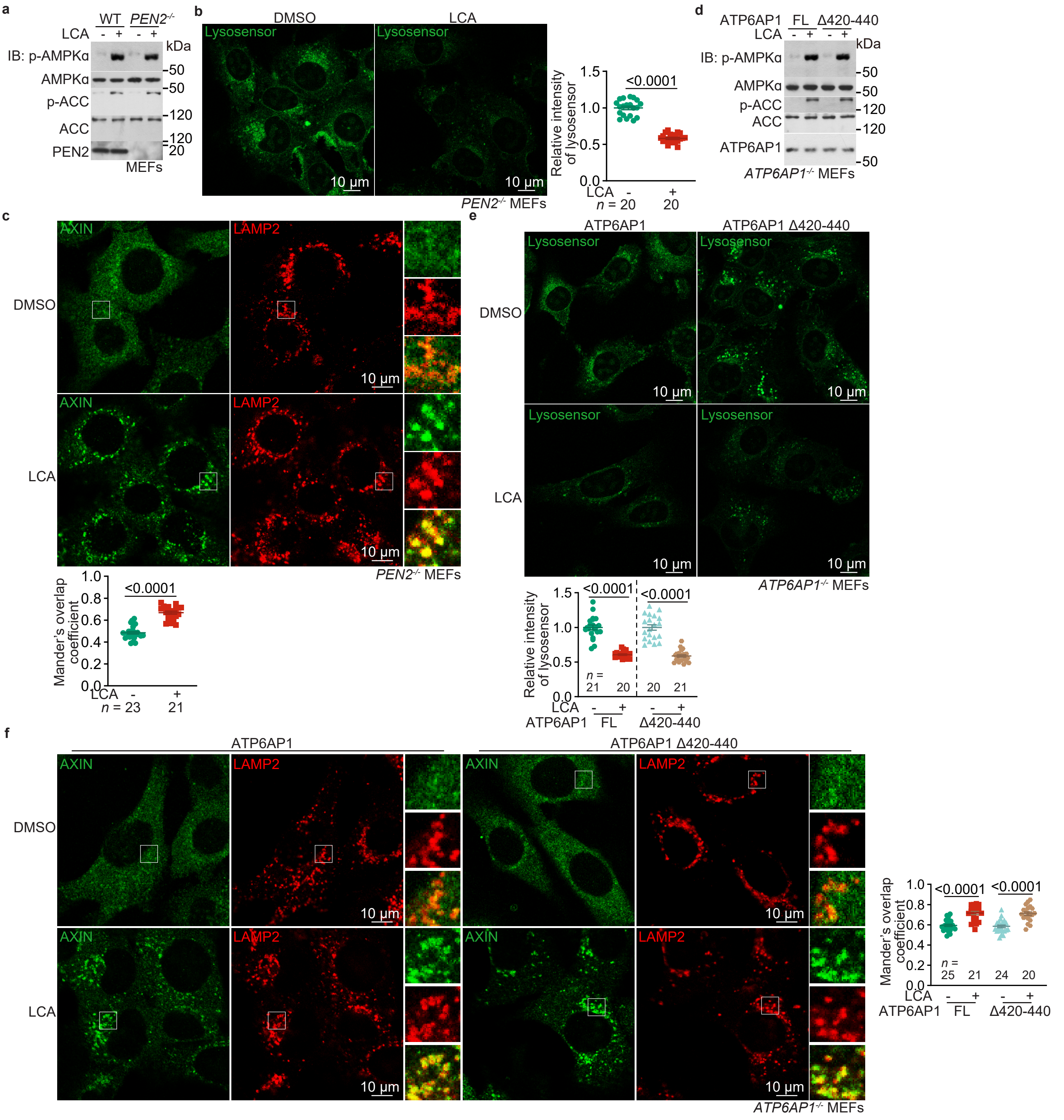


**Extended Data Fig. 2 | LCA triggers the lysosomal AMPK pathway unrelated to the PEN2-ATP6AP1 axis.**
**a-f**, LCA could still activate AMPK in MEFs depleted of the PEN2-ATP6AP1 axis. The *PEN2* knockout (*PEN2*$^{-/-}$) MEFs (**a**-**c**) and the *ATP6AP1* knockout (*ATP6AP1*$^{-/-}$) MEFs with ATP6AP1$^{\Delta 420-440}$ re-introduction (**d**-**f**), both are deficient in the PEN2-ATP6AP1 axis[23], were treated with 1 μM LCA for 4 h. The activation of AMPK (**a**, **d**), the activity of v-ATPase (**b**, **e**), and the lysosomal translocation of AXIN (**c**, **f**) were then determined. Statistical analysis data of **b**, **c**, **e**, **f** are shown as mean ± s.e.m., with **b** and **e** normalised to the DMSO group. The *n* numbers were labelled in each panel, and *P* value by two-sided Student's *t*-test (**c**), two-sided Student's *t*-test with Welch's correction (**b**), two-sided Mann-Whitney test (**e**), or two-way ANOVA followed by Tukey's test (**f**).
Experiments in this figure were performed three times.

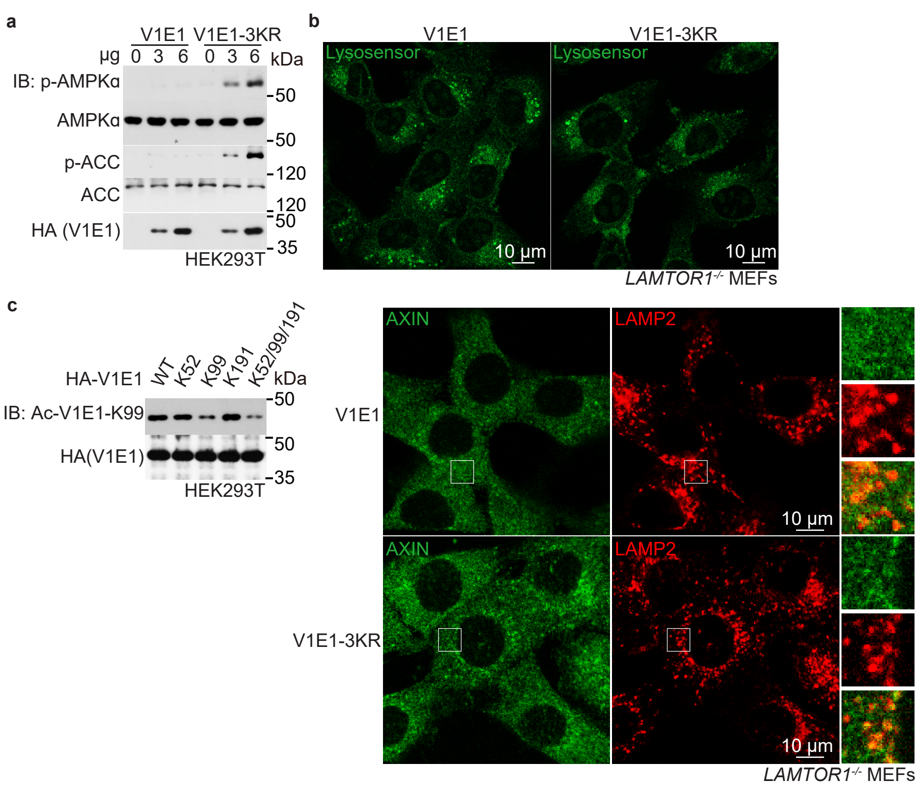


**Extended Data Fig. 3 | LCA triggers the lysosomal AMPK pathway through deacetylation of v-ATPase**.

**a**, **b**, V1E1-3KR renders the lysosomal AMPK pathway constitutively active. HEK293T with ectopic expression of V1E1 or V1E1-3KR were lysed, followed by determining the activity of AMPK (**a**), the activity of v-ATPase (upper panel of **b**), and the translocation of AXIN (lower panel of **b**). The statistical analysis data for **b** are shown in Fig. 2f.

**c**, Validation of the antibody specifically against K99-acetylated V1E1. HEK293T cells were transfected with V1E1-K99R, or V1E1-K52R, V1E1-K191R and the wildtype V1E1 as controls, followed by determining the reaction specificity of the Ac-K99-V1E1 antibody by immunoblotting.

Experiments in this figure were performed three times, except **a** four times.

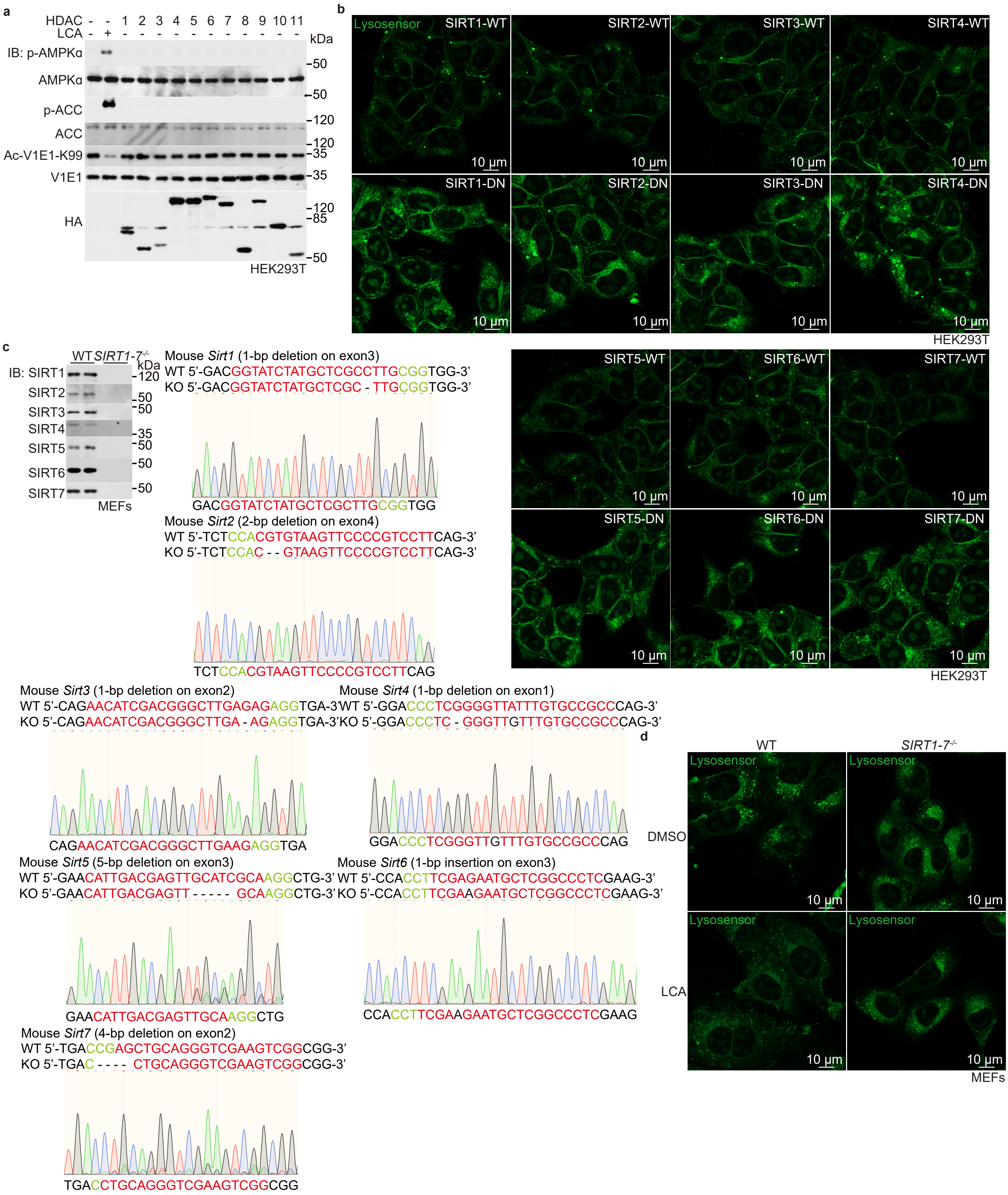


**Extended Data Fig. 4**

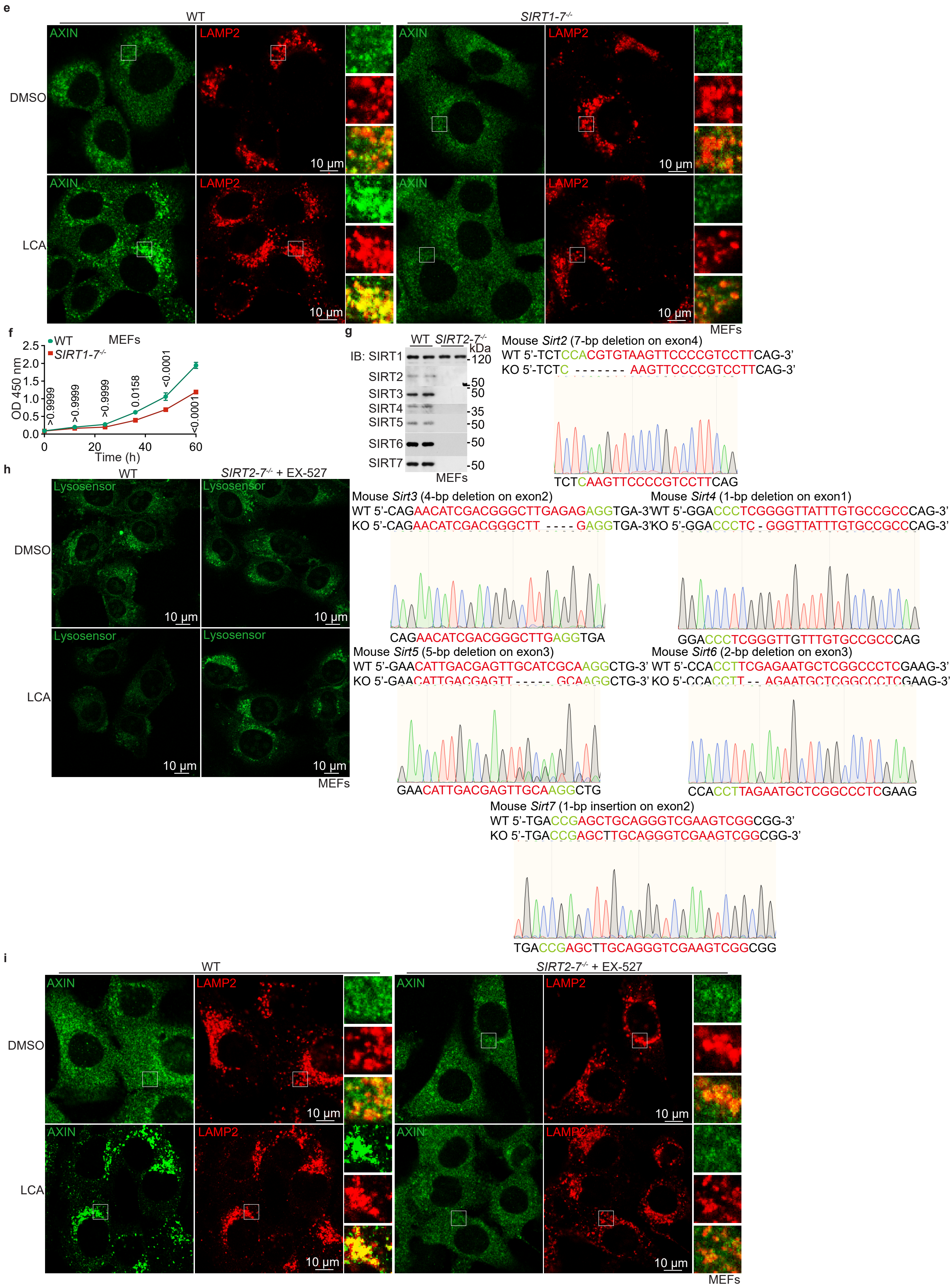


**Extended Data Fig. 4 (cont.)**

**j**

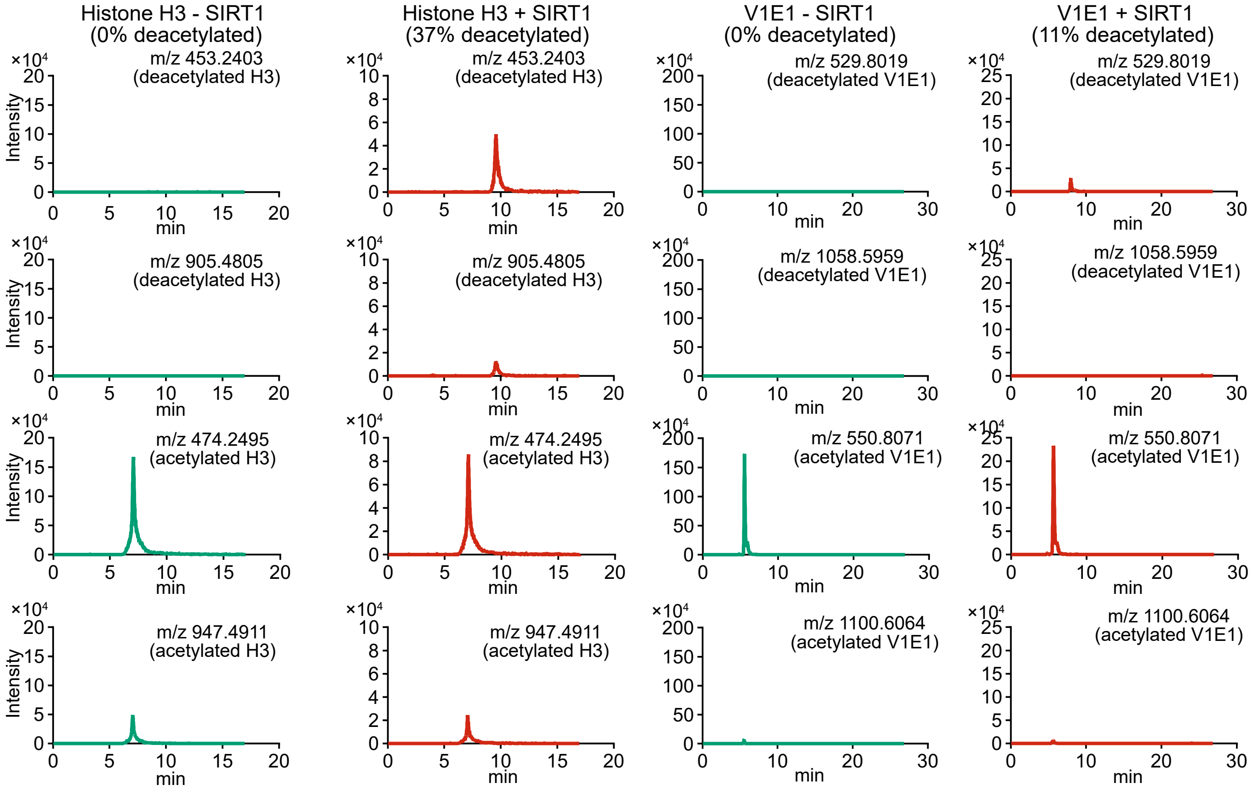


**Extended Data Fig. 4 | LCA promotes sirtuins to deacetylase v-ATPase**.

**a**, Sirtuins, but not HDACs, deacetylate V1E1 and activate AMPK. HEK293T cells were infected with lentiviruses carrying HA-tagged HDAC1 to HDAC11, followed by determining the acetylation of V1E1 and the activation of AMPK. The effects of sirtuins on AMPK activation are shown in Fig. 3a.

**b**, Sirtuins inhibit v-ATPase. HEK293T cells infected with lentiviruses carrying each sirtuin (HA-tagged SIRT1 to SIRT7), or its dominant negative (DN) form, were subjected to determine the activity of v-ATPase by immunostaining (statistical analysis data are shown in Fig. 3b).

**c-e**, Depletion of all members of the sirtuin family blocks the activation of AMPK by LCA. MEFs with hepta-knockout of SIRT1 to SIRT7 (*SIRT1-7*$^{-/-}$; knockout strategy and the validation data are shown in **c**) were treated with 1 µM LCA for 4 h, followed by determining the activity of v-ATPase (**d**; statistical analysis data are shown in Fig. 3d), and the lysosomal translocation of AXIN (**e**; statistical analysis data are shown in Fig. 3e).

**f**, The *SIRT1-7*$^{-/-}$ MEFs grow much slower than wildtype MEFs. The *SIRT1-7*$^{-/-}$ MEFs and the wildtype control MEFs were regularly cultured, followed by determining the proliferation rates by CCK8 assays. Results are shown as mean ± s.e.m.; $n$ = 6 biological replicates, and $P$ value by two-way ANOVA followed by Tukey's test.

**g-i**, Inhibition of SIRT1 in *SIRT2-7*$^{-/-}$ MEFs blocks the activation of AMPK by LCA. The *SIRT2-7*$^{-/-}$ MEFs (validated in **g**) were pre-treated with 10 µM EX-527 for 12 h, followed by treatment with 1 µM LCA for 4 h. The activity of v-ATPase (**h**; and statistical analysis data in Fig. 3g), and the lysosomal translocation of AXIN (**i**; and statistical analysis data in Fig. 3h).

**j**, Validation data for the SIRT1 activity assay. Experiments were performed as described in the "Determination of SIRT1 activity" of Methods section. The representative spectrograms of acetylated histone H3 (left panel) and acetylated V1E1 (right panel) peptides before (-SIRT1) and after (+SIRT1) incubating with SIRT1 are shown. The presence of deacetylated histone H3 (m/z 453.2403 and 905.4805) and V1E1 (m/z 529.8019 and 1058.5959) peaks indicate that SIRT1 effectively catalysed the deacetylation reaction.

Experiments in this figure were performed three times, except **a** four times.

**a**

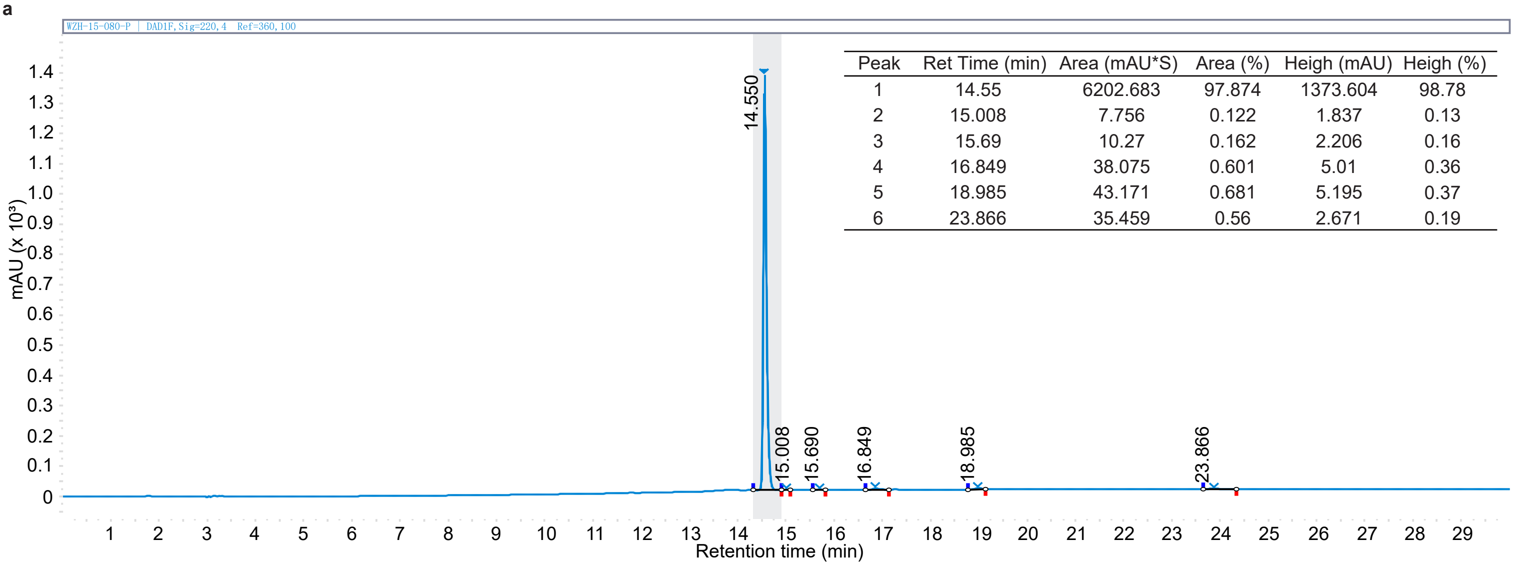


| Peak | Ret Time (min) | Area (mAU*S) | Area (%) | Heigh (mAU) | Heigh (%) |
|---|---|---|---|---|---|
| 1 | 14.55 | 6202.683 | 97.874 | 1373.604 | 98.78 |
| 2 | 15.008 | 7.756 | 0.122 | 1.837 | 0.13 |
| 3 | 15.69 | 10.27 | 0.162 | 2.206 | 0.16 |
| 4 | 16.849 | 38.075 | 0.601 | 5.01 | 0.36 |
| 5 | 18.985 | 43.171 | 0.681 | 5.195 | 0.37 |
| 6 | 23.866 | 35.459 | 0.56 | 2.671 | 0.19 |

**b**

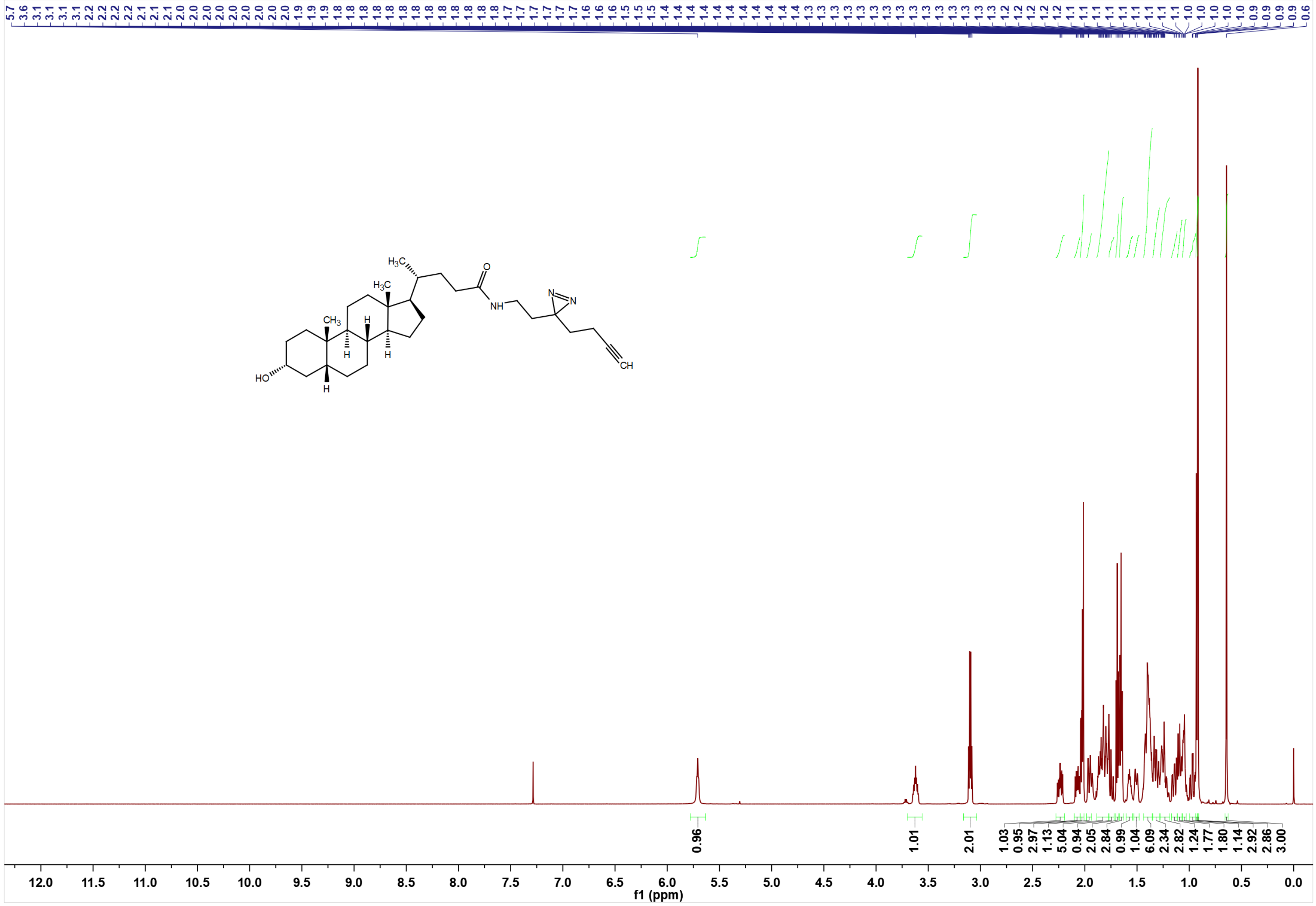


**Extended Data Fig. 5**

c

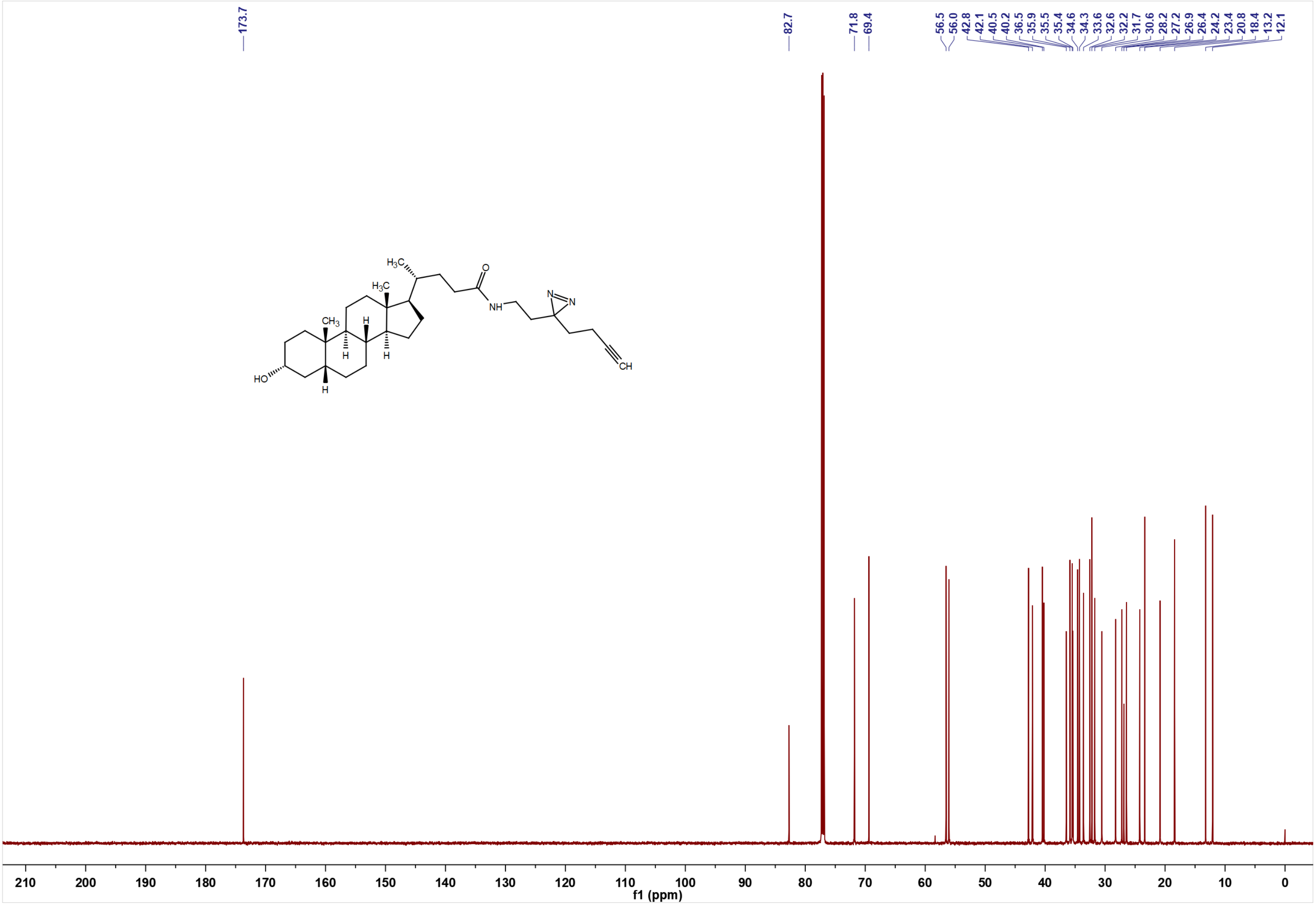


d

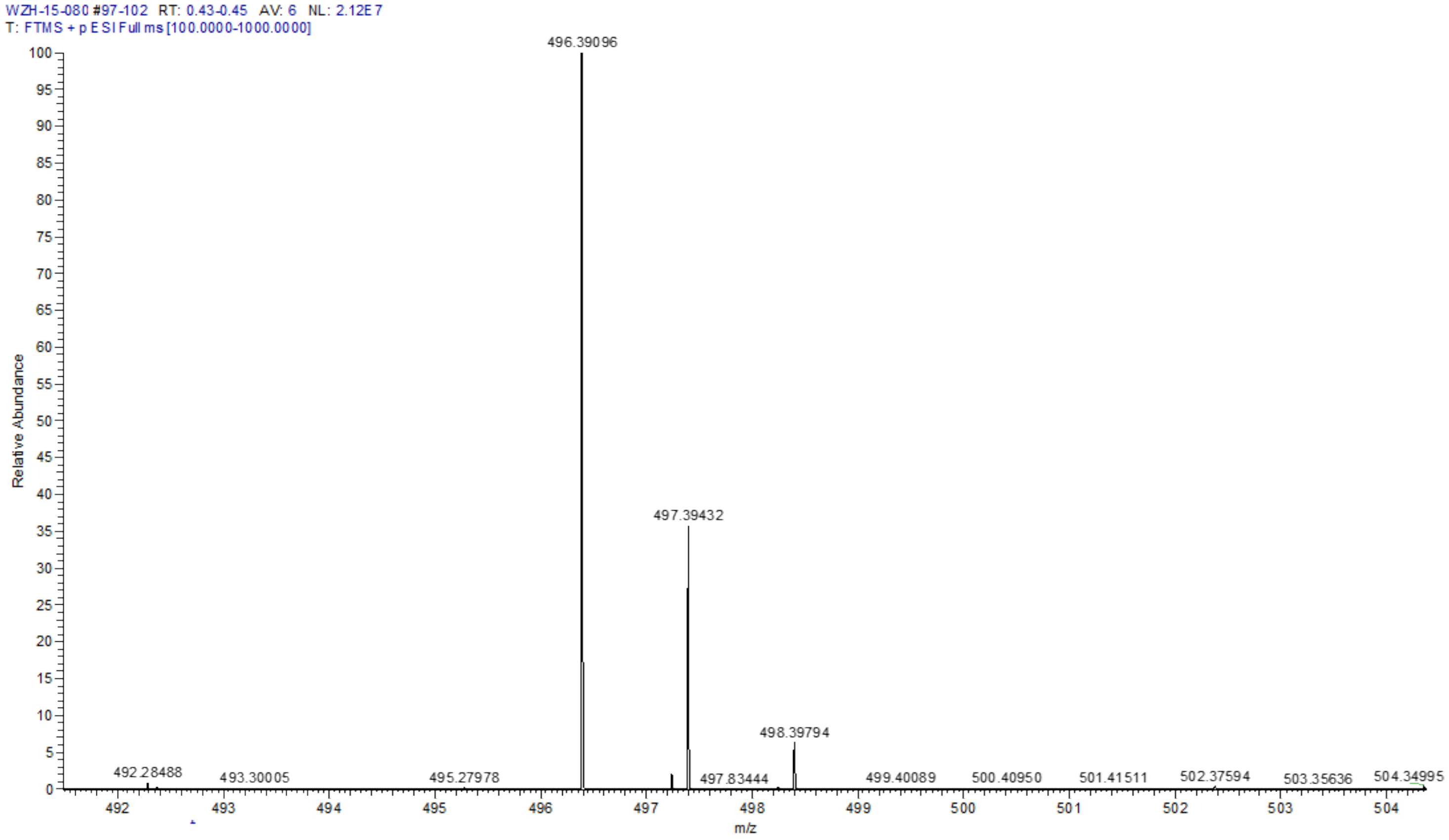


m/z= 491.39096-501.39096

| m/z | Theo. Mass | Delta (ppm) | RDB equiv. | Composition |
|---|---|---|---|---|
| 496.39096 | 496.38975 | 2.43 | 8.5 | $C_{31}H_{50}O_{2}N_{3}$ |
| | 496.39294 | -3.99 | 0.5 | $C_{20}H_{50}O_{5}N_{9}$ |

**Extended Data Fig. 5 (cont.)**

**e**

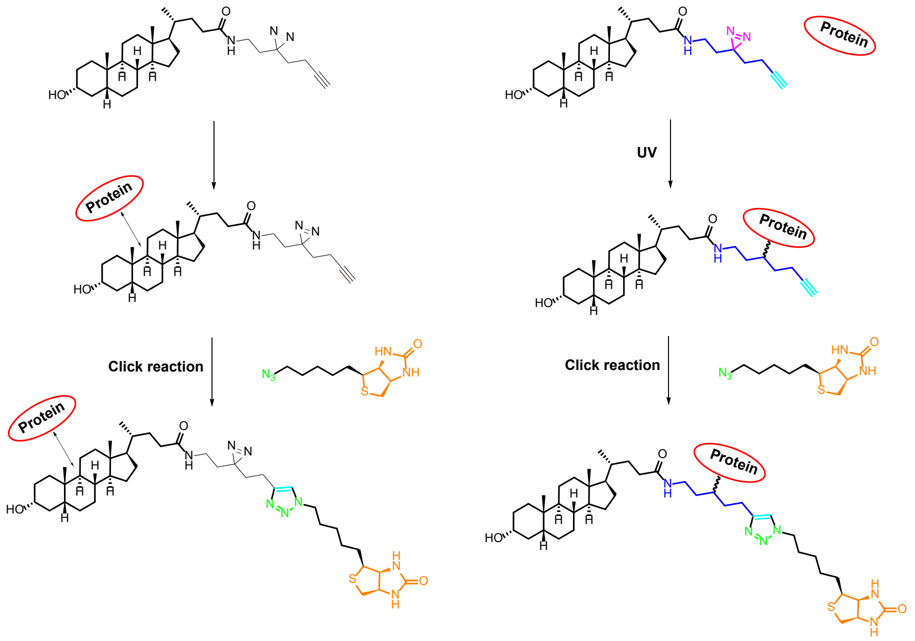


**Extended Data Fig. 5 | Identification of binding partners for the LCA probe**.
**a**-**d**, Synthesis and purification of a photoactive LCA probe (WZH-15-080). Reactions for conjugating 2-(3-(but-3-yn-1-yl)-3H-diazirin-3-yl)ethan-1-amine to LCA that introduced a diazirine with a terminal alkyne moiety the carboxyl group of LCA (**a**; see detailed procedures in the "synthesis of LCA probe" of Methods section). The product was then purified on a preparative HPLC (**a**), and the spectrums of $^{1}H$ NMR (**b**), $^{13}C$ NMR (**c**), and the HSMS (**d**) are shown.
**e**, Procedures for forming the complexes of biotinylated LCA probe and cellular proteins. To determine the binding affinity of LCA for TULP3 by the competition assay (left panel), the LCA probe was first biotinylated by mixing with Cu(II) salt which catalyses a [3 + 2] azide-alkyne cycloaddition with a biotin-azide linker, followed by incubating with Streptavidin beads, and then the TULP3 protein. The TULP3 protein captured by the biotinylated LCA probe-bound Streptavidin beads was then competitively eluted by different concentrations of LCA. To determine the potential binding site of LCA on TULP3 (right panel), the TULP3 protein was incubated with the LCA probe, followed by exposure to UV light. The LCA probe-protein conjugates were then mixed with Cu(II) salt and the biotin-azide linker, thus biotinylating probe-target complexes, allowing for the pull-down of such complexes for MS analysis with Streptavidin beads.
Experiments in this figure were performed three times.

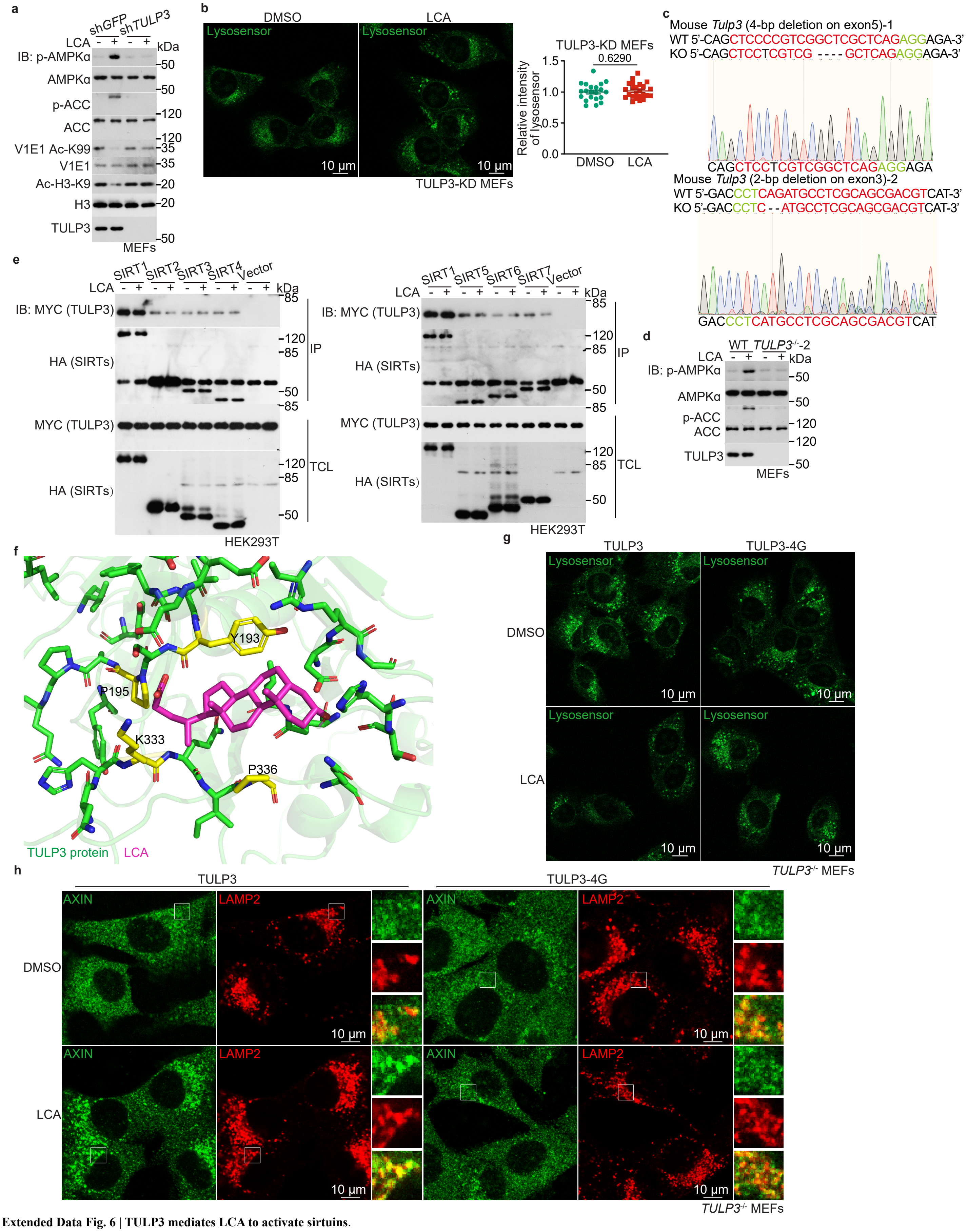


**Extended Data Fig. 6 | TULP3 mediates LCA to activate sirtuins.**

**a**, **b**, TULP3 is required for the LCA-mediated activation of SIRT1 and AMPK. MEFs with *TULP3* knocked down were treated with 1 μM LCA for 4 h, followed by determining the activity of AMPK (**a**), the acetylation of V1E1 (**a**), and the activity of v-ATPase (**b**). Statistical analysis data of **b** are shown as mean ± s.e.m., normalised to the DMSO group; *n* = 21 (DMSO) or 24 (LCA) cells, and *P* value by two-sided Student's *t*-test.

**c**, Knockout strategy and validation data of *TULP3*$^{-/-}$ MEFs (both clone #1 and #2).

**d**, Knockout of TULP3 abrogates the LCA-induced activation of SIRT1 and AMPK. The clone #2 of *TULP3*$^{-/-}$ MEFs were treated with 1 μM LCA for 4 h, followed by determining the activity of AMPK by immunoblotting.

**e**, TULP3 interacts with all 7 members of the sirtuin family independently of LCA. HEK293T cells were transfected with Myc-tagged TULP3 and each HA-tagged sirtuin (SIRT1 to SIRT7). Cells were then lysed, and a concentration of 5 μM LCA was added to the lysate. After incubation for 1 h, TULP3 was immunoprecipitated, and the co-immunoprecipitated sirtuins were probed through immunoblotting.

**f**, In silico modelling of LCA bound to TULP3. Modelling was performed according to the results of MS on purified TULP3 conjugated to the LCA-probe, and the AlphaFold-predicted TULP3 structure (https://alphafold.ebi.ac.uk/entry/O75386; coloured in green). The Y193, P195, K333 and P336 residues (coloured in yellow) that comprise a hydrophobic pocket for LCA (coloured in red) binding as indicated were mutated to glycine (TULP3-4G) to create a TULP3 mutant that is defective in binding to LCA.

**g**, **h**, TULP3-4G, unable to bind LCA, blocked the LCA-mediated activation of AMPK. *TULP3*$^{-/-}$ MEFs were infected with lentivirus carrying TULP3-4G mutant or its wildtype control, followed by treatment with 1 μM LCA for 4 h. The activity of v-ATPase (**g**) and the lysosomal translocation of AXIN (**h**) were then determined. The statistical analysis data of **g** and **h** are shown in Fig. 4k and 4l.

Experiments in this figure were performed three times.

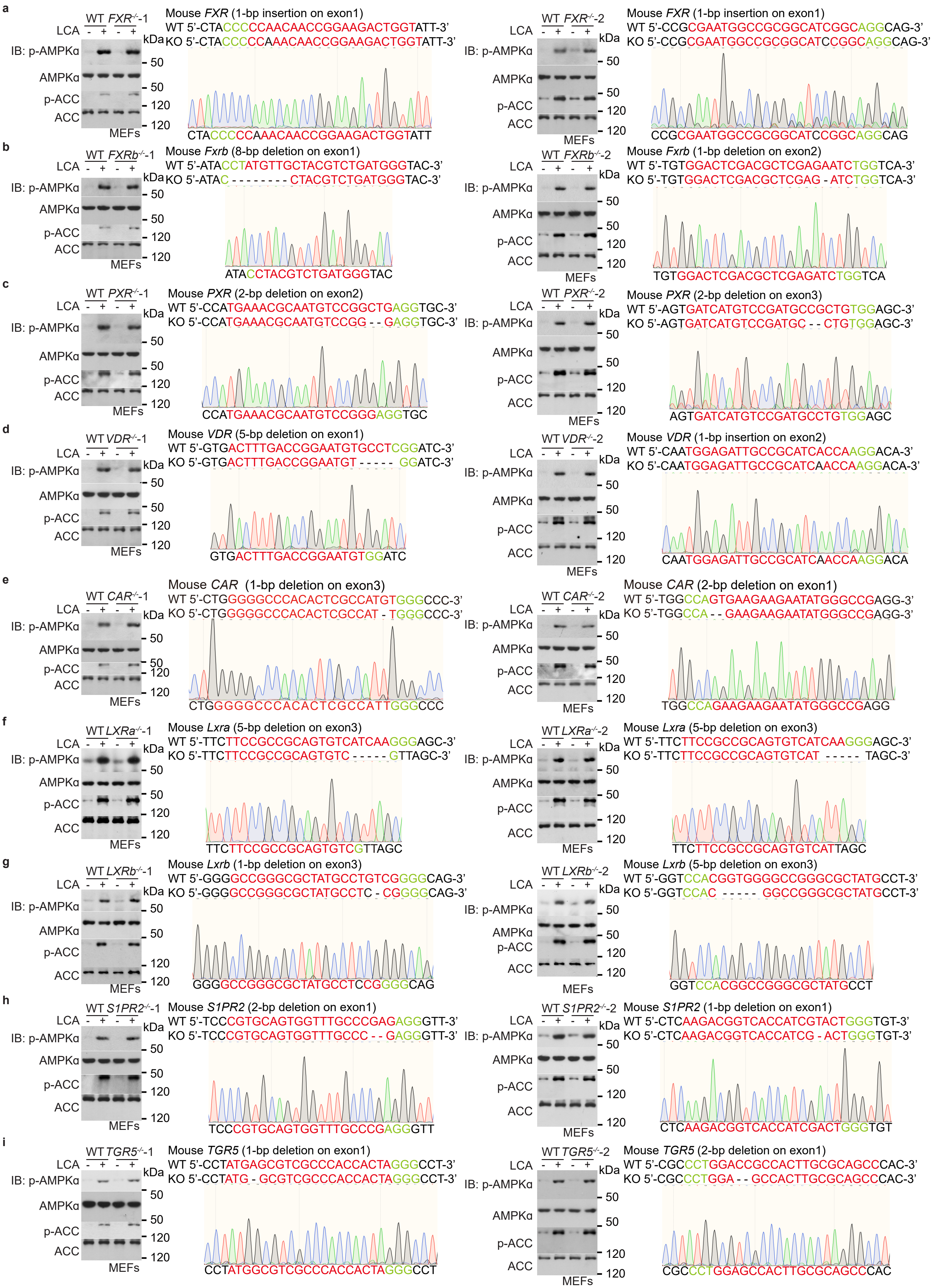


**Extended Data Fig. 7**

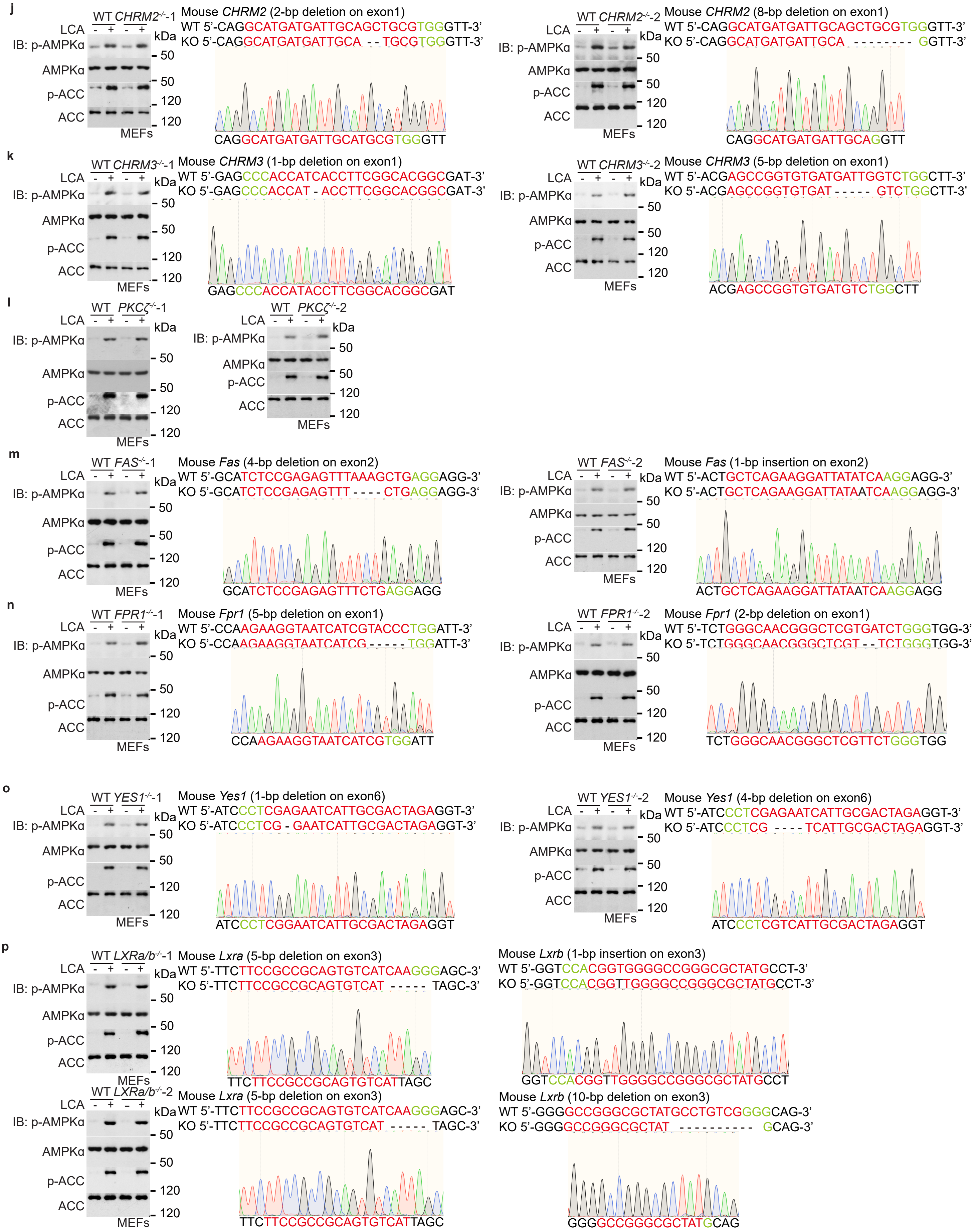


**Extended Data Fig. 7 (cont.)**

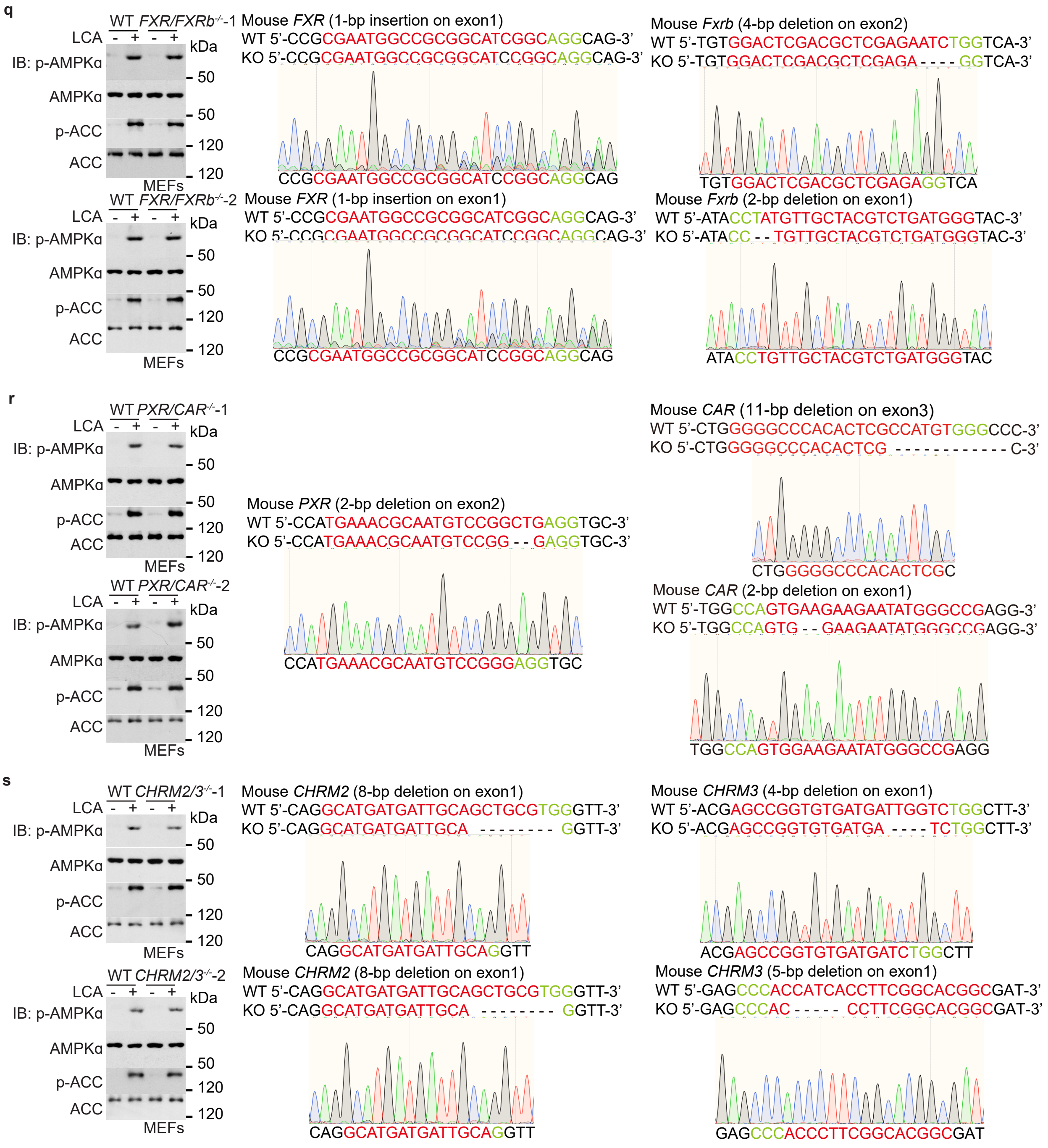


**Extended Data Fig. 7 | Other known binding partners of LCA are not required for the activation of AMPK.**
**a**-**s**, MEFs with *FXR* (**a**, clone #1 on the left or upper panel, and clone #2 right or lower, and the same hereafter in this figure), *FXRb* (**b**), *PXR* (**c**), *VDR* (**d**), *CAR* (**e**), *LXRa* (**f**), *LXRb* (**g**), *S1PR2* (**h**), *TGR5* (**i**), *CHRM2* (**j**), *CHRM3* (**k**), *PKCζ* (**l**), *FAS* (**m**), *FPR* (**n**), or *YES* (**o**) knockout, or with *LXRa* and *LXRb* (**p**), *FXRa* and *FXRb* (**q**), *PXR* and *CAR* (**r**), or *CHRM2* and *CHRM3* (**s**) double knockout, were treated with 1 μM LCA for 4 h, followed by determining the activity of AMPK. Experiments in this figure were performed three times.

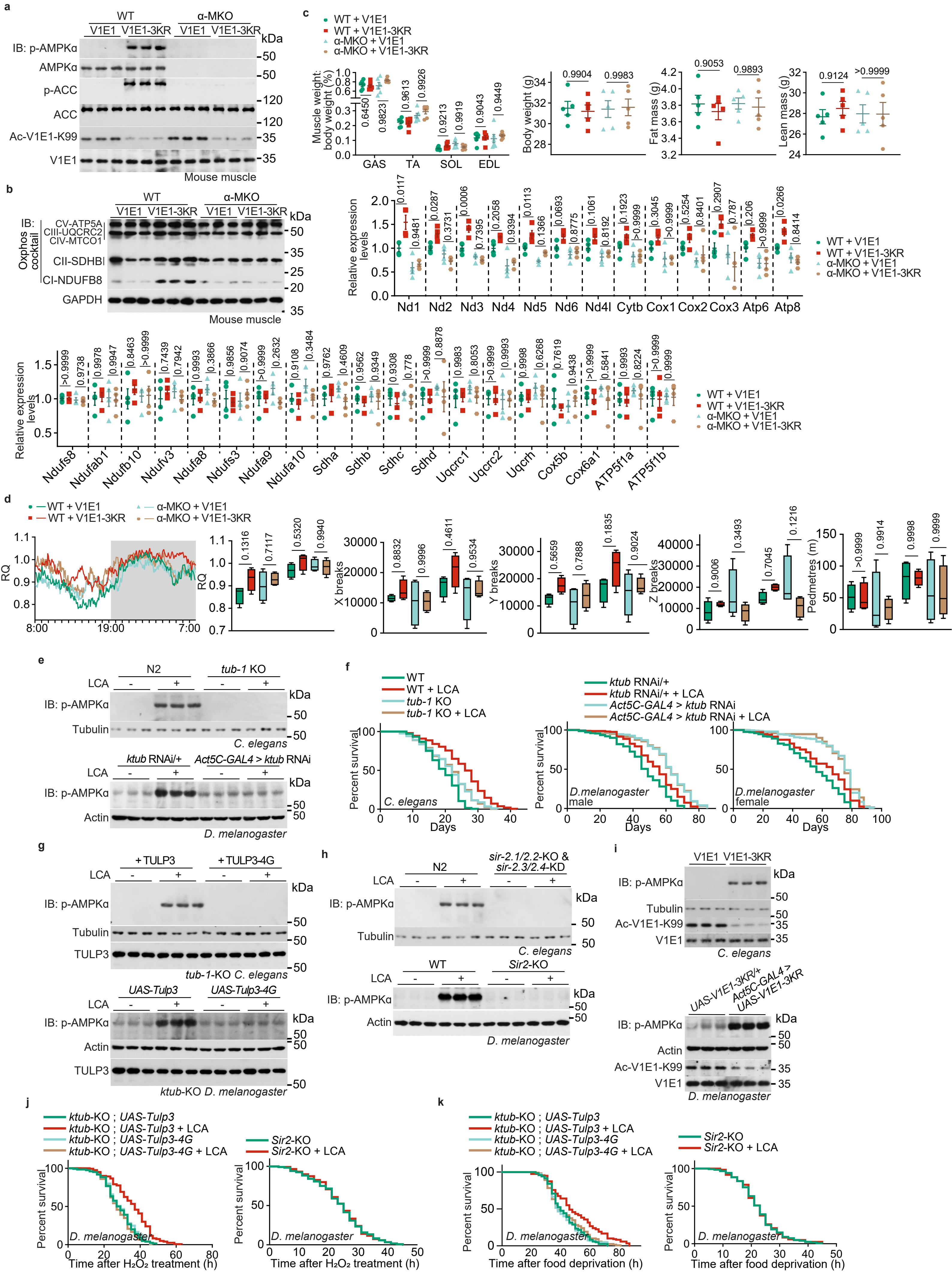


**Extended Data Fig. 8**

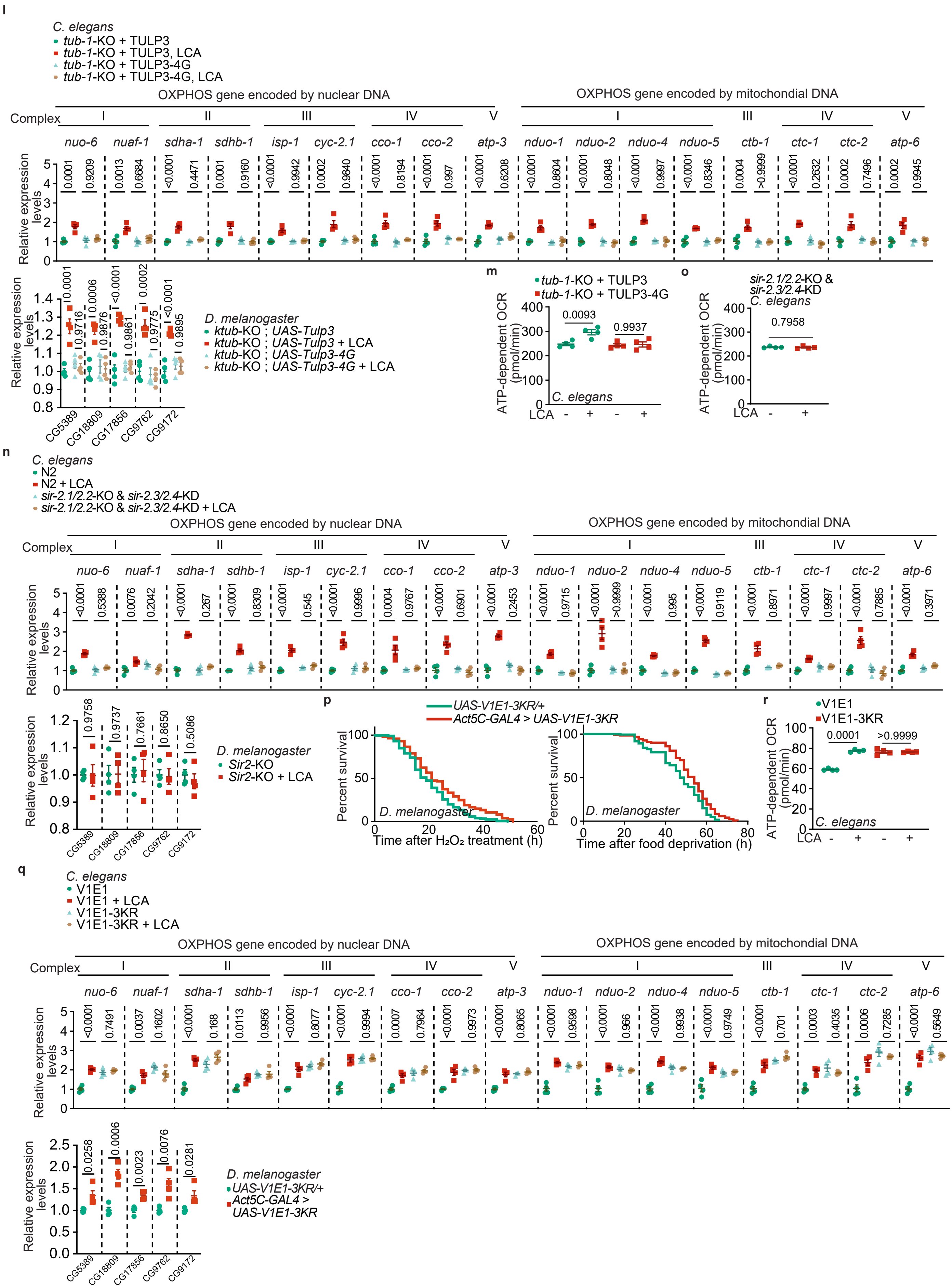


**Extended Data Fig. 8 (cont.)**

**s**

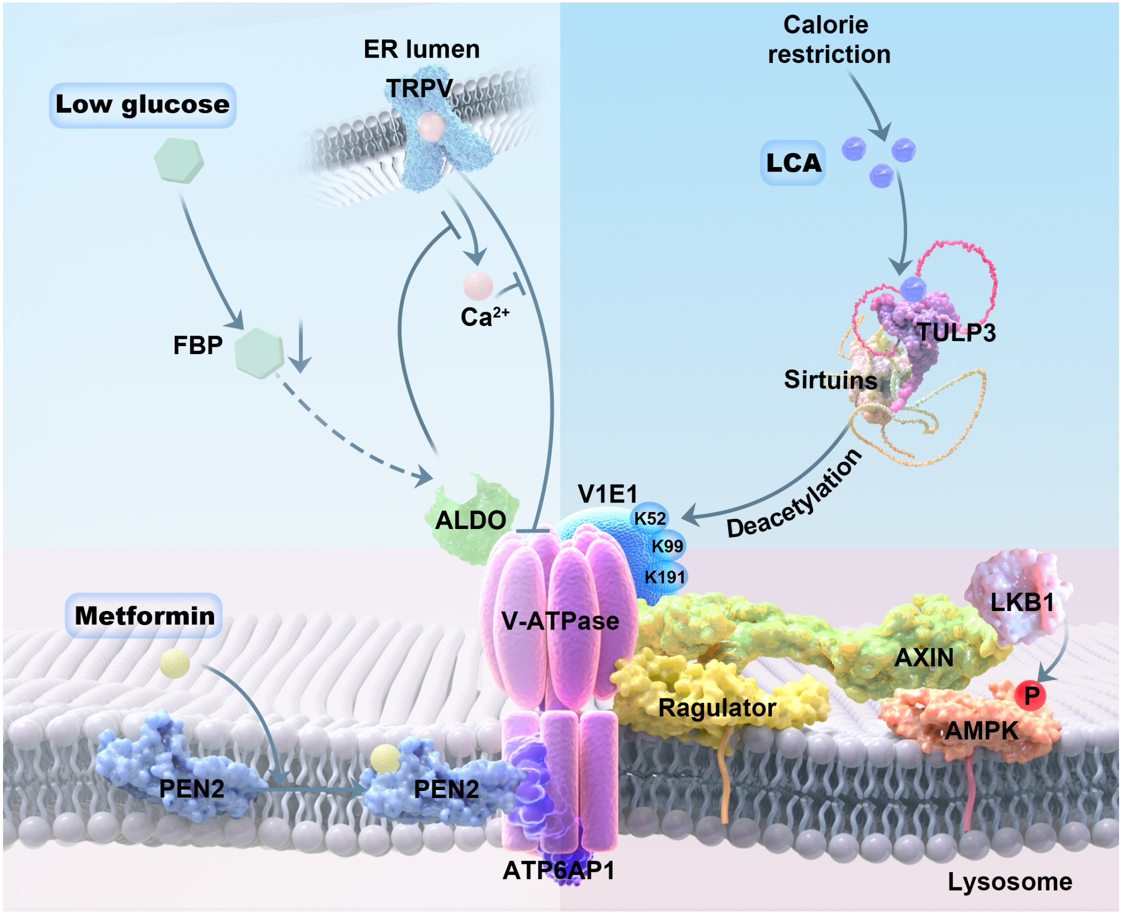


**Extended Data Fig. 8 | The LCA-TULP3-sirtuins-v-ATPase axis retards ageing**.

**a-d**, V1E1-3KR improves muscle function in aged mice. WT or muscle-specific *AMPKα* knockout (α-MKO) mice with muscle V1E1-3KR or wildtype V1E1 expression (induced by tamoxifen at 18 months old, for 4 weeks) were subjected to the analysis of muscular AMPK activation (**a**), the mRNA and protein levels of the OXPHOS complex (**b**), the body composition (**c**), and the RQ and ambulatory activity (**d**). Data in **b** are shown as mean ± s.e.m., normalised to the WT + V1E1 group ($n$ = 4 mice for each genotype, and $P$ value by two-way ANOVA followed by Tukey's test); in **c** as mean ± s.e.m. ($n$ = 5 mice for each genotype, and $P$ value by two-way ANOVA followed by Tukey's test); and in **d** as mean (at 5-min intervals during a 24-h course; $n$ = 4 mice for each genotype), and the others as box-and-whisker plots ($n$ = 4 mice for each genotype, and $P$ value by two-way ANOVA followed by Tukey's test).

**e-i**, The TULP3-sirtuins-v-ATPase axis is required for LCA to activate AMPK and extend lifespan in nematodes and flies. Nematodes and flies, either with *TULP3* knockout (upper panel of **e**, and left panel of **f**) or knockdown (*Act5C-GAL4* > *ktub* RNAi; lower panel of **e**, and right panel of **f**), with TULP3-4G reintroduction into the *TULP3*-knockout background (**g**), with sirtuin depletion (**h**), or with V1E1-3KR expression (**i**) were cultured in the medium containing LCA at 100 μM, followed by determining the activation of AMPK (**e**, **g**, **h**, **i**) and lifespan (**f**; shown as Kaplan-Meier curves).

**j**, **k**, **p**, The TULP3-sirtuins-v-ATPase axis is required for LCA to promote resistance to oxidative stress and starvation. Flies, either with TULP3-4G reintroduction into the *TULP3*-knockout background (left panel of **j**, **k**), with sirtuin depletion (right panel of **j**, **k**), or with V1E1-3KR expression (**p**) were cultured in the medium containing LCA at 100 μM, followed by transferring to media containing 5% $H_2O_2$ (**j**, and left panel of **p**) or deprived of food (**k**, and right panel of **p**). Lifespan data are shown as Kaplan-Meier curves.

**l-o**, **q**, **r**, The TULP3-sirtuins-v-ATPase axis elevates mitochondrial contents and respiratory functions in nematodes and flies. Nematodes and flies with TULP3-4G re-introduction in the background of *TULP3*-knockout (**l**, **m**), with sirtuins depletion (**n**, **o**), or with V1E1-3KR expression (**q**, **r**) were treated with LCA, followed by determining the mRNA levels of OXPHOS complex (**l**, **n**, **q**; results are mean ± s.e.m.; $n$ = 4 samples for each genotype/treatment, and $P$ value by two-way ANOVA followed by Tukey's test, except those of flies in **n**, **q** two-sided Student's $t$-test), and OCR (**m**, **o**, **r**; results are mean ± s.e.m.; $n$ = 4 samples for each genotype/treatment, and $P$ value by two-way ANOVA followed by Tukey's test, except **o** two-sided Student's $t$-test).

**s**, Schematic diagram showing that the v-ATPase serves as a common entry into the lysosomal AMPK pathway. LCA, elevated after CR, binds to TULP3 and activates sirtuins, which in turn deacetylate the V1E1 subunit of v-ATPase (on K52, K99 and K191 residues) and inhibits v-ATPase, which along with Ragulator allows for AXIN/LKB1 to translocate to the surface of the lysosome, where LKB1 phosphorylates and activates AMPK. In response to low glucose, the FBP-unoccupied aldolase interacts and inhibits the cation channel TRPV, which in turn interacts and reconfigures v-ATPase, allowing for AXIN/LKB1 to translocate to the lysosome to activate AMPK. Likewise, metformin binds to its target PEN2, and the metformin-bound PEN2 is recruited to v-ATPase via interacting with ATP6AP1, thereby inhibiting v-ATPase. The v-ATPase hence acts as a common node for the signalling of low glucose, metformin and LCA to intersect, all leading to AMPK activation and manifestation of anti-aging effects.

Experiments in this figure were performed three times.

| Subunit | Acetylated sites |
|---|---|
| **V1 domain** | |
| V1A | K14, K84, K132, K139, K149, S152, K155, K157, K200, K220, K256, K265, K301, S304, K307, K393, K443, K456, K467, K480, K498, K506, K513, K516, K564, K585, K587, K591, K596, K598 |
| V1B2 | K48, K108, K109, K430 |
| V1C1 | K21, K28, K37, K70, K84, K111, K115, K119, K137, K147, K156, K232, K245, K259, K262, K274, K325 |
| V1C2 | K28, K36, K83, K85, K114, K126, K136, K146, K154, K155, K213, K219, K231, K237, K239, K241, K244, K255, K271, K272, K324 |
| V1D | K24, K89, K93, K133, K215, K240 |
| V1E1 | K52, K59, K60, K68, K69, K99, K104, K138, K145, K150, K156, K160, K189, K191, K223 |
| V1E2 | K52, K59, K60, K69, K99, K118, K191, K207 |
| V1F | K6, K30, K41, K94, K102 |
| V1G1 | K16, K21, K27, K34, K37, K53, K56, K58, K75, K80 |
| V1G2 | K16, K21, K27, K34, K37, K58 |
| V1G3 | K21, K34, K37, K61, K80, K107 |
| V1H | K22, K30, K56, K62, K75, K86, K119, K147, K153, K169, K176, K234, K278, K280, K295, K313, K316, K325, K336, K341, K360, K371, K374, K386, K392, K396, K434, K458, K469 |
| **V0 domain** | |
| V0A4 | K38, K50, K83, K102, K118, K128, K133, K160, K169, K200, K214, K219, K234, K237, K298, K301, K303, K336, K360, K371, K436, K472, K532, K540, K542, K668, K816, K824, K831 |
| V0A2 | K32, K50, K56, K92, K103, K113, K115, K117, K120, K172, K184, K190, K197, K237, K239, K240, K279, K290, K303, K304, K306, K322, K363, K374, K474, K499, K500, K551, K553, K579, K580, K608, K637, K667, K693, K828, K836, K848 |
| V0B | ND |
| V0C | K5, K36, K54, K88 |
| V0D1 | K24, K46, K77, K128, K195, K234, K239, K265, K275, K288, K300, K303, K322, K343 |
| V0D2 | K24, K46, K69, K76, K119, K120, K123, K128, K184, K187, K195, K195, K234, K246, K265, K275, K287, K319, K321 |
| V0E1 | ND |
| AP1 | ND |
| AP2 | K238, K246 |

**Extended Data Table 1 | A summary of acetylated residues of v-ATPase**. The table contains the acetylated residues for each of the 21 subunits of v-ATPase (ATP6V1A, ATP6V1B2, ATP6V1C1, ATP6V1C2, ATP6V1D, ATP6V1E1, ATP6V1E2, ATP6V1F, ATP6V1G1, ATP6V1G2, ATP6V1G3, ATP6V1H, ATP6V0A4, ATP6V0A2, ATP6V0B, ATP6V0C, ATP6V0D1, ATP6V0D2, ATP6V0E1, ATP6AP1 and ATP6AP2) detected.

**Extended Data Table 2 | Summary of lifespan and analysis in worms[a,b]**

| Genotypes/ treatments | Mean life span (days): Estimated life span ± s.e.m. | Mean: 95% confidence interval Lower bound | Mean: 95% confidence interval Upper bound | Median life span (days): Estimated life span ± s.e.m. | Median: 95% confidence interval Lower bound | Median: 95% confidence interval Upper bound | N[c] | N[d] | N[e] | P-value Vs saline control within each genotype (Mantel-CoX) |
|---|---|---|---|---|---|---|---|---|---|---|
| | | | | | Fig. 5h | | | | | |
| *tub-1-KO* + TULP3 | 18.785 ± 0.395 | 18.010 | 19.559 | 18.000 ±0.770 | 16.491 | 19.509 | 184 | 16 | 200 | N/A |
| *ttub-1-KO* + TULP3, LCA | 22.886 ± 0.531 | 21.846 | 23.925 | 24.000 ±0.596 | 22.833 | 25.167 | 187 | 13 | 200 | <0.001 |
| *tub-1-KO* + TULP3-4G | 18.931 ± 0.394 | 18.160 | 19.703 | 20.000 ±0.964 | 18.111 | 21.889 | 186 | 14 | 200 | N/A |
| *tub-1-KO* + TULP3-4G, LCA | 19.188 ± 0.406 | 18.393 | 19.983 | 20.000 ±0.683 | 18.660 | 21.340 | 187 | 13 | 200 | 0.536 |
| | | | | | Fig. 5i | | | | | |
| *sir-2.1/2.2*-KO & *sir-2.3/2.4*-KD | 18.842 ± 0.391 | 18.076 | 19.608 | 18.000 ±0.697 | 16.633 | 19.367 | 188 | 12 | 200 | N/A |
| *sir-2.1/2.2*-KO & *sir-2.3/2.4*-KD + LCA | 18.785 ± 0.394 | 18.012 | 19.558 | 18.000 ±0.734 | 16.562 | 19.438 | 186 | 14 | 200 | 0.971 |
| | | | | | Fig. 5j | | | | | |
| V1E1 | 22.224 ± 0.508 | 21.228 | 23.220 | 24.000 ±0.632 | 22.761 | 25.239 | 184 | 16 | 200 | N/A |
| V1E1 + LCA | 24.306 ± 0.540 | 23.247 | 25.365 | 26.000 ±0.734 | 24.562 | 27.438 | 184 | 16 | 200 | <0.001 |
| V1E1-3KR | 24.794 ± 0.543 | 23.730 | 25.858 | 26.000 ±0.559 | 24.905 | 27.095 | 186 | 14 | 200 | N/A |
| V1E1-3KR + LCA | 24.776 ± 0.534 | 23.730 | 25.822 | 26.000 ±0.785 | 24.462 | 27.538 | 183 | 17 | 200 | 0.866 |
| | | | | | Extended Data Fig. 8f | | | | | |
| WT | 19.082 ± 0.375 | 18.347 | 19.816 | 20.000 ±0.761 | 18.508 | 21.492 | 183 | 17 | 200 | N/A |
| WT + LCA | 25.459 ± 0.548 | 24.385 | 26.534 | 26.000 ±0.647 | 24.732 | 27.492 | 187 | 13 | 200 | <0.001 |
| *tub-1* KO | 21.132 ± 0.522 | 20.109 | 22.154 | 22.000 ±0.650 | 20.727 | 23.273 | 187 | 13 | 200 | N/A |
| *tub-1* KO+ LCA | 21.306 ± 0.504 | 20.317 | 22.294 | 22.000 ±0.586 | 20.852 | 23.148 | 187 | 13 | 200 | 0.961 |

**Extended Data Table 2 | Summary of healthspan and analysis in worms[a,b]**

| Genotypes/ treatments | Mean health span (hours): Estimated life span ± s.e.m. | Mean: 95% confidence interval Lower bound | Mean: 95% confidence interval Upper bound | Median health span (hours): Estimated life span ± s.e.m. | Median: 95% confidence interval Lower bound | Median: 95% confidence interval Upper bound | N[c] | N[d] | N[e] | P-value Vs saline control within each genotype (Mantel-CoX) |
|---|---|---|---|---|---|---|---|---|---|---|
| | | | | | Fig. 5k | | | | | |
| *tub-1-KO* + TULP3 | 9.967 ± 0.576 | 8.837 | 11.096 | 10.000 ± 1.073 | 7.897 | 12.103 | 52 | 8 | 60 | N/A |
| *ttub-1-KO* + TULP3, LCA | 13.104 ± 0.743 | 11.648 | 14.559 | 14.000 ± 1.345 | 11.363 | 16.637 | 53 | 7 | 60 | <0.001 |
| *tub-1-KO* + TULP3-4G | 9.436 ± 0.508 | 8.441 | 10.431 | 11.000 ± 0.770 | 9.490 | 12.510 | 54 | 6 | 60 | N/A |
| *tub-1-KO* + TULP3-4G, LCA | 9.134 ± 0.526 | 8.104 | 10.165 | 10.000 ± 1.001 | 8.038 | 11.962 | 55 | 5 | 60 | 0.84 |
| | | | | | Fig. 5l | | | | | |
| *sir-2.1/2.2*-KO & *sir-2.3/2.4*-KD | 9.183 ± 0.547 | 8.111 | 10.255 | 9.000 ± 1.140 | 6.766 | 11.234 | 48 | 12 | 60 | N/A |
| *sir-2.1/2.2*-KO & *sir-2.3/2.4*-KD + LCA | 8.835 ± 0.477 | 7.900 | 9.771 | 9.000 ± 0.703 | 7.623 | 10.377 | 50 | 10 | 60 | 0.276 |
| | | | | | Fig. 5q | | | | | |
| V1E1 | 13.652 ± 0.650 | 12.378 | 14.926 | 14.000 ± 1.188 | 11.671 | 16.329 | 54 | 6 | 60 | N/A |
| V1E1 + LCA | 18.304 ± 1.018 | 16.310 | 20.299 | 22.000 ± 1.786 | 18.499 | 25.501 | 54 | 6 | 60 | <0.001 |
| V1E1-3KR | 17.735 ± 0.972 | 15.829 | 19.640 | 21.000 ± 1.241 | 18.567 | 23.433 | 56 | 4 | 60 | N/A |
| V1E1-3KR + LCA | 17.801 ± 0.978 | 15.885 | 19.718 | 20.000 ± 1.211 | 17.626 | 22.374 | 57 | 3 | 60 | 0.024 |

[a]Independent repeats of each lifespan and healthspan experiment were performed. Data from representative experiments are shown.
[b]Data sets within each panel of this table were done in parallel and statistical analyses were done within the data set.
[c]Number of worms scored (death events).
[d]Number of worms censored.
[e]Total number of worms.

**Extended Data Table 2 | Summary of lifespan and analysis in flies[a,b]**

| Genotypes/ treatments | Mean life span (days) | | | Median life span (days) | | | | | | P-value Vs saline control within each genotype (Mantel-CoX) |
|---|---|---|---|---|---|---|---|---|---|---|
| | | 95% confidence interval | | | 95% confidence interval | | | | | |
| | Estimated life span ± s.e.m. | Lower bound | Upper bound | Estimated life span ± s.e.m. | Lower bound | Upper bound | N[c] | N[d] | N[e] | |
| **Fig. 5h male** | | | | | | | | | | |
| *ktub KO ; UAS-Tulp3* | 50.840 ± 1.044 | 48.793 | 52.887 | 53.000 ±0.914 | 51.208 | 54.792 | 200 | 0 | 200 | N/A |
| *ktub KO ; UAS-Tulp3*+LCA | 56.405 ± 1.009 | 54.428 | 58.382 | 59.000 ±1.445 | 56.169 | 61.831 | 200 | 0 | 200 | <0.001 |
| *ktub KO ; UAS-Tulp3-4G* | 49.055 ± 1.037 | 47.022 | 51.088 | 50.000 ±0.866 | 48.303 | 51.697 | 200 | 0 | 200 | N/A |
| *ktub KO ; UAS-Tulp3-4G*+LCA | 49.150 ± 1.027 | 47.137 | 51.163 | 50.000 ±0.866 | 48.303 | 51.697 | 200 | 0 | 200 | 0.945 |
| **Fig. 5h female** | | | | | | | | | | |
| *ktub KO ; UAS-Tulp3* | 55.390 ± 1.124 | 53.186 | 57.594 | 59.000 ±1.112 | 56.821 | 51.179 | 200 | 0 | 200 | N/A |
| *ktub KO ; UAS-Tulp3*+LCA | 62.475 ± 1.071 | 60.375 | 64.575 | 68.000 ±0.696 | 66.637 | 69.363 | 200 | 0 | 200 | <0.001 |
| *ktub KO ; UAS-Tulp3-4G* | 53.005 ± 1.101 | 50.847 | 55.163 | 56.000 ±1.248 | 53.555 | 58.445 | 200 | 0 | 200 | N/A |
| *ktub KO ; UAS-Tulp3-4G*+LCA | 52.595 ± 1.218 | 50.208 | 54.982 | 56.000 ±1.586 | 52.892 | 59.108 | 200 | 0 | 200 | 0.218 |
| **Fig. 5i male** | | | | | | | | | | |
| *Sir2*-KO | 52.185 ± 0.713 | 50.787 | 53.583 | 53.000 ±1.031 | 50.980 | 55.020 | 200 | 0 | 200 | N/A |
| *Sir2*-KO+LCA | 51.575 ± 0.758 | 50.090 | 53.060 | 53.000 ±0.901 | 51.234 | 54.766 | 200 | 0 | 200 | 0.385 |
| **Fig. 5i female** | | | | | | | | | | |
| *Sir2*-KO | 70.120 ± 1.104 | 67.955 | 72.285 | 74.000 ±0.701 | 72.626 | 75.375 | 200 | 0 | 200 | N/A |
| *Sir2*-KO+LCA | 69.515 ± 1.085 | 67.389 | 71.641 | 74.000 ±0.805 | 72.421 | 75.579 | 200 | 0 | 200 | 0.229 |
| **Fig. 5j male** | | | | | | | | | | |
| *UAS-V1E1-3KR/+* | 32.150 ± 0.713 | 30.753 | 33.547 | 33.000 ±0.785 | 31.462 | 34.538 | 200 | 0 | 200 | N/A |
| *Act5C-GAL4>UAS-V1E1-3KR* | 49.615 ± 1.100 | 47.458 | 51.772 | 52.000 ±1.367 | 49.321 | 54.679 | 200 | 0 | 200 | <0.001 |
| **Fig. 5j female** | | | | | | | | | | |
| *UAS-V1E1-3KR/+* | 46.360 ± 1.124 | 44.156 | 48.564 | 52.000 ±0.467 | 51.084 | 52.916 | 200 | 0 | 200 | N/A |
| *Act5C-GAL4>UAS-V1E1-3KR* | 57.520 ± 1.056 | 55.450 | 59.590 | 62.000 ±0.914 | 60.208 | 63.792 | 200 | 0 | 200 | <0.001 |
| **Extended Data Fig. 8f male** | | | | | | | | | | |
| *ktub RNAi/+* | 46.815 ± 1.043 | 44.772 | 48.858 | 49.000 ±1.113 | 46.819 | 51.181 | 200 | 0 | 200 | N/A |
| *ktub RNAi/+* + LCA | 53.815 ± 1.004 | 51.848 | 55.782 | 56.000 ±1.534 | 52.994 | 59.006 | 200 | 0 | 200 | <0.001 |
| *Act5C-GAL4>ktub RNAi* | 61.120 ± 1.040 | 59.081 | 63.159 | 63.000 ±0.925 | 61.188 | 64.812 | 200 | 0 | 200 | N/A |
| *Act5C-GAL4>ktub RNAi* + LCA | 60.460 ± 1.125 | 58.255 | 62.665 | 63.000 ±0.982 | 61.076 | 64.924 | 200 | 0 | 200 | 0.913 |
| **Extended Data Fig. 8f female** | | | | | | | | | | |
| *ktub RNAi/+* | 53.360 ± 1.358 | 50.698 | 56.022 | 56.000 ±2.097 | 51.890 | 60.110 | 200 | 0 | 200 | N/A |
| *ktub RNAi/+* + LCA | 60.190 ± 1.439 | 57.370 | 63.010 | 67.000 ±2.856 | 61.402 | 72.598 | 200 | 0 | 200 | <0.001 |
| *Act5C-GAL4>ktub RNAi* | 71.650 ± 1.138 | 69.419 | 73.881 | 75.000 ±0.602 | 73.821 | 76.179 | 200 | 0 | 200 | N/A |
| *Act5C-GAL4>ktub RNAi* + LCA | 72.640 ± 1.083 | 70.517 | 74.763 | 75.000 ±0.782 | 73.468 | 76.532 | 200 | 0 | 200 | 0.584 |

[a]Independent repeats of each lifespan experiment were performed. Data from representative experiments are shown.
[b]Lifespan data sets within each panel of this table were done in parallel and statistical analyses were done within the data set.
[c]Number of flies scored (death events).
[d]Number of flies censored.
[e]Total number of flies.

**Extended Data Table 2 | Summary of healthspan and analysis in flies[a,b]**

| Genotypes/ treatments | Mean health span (hours) | | | Median health span (hours) | | | | | | |
|---|---|---|---|---|---|---|---|---|---|---|
| | Estimated life span ± s.e.m. | 95% confidence interval | | Estimated life span ± s.e.m. | 95% confidence interval | | | | | P-value Vs saline control within each genotype (Mantel-CoX) |
| | | Lower bound | Upper bound | | Lower bound | Upper bound | N[c] | N[d] | N[e] | |
| **Fig. 5k** | | | | | | | | | | |
| *ktub KO ; UAS-Tulp3* | 19.092 ± 0.401 | 18.306 | 19.878 | 18.000 ± 0.449 | 17.121 | 18.879 | 120 | 0 | 120 | N/A |
| *ktub KO ; UAS-Tulp3*+LCA | 22.242 ± 0.424 | 21.411 | 23.072 | 23.500 ± 0.671 | 22.185 | 24.815 | 120 | 0 | 120 | <0.001 |
| *ktub KO ; UAS-Tulp3-4G* | 17.604 ± 0.342 | 16.935 | 18.274 | 18.000 ± 0.285 | 17.442 | 18.558 | 120 | 0 | 120 | N/A |
| *ktub KO ; UAS-Tulp3-4G*+LCA | 18.017 ± 0.367 | 17.298 | 18.736 | 18.000 ± 0.413 | 17.190 | 18.810 | 120 | 0 | 120 | 0.367 |
| **Fig. 5l** | | | | | | | | | | |
| *Sir2*-KO | 31.783 ± 0.640 | 30.530 | 33.037 | 33.000 ± 0.755 | 31.519 | 34.481 | 120 | 0 | 120 | N/A |
| *Sir2*-KO+LCA | 31.467 ± 0.772 | 29.954 | 32.980 | 33.000 ± 1.147 | 30.751 | 35.249 | 120 | 0 | 120 | 0.404 |
| **Fig. 5q** | | | | | | | | | | |
| *UAS-V1E1-3KR/+* | 22.667 ± 0.585 | 21.521 | 23.813 | 25.000 ± 0.763 | 23.505 | 26.495 | 120 | 0 | 120 | N/A |
| *Act5C-GAL4>UAS-V1E1-3KR* | 38.408 ± 1.176 | 36.104 | 40.713 | 40.000 ± 1.965 | 36.149 | 43.851 | 120 | 0 | 120 | <0.001 |
| **Extended Data Fig. 8j** | | | | | | | | | | |
| *ktub KO ; UAS-Tulp3* | 28.525 ± 0.806 | 26.946 | 30.104 | 27.000 ± 1.461 | 24.137 | 29.863 | 120 | 0 | 120 | N/A |
| *ktub KO ; UAS-Tulp3*+LCA | 35.792 ± 0.975 | 33.880 | 37.704 | 37.000 ± 1.637 | 33.791 | 40.209 | 120 | 0 | 120 | <0.001 |
| *ktub KO ; UAS-Tulp3-4G* | 29.117 ± 0.800 | 27.548 | 30.685 | 29.000 ± 1.561 | 25.940 | 32.060 | 120 | 0 | 120 | N/A |
| *ktub KO ; UAS-Tulp3-4G*+LCA | 37.946 ± 0.731 | 26.513 | 29.378 | 27.000 ± 1.112 | 24.821 | 29.179 | 120 | 0 | 120 | 0.207 |
| **Extended Data Fig. 8j** | | | | | | | | | | |
| *Sir2*-KO | 23.817 ± 0.810 | 22.228 | 25.405 | 25.000 ± 1.029 | 22.984 | 27.016 | 120 | 0 | 120 | N/A |
| *Sir2*-KO+LCA | 24.167 ± 0.771 | 22.655 | 25.678 | 25.000 ± 1.034 | 22.973 | 27.027 | 120 | 0 | 120 | 0.920 |
| **Extended Data Fig. 8k** | | | | | | | | | | |
| *ktub KO ; UAS-Tulp3* | 43.004 ± 1.058 | 40.931 | 45.077 | 40.000 ± 1.880 | 36.314 | 43.686 | 120 | 0 | 120 | N/A |
| *ktub KO ; UAS-Tulp3*+LCA | 50.579 ± 1.570 | 47.502 | 53.656 | 46.000 ± 2.347 | 41.400 | 50.600 | 120 | 0 | 120 | <0.001 |
| *ktub KO ; UAS-Tulp3-4G* | 42.283 ± 1.064 | 40.199 | 44.368 | 38.000 ± 1.171 | 35.705 | 40.295 | 120 | 0 | 120 | N/A |
| *ktub KO ; UAS-Tulp3-4G*+LCA | 43.788 ± 1.211 | 41.413 | 46.162 | 38.000 ± 2.317 | 33.459 | 42.541 | 120 | 0 | 120 | 0.325 |
| **Extended Data Fig. 8k** | | | | | | | | | | |
| *Sir2*-KO | 23.817 ± 0.810 | 22.228 | 25.405 | 25.000 ± 1.029 | 22.984 | 27.016 | 120 | 0 | 120 | N/A |
| *Sir2*-KO+LCA | 24.167 ± 0.771 | 22.655 | 25.678 | 25.000 ± 1.034 | 22.973 | 27.027 | 120 | 0 | 120 | 0.920 |
| **Extended Data Fig. 8p** | | | | | | | | | | |
| *UAS-V1E1-3KR/+* | 20.333 ± 0.862 | 18.643 | 22.023 | 19.000 ± 1.313 | 16.427 | 21.573 | 120 | 0 | 120 | N/A |
| *Act5C-GAL4>UAS-V1E1-3KR* | 24.250 ± 1.088 | 22.118 | 26.382 | 21.000 ± 1.427 | 18.203 | 23.797 | 120 | 0 | 120 | 0.002 |
| **Extended Data Fig. 8p** | | | | | | | | | | |
| *UAS-V1E1-3KR/+* | 46.517 ± 1.092 | 44.377 | 48.657 | 47.000 ± 1.217 | 44.614 | 49.386 | 120 | 0 | 120 | N/A |
| *Act5C-GAL4>UAS-V1E1-3KR* | 51.567 ± 1.063 | 49.483 | 53.650 | 54.000 ± 1.429 | 51.200 | 56.800 | 120 | 0 | 120 | 0.001 |

[a]Independent repeats of each healthspan experiment were performed. Data from representative experiments are shown.
[b]Healthspan data sets within each panel of this table were done in parallel and statistical analyses were done within the data set.
[c]Number of flies scored (death events).
[d]Number of flies censored.
[e]Total number of flies.
