## Supplementary material for "Lithocholic acid targets TULP3 to activate sirtuins and AMPK to retard ageing": Uncropped gel images

#### Fig. 1b

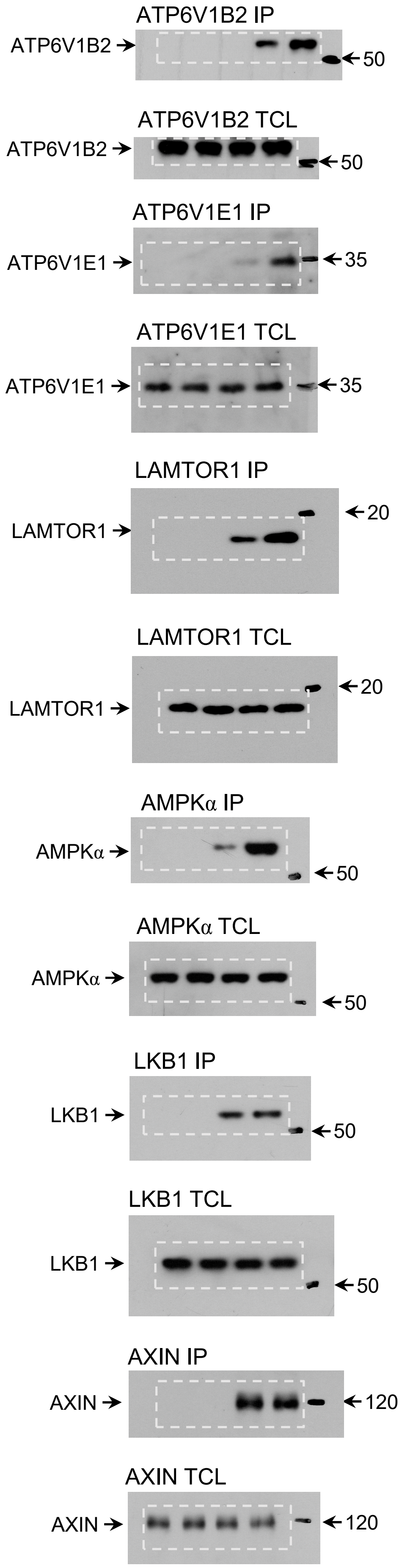


#### Fig. 1d

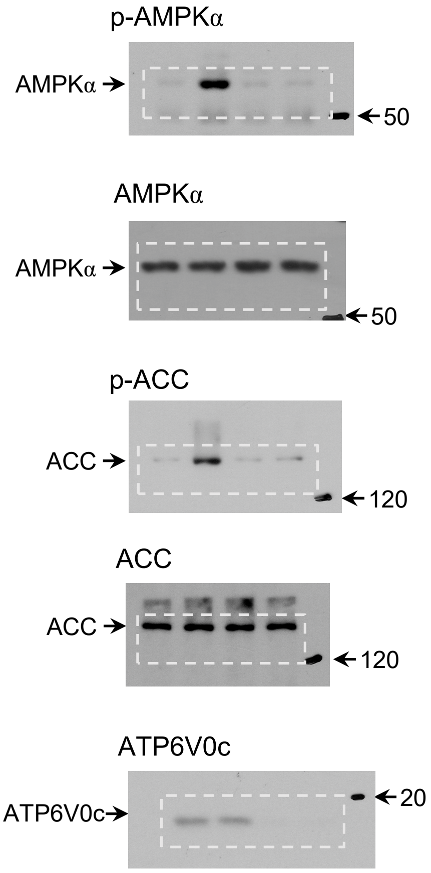


#### Fig. 1e

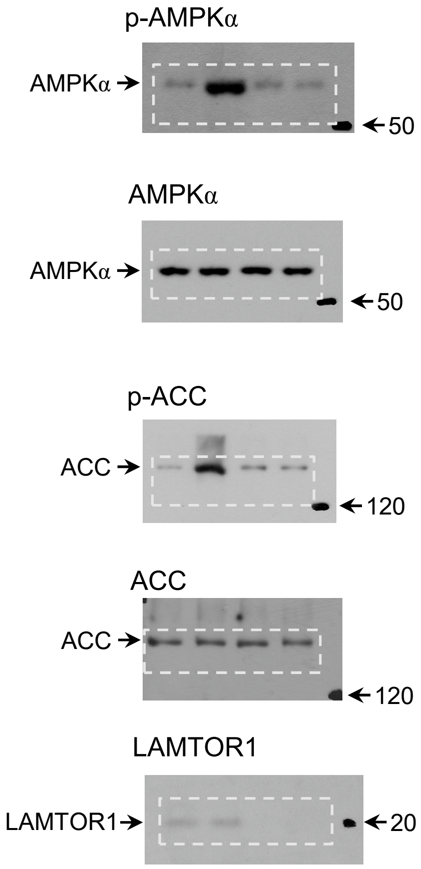

#### Fig. 1g

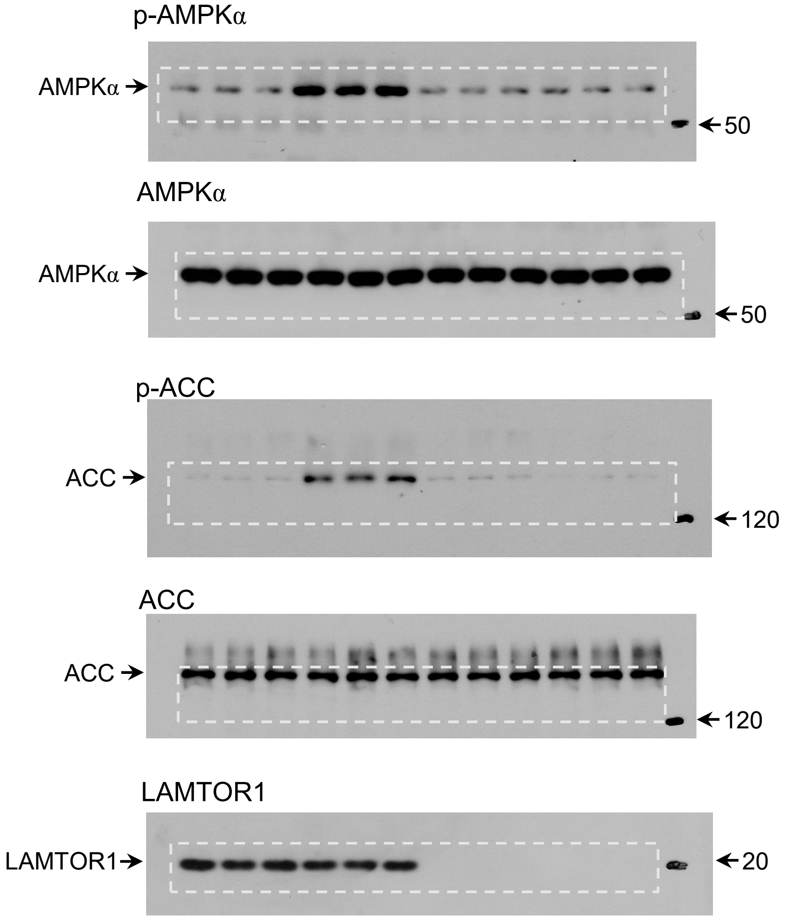
p-AMPKα
AMPKα
50
AMPKα
AMPKα
50
p-ACC
ACC
120
ACC
ACC
120
LAMTOR1
LAMTOR1
20

#### Fig. 1h

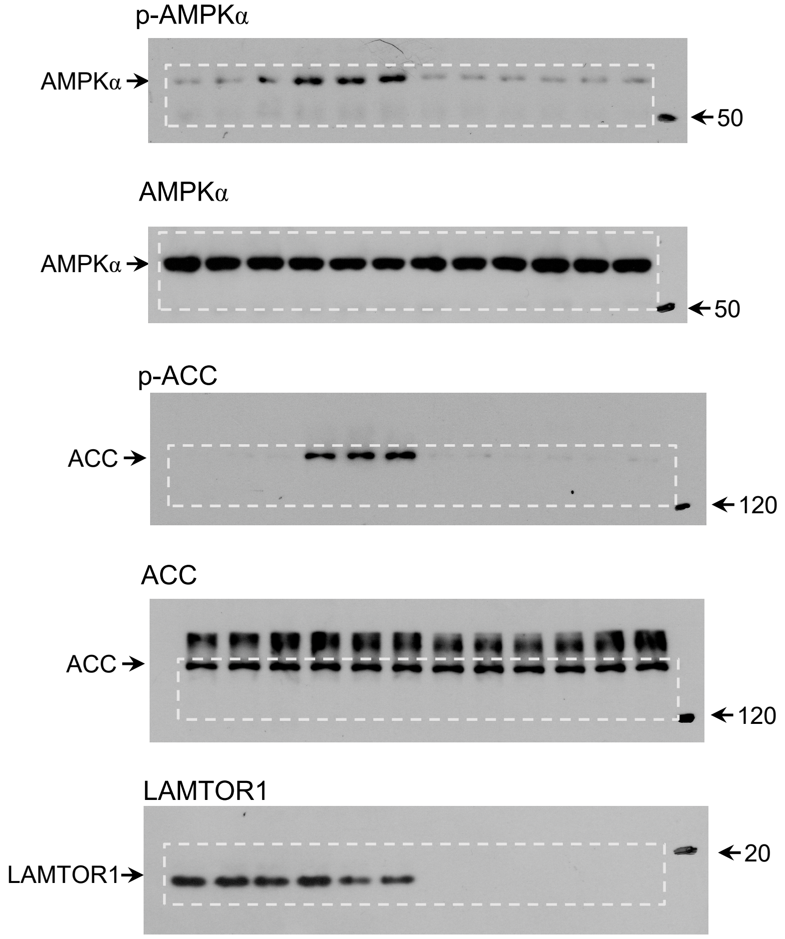
p-AMPKα
AMPKα
50
AMPKα
AMPKα
50
p-ACC
ACC
120
ACC
ACC
120
LAMTOR1
LAMTOR1
20

**Fig. 1i**

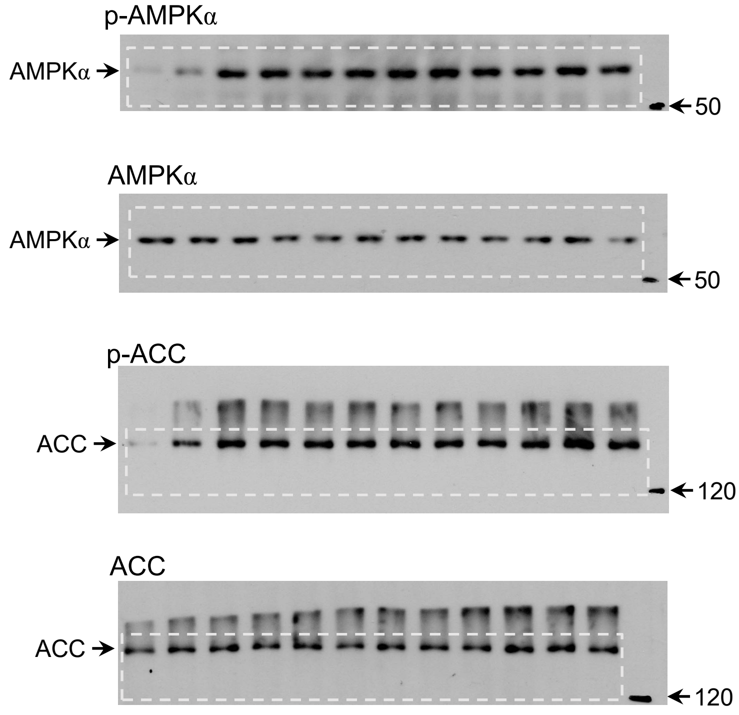


**Fig. 1j**

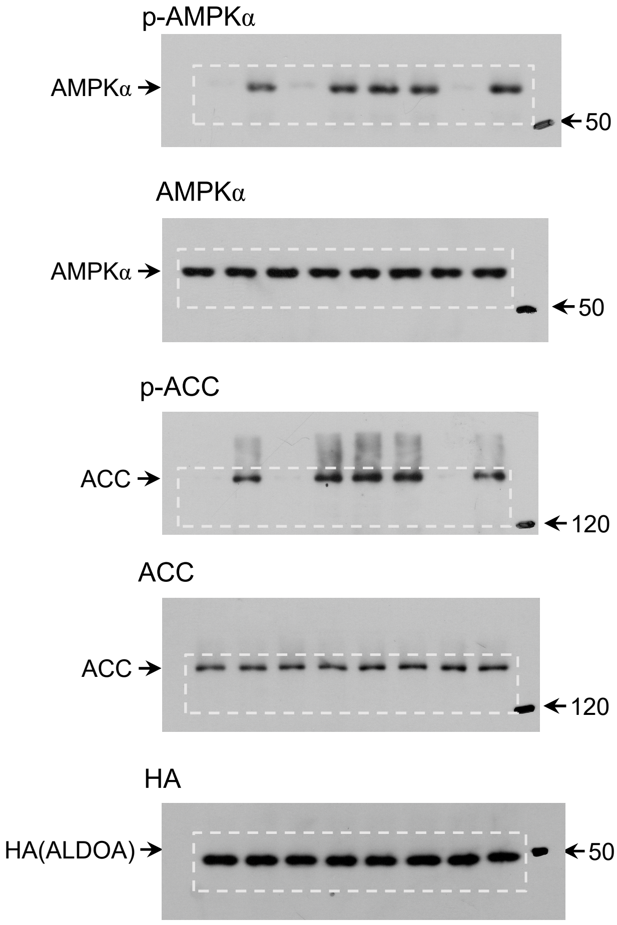


**Fig. 1m**

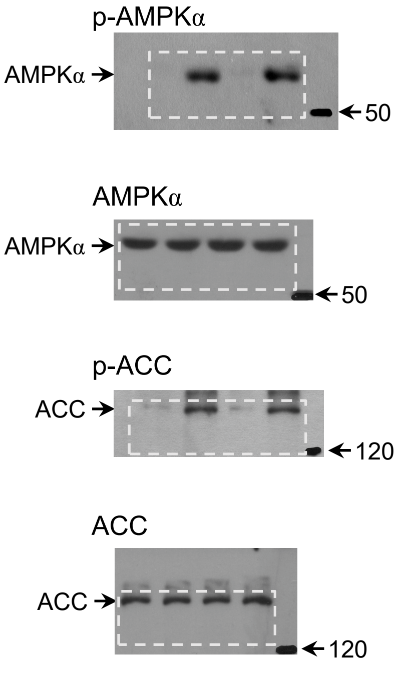

**Fig. 2a**

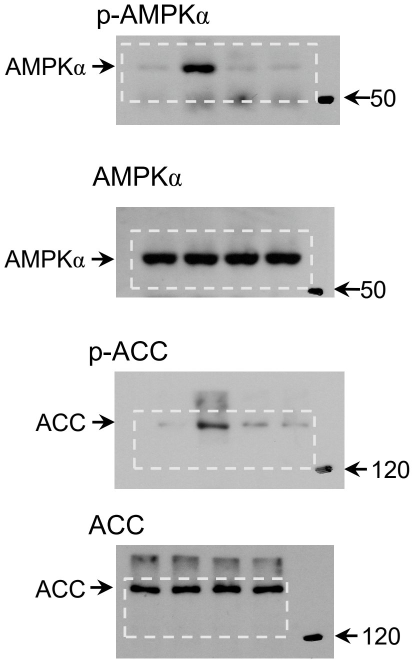


**Fig. 2b**

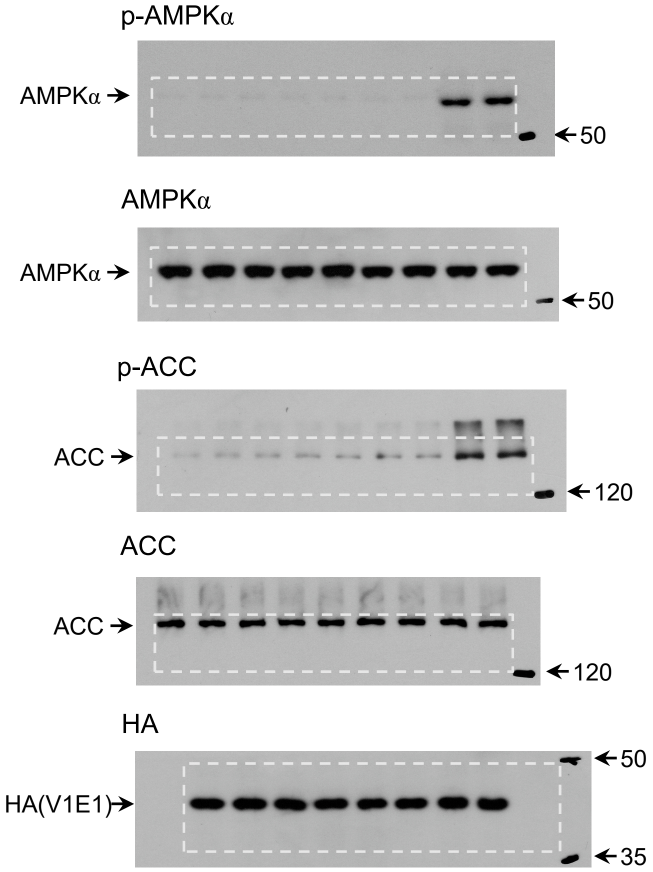


**Fig. 2e**

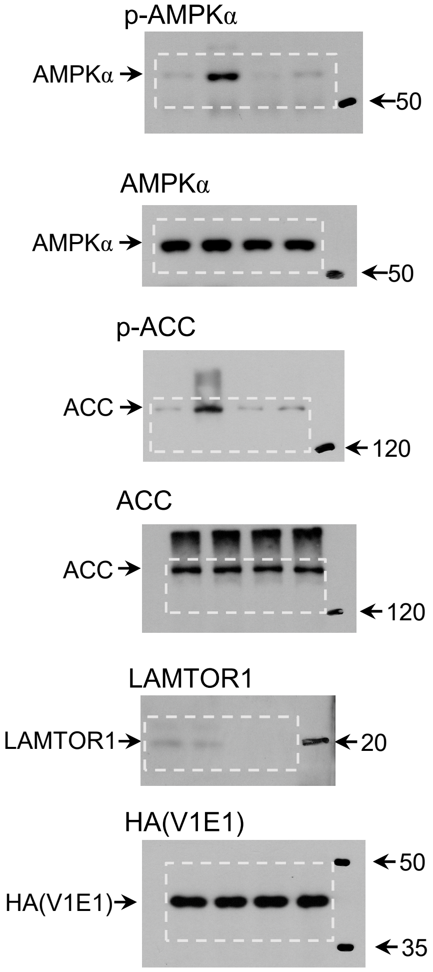

#### Fig. 2g

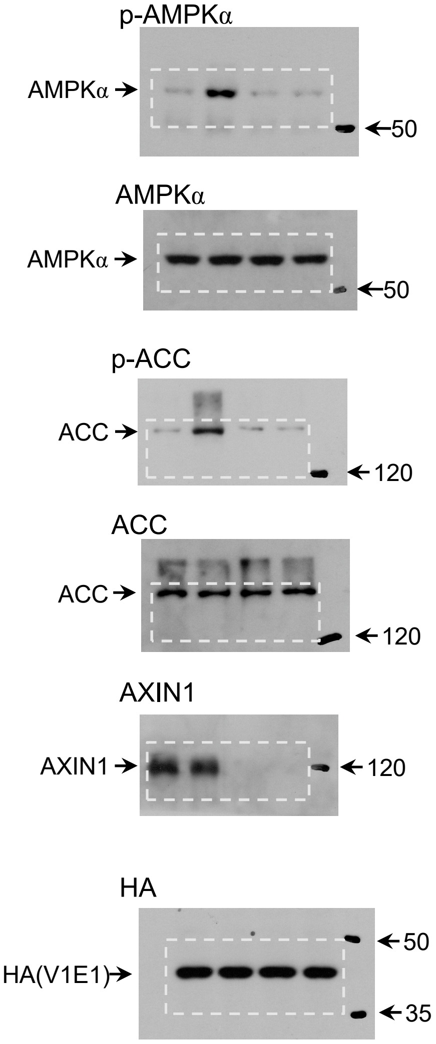


#### Fig. 2h

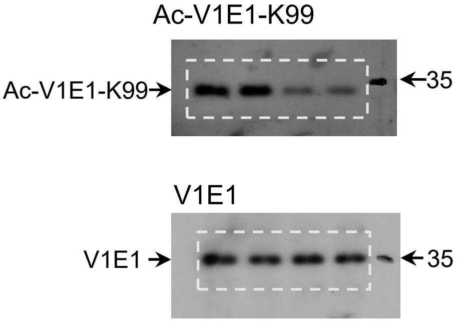


#### Fig. 2i

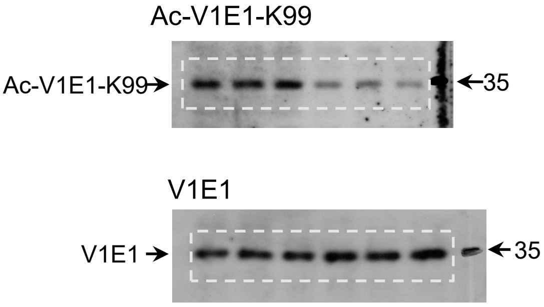

#### Fig. 2j

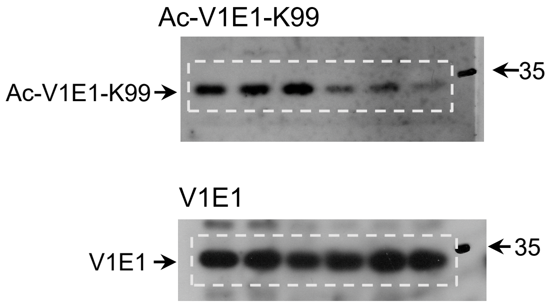


#### Fig. 2k

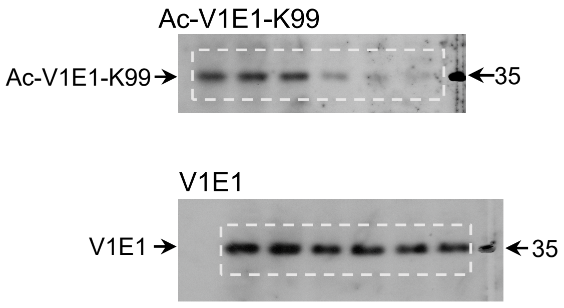


#### Fig. 2l

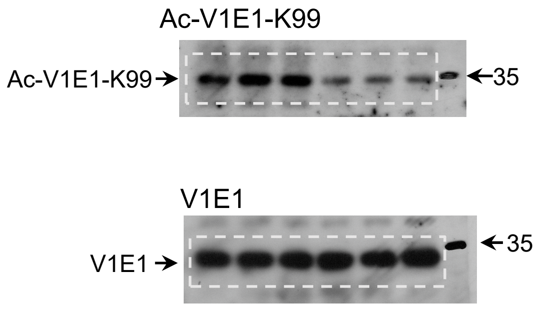


#### Fig. 2m

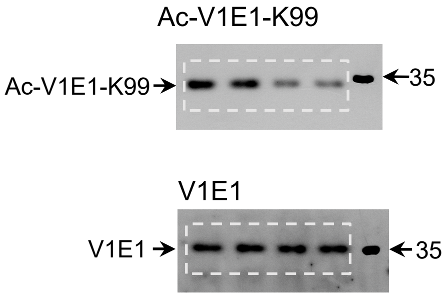

#### Fig. 3a (left)

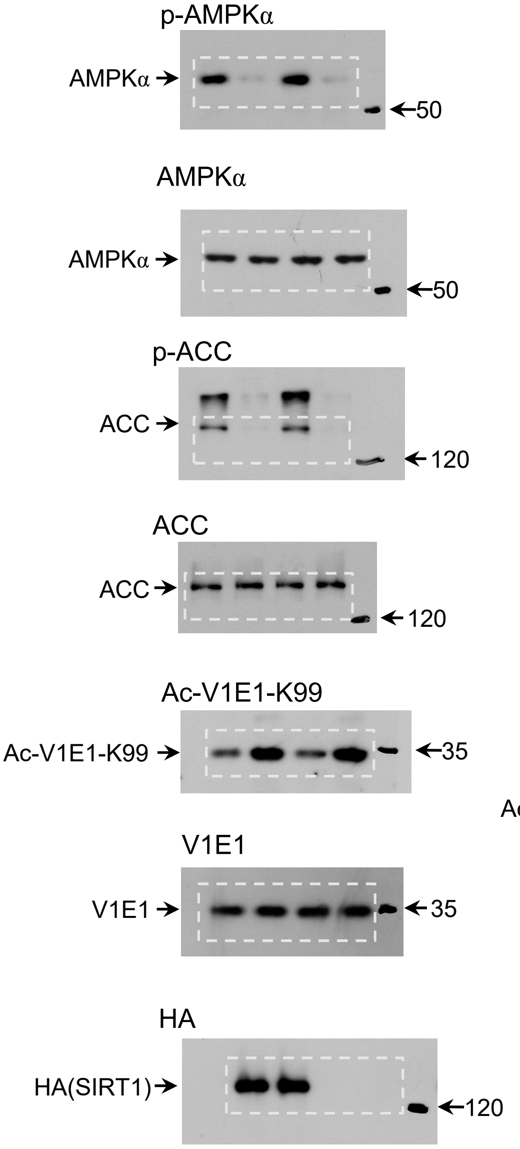


#### Fig. 3a (right)

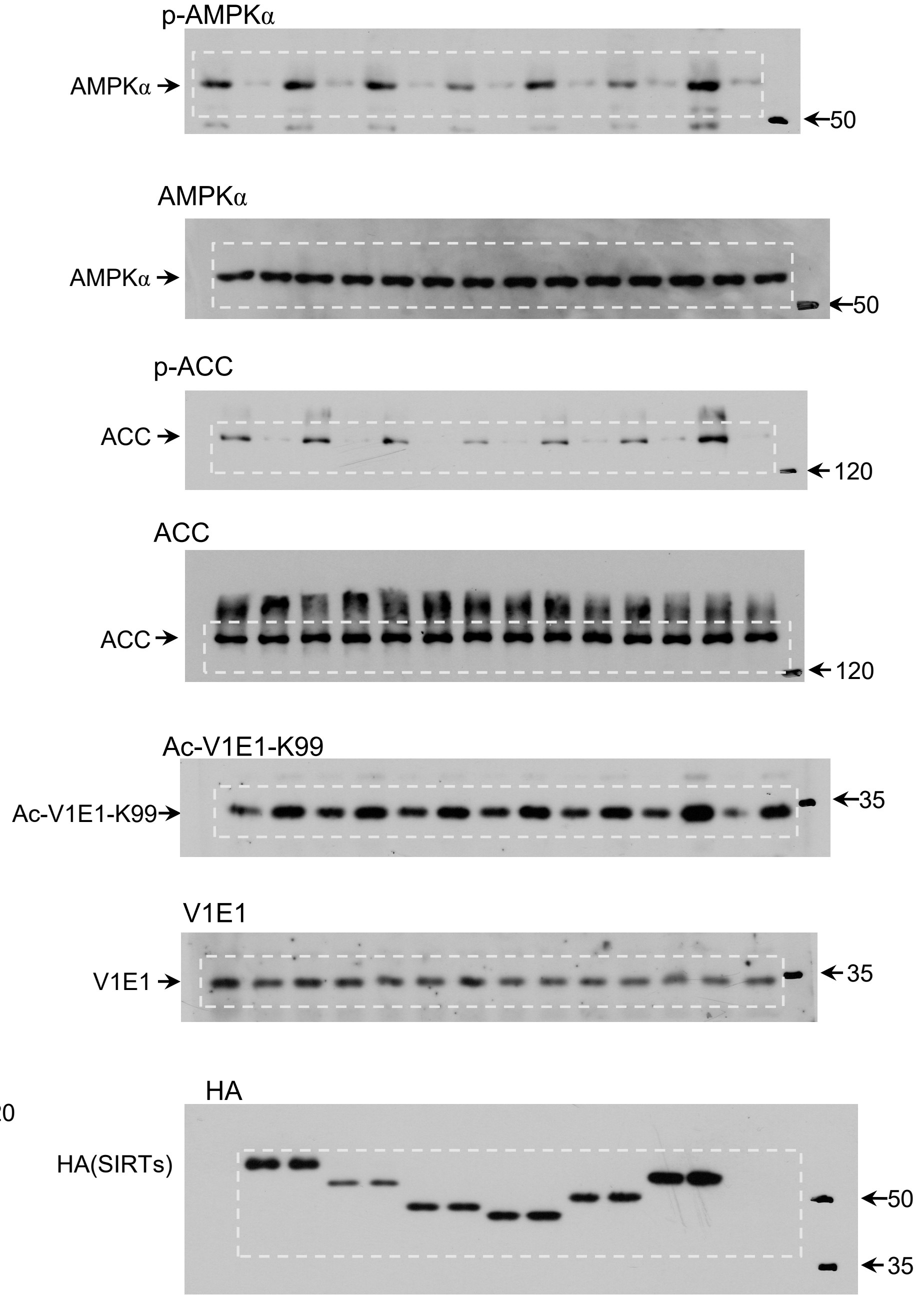

**Fig. 3c**

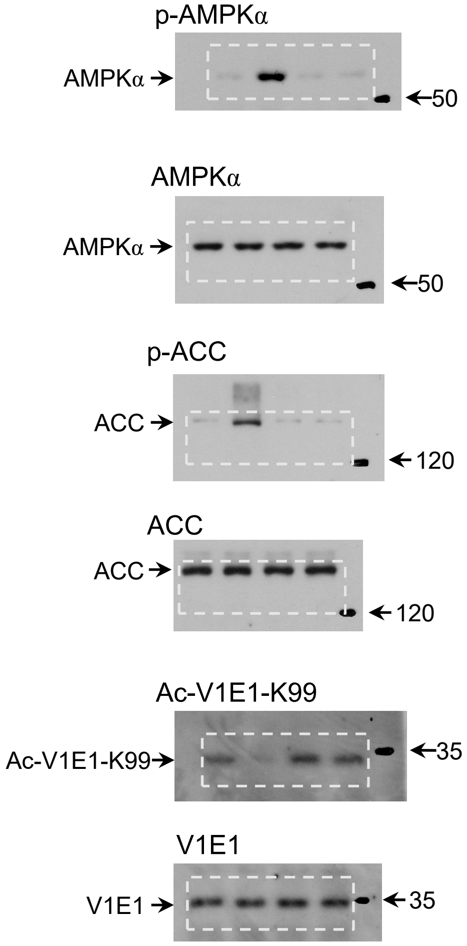
p-AMPKα
AMPKα
50
AMPKα
AMPKα
50
p-ACC
ACC
120
ACC
ACC
120
Ac-V1E1-K99
Ac-V1E1-K99
35
V1E1
V1E1
35

**Fig. 3f**

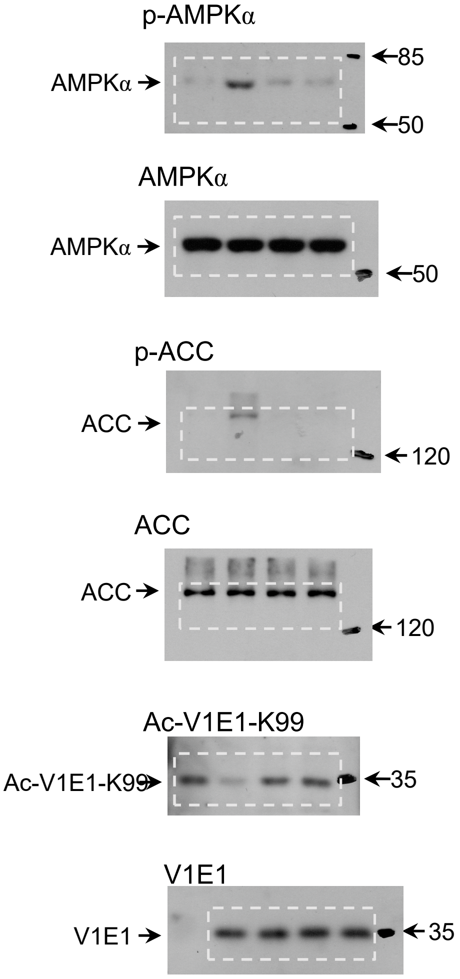
p-AMPKα
AMPKα
85
50
AMPKα
AMPKα
50
p-ACC
ACC
120
ACC
ACC
120
Ac-V1E1-K99
Ac-V1E1-K99
35
V1E1
V1E1
35

#### Fig. 3i

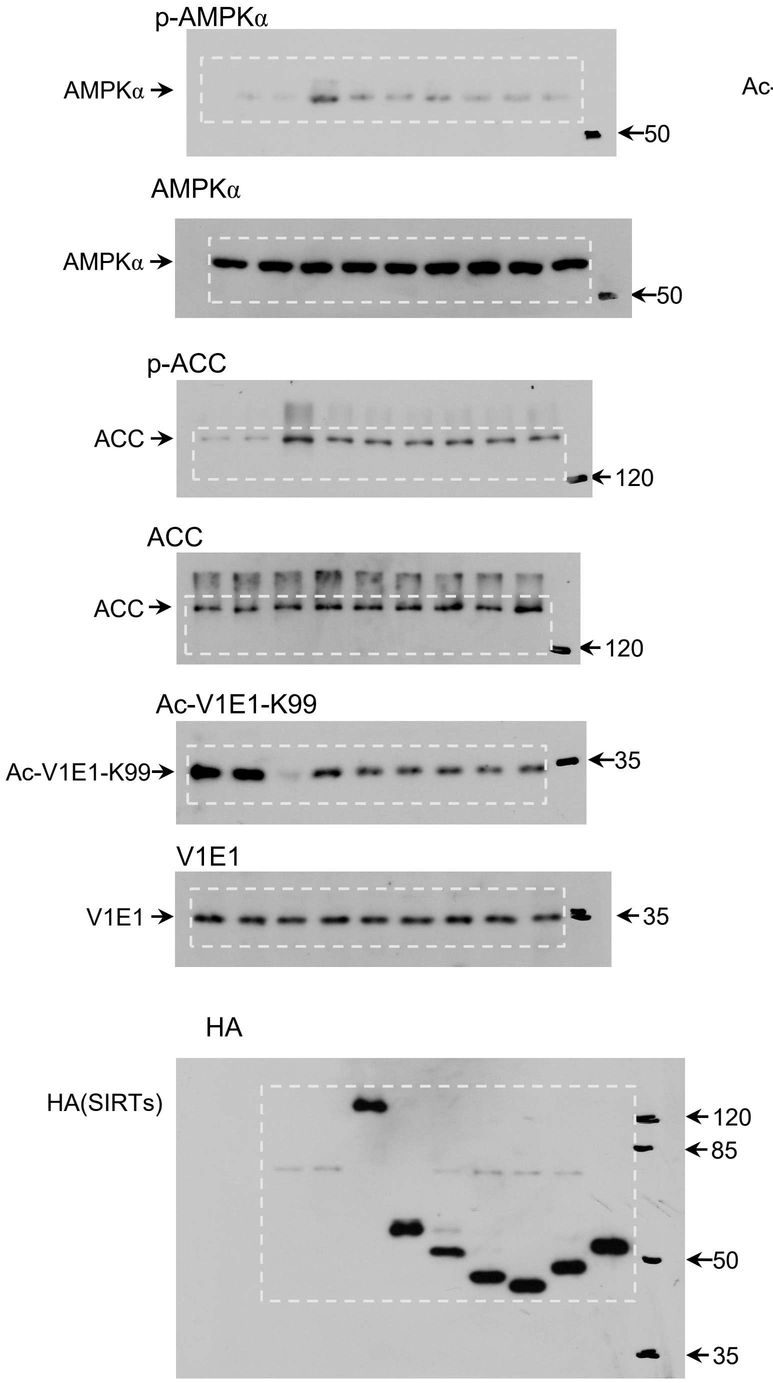
p-AMPKα
AMPKα
50
AMPKα
AMPKα
50
p-ACC
ACC
120
ACC
ACC
120
Ac-V1E1-K99
Ac-V1E1-K99
35
V1E1
V1E1
35
HA
HA(SIRTs)
120
85
50
35

#### Fig. 3j

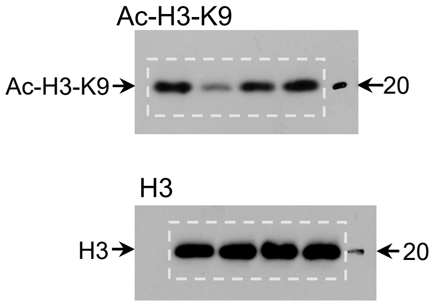
Ac-H3-K9
Ac-H3-K9
20
H3
H3
20

**Fig. 3l**

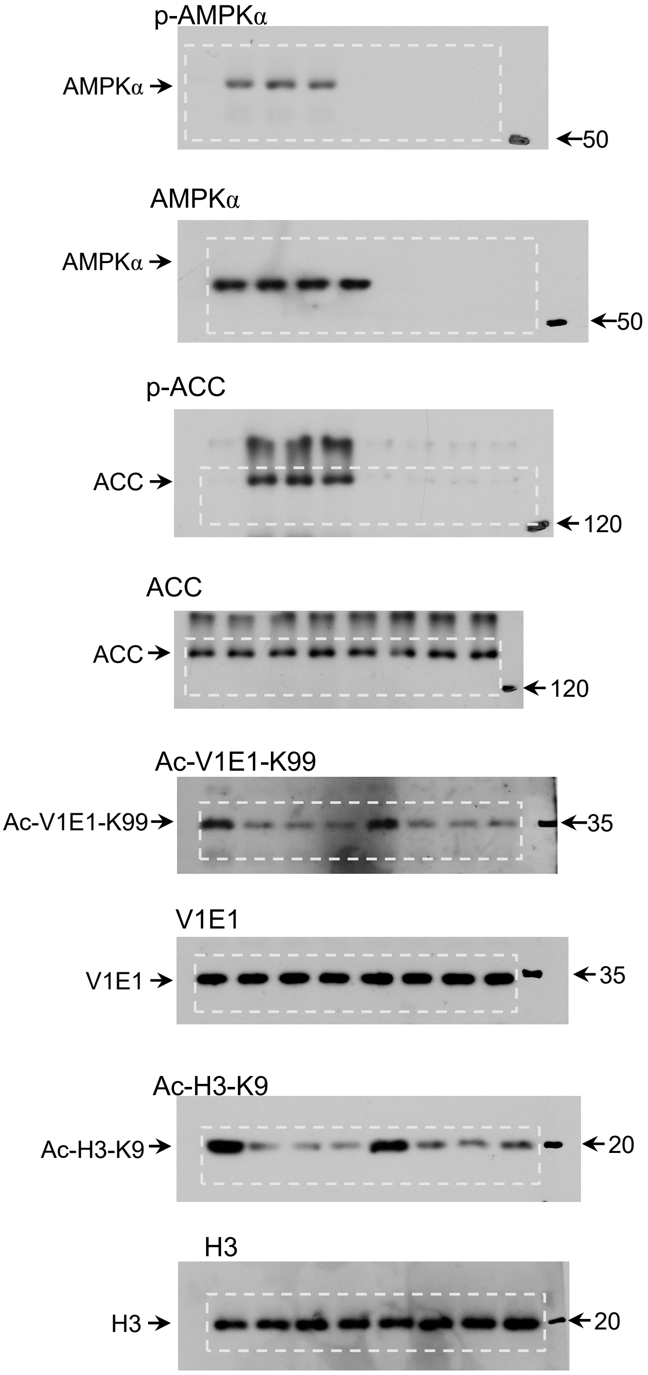

p-AMPKα
AMPKα
50
AMPKα
AMPKα
50
p-ACC
ACC
120
ACC
ACC
120
Ac-V1E1-K99
Ac-V1E1-K99
35
V1E1
V1E1
35
Ac-H3-K9
Ac-H3-K9
20
H3
H3
20

#### Fig. 4b

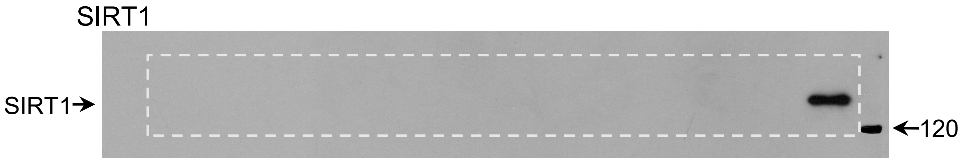


#### Fig. 4c

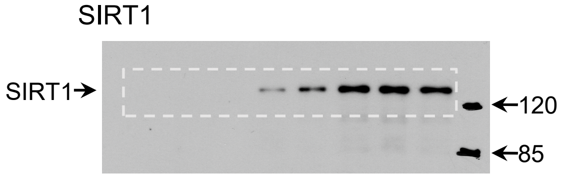


#### Fig. 4d

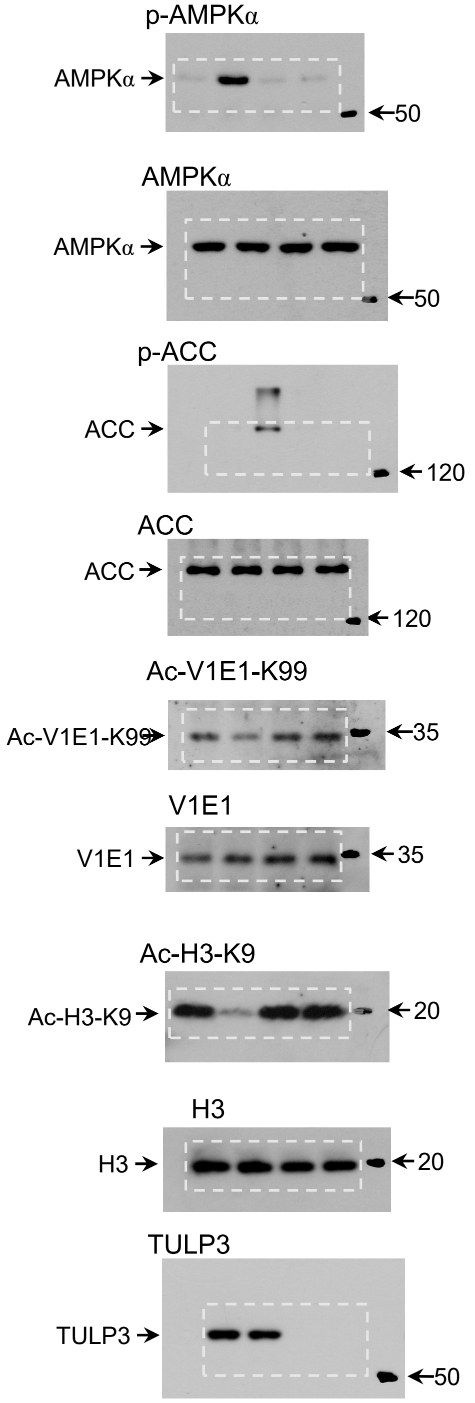

#### Fig. 4f (left)

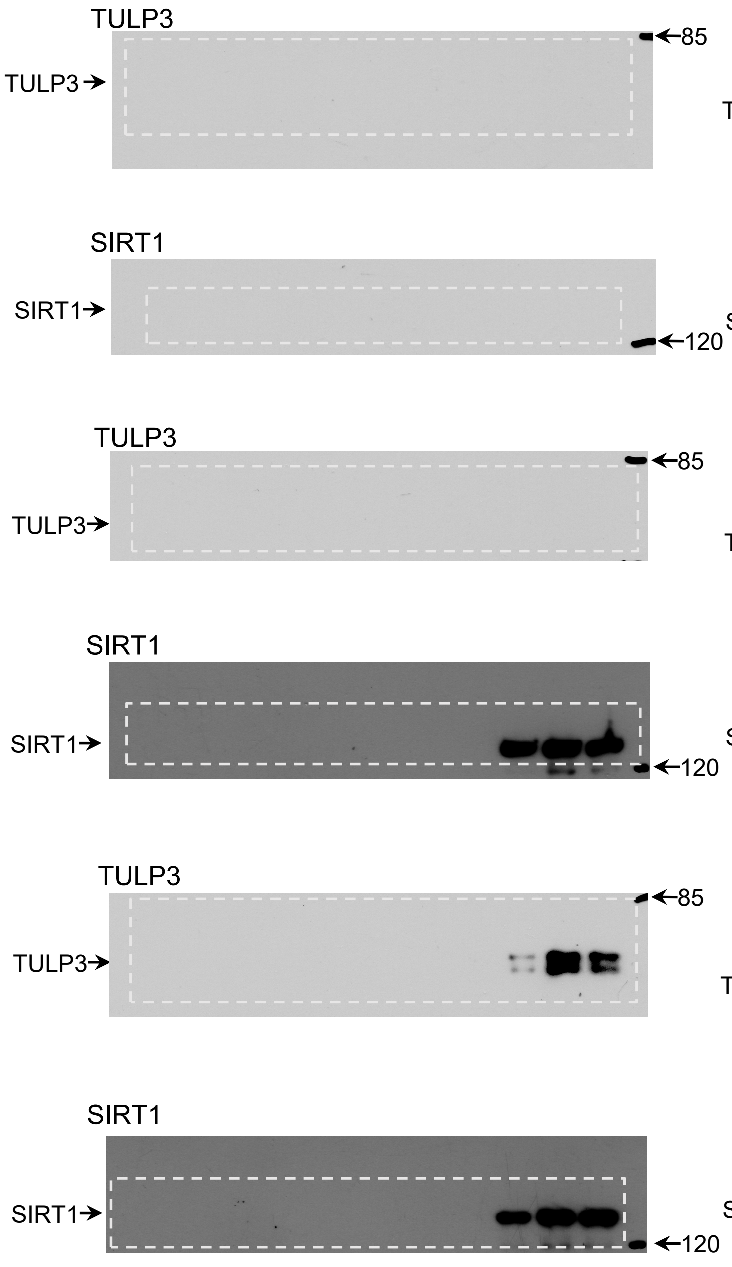


#### Fig. 4f (right)

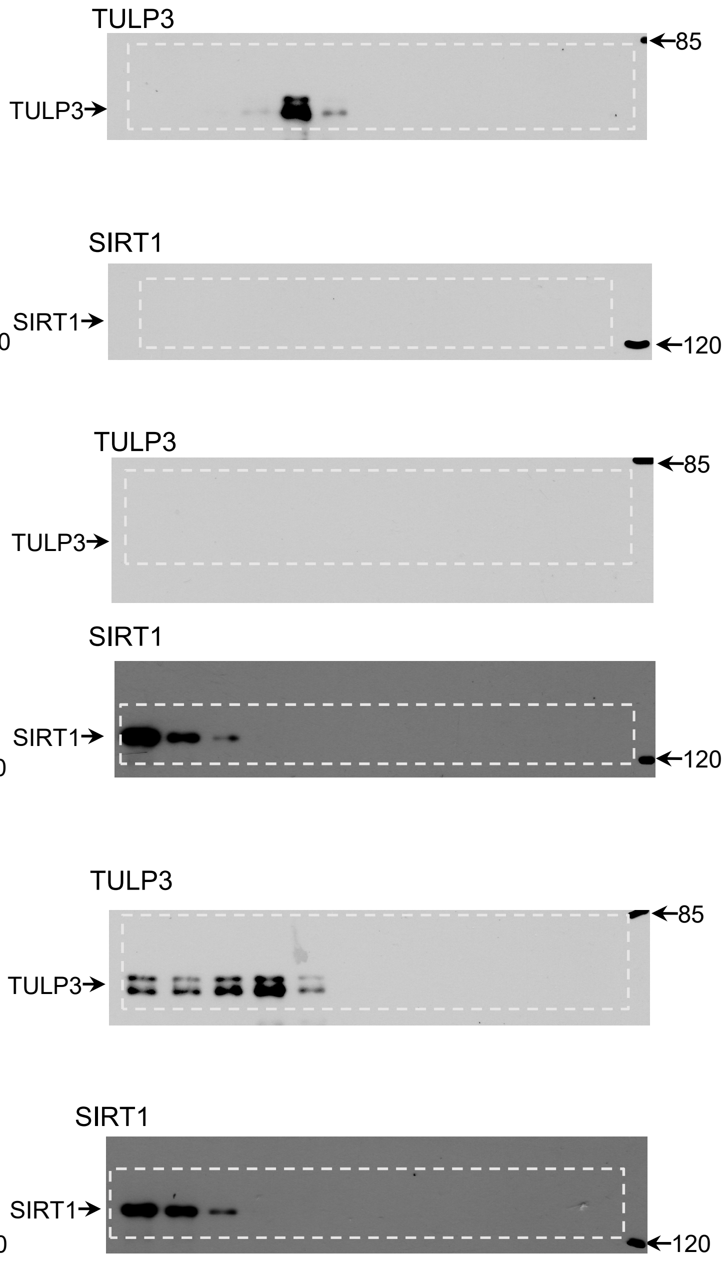

**Fig. 4h**

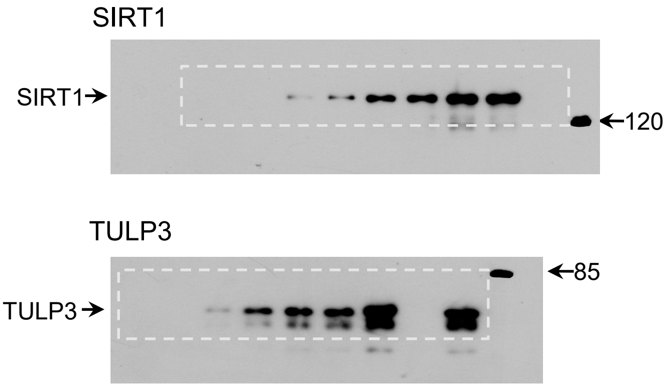


**Fig. 4i**

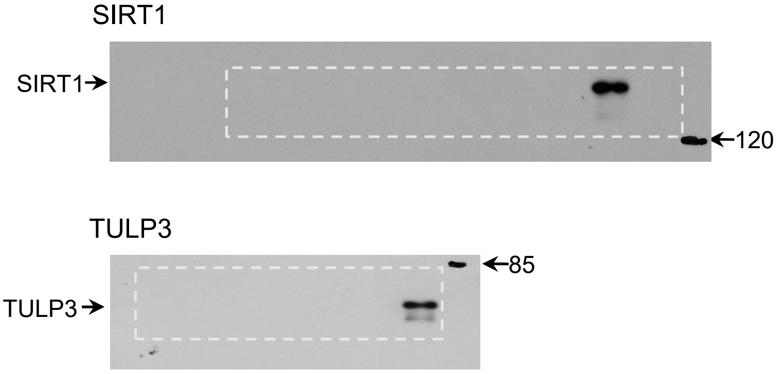


**Fig. 4j**

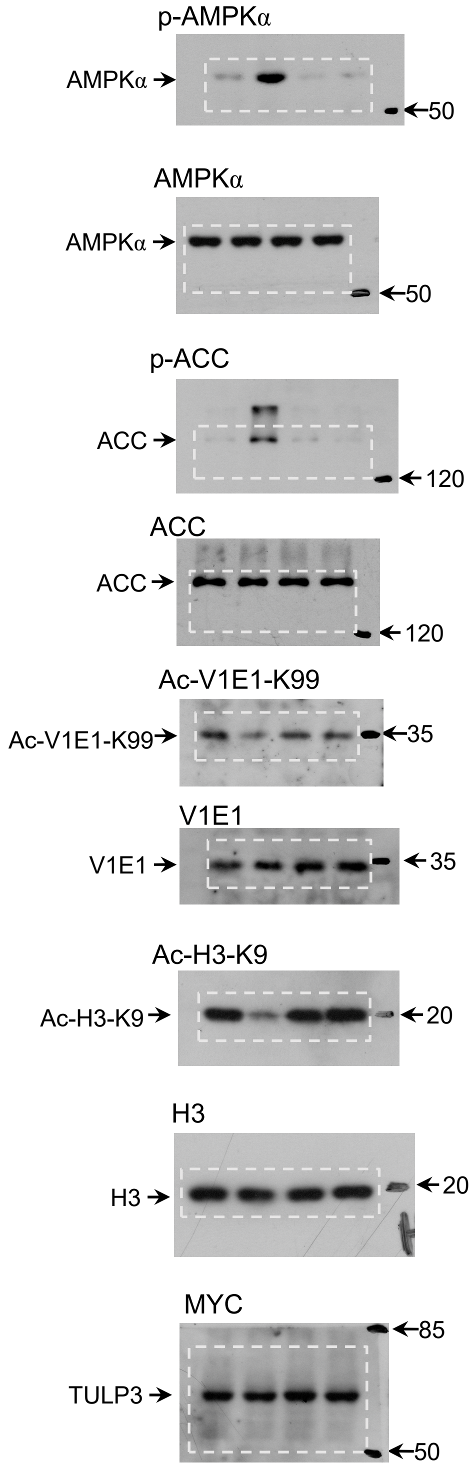

#### Extended Data Fig. 1a

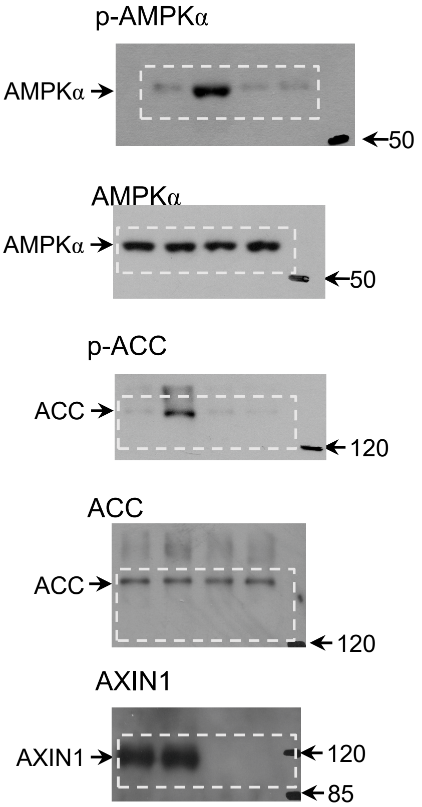


#### Extended Data Fig. 1d

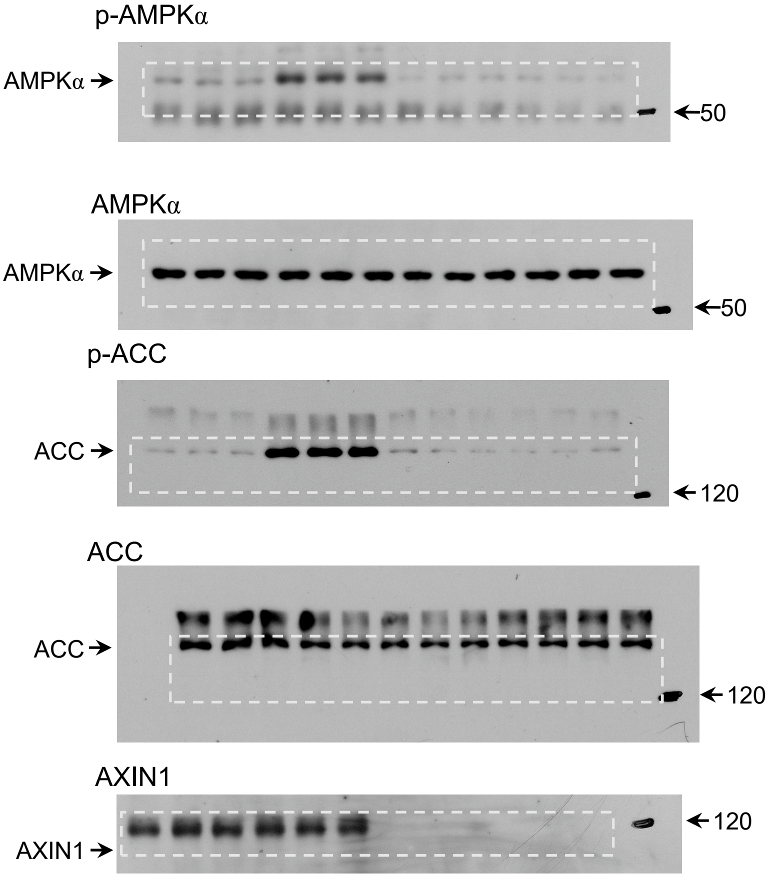

#### Extended Data Fig. 1e

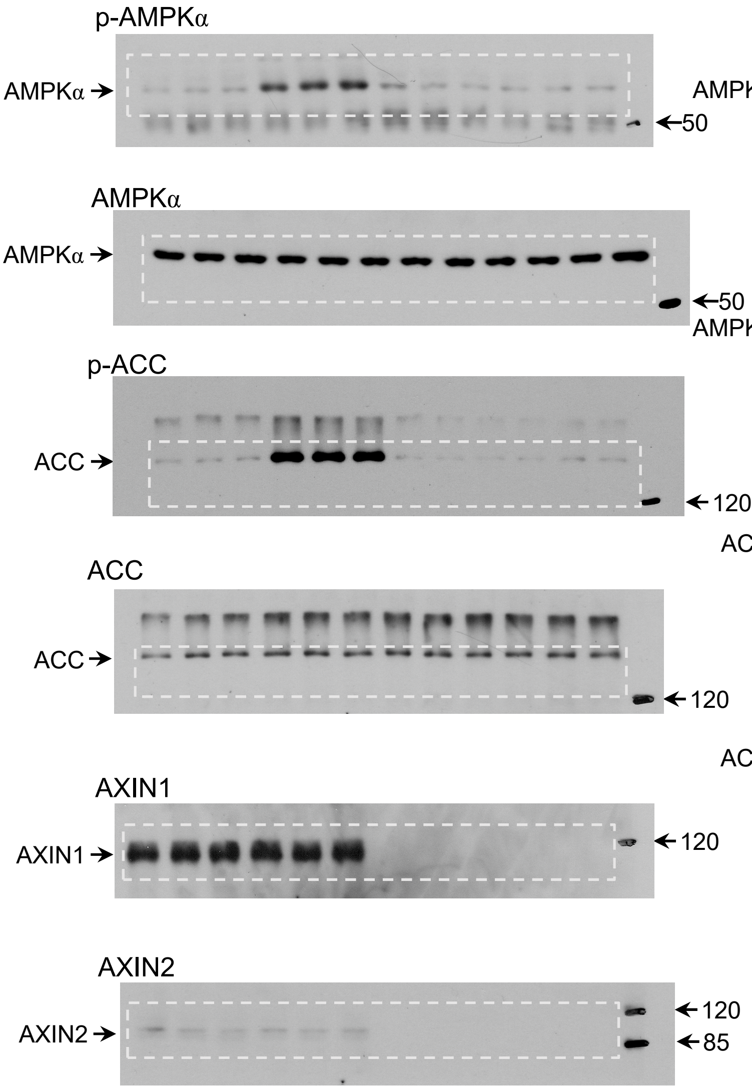


#### Extended Data Fig. 1f

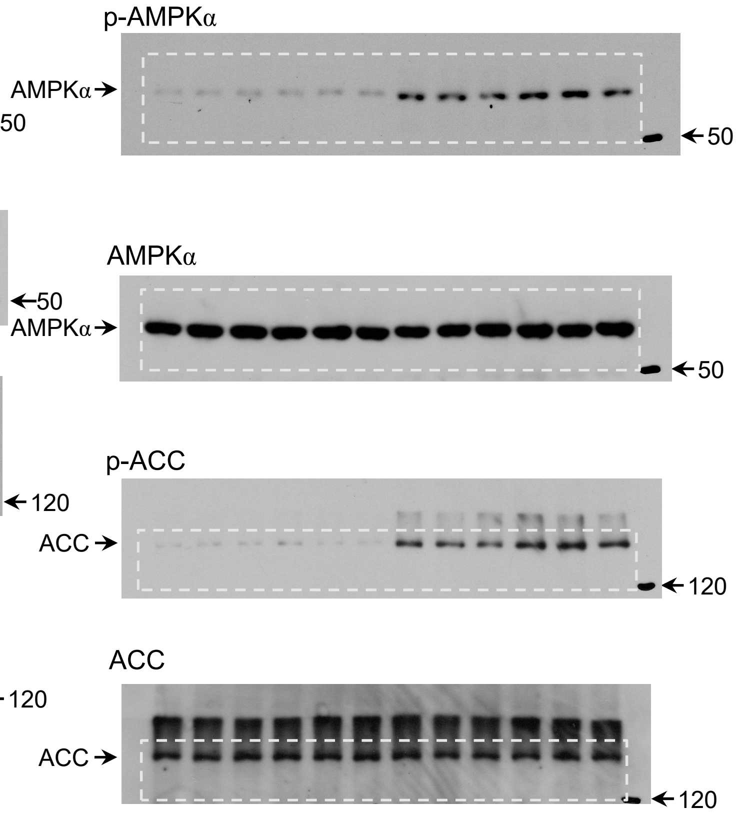

#### Extended Data Fig. 2a

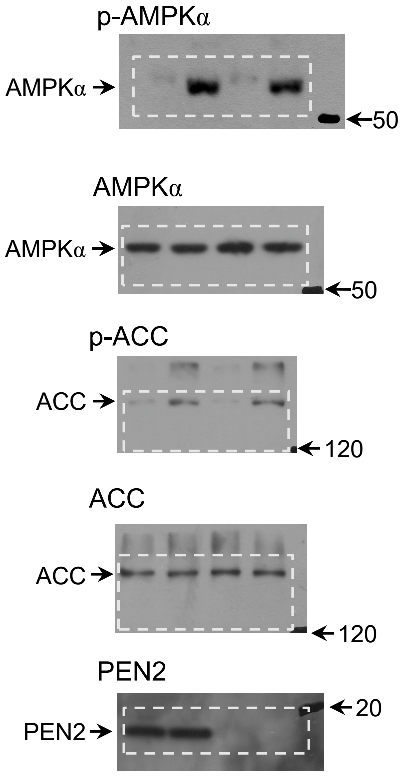


#### Extended Data Fig. 2d

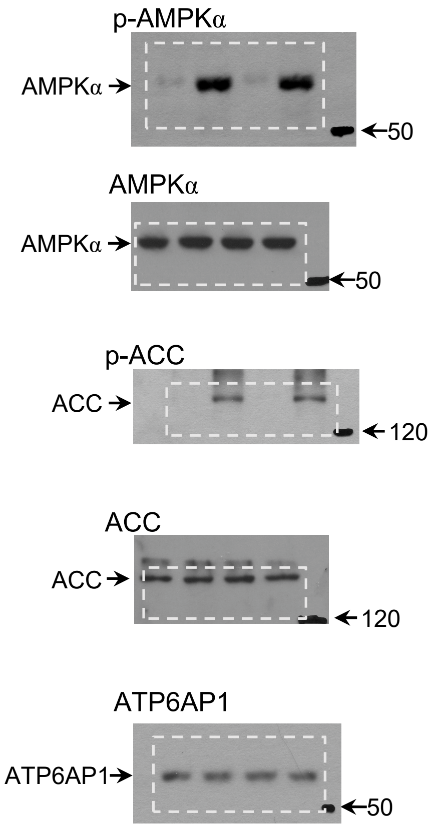

#### Extended Data Fig. 3a

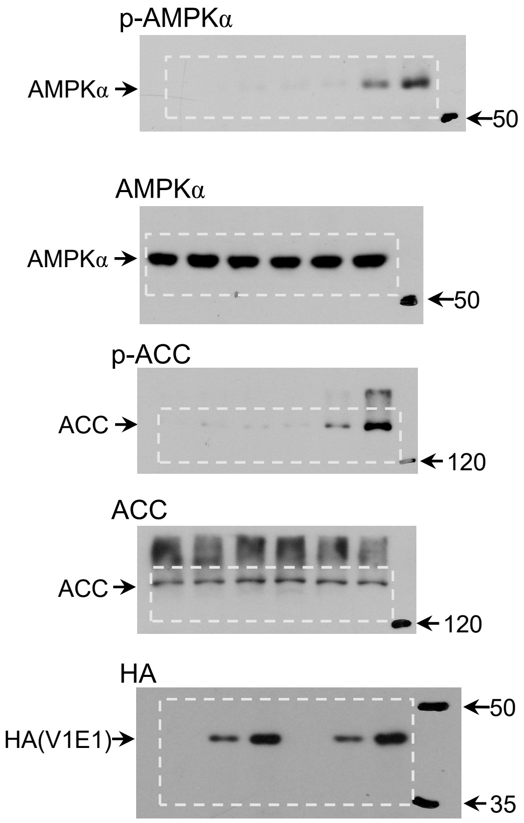


#### Extended Data Fig. 3c

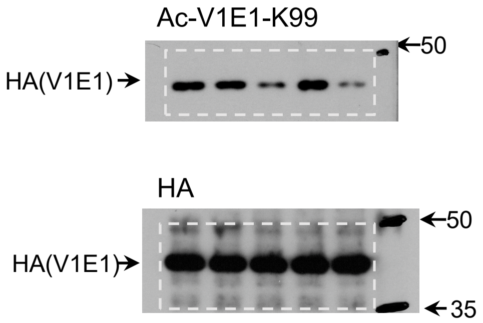

#### Extended Data Fig. 4a

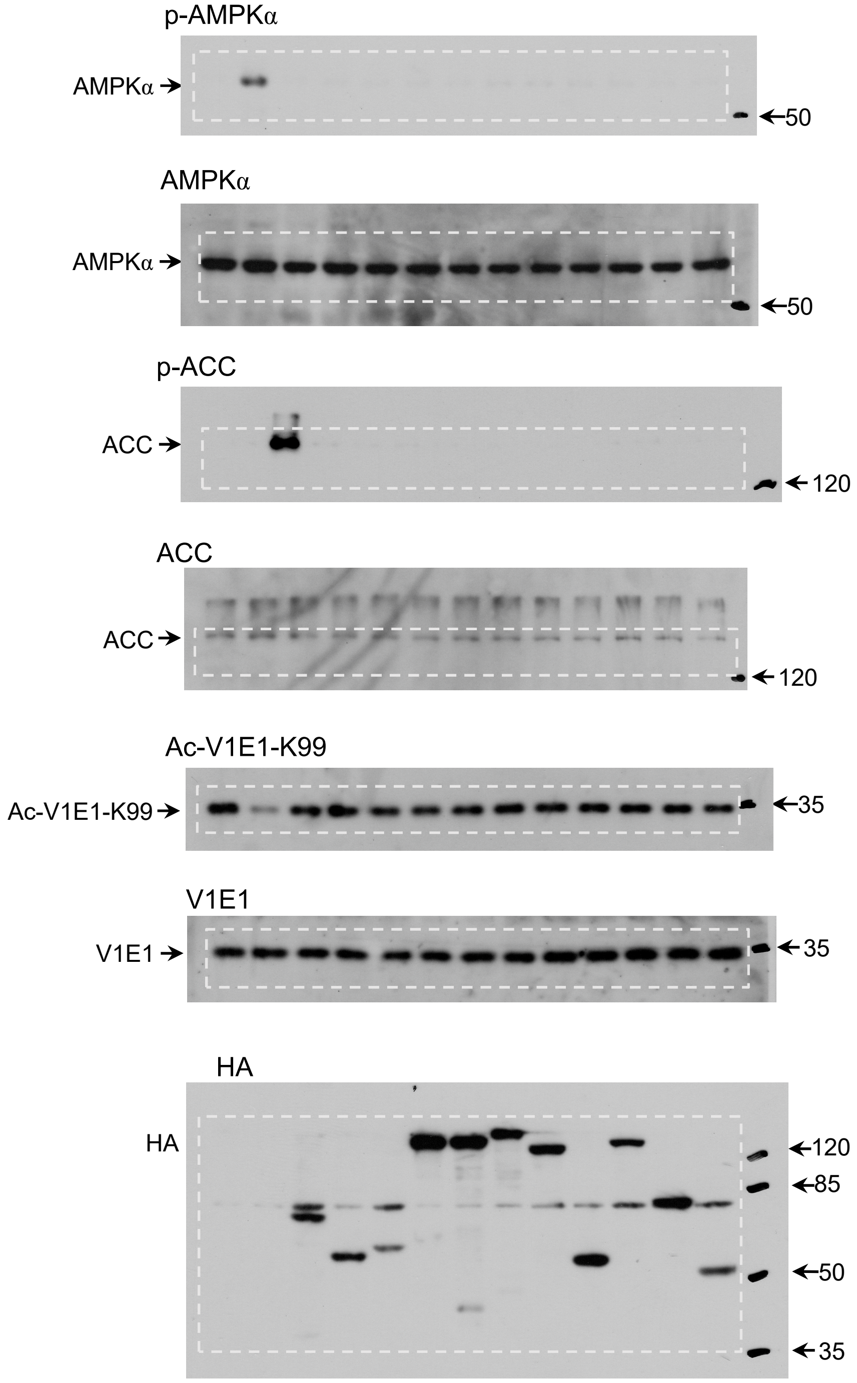

#### Extended Data Fig. 4c

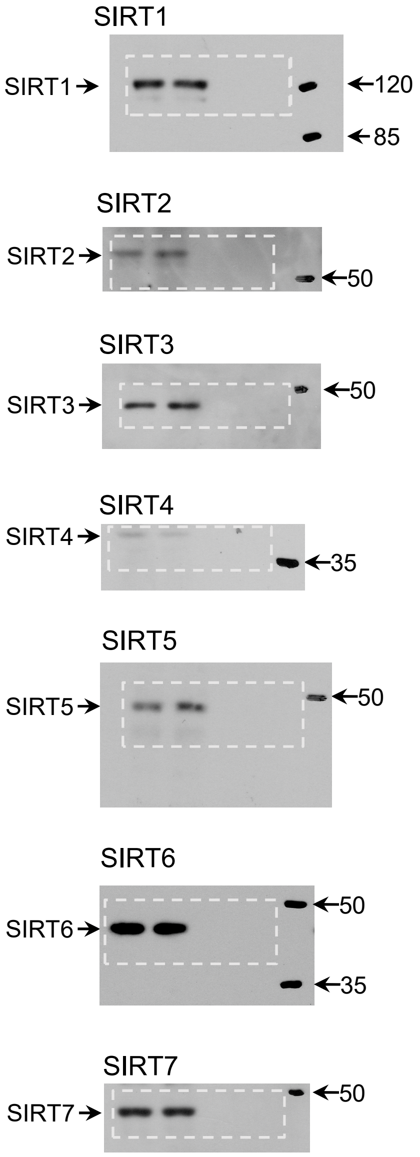


#### Extended Data Fig. 4g

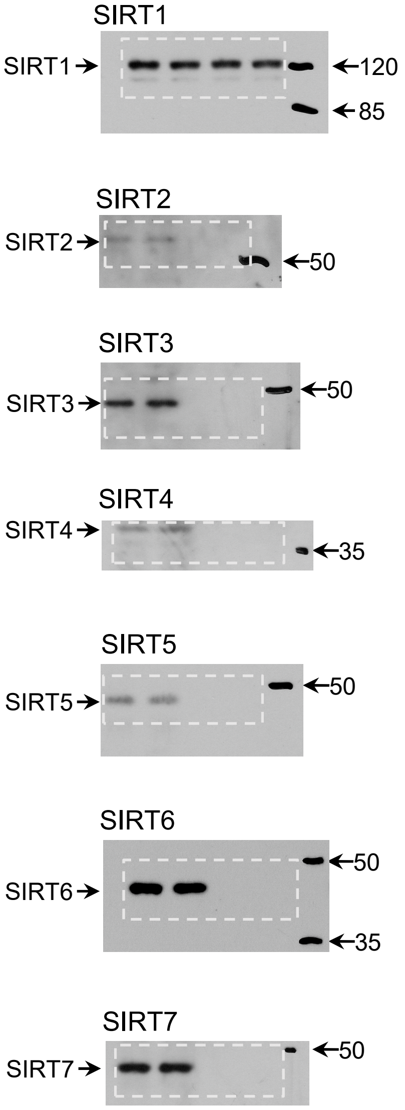

### Extended Data Fig. 6a

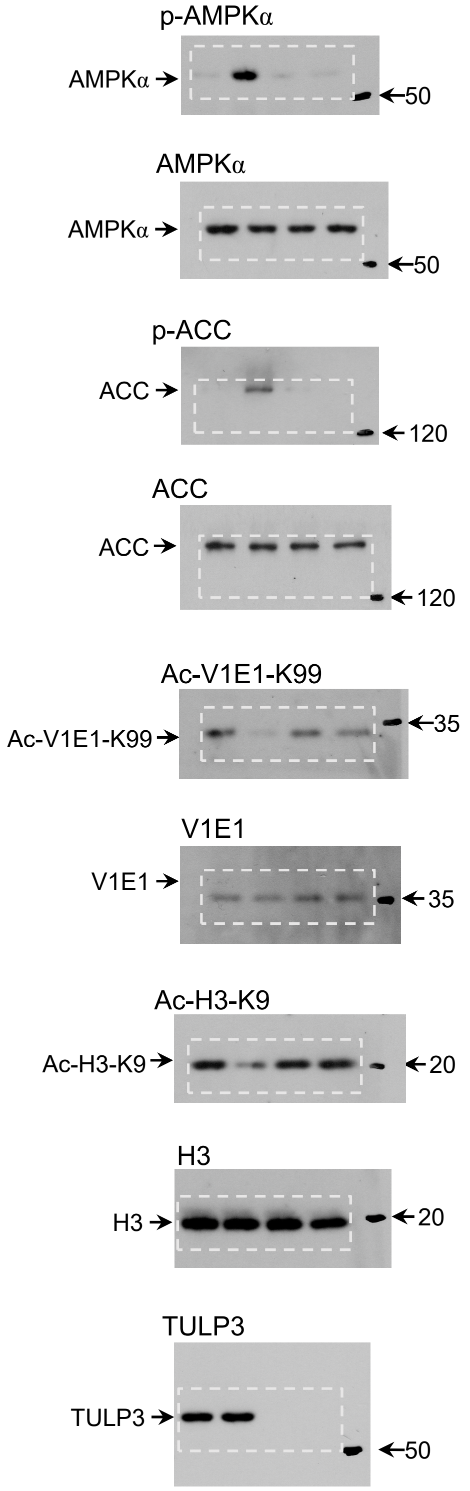

#### Extended Data Fig. 6d

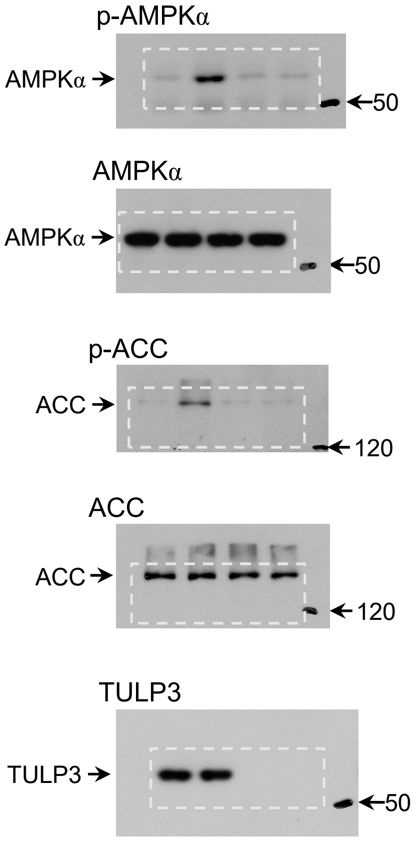

#### Extended Data Fig. 6e (left)

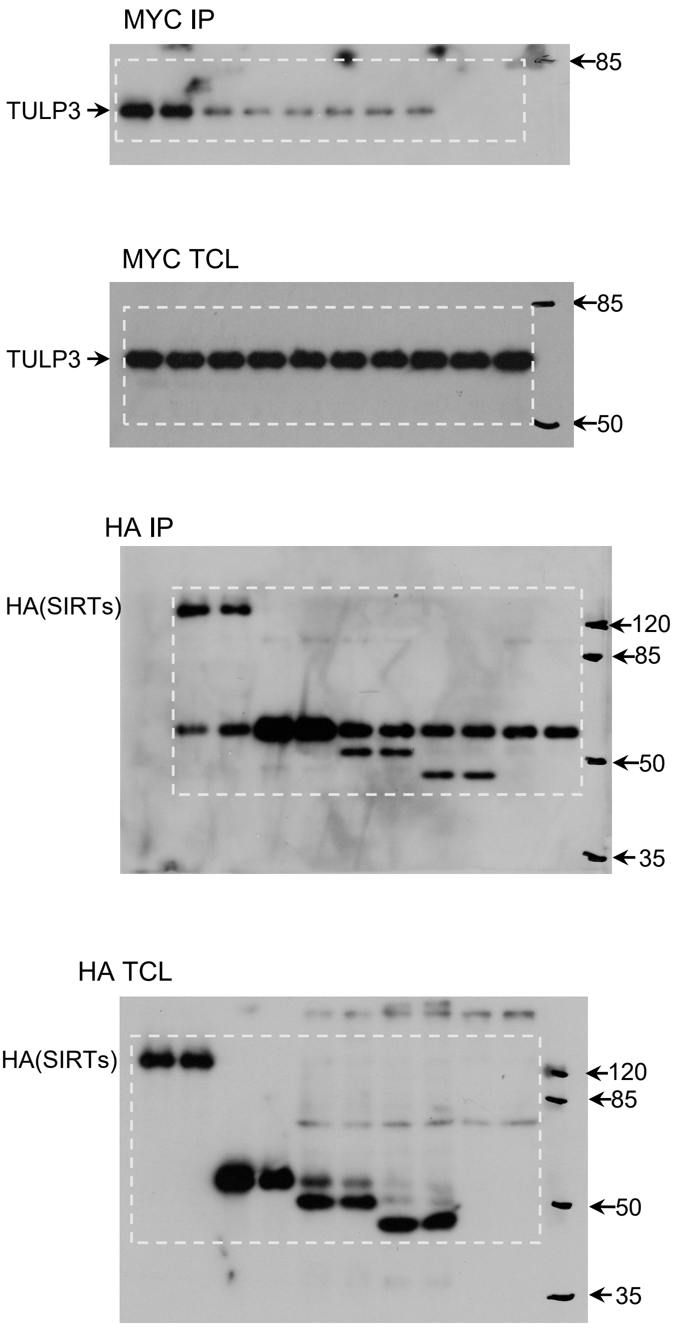


#### Extended Data Fig. 6e (right)

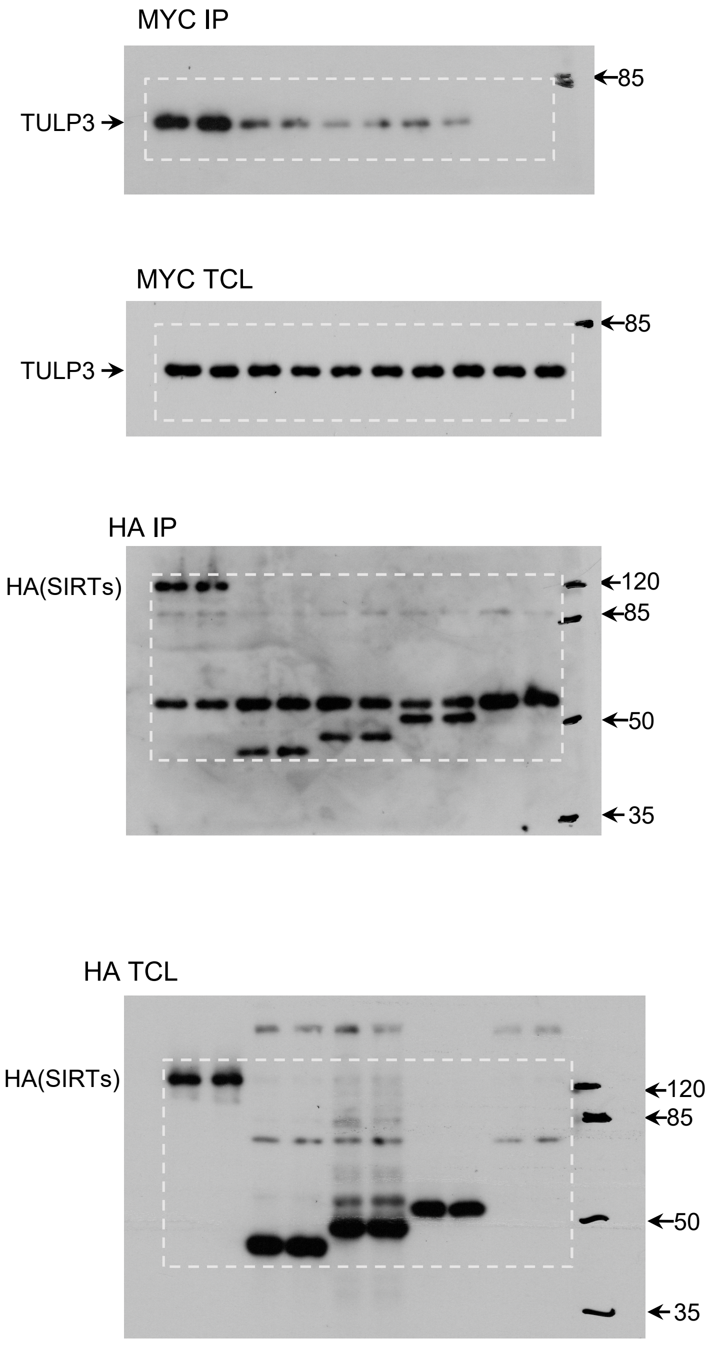

#### Extended Data Fig. 7a (left)

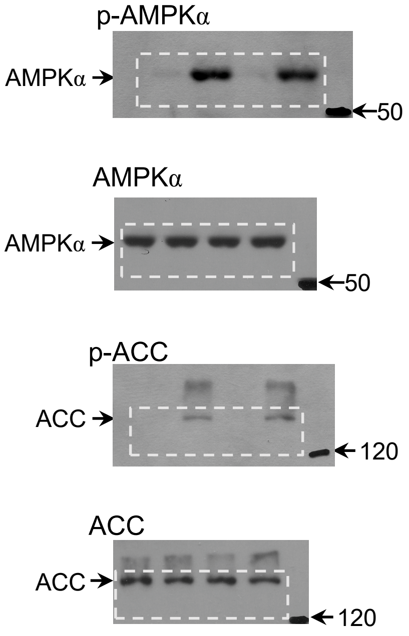


#### Extended Data Fig. 7a (right)

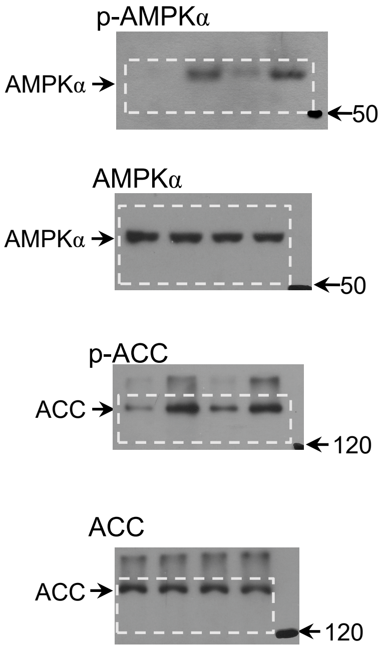


#### Extended Data Fig. 7b (left)

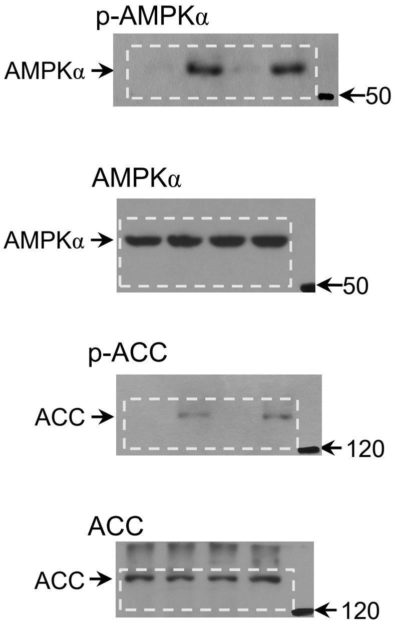


#### Extended Data Fig. 7b (right)

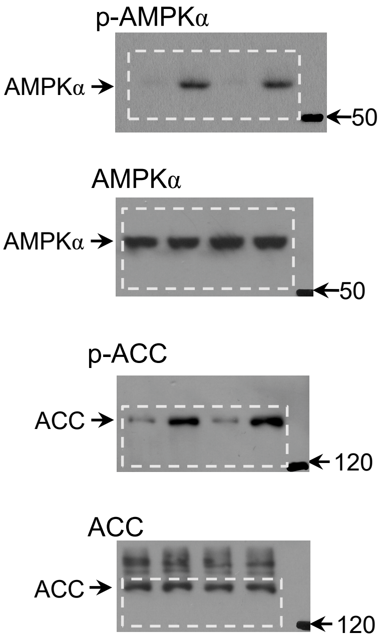

#### Extended Data Fig. 7c (left)

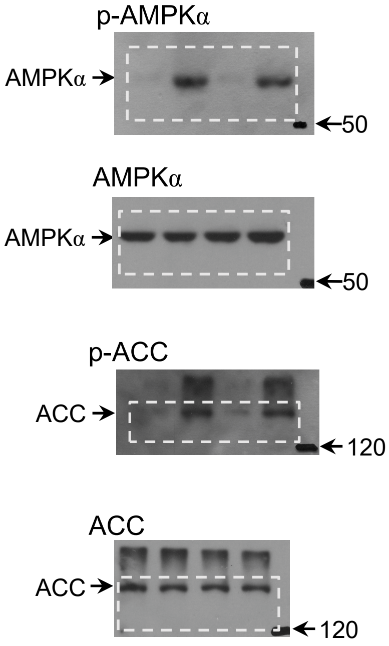


#### Extended Data Fig. 7c (right)

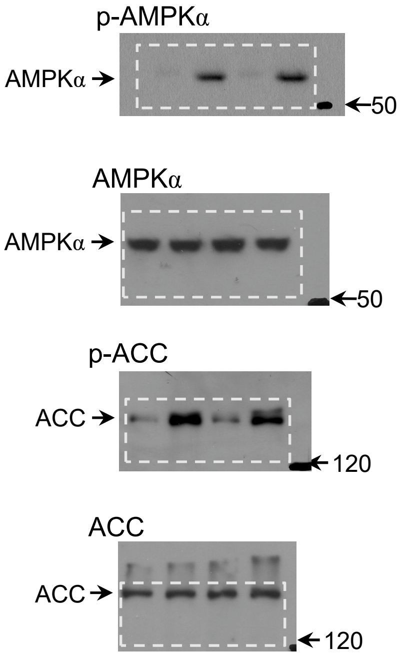


#### Extended Data Fig. 7d (left)

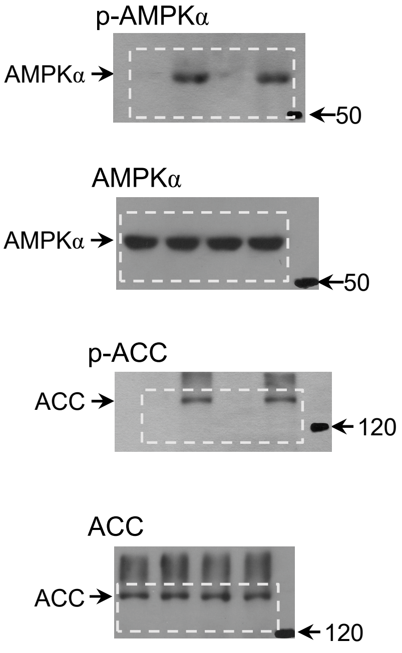


#### Extended Data Fig. 7d (right)

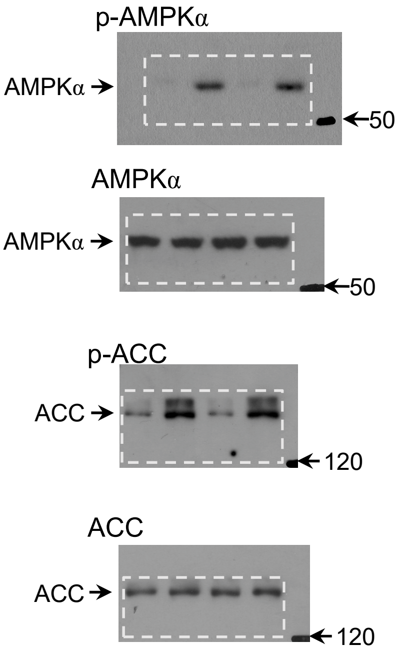

#### Extended Data Fig. 7e (left)

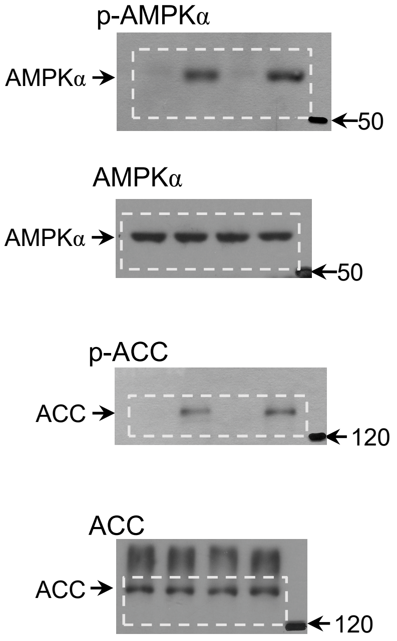


#### Extended Data Fig. 7e (right)

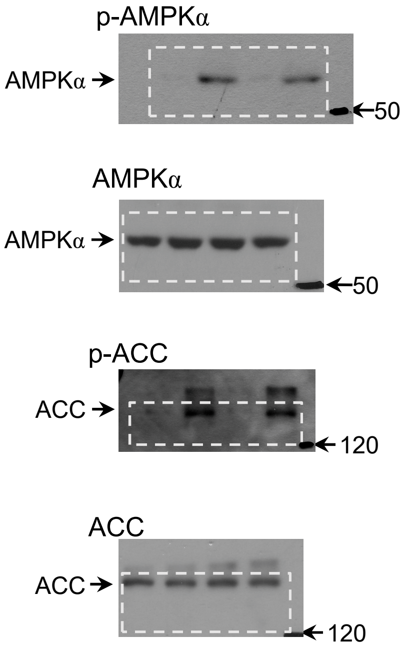


#### Extended Data Fig. 7f (left)

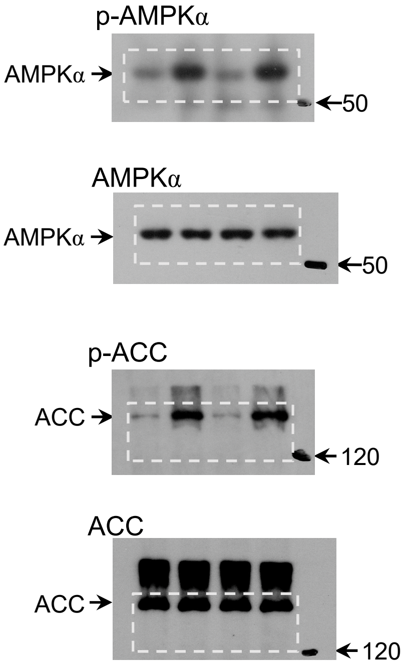


#### Extended Data Fig. 7f (right)

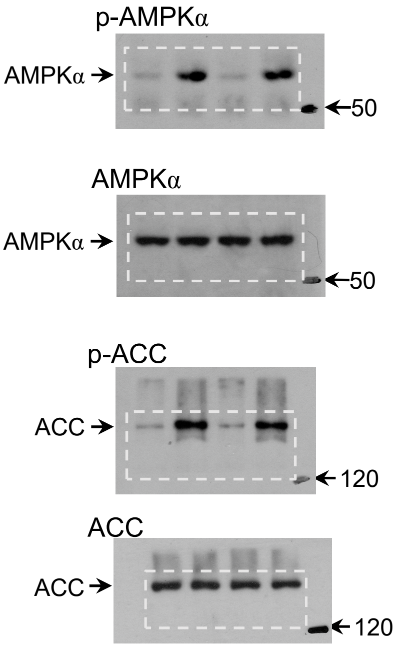

#### Extended Data Fig. 7g (left)

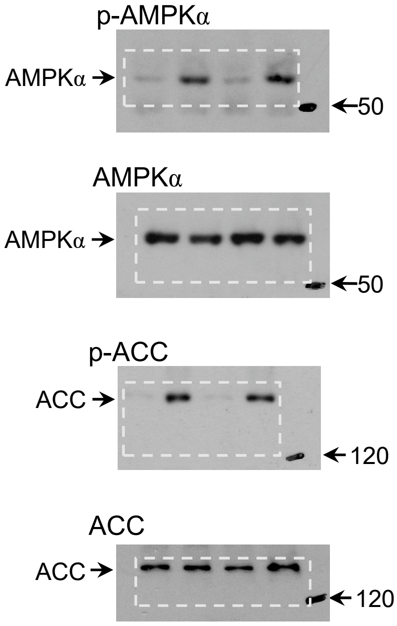


#### Extended Data Fig. 7g (right)

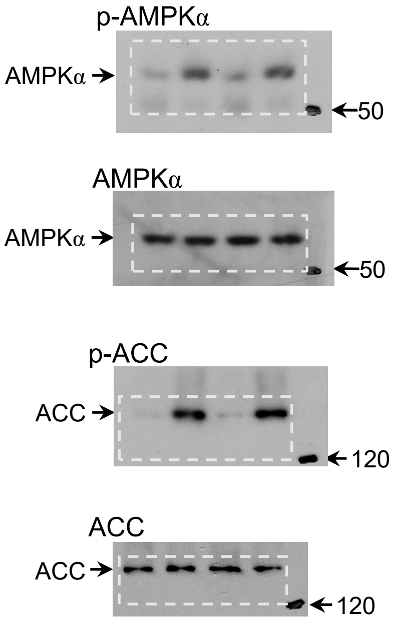


#### Extended Data Fig. 7h (left)

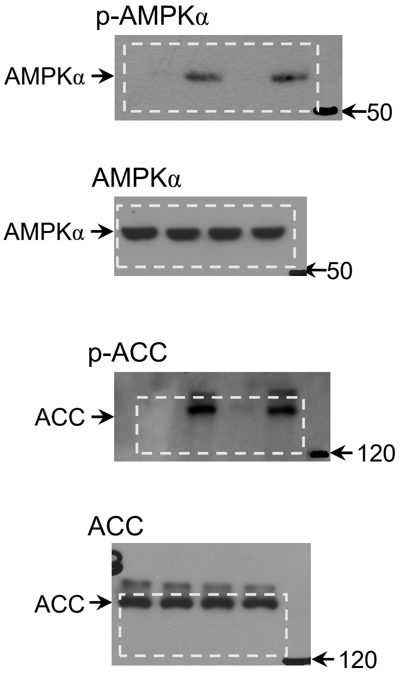


#### Extended Data Fig. 7h (right)

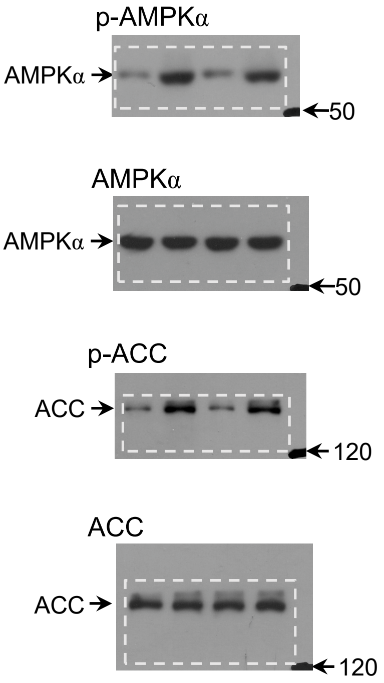

#### Extended Data Fig. 7i (left)

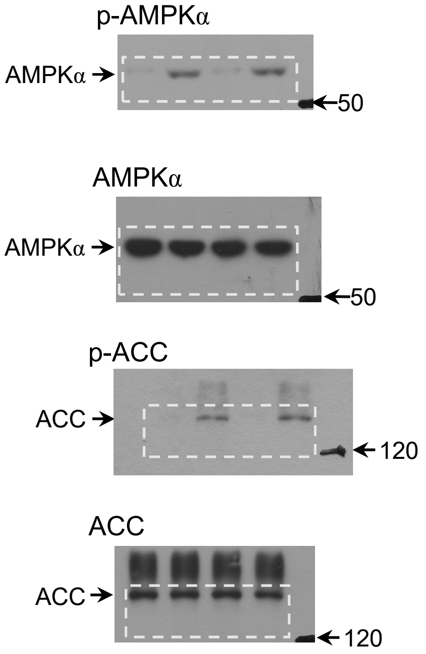


#### Extended Data Fig. 7i (right)

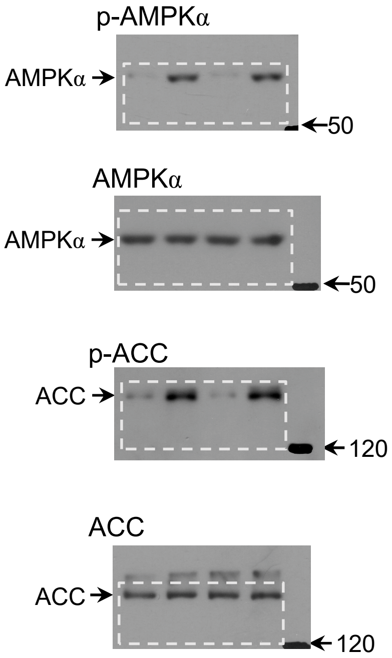


#### Extended Data Fig. 7j (left)

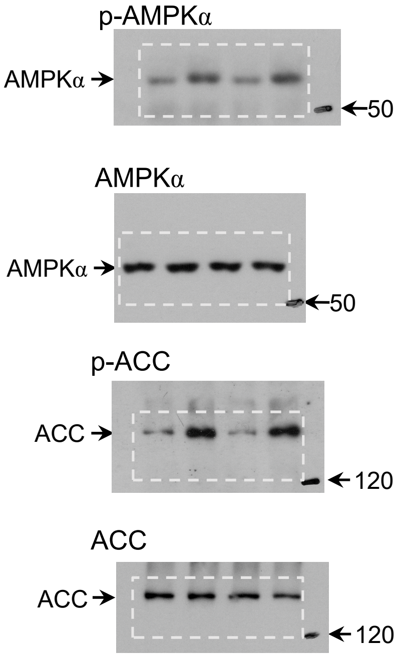


#### Extended Data Fig. 7j (right)

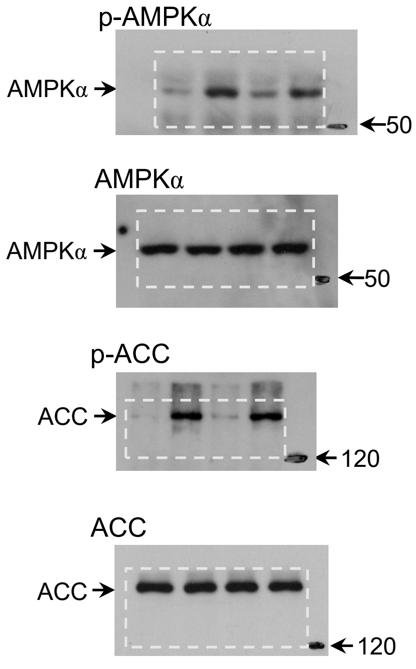

#### Extended Data Fig. 7k (left)

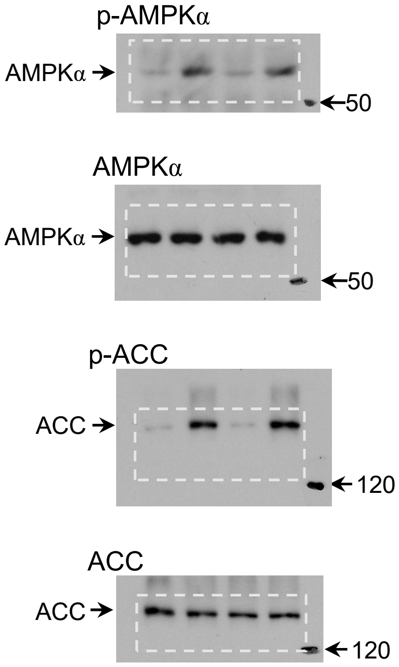


#### Extended Data Fig. 7k (right)

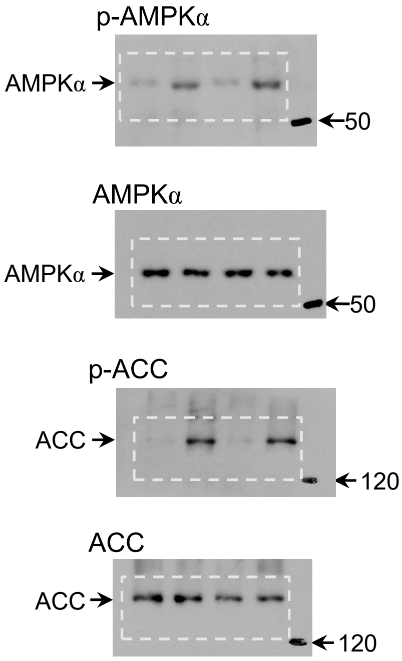


#### Extended Data Fig. 7l (left)

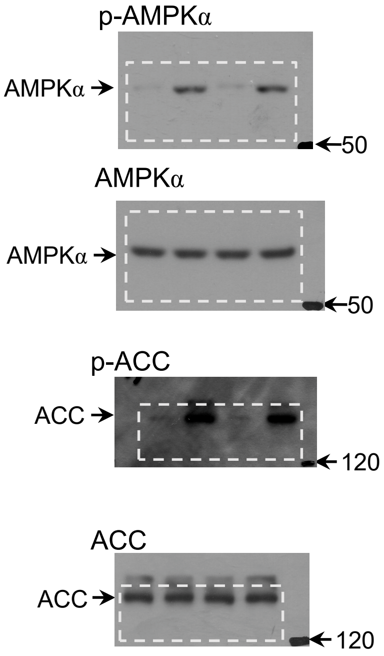


#### Extended Data Fig. 7l (right)

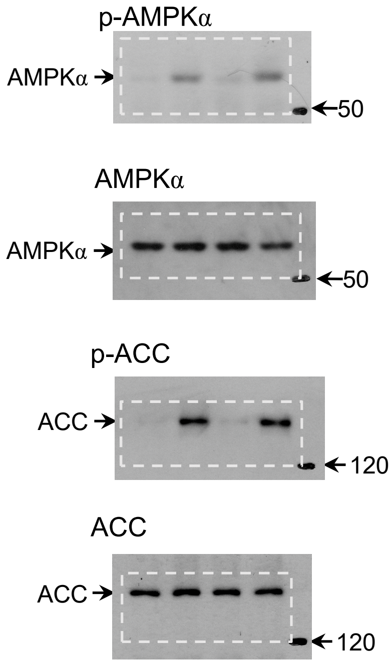

#### Extended Data Fig. 7m (left)

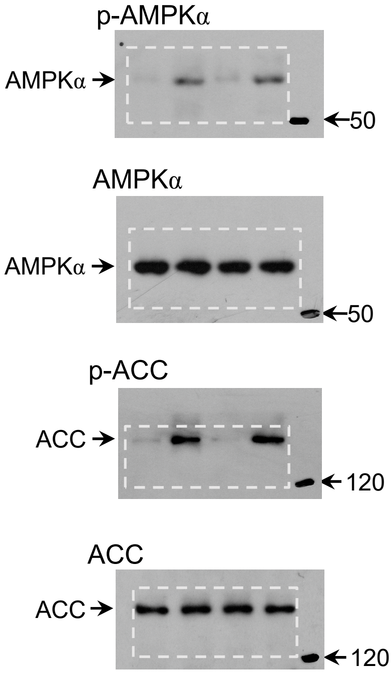


#### Extended Data Fig. 7m (right)

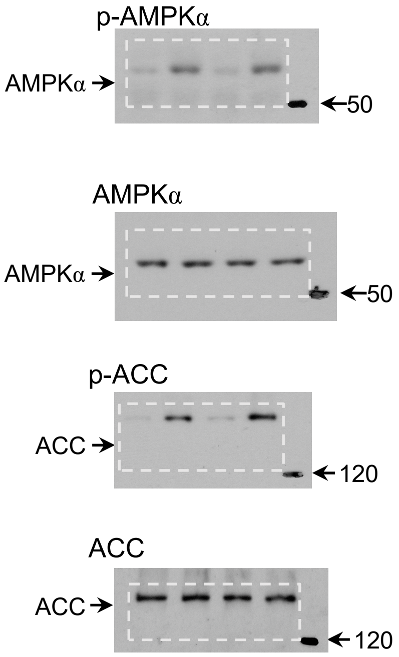


#### Extended Data Fig. 7n (left)

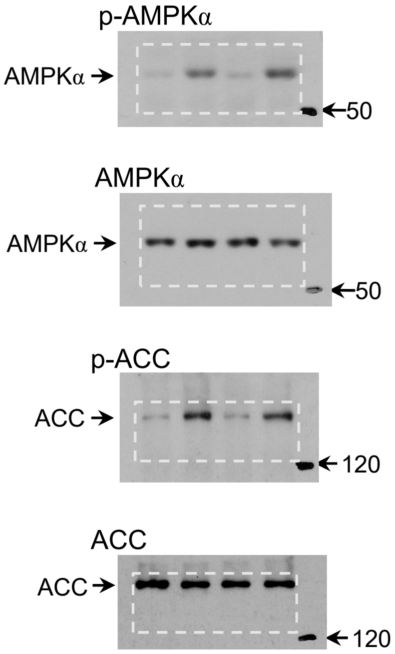


#### Extended Data Fig. 7n (right)

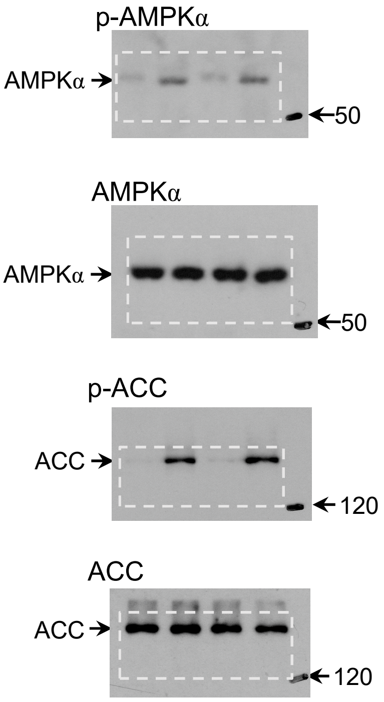

#### Extended Data Fig. 7o (left)

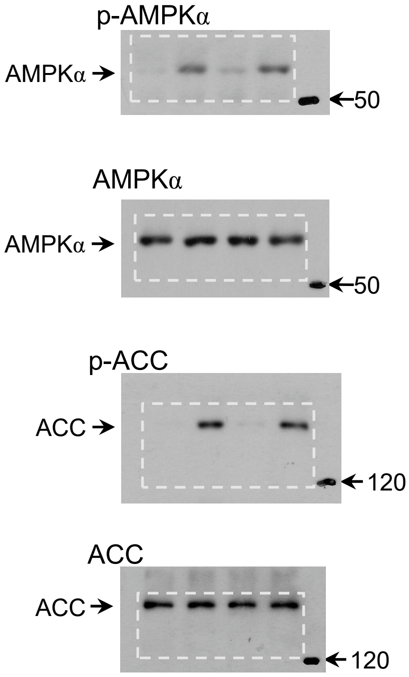


#### Extended Data Fig. 7o (right)

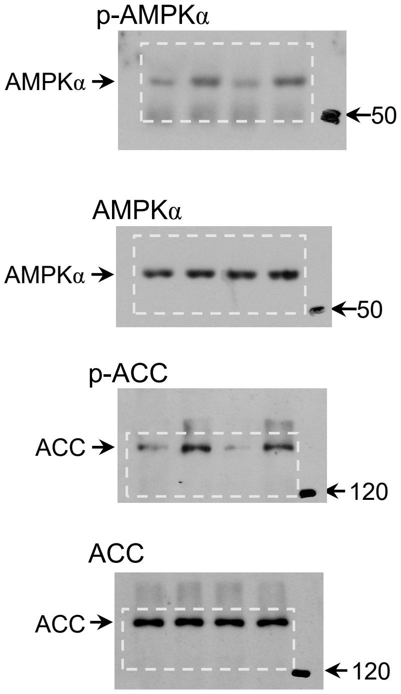


#### Extended Data Fig. 7p (left)

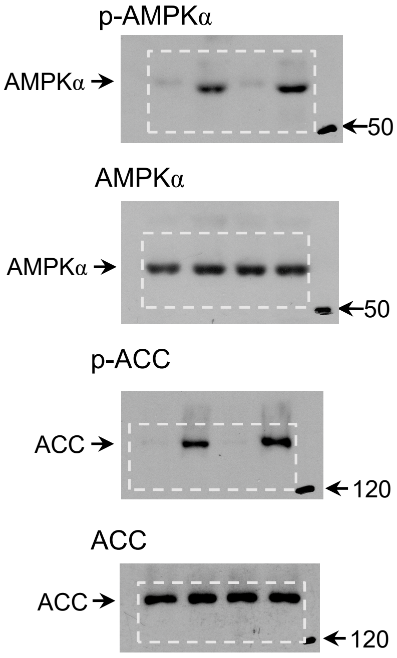


#### Extended Data Fig. 7p (right)

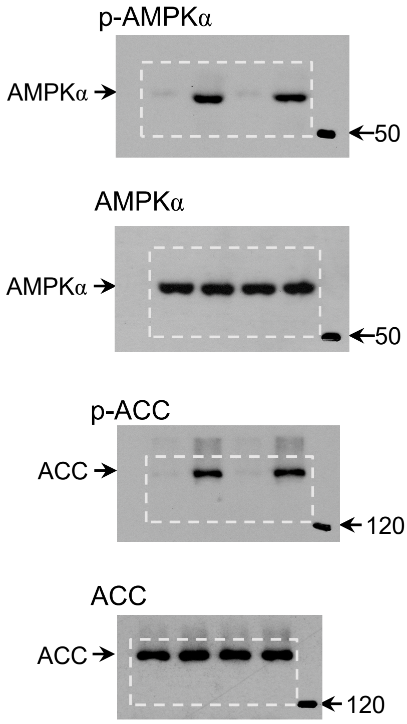

#### Extended Data Fig. 7q (left)

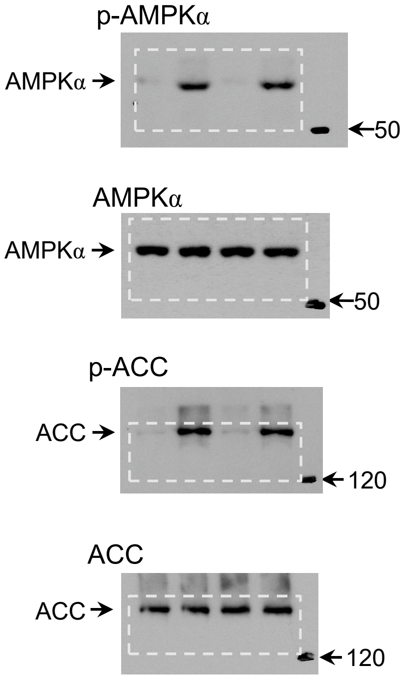


#### Extended Data Fig. 7q (right)

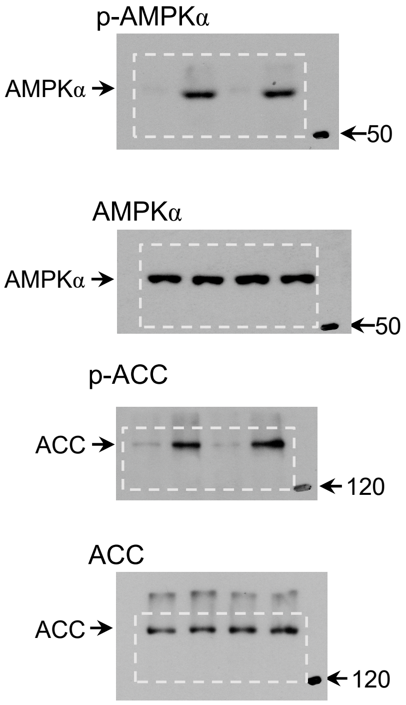


#### Extended Data Fig. 7r (upper)

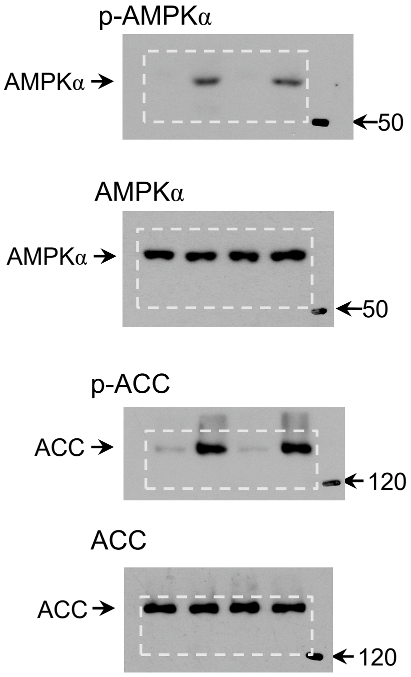


#### Extended Data Fig. 7r (lower)

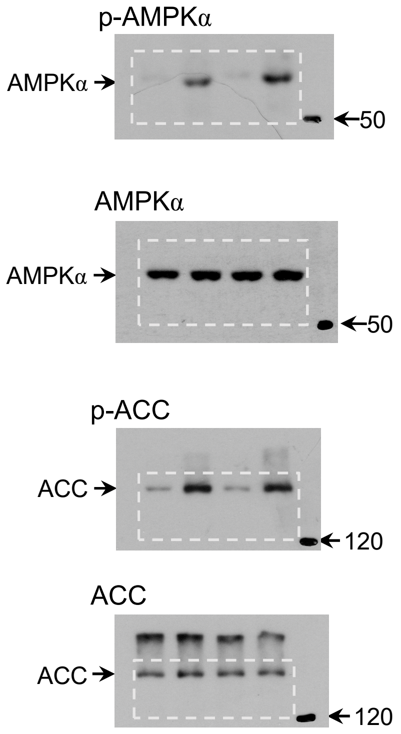

#### Extended Data Fig. 7s (left)

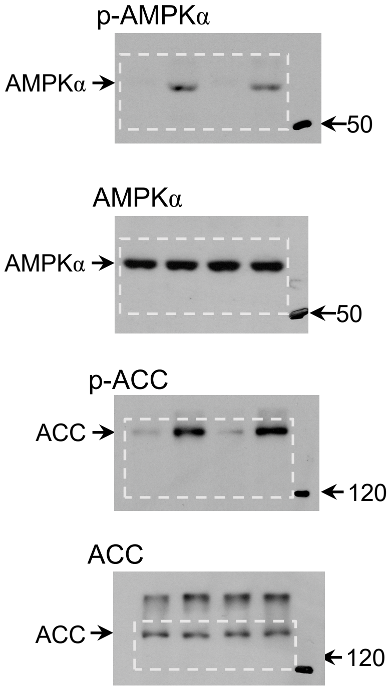


#### Extended Data Fig. 7s (right)

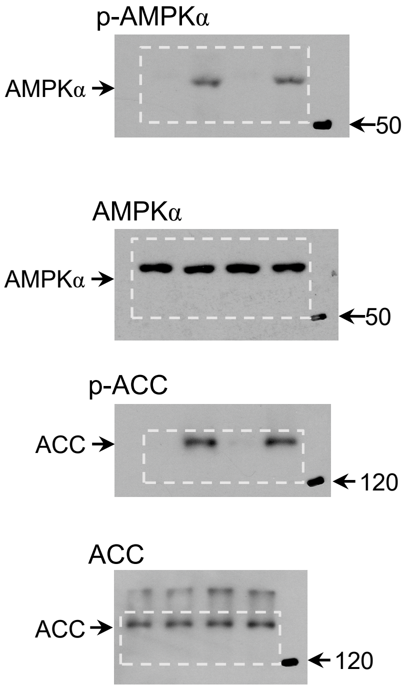

### Extended Data Fig. 8a

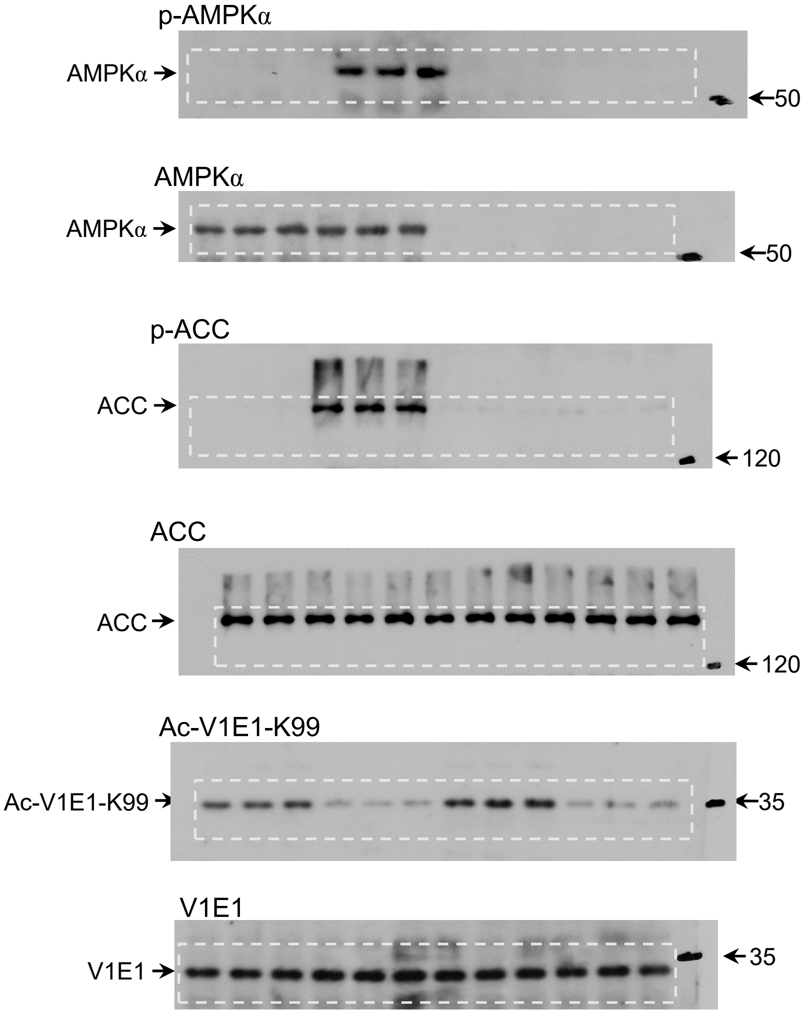


### Extended Data Fig. 8b

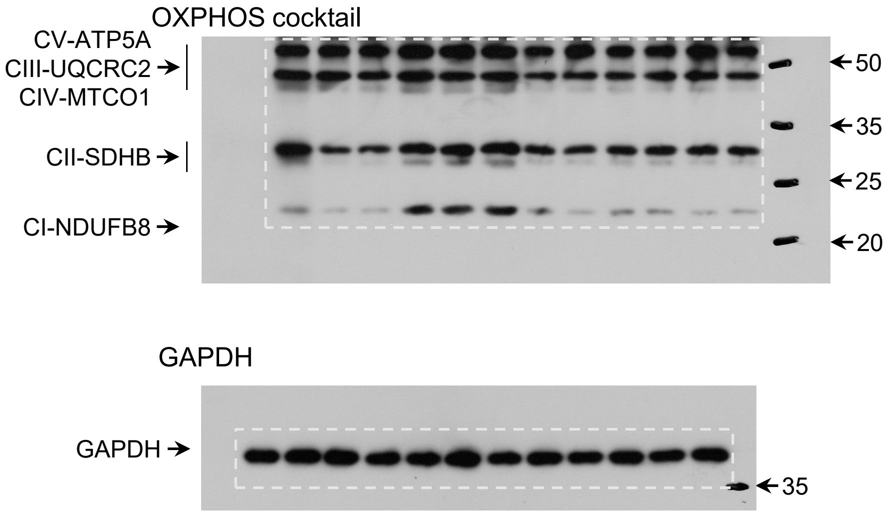

#### Extended Data Fig. 8e (upper)

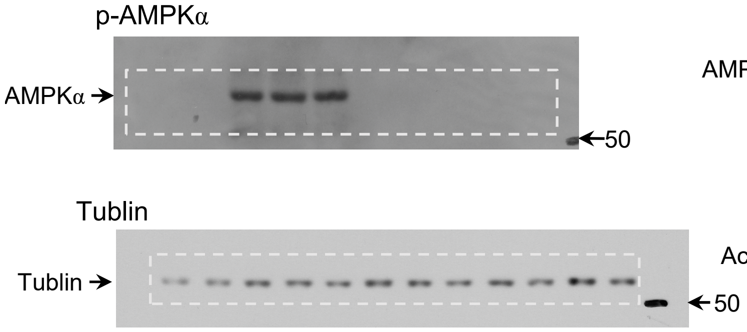


#### Extended Data Fig. 8e (lower)

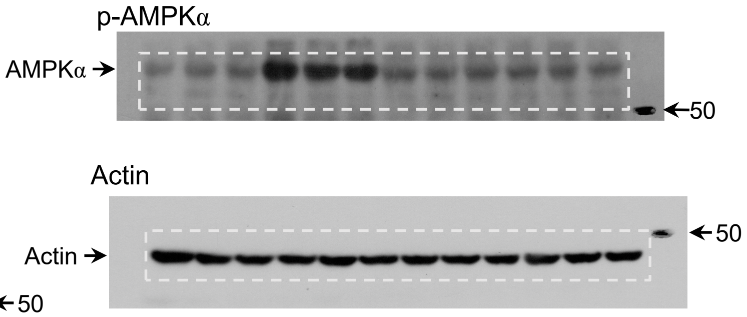


#### Extended Data Fig. 8g (upper)

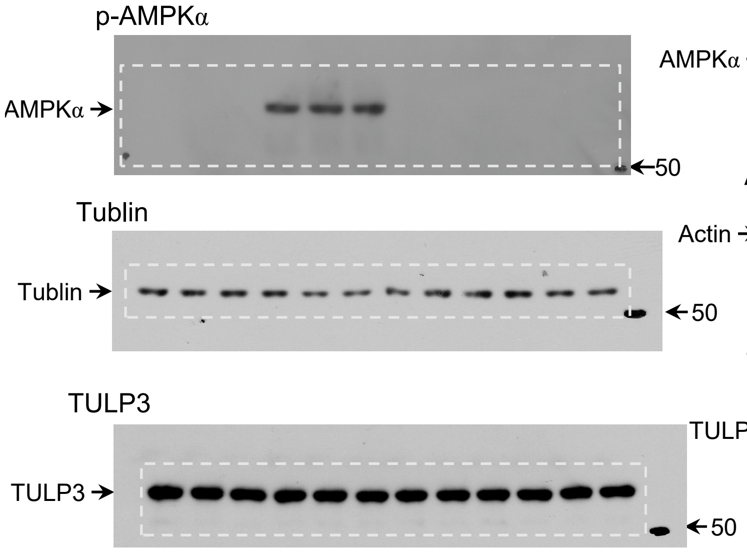


#### Extended Data Fig. 8g (lower)

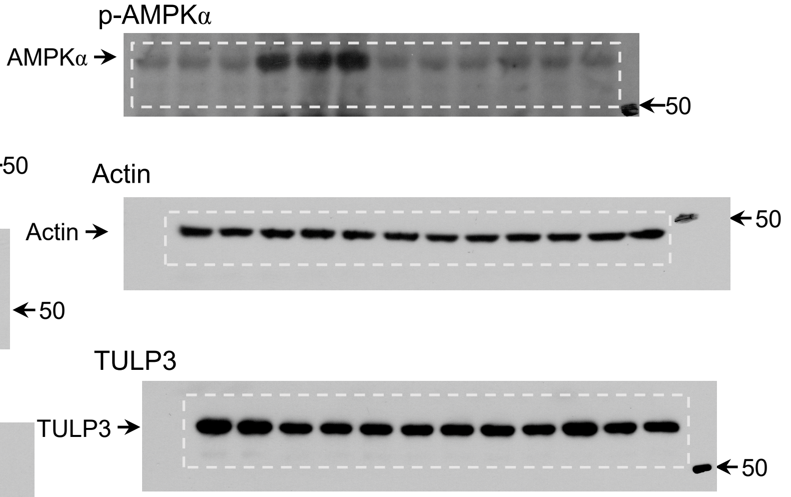

#### Extended Data Fig. 8h (upper)

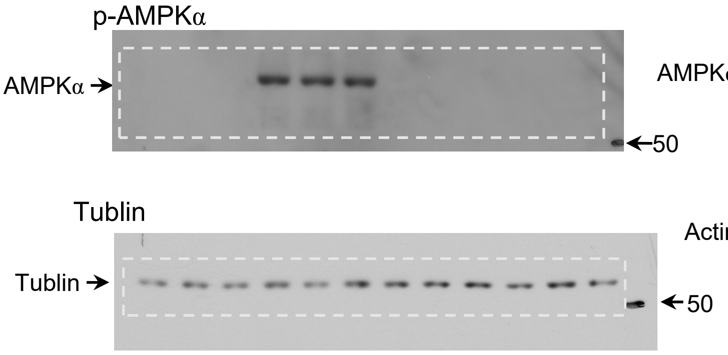


#### Extended Data Fig. 8h (lower)

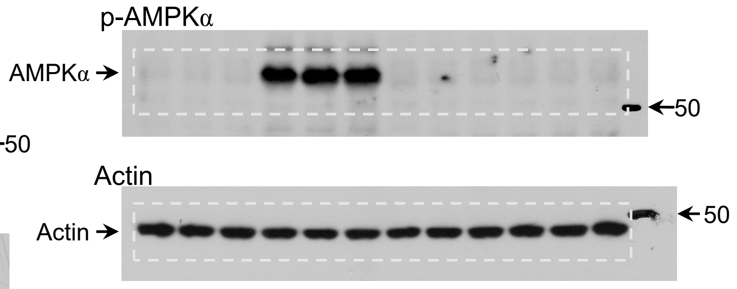


#### Extended Data Fig. 8i (upper)

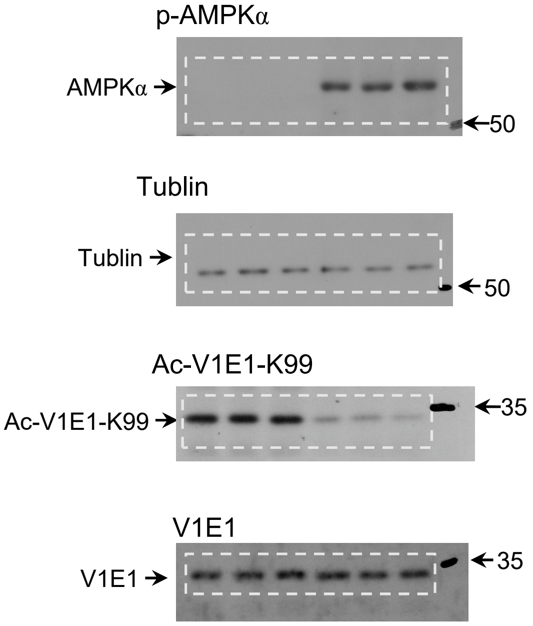


#### Extended Data Fig. 8i (lower)

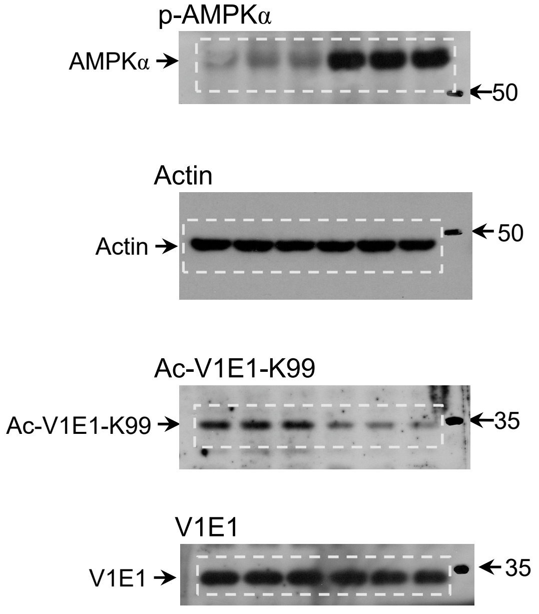
